# Tipping the balance: Synthesis and evaluation of centrinone-based degraders of polo-like kinase 4

**DOI:** 10.64898/2026.01.31.703010

**Authors:** Andrej Kovacevic, Aleksandar Salim, Crisálida Borges, Patrick Meraldi, Sascha Hoogendoorn

## Abstract

Polo-like kinase 4 (PLK4) is a serine/threonine-protein kinase that plays a pivotal role in centriole biogenesis and, as such, represents a master regulator of centriole duplication. Due to its importance in cancer development and progression, PLK4 represents an attractive target for the development of novel therapeutics. Herein, we present a series of molecular degraders of PLK4, based on the highly selective PLK4 inhibitor centrinone, with the aim of targeting PLK4 for degradation via the ubiquitin-proteasome system. While all synthesized degraders retained low nanomolar binding affinities to the kinase domain of PLK4, large differences were found with respect to their ability to change cellular PLK4 levels. We uncover a complex pharmacological profile of the most potent degraders, **D6** and **D10**, consisting of concomitant lowering of PLK4 levels through degradation, and enhancing PLK4 levels through inhibition of its autoregulation – dependent on its localization at the centrioles.

## INTRODUCTION

Centrosomes (Figure 1A) are non-membrane bound organelles comprised of two centrioles, surrounded by the pericentriolar material (PCM).^1^ Centrioles, approximately 500 nm in length and 250 nm in width, exhibit a characteristic barrel-shaped structure, arising from a nine-fold symmetrical arrangement of microtubule triplets.^2,3^ Centrosomes serve as microtubule-organizing centers and as such provide a scaffold for trafficking and organization of organelles and vesicles, while, during cell division, they facilitate microtubule spindle assembly, ensuring accurate chromosomal segregation.^4^

**Figure 1.**
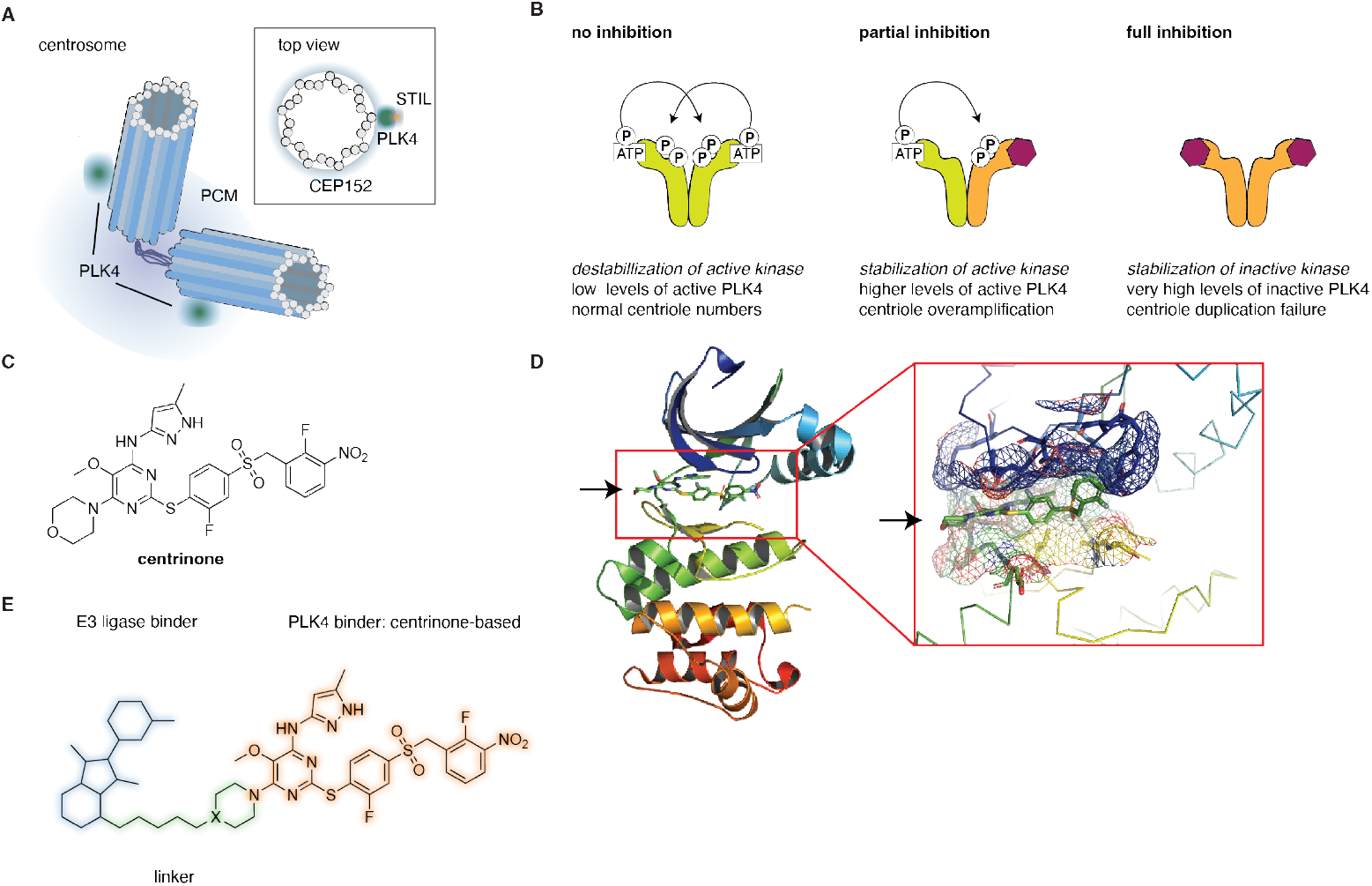
PLK4 at centrosomes and its inhibition by centrinone. (A) Schematic representation of the centrosome. (B) Cellular levels of PLK4 are tightly controlled through autoregulation, ensuring faithful centriole duplication. Partial inhibition by centrinone leads to stabilization of the active kinase and centriole overduplication, whereas full inhibition results in accumulation of inactive PLK4 and centriole duplication failure. (C) Structure of centrinone. (D) Crystal structure of centrinone bound to the PLK4 kinase domain (PDB: 4YUR). (E) General structure of centrinone-based molecular degraders.

Proper cell division requires tight control of centriole numbers - with precisely 4 centrioles at the onset of mitosis, following a single centriole duplication event at G1/S^.5^ A protein with a critical role in this process of centriole biogenesis is Polo-like kinase 4 (PLK4), a serine/threonine kinase localized at the proximal region of centrioles, on a CEP152-containing torus (Figure 1A).^6,7^ To achieve centriole duplication rather than overamplification, activity and levels of PLK4 must be tightly regulated. PLK4 exists as a homodimer, with its activity heavily regulated by phosphorylation.^8,9^ One of the critical phosphorylation events is the trans-auto phosphorylation of the degron region, leading to the recruitment of the SCFβ-TRCP E3 ligase, ubiquitination and its subsequent degradation by the proteasome, keeping the levels of PLK4 low at centrioles and preventing overamplification (Figure 1B).^10,11^

As a key regulator of centriole duplication, any defects in the function or expression level of PLK4 can lead to detrimental consequences. Indeed, overexpression of PLK4 in mammalian cells causes centrosome amplification, promoting aneuploidy and spontaneous tumorigenesis.^12^ Furthermore, several studies have reported PLK4 overexpression in many types of cancers, with its overexpression being a biomarker for poor prognosis in patients, as well as poor response to therapy.^13–16^ Therefore, PLK4 represents an attractive therapeutic target for new chemotherapeutic drugs. Thus far, only one PLK4 inhibitor, CFI-400945, has entered Phase II clinical trials for the treatment of several malignancies.^17,18^ However, CFI-400945 lacks PLK4-selectivity,^19^ which led to the endeavours to create more selective binders, resulting in the discovery of centrinone (Ki=0.16 nM), a highly selective reversible inhibitor of PLK4 (Figure 1C).^20^ Of note, partial inhibition of PLK4 results in stabilization of the active kinase and therefore centriole overamplification, whereas full inhibition results in accumulation of the inactive kinase and centriole dilution after each cell division (Figure 1B).^18,21,22^

Here, we aimed to take an alternative approach to classical inhibition, by converting centrinone into a proteolysis-targeting chimera (PROTAC). PROTACs are bifunctional molecules, composed of a ligand for a protein of interest (POI) covalently attached to an E3 ligase ligand via a small linker, and as such are capable to hijack the ubiquitin-proteasome system to induce degradation of a POI.^23–25^ In the last two decades, PROTACs emerged as powerful modalities in the treatment of various diseases, largely due to their different mode of action compared to classical inhibitors. Given the particular mode of action of centrinone, yielding high cellular levels of inactive protein, we envisioned that a centrinone-based PROTAC would provide a valuable chemical biology tool to investigate the potential roles of PLK4 beyond its main enzymatic function, which is the phosphorylation of STIL (Figure 1A).^26,27^ We present the synthesis of 13 molecular degraders targeting PLK4 (Figure 1E), using centrinone as the targeting ligand. The in-vitro binding affinities of all synthesized molecules were in the low nM range, comparable to centrinone, yet large differences in cellular PLK4 levels were observed. To understand the mechanism of action we selected two distinct molecules, **D6** and **D10**, for further evaluation, revealing an intricate balance between the intrinsic propensity of centrinone to increase PLK4 levels, and the ability of the degraders to target PLK4 for destruction, especially within the apparently shielded environment at the centrioles.

## RESULTS AND DISCUSSION

### Synthesis and *In Vitro* Evaluation of Centrinone-Based Molecular Degraders

As a starting point, we analyzed the crystal structure of the PLK4 kinase domain, with centrinone bound to it (PDB: 4YUR), to identify the solvent-exposed morpholine ring as the optimal position for structural modifications (Figure 1D). This approach was already verified in the design of fluorogenic probes which effectively target PLK4.^22^ We envisioned eleven PROTACs based on centrinone, tethered to a CRBN-binding ligand (thalidomide or its analogues) via linkers of varying lengths and flexibilities (Figure 1E). Within this group, **D1-7** contained flexible polyethylene-glycol or aliphatic chain linkers, while **D8-D11** included more rigid piperazine-based linkers. As an alternative to CRBN targeting with thalidomide, we sought to incorporate a recently designed covalent fumarate handle, reported to target RNF126 E3 ligase,^28^ resulting in **D12** and **D13**. The common centrinone precursor for all the probes, **14**, containing an electrophilic chloropyrimidine moiety suitable for nucleophilic aromatic substitution, was synthesized as previously reported, with minor modifications (Supplementary Scheme 1). Compound **14** was then converted into piperidine **16** (X=C) and piperazine **18** (X=N) (Supplementary Scheme 2), and these were utilized for the synthesis of **D1-7, D9-11** and **D8, D12-13**, respectively (Supplementary Schemes 3-9, Figure 2C). Thalidomide-linker fragments were synthesized separately, starting either from 4-hydroxythalidomide for **D1-8**, or 3-/4-fluorothalidomide for **D9-11** (Supplementary Schemes 3-8). **D12-13** were obtained by regular amide coupling of phenyl-fumarate units, **40** and **42**, with **16** and **18**, respectively (Supplementary Scheme 9).

**Figure 2.**
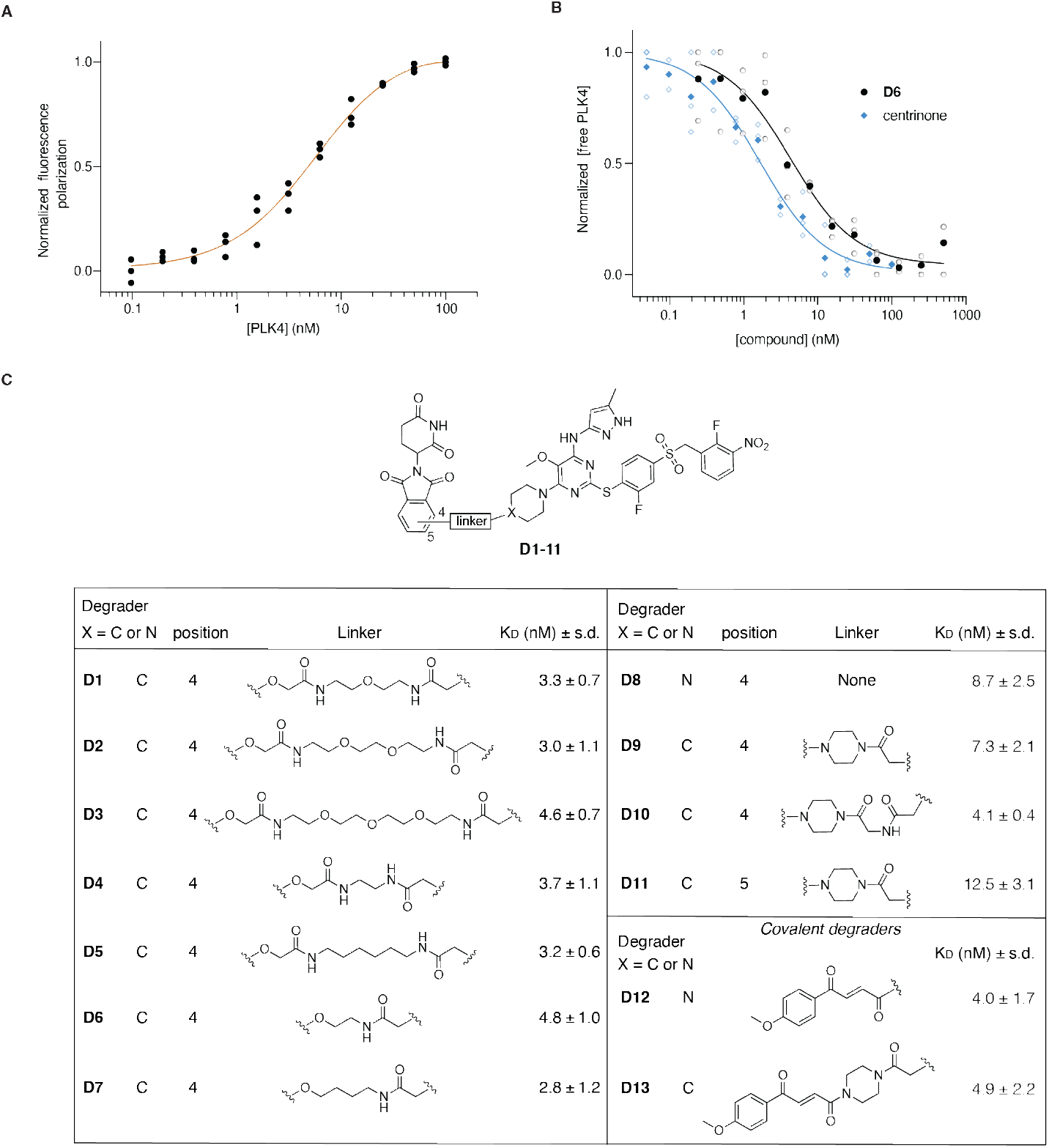
Evaluation of the binding affinities of molecular degraders to PLK4-KD using fluorescence polarization. (A) Representative binding curve of TMR-centrinone tracer **45** to PLK4-KD (technical replicates shown). (B) Full binding curves of centrinone (blue) and **D6** (black) to PLK4-KD, solid symbols present the mean of 3 independent curves (open symbols). (C) Measured binding affinities (mean +/- s.d., N = 3 independent experiments) for all the synthesized PROTACs with the structures shown.

To assess the binding affinities of the synthesized molecules to PLK4, we expressed and purified the 6xHistagged PLK4 kinase domain (PLK4-KD) and measured the dissociation constants using fluorescence polarization. For this purpose, we synthesized a fluorescent centrinone-based tracer **45** (Supplementary Scheme 10), by coupling **16** to a tetramethylrhodamine (TMR) fluorophore via a short linker. Tracer **45** exhibited high-affinity binding to the purified PLK4-KD (K_d_=2.34 ± 0.80 nM), comparable to centrinone (Figure 2A, B). We then used this probe in a competitive binding assay to quantify the binding affinities of the PROTACs to PLK4-KD (Figure 2B, C). Satisfyingly, all of the synthesized molecules showed high binding affinities for PLK4 *in vitro*, with K_D_ values in the nanomolar range (Figure 2C), similar to centrinone (K_d_=2.26 ± 0.49 nM), indicating that our structural modifications did not impact the binding to PLK4 and allowing us to proceed with in-cell experiments.

### Biological Evaluation of PLK4 Degradation

Following the fluorescence polarization results, we tested the abilities of the probes to degrade PLK4 *in cellulo*. Breast cancer MDA-MB-231 cells, which show high PLK4 expression levels, were treated with 1 µM of each of the compounds and the presence of PLK4 was probed by western blot. We determined 24 h to be an optimal time of treatment to observe effects of the molecules on PLK4 levels (Supplementary Figure 1). PLK4 inhibition by centrinone causes elevation of PLK4 levels, due to the inhibition of autophosphorylation, which in turn prevents its degradation (Figure 1B, 3A). However, we found a high heterogeneity in PLK4 levels of cells treated with the degraders. Several of the synthesized probes induced PLK4 accumulation similar to centrinone (Figure 3A, -HyThal conditions), most evident for **D2, D3, D5, D7, D12** and **D13**, whereas others showed no or very little change compared to DMSO control (**D1, D4, D6, D8**). To understand this mix of behaviours, which could be the result of pure inhibition without degradation, degradation alone, or inefficient degradation/inhibition, we probed the efficacy of the probes in the presence of 4-hydroxythalidomide (HyThal). In this competition assay HyThal was used at a 10-fold excess (10 µM) to block all available CRBN E3 ligases (Figure 3A) and prevent ternary complex formation. In this case, the sole biological effect induced by the degraders would be inhibition of PLK4, which should lead to its accumulation. Indeed, this proved true for seven of the PROTACs (**D5**-**D11**, Figure 3B,C), with the most pronounced increase of approximately 2-fold upon HyThal co-treatment for **D6** and **D10** (Figure 3C). While the fold increase for these molecules was similar, the absolute induction differed greatly. In the absence of HyThal, **D6** did not induce any accumulation of PLK4, whereas **D10** already showed a 1.5-fold increase compared to DMSO. Indeed, the level of PLK4 in **D10** +HyThal-treated cells reached almost that induced by centrinone, suggesting that **D10** more potently inhibited PLK4 than **D6**. We therefore selected both **D6** and **D10** for further investigation, as both showed prominent degradation, with **D6** appearing as a poor inhibitor and **D10** showing potent inhibition at 1 µM. To determine their *in cellulo* potencies, we tested **D6** and **D10** in a range of concentrations (10 µM to 100 nM) in both MDA-MB-231 and HeLa (cervical cancer) Centrin1-GFP^29^ cell lines in the absence or presence of an excess of HyThal (Figure 4, Supplementary Figure 2).

**Figure 3.**
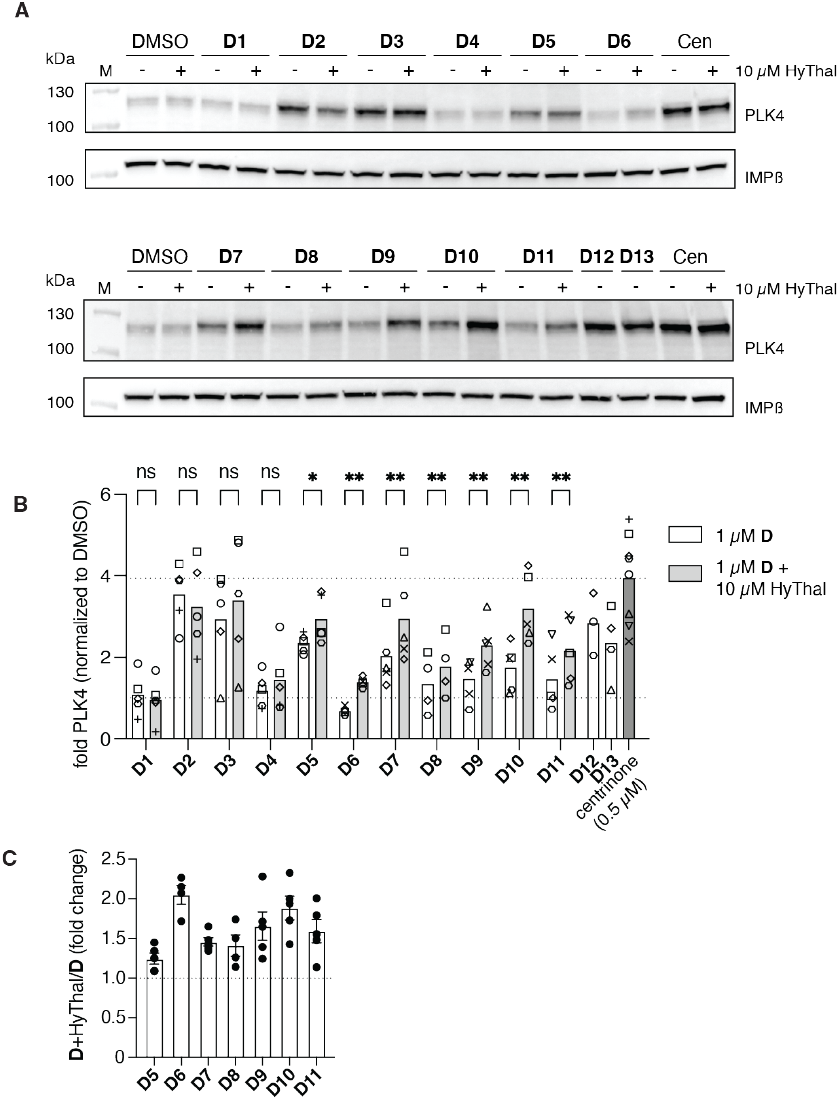
Degradation capabilities of **D1-D13** (A) Representative immunoblot showing the effect of 1 µM **D1-D13** on PLK4 levels in the MDA-MB-231 cell line in the presence or absence of 10 µM HyThal. IMPβ: loading control. (B) Quantification of immunoblot PLK4 band intensity of four independent experiments (symbols matched for experiment; bar represents mean). A mixed-effects model with Fisher’s LSD, without multiple comparisons, was used to calculate the p-values between D and D+HT. * : p<0.0332, ** : p<0.0021. (C) Ratio of PLK4 band intensities with and without 4-hydroxythalidomide (four independent experiments).

**Figure 4.**
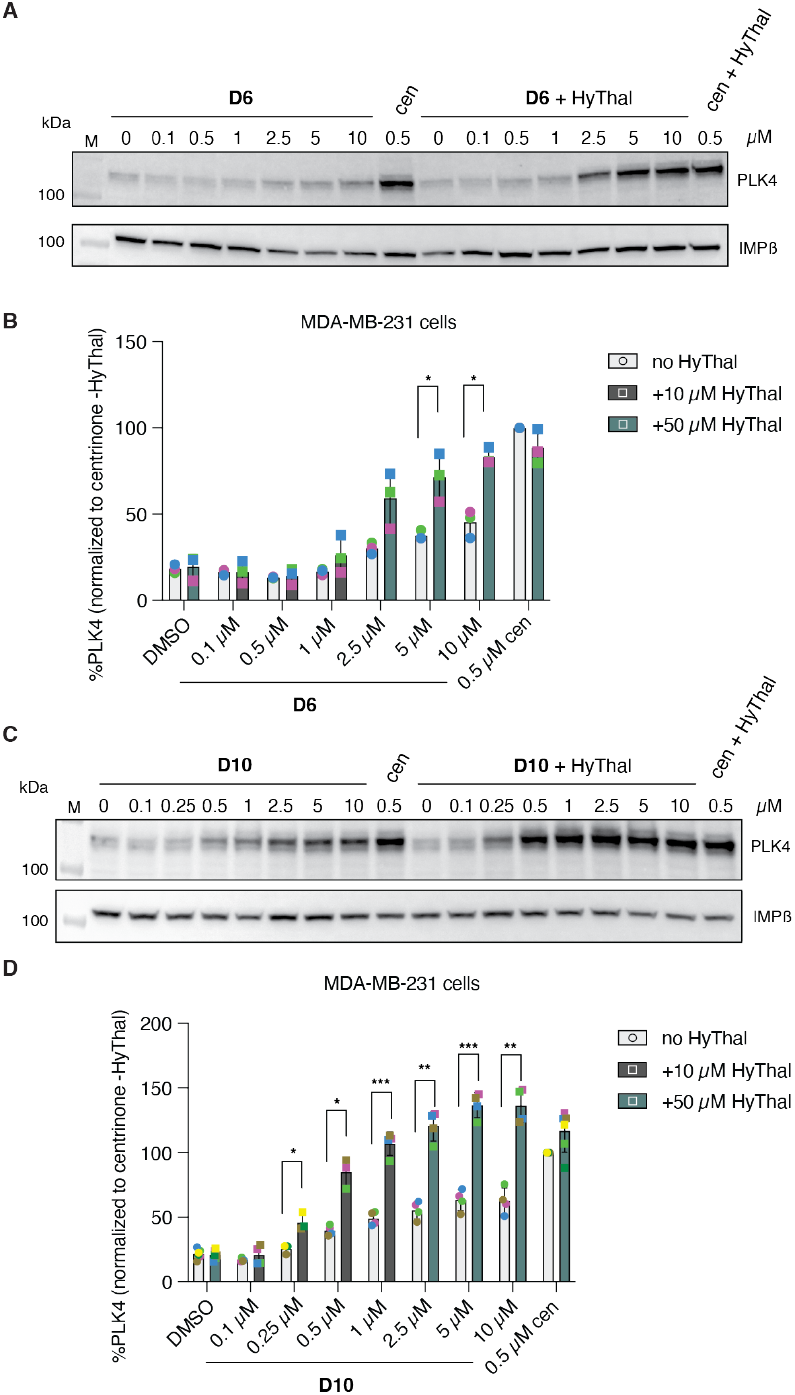
Dose-dependent changes in PLK4 levels in MDA-MB-231 cells. (A,C) Representative immunoblots of cells treated with the indicated concentration of **D6** (A) or **D10** (C) in the presence or absence of 10 or 50 µM HyThal. IMPβ: loading control (B,D) Bargraphs showing the PLK4 quantification of 3 (B) or 4 (D) independent experiments (symbols). Bars represent means. Unpaired t-tests, * : p<0.0332, ** : p<0.0021, *** : p<0.0002

These experiments revealed several things. First, while western blot quantification of low levels of PLK4 proved not accurate enough to obtain statistical significance for all comparisons, in both cell lines, and for both compounds, there was a clear upward trend with HyThal competition, indicative of compound-induced degradation of PLK4 (Figure 4A-D, Supplementary Figure 2). The ability of both compounds to inhibit, and thus accumulate PLK4, was stronger in HeLa cells (Supplementary Figure 2) compared to MDAMB-231 cells (Figure 4), which may be attributed to differences in basal levels of PLK4 between these cells and/or cell permeability affecting intracellular compound concentrations. Second, **D10** was the more potent inhibitor of the two, resulting in accumulation of PLK4 in HeLa cells starting at the 0.25 µM concentration, whereas for **D6** about 10-fold more compound was needed to detect an increase in PLK4 by western blot. These results highlight the complexity of the biology induced by both **D6** and **D10** and that the resulting PLK4 levels are a product of two concurrent inseparable events: inhibition and degradation. Moreover, it is not possible to distinguish by western blot the distribution of uninhibited, partially, and fully inhibited PLK4, and thus band intensities may not be reflective of the induced phenotypes.

To assess this, we performed centriole scoring experiments. The recruitment of PLK4 to centrioles is strictly regulated, and it is known that centrinone inhibition of PLK4 leads to prevention of procentriole formation, resulting in centriole depletion in dividing cells. Contrastingly, partial inhibition, by lower concentrations of centrinone, causes centriole overamplification, through inhibition of degradation-inducing autophosphorylation (Figure 1B).^18,21,22^ We treated HeLa centrin-GFP cells with varying concentrations of **D6** and **D10** in the presence and absence of HyThal and counted the number of centrioles present after 24 h of treatment (Supplementary Figure 3). In accordance with the inhibitory potencies (Supplementary Figure 2), **D10** treatment resulted in more potent centriole depletion compared to **D6**, whereas for **D6** centriole overamplification was the predominant phenotype. No significant differences were found in the induced phenotypes in the presence of HyThal for the stronger inhibitor **D10**. For **D6**, at the 5 µM concentration there was a slight shift from overamplification to centriole depletion in the presence of HyThal, indicative of stronger inhibition in the absence of degradation. To exclude the possibility that this was the result of a higher effective concentration of the probe (as no E3 ligase is bound), rather than a reflection of a shift in degradation/inhibition, we sought to further confirm the ability of **D6** and **D10** to degrade PLK4 in a more direct manner.

### D6 and D10 Potently Degrade an Autoregulation-Deficient PLK4 Mutant and Non-Centriolar Endogenous PLK4

As increasing levels of PLK4 upon inhibition prevented a proper assessment of the degradation capacities of **D6** and **D10**, we overexpressed a mutant PLK4, PLK4(K41M), that is incapable of autoregulation in HeLa cells, preventing the complex phenotype as observed with endogenous PLK4.^10^ We observed very potent degradation of overexpressed GFP-PLK4(K41M)-FLAG by both **D6** and **D10** by western blot (Figure 5A,B, Supplementary Figure 4A,B). However, degradation plateaued at ∼30% PLK4(K41M) remaining, and so we turned to fluorescence microscopy to understand where the remaining PLK4 was localized.

**Figure 5.**
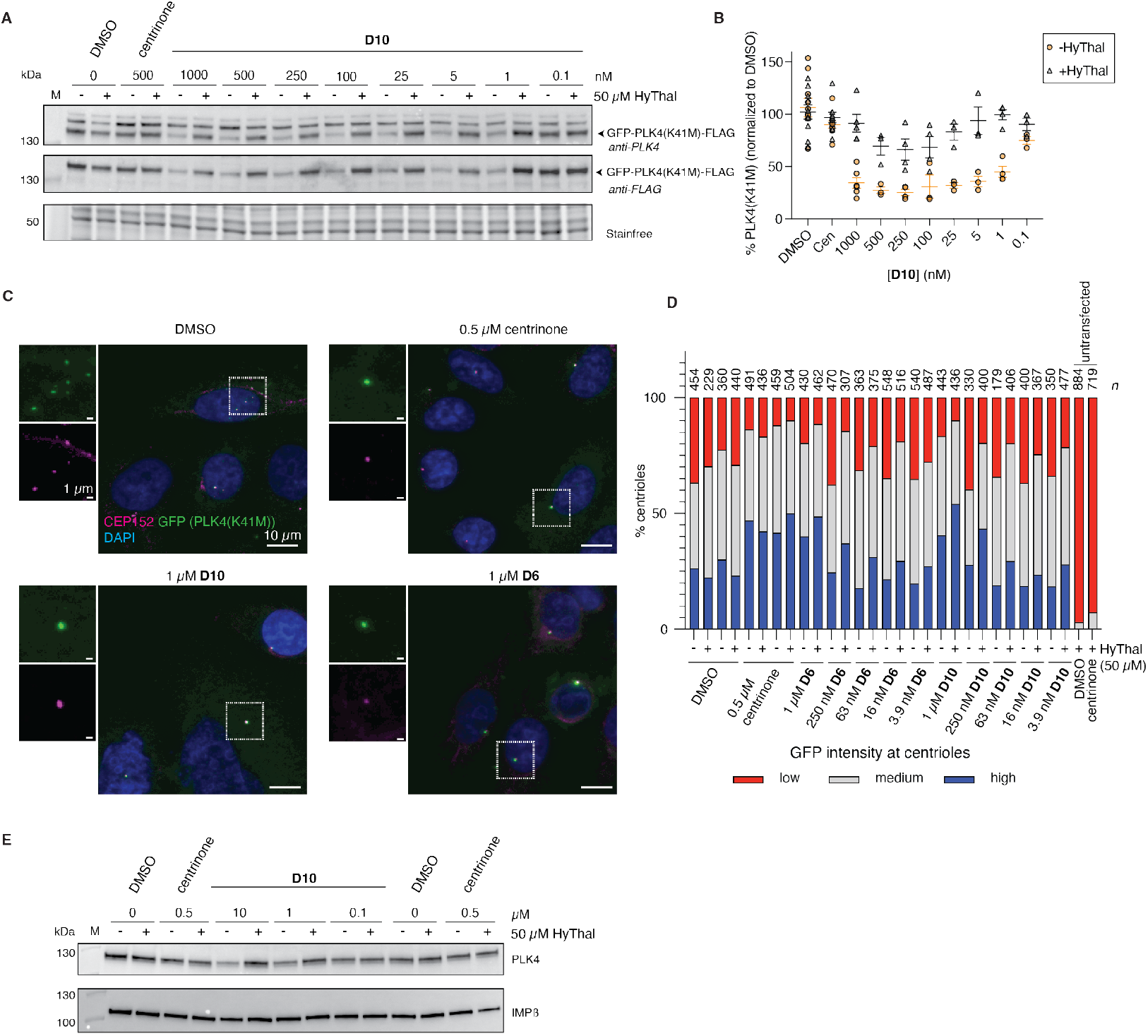
GFP-PLK4(K41M)-FLAG and non-centriolar PLK4 are degraded by **D6** and **D10.** (A,B) HeLa cells overexpressing GFP-PLK4(K41M)-FLAG were treated with the indicated concentration of **D10** with or without 50 µM HyThal for 24 h and probed for the presence of PLK4 and FLAG by western blot. (A) representative WB and (B) quantification of PLK4 immunoblots of N≥3 independent experiments (indicated by symbol), line and error bars indicate mean +/-SEM. (C) Representative immunofluorescence images and (D) quantification of GFP intensity at the centrioles of HeLa cells expressing GFP-PLK4(K41M)-FLAG. n = number of centrioles analyzed, data shown is from one out of two independent experiments. (E) Acentriolar RPE-1 USP28^-/-^ cells were treated with DMSO, 500 nM centrinone or compound **D10** with or without 50 µM HyThal and the degradation of endogenous PLK4 was assessed by western blot. Representative blot of two independent experiments.

As overexpression was done transiently, we observed high heterogeneity in expression levels, with a small percentage (typically ∼5%) of highly transfected cells having predominant cytoplasmic GFP signal, and lower expressing cells showing both cytoplasmic and centriolar GFP signal. We found that the overall percentage of highly overexpressing cells was reduced by **D10**, in line with the western blot result (Supplementary Figure 4C), which could be competed with HyThal to levels observed with centrinone. Of note, centrinone-treated cells also had fewer bright cells than DMSO control, because of stringent masking criteria in the analysis, excluding bright round cells. However, for centrinone, in contrast to **D10**, this effect was independent of the presence of HyThal and likely reflects a mild toxicity. As expression levels found by western blot for DMSO and centrinone (Fig 5A,B) treatments were similar, highly transfected cells likely contributed greatest to the overall signal detected by blot, and we assume that those cells are more easily washed away during immunofluorescence protocols, compared to direct lysis of the cells for western blot. At all concentrations tested, we could observe a dot of PLK4 at the centrioles, both using a PLK4 antibody (detecting mutant and endogenous PLK4, Supplementary Figure 4D) and a GFP antibody (solely detecting mutant PLK4, Figure 5C,D). The intensity of PLK4 at centrioles increased using centrinone, or high (1 µM) concentration **D6** or **D10** in the presence of HyThal, consistent with inhibition and stabilization of endogenous PLK4 (Supplementary Figure 4D). The fraction of GFP positive centrioles also increased under the same conditions, consistent with a model where endogenous PLK4 can phosphorylate and thereby stabilize the mutant (Figure 5D).^10^ While the difference between with and without HyThal indicates that there is some degradation occurring which prevents levels at the centrioles from increasing in the absence of HyThal, similar to what we found for overall amounts of endogenous PLK4 by western blot (Figure 4, Supplementary Figure 2), these levels did not go below that found for DMSO-treated cells. Together, this raises the possibility that a large proportion of PLK4 at the centrioles is in some way shielded to small molecule-mediated degradation, yet not to inhibition. This would also provide an explanation as to why there was little to no phenotypic shift observed upon HyThal treatment in the centriole scoring experiment (Supplementary Figure 3).

To further test this hypothesis, we next used RPE-1 (non-transformed retina pigment epithelial cells) USP28-/- cells, that were chronically treated with centrinone to fully deplete centrioles while remaining cycling-competent (USP28 is part of a mitotic surveillance mechanism that arrests the cell cycle of cells lacking centrosomes^30^). Because of chronic centrinone treatment, we found that the PLK4 levels in these cells are relatively high, enabling detection by western blot, yet its localization is non-centriolar as the cells lack centrioles. We found that PLK4 levels quickly decreased to barely detectable levels upon centrinone washout (Supplementary Figure 5A), and for that reason cells were maintained with centrinone until the start of the experiment, where they were switched to medium without centrinone after a single wash. Under these experimental conditions it is likely that low concentrations of centrinone remained present inside the cells, preventing the centrioles from re-forming and keeping PLK4 levels stable. Indeed, 24 h after washing out centrinone, DMSO-, centrinone- or **D10**-treated cells were still acentriolar (as judged by CEP152 staining, Supplementary Figure 5B). PLK4 levels were stable between DMSO- or centrinone-treated cells but decreased in **D10**- or **D6**- treated cells in a dose-dependent manner (Figure 5E, Supplementary Figure 5C). Protein levels could be fully restored in the presence of HyThal, illustrating CRBN-dependent degradation of endogenous PLK4 induced by **D6** and **D10**. Importantly, no conclusions about degradation potency should be drawn in this experimental setup, as the starting point is centrinone-bound PLK4, which requires an additional displacement step before the degraders can bind. Nevertheless, this experiment conclusively proves the ability of the PROTACs to degrade endogenous PLK4 in the absence of centrioles.

## Discussion

The selective PLK4 inhibitor centrinone has enabled the controlled depletion of centrioles from mammalian cells, but, through its mechanism of action, leads to high cellular levels of inhibited PLK4. Here, we sought to leverage the high affinity binding and selectivity of centrinone to prepare a library of molecular degraders that would enable cellular PLK4 depletion. Towards the same goal, Sun et al. recently synthesized a PROTAC,^31^ named SP27, based on another PLK4 inhibitor, CZS-034. Surprisingly, while CZS-034 has been reported as a type I (ATP-competitive)-inhibitor, like centrinone, no PLK4 stabilization has been shown for this compound.^32^ To the best of our knowledge, there is no experimental evidence why this is the case, but possibly the off-rate for this compound is fast, enabling occasional autophosphorylation. SP27-mediated PLK4 degradation was potent but also incomplete. A detailed mechanistic investigation of SP27 or CZS-034 on centrioles, or PLK4 localization to centrioles, has not been performed, in contrast to the well-studied effects of centrinone on PLK4 stability and centriole duplication. Therefore, we decided to employ centrinone as the ligand of choice for our PROTAC design. Satisfyingly, our synthesized centrinone-based degraders retained low nanomolar affinity for the kinase domain of PLK4, illustrating that the scaffold is tolerant to chemical changes, in agreement with what has been described for a centrinone-based fluorescent probe.^22^ Upon cellular evaluation, however, we found large differences between the different degraders, ranging from pure inhibition to selective degradation. For the two most potent degraders, **D6** and **D10**, we discovered a cellular phenotype resembling that of centrinone, but especially for **D10**, the balance between inhibition and degradation remained strongly at inhibition. To circumvent the autoregulation of PLK4, a phosphorylation deficient mutant provided the means to conclusively prove the ability of these compounds to degrade PLK4. Under the conditions tested, we could not, however, degrade endogenous PLK4 beyond basal levels, nor could we remove PLK4(K41M) from the centrioles, prompting us to hypothesize that PLK4 at the centrioles may be shielded from degradation. The different capacity of **D6** and **D10** to inhibit PLK4 further illustrates that there may be differential accessibility for these compounds to PLK4 at the centrioles, as both compounds were equally potent in their degradation of a cytoplasmic PLK4 mutant and showed the same binding affinity to purified PLK4 in vitro. PLK4 at centrioles has been shown to have strong binding interactions with both scaffolding proteins (CEP152)^6,7^ and phosphorylation substrates (STIL).^26,27^ Centrinone binding, while preventing the PLK4-STIL interaction, does not interfere with the PLK4-CEP152 interaction, but rather increases the number of interaction sites to 9 over prolonged exposure times.^33^ Moreover, Scott *et al*. have shown that centrinone-mediated inhibition of PLK4 leads to much reduced turnover time (and thus accumulation) of PLK4 at centrioles, more so than was observed with inhibition of proteosomal degradation.^33^ Possibly, this PLK4-CEP152 interaction blocks the accessibility to CRBN - a requirement for functional PROTAC-mediated degradation. For example, Gasic *et al*. have shown that tubulin is able to evade CRBN targeted degradation, conceivably because of its association with other proteins in tubulin dimers or microtubules.^34^ Satisfyingly, we were able to conclusively show that endogenous PLK4 could be degraded in acentriolar cells, strengthening the hypothesis that PLK4 at centrioles is shielded. However, even in the absence of centrioles, degradation was incomplete. This could be due to the imperfect experimental system, employing centrinone to deplete cells of centrioles and artificially enhance PLK4 levels, which necessitates a competition between the probes and centrinone. In addition, it is unknown if the cytoplasmic PLK4 retains some level of interaction with scaffolding proteins even in the absence of centrioles, which could be an alternative explanation of incomplete degradation. We envision that our probes will be valuable tools to further decipher the accessibility of PLK4, when combined with genetic perturbation of key PLK4 interactors.

## Conclusions

In this study, we report the successful synthesis of a series of molecular degraders of PLK4, with low nanomolar in vitro binding affinities. PROTACs **D6** and **D10**, while both able to degrade PLK4, differed severely in their ability to inhibit PLK4, with **D6** being a poor inhibitor and **D10** a strong inhibitor. Both compounds were equally potent to degrade a PLK4 mutant and endogenous PLK4 in acentriolar RPE-1 cells. Together, our study uncovers that PLK4 accessibility at centrioles is limited for targeted degraders, enabling binding and inhibition, but largely preventing degradation.

## Author contributions

Conceptualization: A.S., S.H.; Formal analysis: A.S., A.K., S.H.; Funding acquisition: S.H., P.M.; Investigation: A.K., A.S., C.B.; Resources: P.M., S.H.; Supervision: A.S., P.M., S.H.; Visualization: A.S., S.H.; Writing – original draft: A.K., S.H.; Writing – review & editing: A.K., A.S., C.B., P.M., S.H.

## Data availability

The raw experimental data that support the findings of this study are available in Zenodo with the identifier 10.5281/zenodo.17738398. Supplementary Information contains Supplementary Figures 1-5, Supplementary Schemes 1-10, full experimental procedures (biology and chemistry) and analytical data (NMR spectra)

## Acknowledgements

We would like to thank the ACCESS Geneva and Bio-imaging Center-Photonic facilities for their help with microscopy experiments, and particularly Dimitri Moreau, for their assistance with fluorescence microscopy analyses. We thank Karen Oegema and Arshad Desai (UCSD) for the RPE1 USP28^-/-^cells. This research was funded by the University of Geneva, the Swiss National Science Foundation (project grants 189246 and 10000608 to S. H, and 310030_208052 to P. M.) and supported by funding from the European Research Council (ERC) under the European Union’s Horizon 2020 research and innovation programme (grant agreement n° 948750, DestCilia, to S.H.).

