## Supplementary information for "Tipping the balance: Synthesis and evaluation of centrinone-based degraders of polo-like kinase 4"

Table of contents

1. Supplementary Figures 1-5
2. Supplementary Schemes 1-10
3. Biology
4. Chemistry
5. References
6. Full western blots
7. NMR spectra

#### 1. Supplementary Figures

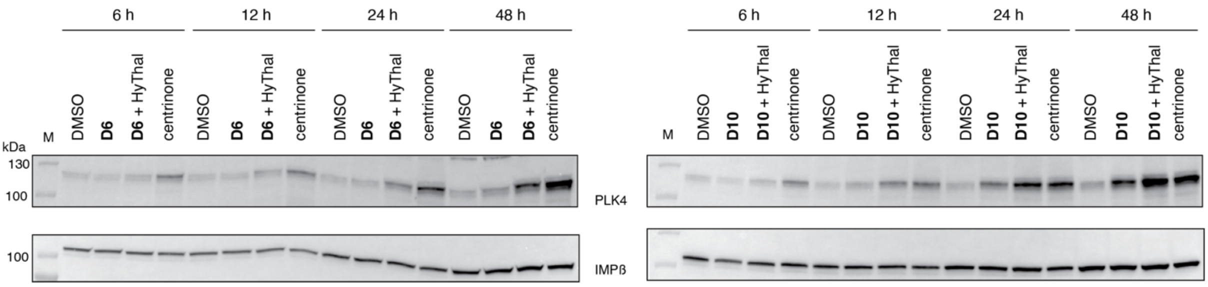


**Supplementary Figure 1** HeLa cells were treated with 500 nM centrinone or 2.5 µM **D6** or **D10** in the presence of absence of 50 µM HyThal for the indicated timepoints and probed for PLK4. IMPβ, loading control.

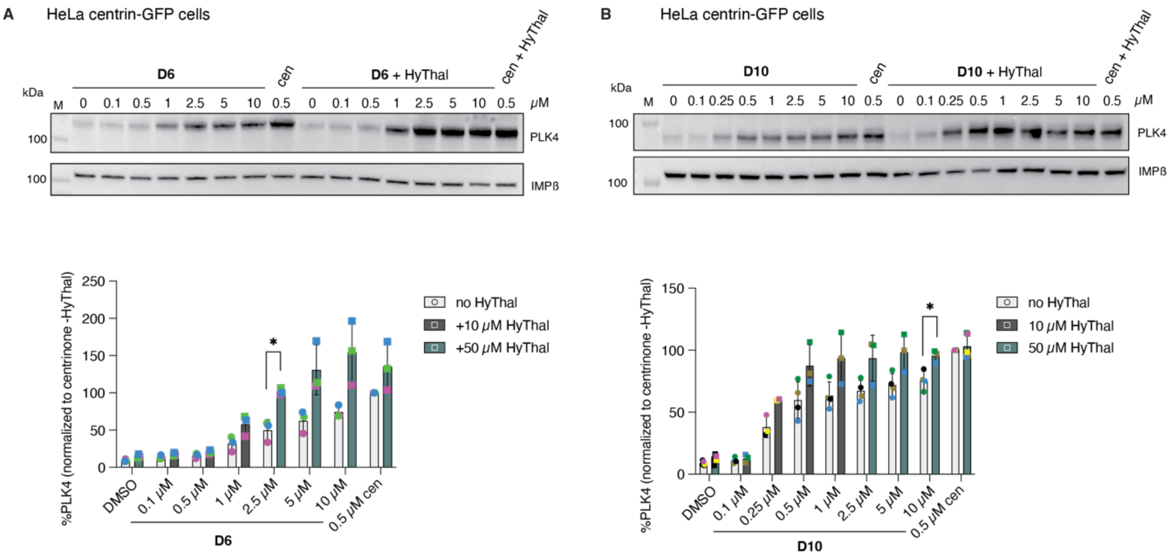


**Supplementary Figure 2.** Dose-dependent changes in PLK4 levels in HeLa centrin-GFP cells (extended Figure 4). (A,B) (top) Representative immunoblots of cells treated with the indicated concentration of **D6** (A) or **D10** (B) in the presence or absence of HyThal with (below) bargraphs showing the PLK4 quantification of 3 (A) or 4 (B) independent experiments (symbols). Bars represent means. Unpaired t-tests, *: p<0.0332, **: p<0.0021, ***: p<0.0002

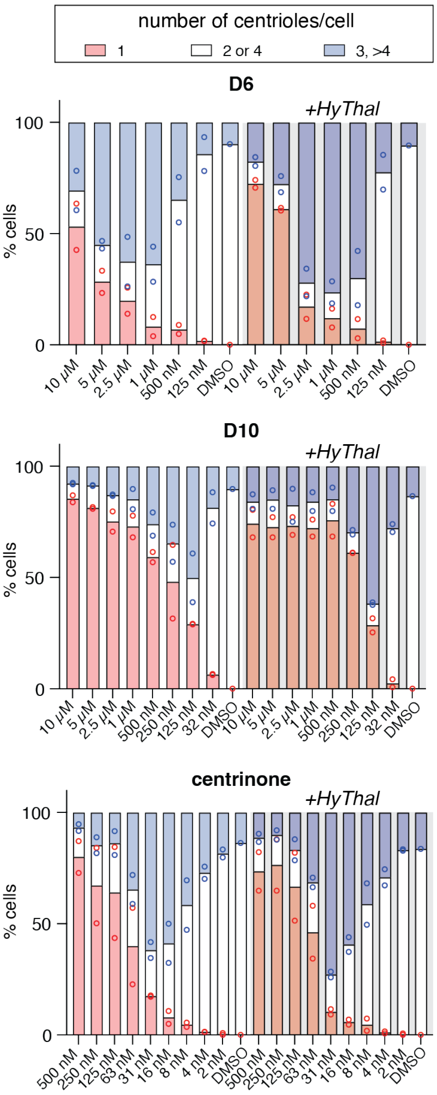


**Supplementary Figure 3.** Dose-dependent loss of centrioles. HeLa centrin-GFP cells were treated with the indicated concentrations of compounds in the presence (right side of graph, dark colors) or absence (left side of graphs, clear colors) of 50 µM (for concentrations > 1 µM) or 10 µM HyThal for 24 h, before the number of centrioles was scored based on the GFP signal. 200-300 cells were scored per experiment, 2 independent experiments (mean of experiment represented by symbol). Bar represents mean of both experiments.

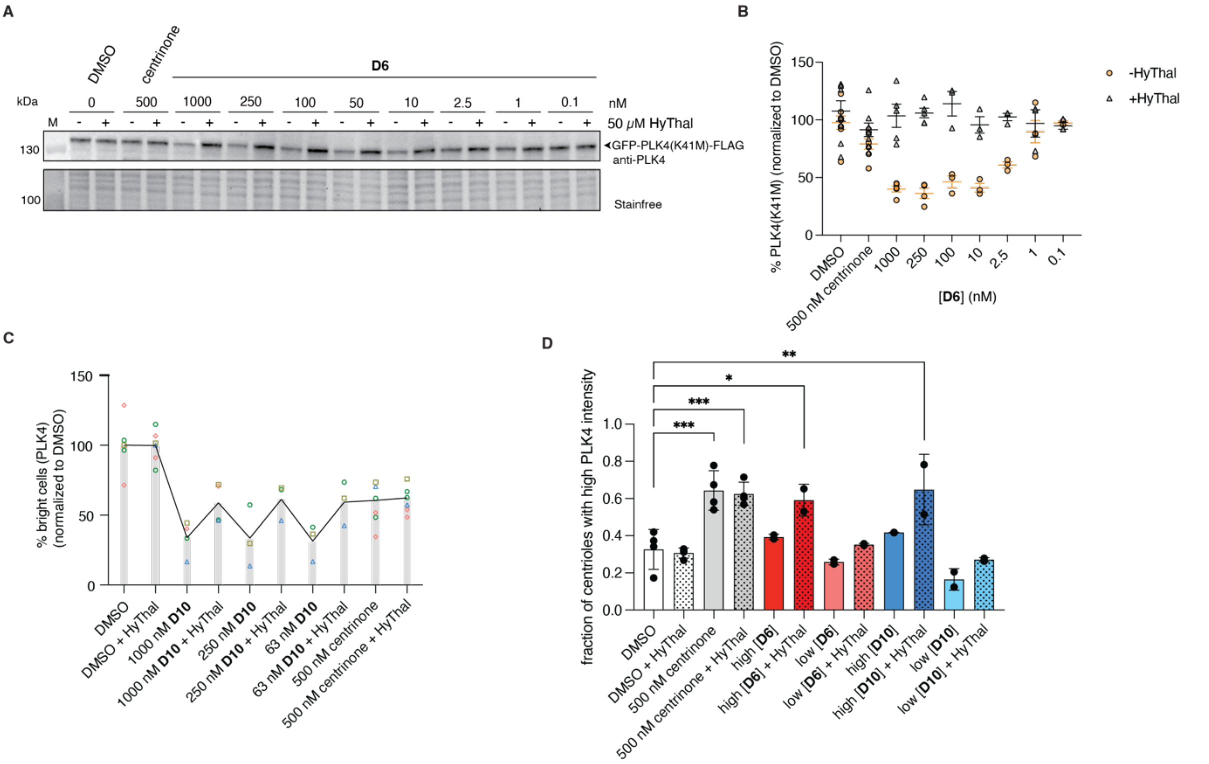

**Supplementary Figure 4.** GFP-PLK4(K41M)-FLAG is degraded by **D6** and **D10** (extended Figure 5). (A,B) HeLa cells overexpressing GFP-PLK4(K41M)-FLAG were treated with the indicated concentration of **D6** with or without 50 µM HyThal for 24 h and probed for the presence of PLK4 by western blot. (A) representative WB and (B) quantification of N≥3 independent experiments (indicated by symbol), line and error bars indicate mean +/- SEM. (C) Quantification of bright HeLa cells expressing GFP-PLK4(K41M)-FLAG (integrated cell PLK4 intensity), normalized to the respective DMSO (usually technical duplicate per experiment), of at least 3 independent experiments. Symbol colors refer to the same experiment. (D) Quantification of the fraction of PLK4 positive centrioles (defined as: signal intensity> non-transfected cells) under the different treatments. Two independent experiments with 400-500 centrioles quantified per condition/experiment. One-way ANOVA with Dunnett's multiple comparisons test, with a single pooled variance. *: $p<0.0332$, **: $p<0.0021$, ***: $p<0.0002$

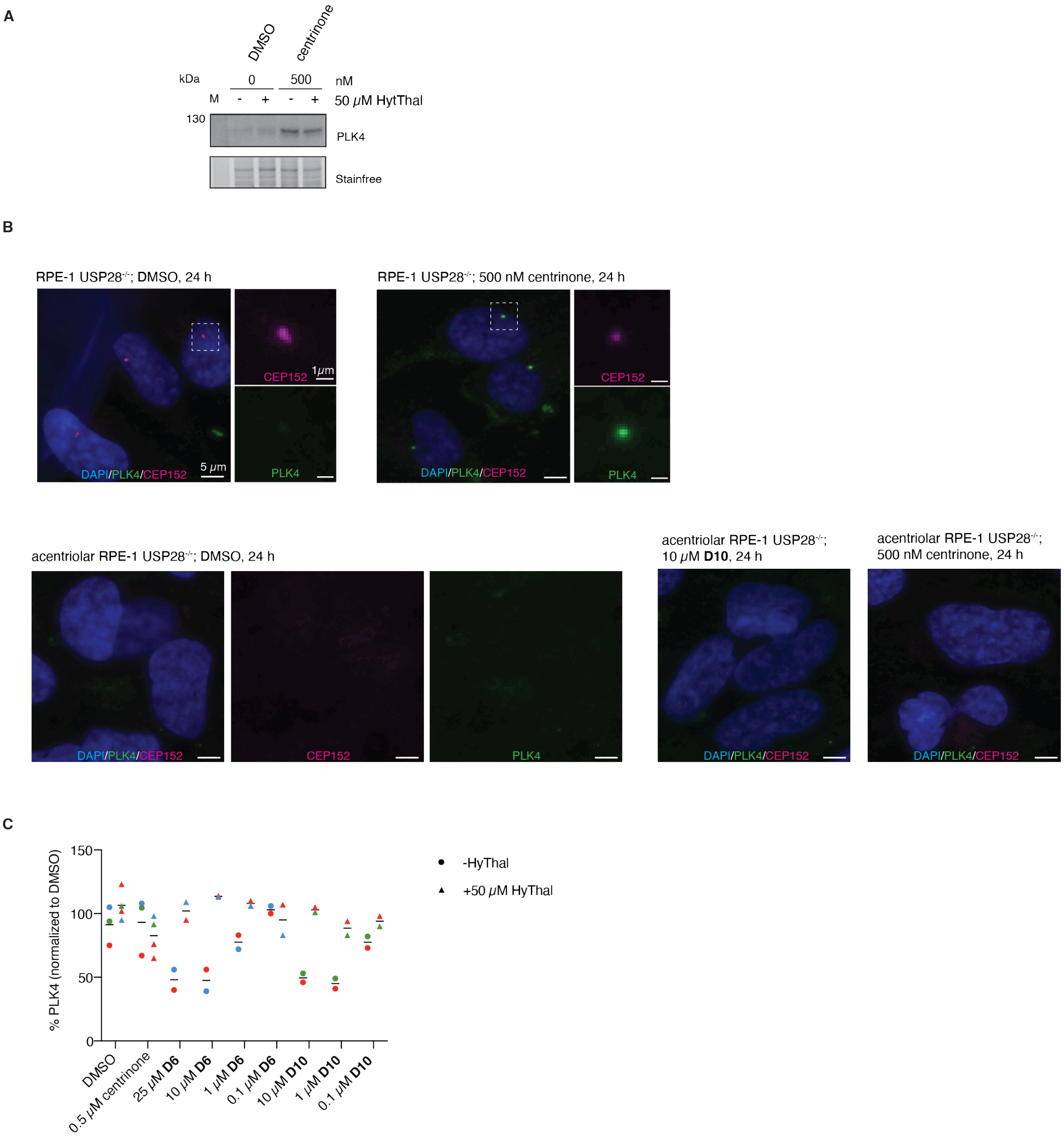


**Supplementary Figure 5.** (A) Acentriolar RPE-1 USP28$^{-/-}$ cells rapidly loose PLK4 upon centrinone wash-out as indicated by western-blot 42 h post-seeding with or without centrinone. (B) representative micrographs showing the presence of CEP152$^{+}$, PLK4$^{-}$ (below level of detection) centrioles in DMSO-treated parental RPE-1 USP28$^{-/-}$ cells, as well as a single CEP152$^{+}$, PLK4$^{+}$ centriole upon centrinone treatment. In contrast, upon chronic exposure to centrinone (lower panels), no centrioles are detected 24 h after incomplete centrinone washout (DMSO condition) or added **D10** or centrinone. (C) Acentriolar RPE-1 cells were treated with the indicated compounds and PLK4 levels were detected by western blot. Quantification of endogenous PLK4 of two independent experiments.

#### 2. Supplementary Schemes

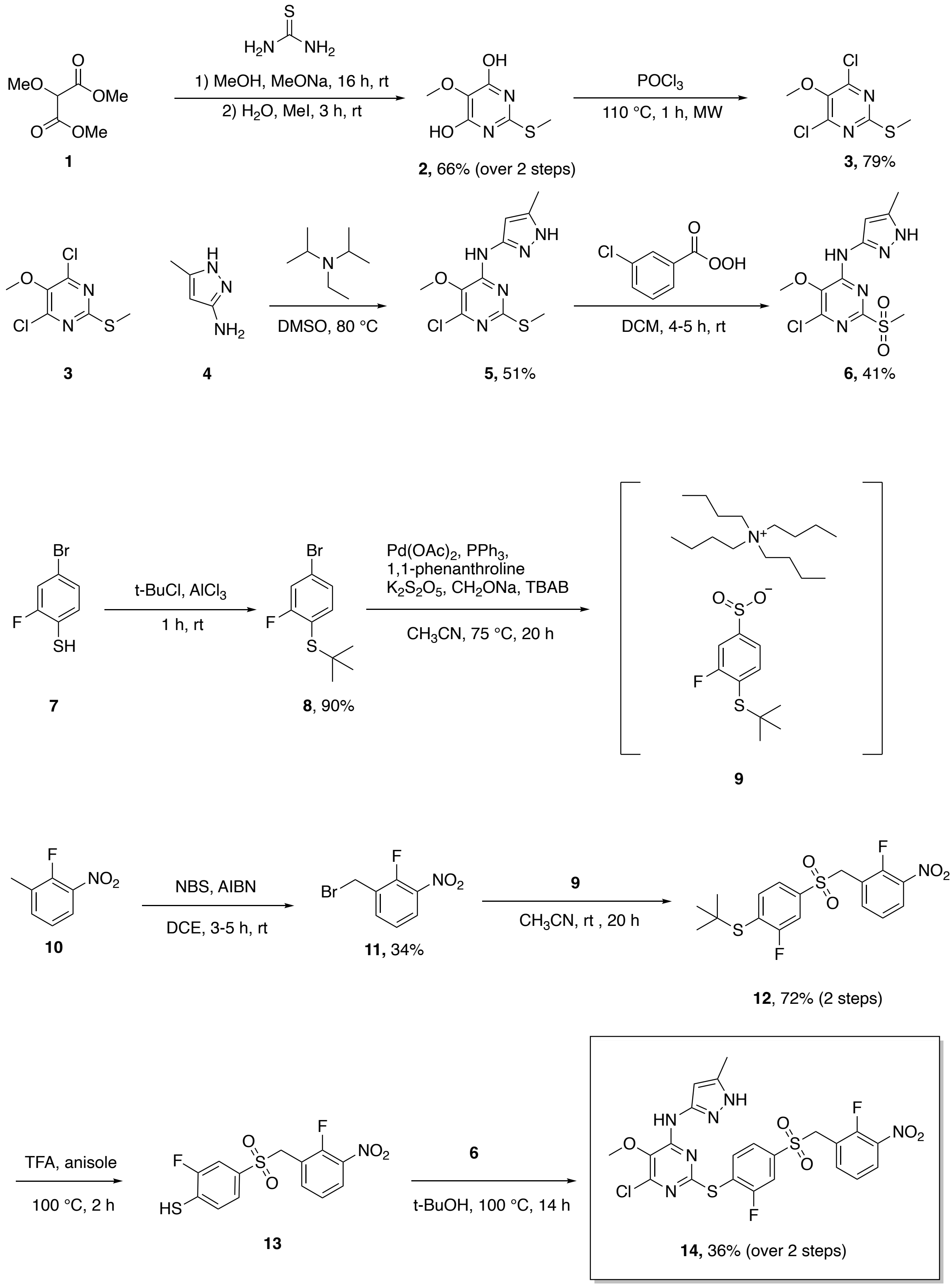


**Supplementary Scheme 1.** Synthesis of centrinone precursor **14**.

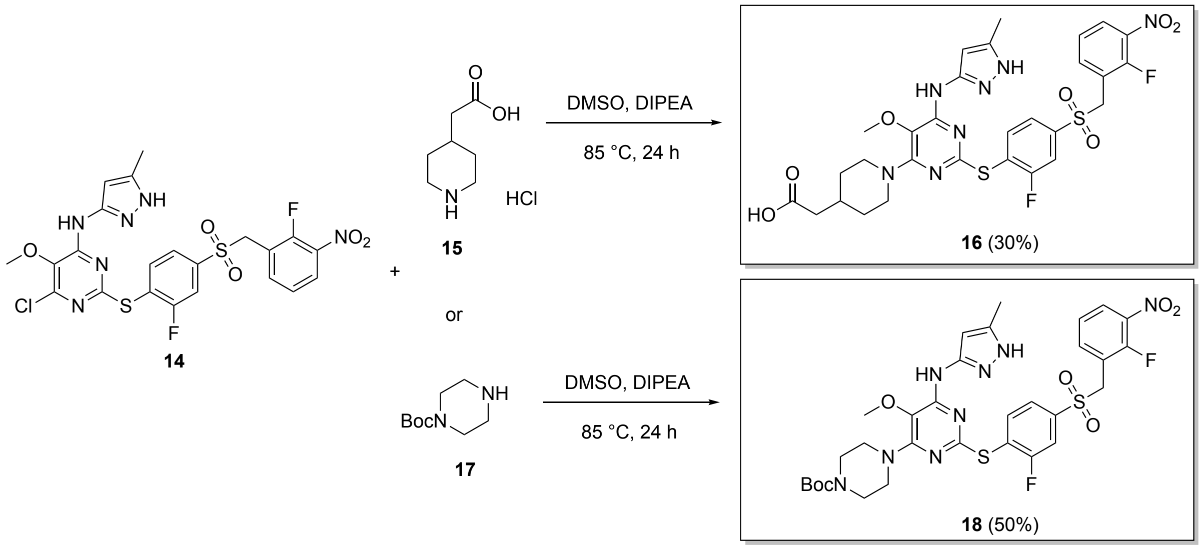


**Supplementary Scheme 2**. Synthesis of functionalized centrinone analogues **16** and **18**.

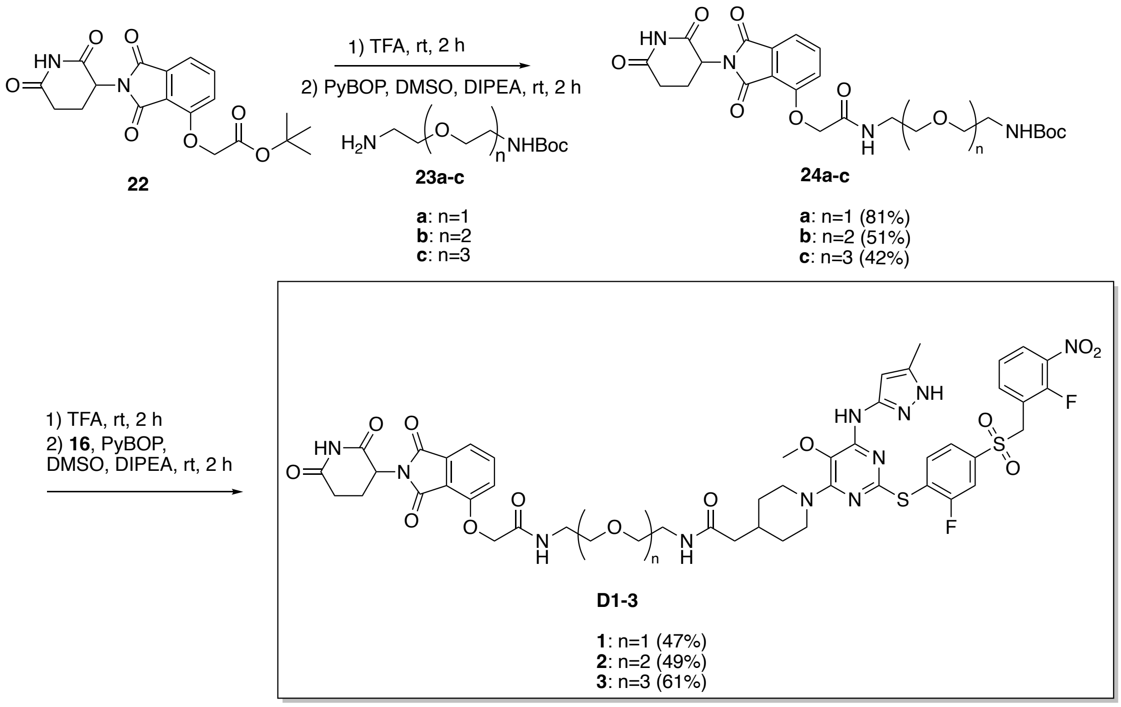


**Supplementary Scheme 3**. Synthesis of **D1-3**.

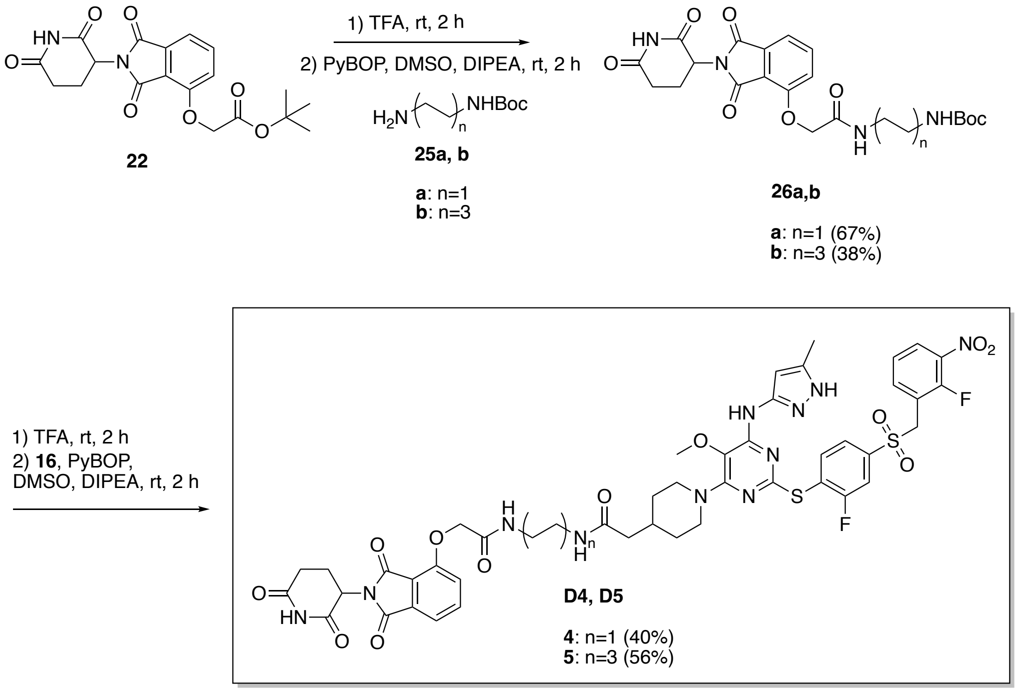


**Supplementary Scheme 4**. Synthesis of **D4** and **D5**.

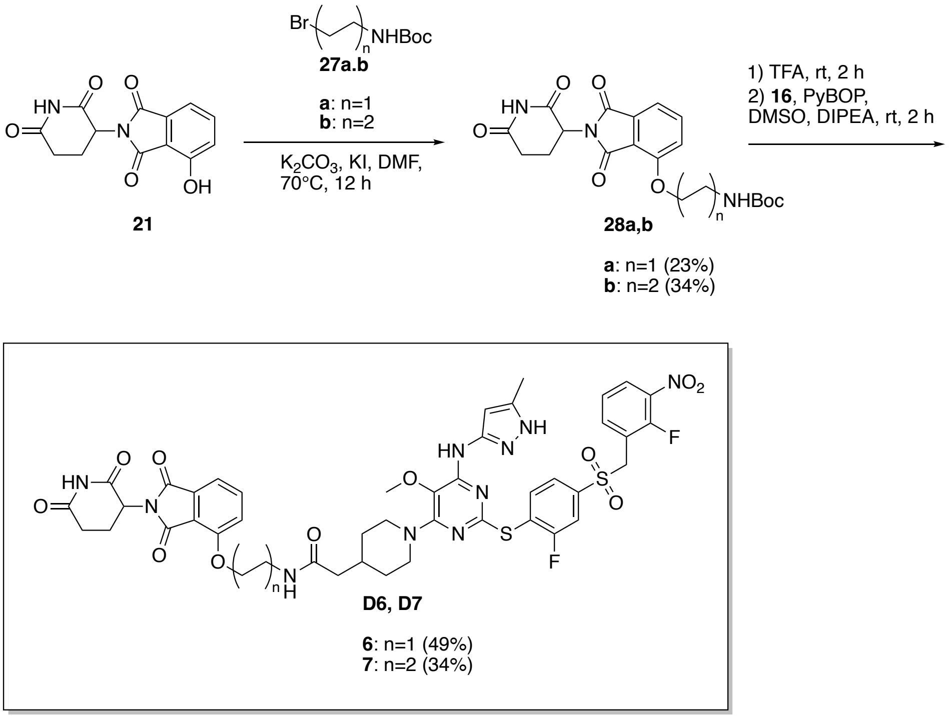


**Supplementary Scheme 5**. Synthesis of **D6** and **D7**.

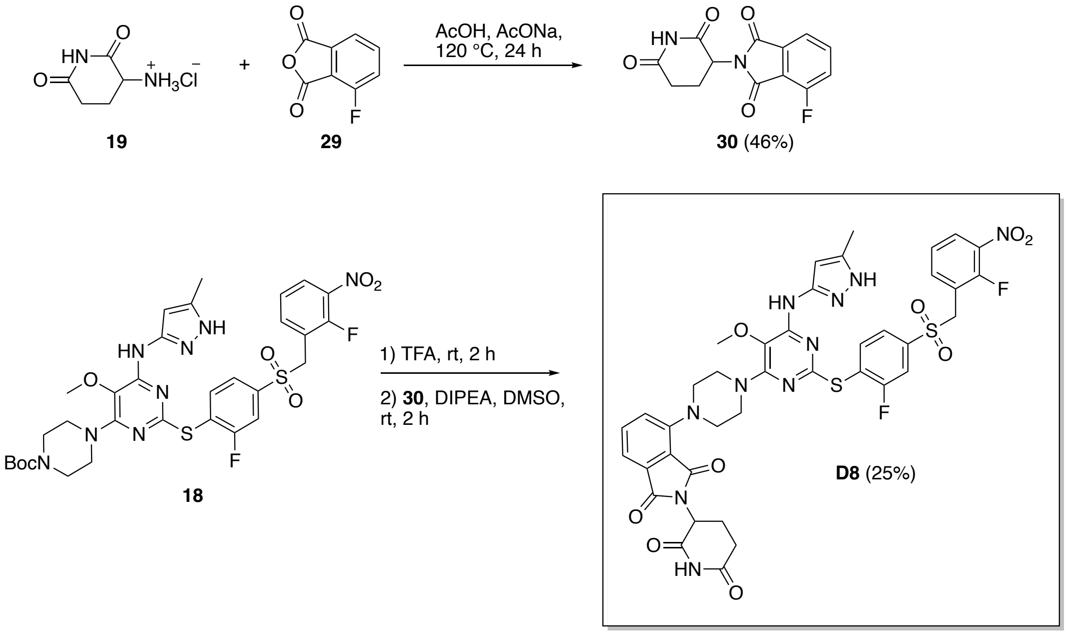


**Supplementary Scheme 6**. Synthesis of **D8**.

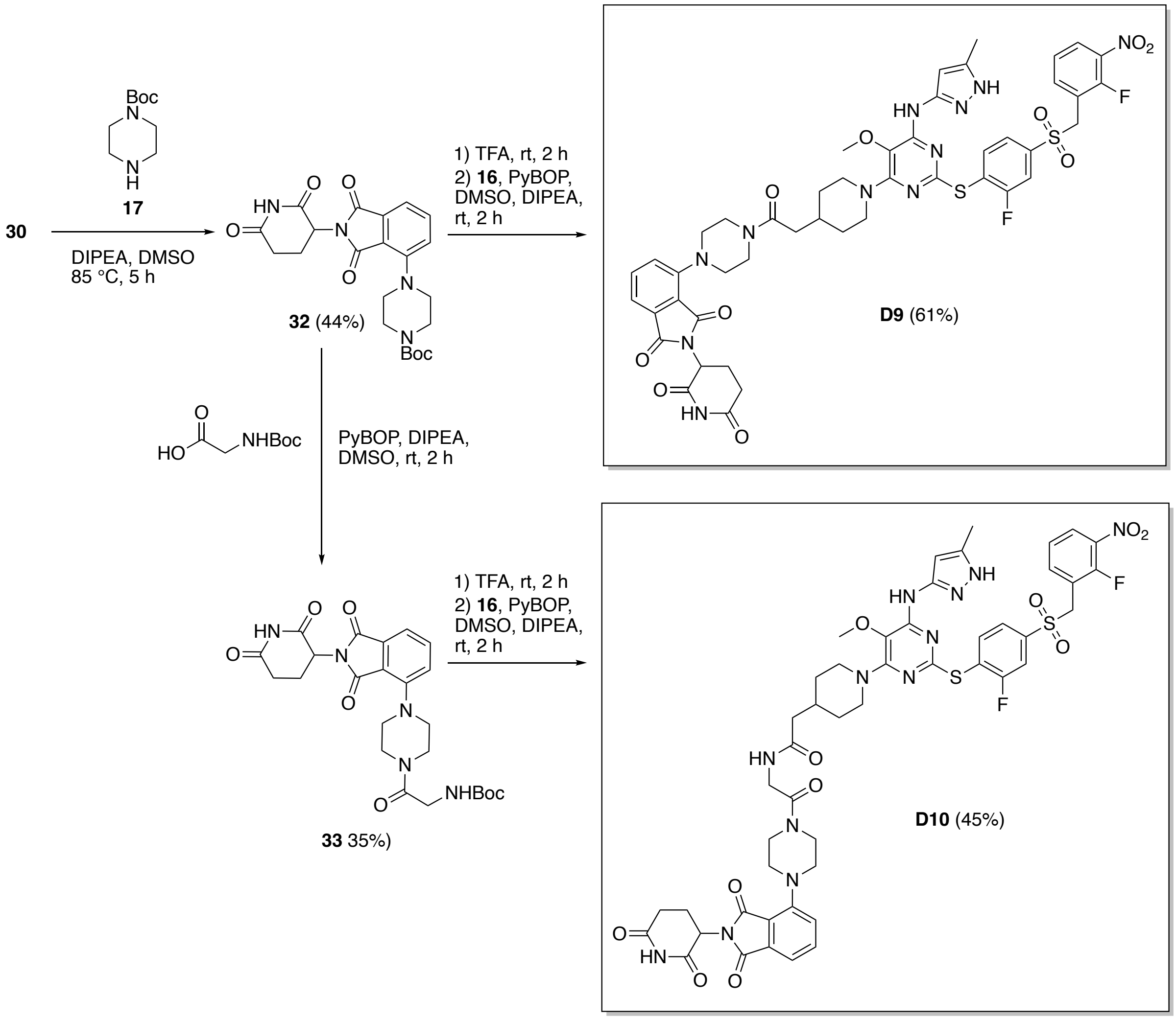


**Supplementary Scheme 7**. Synthesis of **D9** and **D10**.

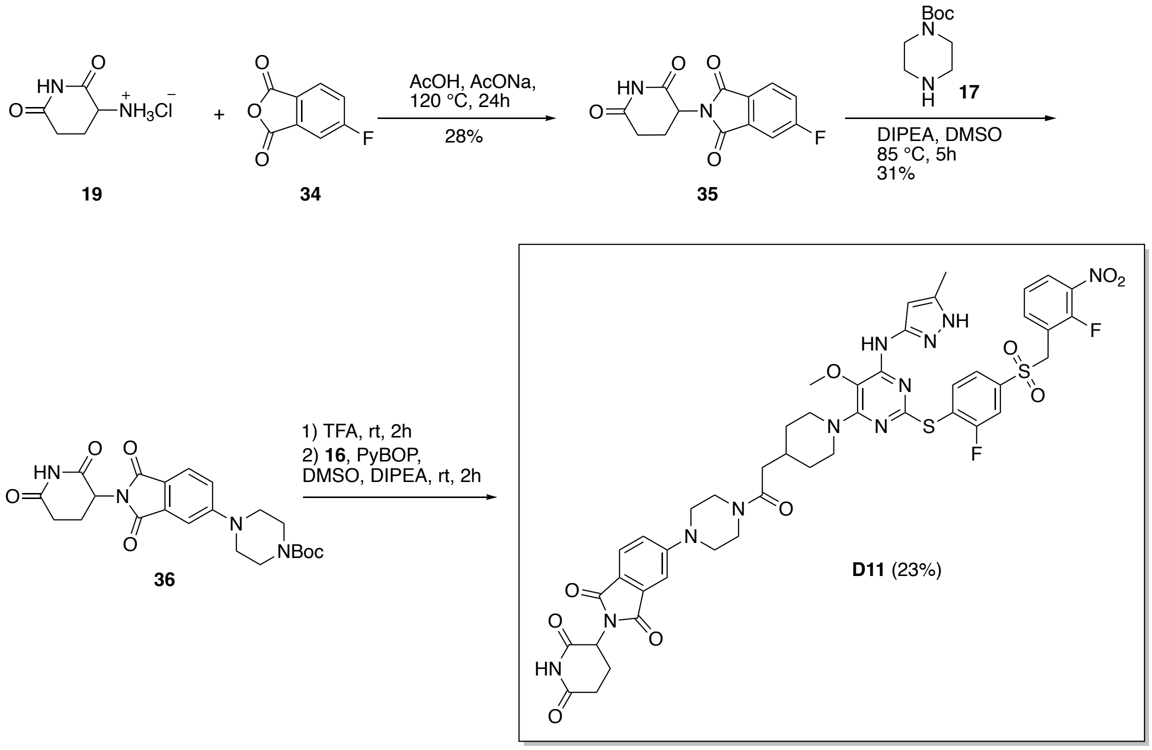


**Supplementary Scheme 8**. Synthesis of **D11**.

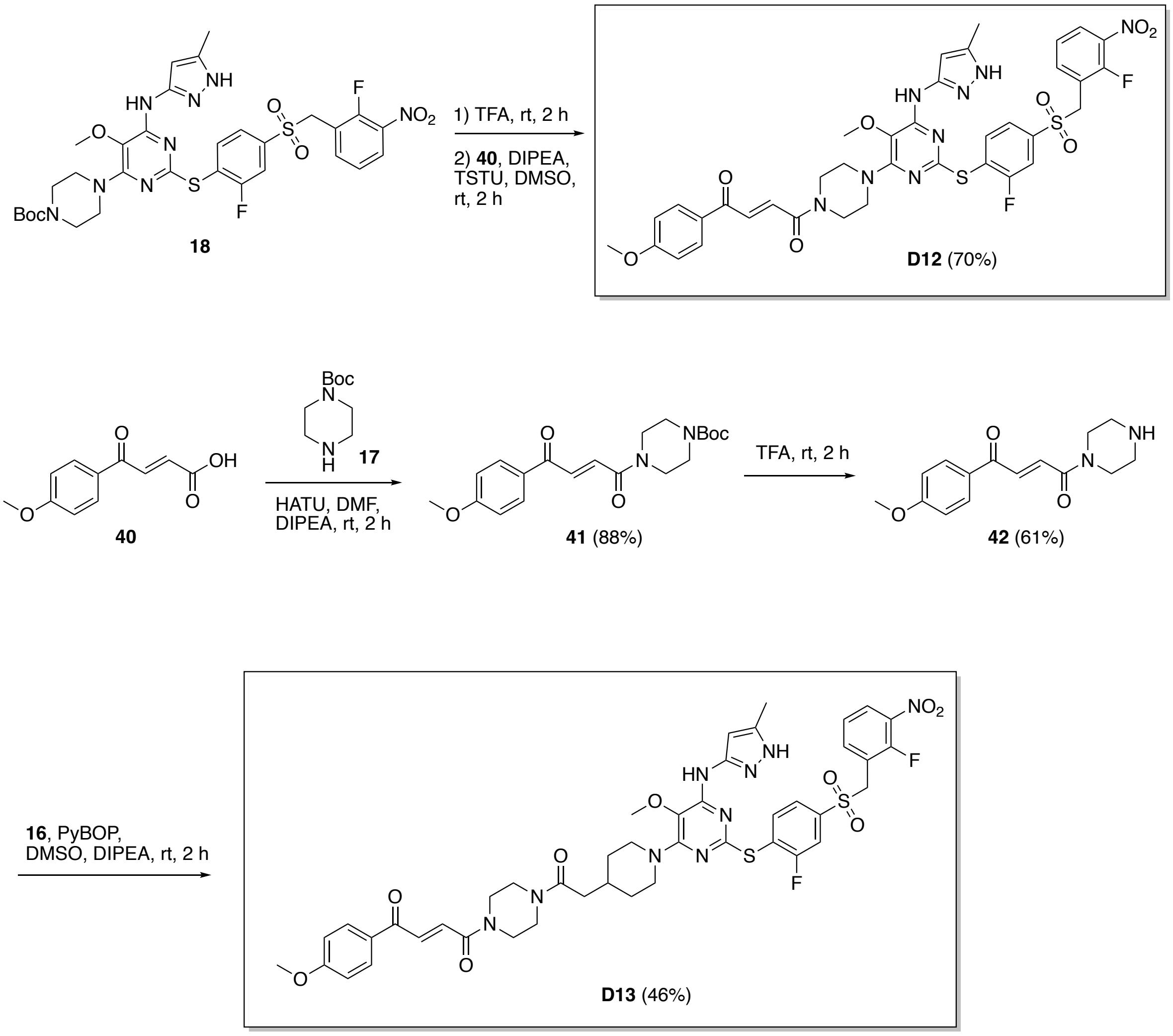


**Supplementary Scheme 9**. Synthesis of **D12** and **D13**.

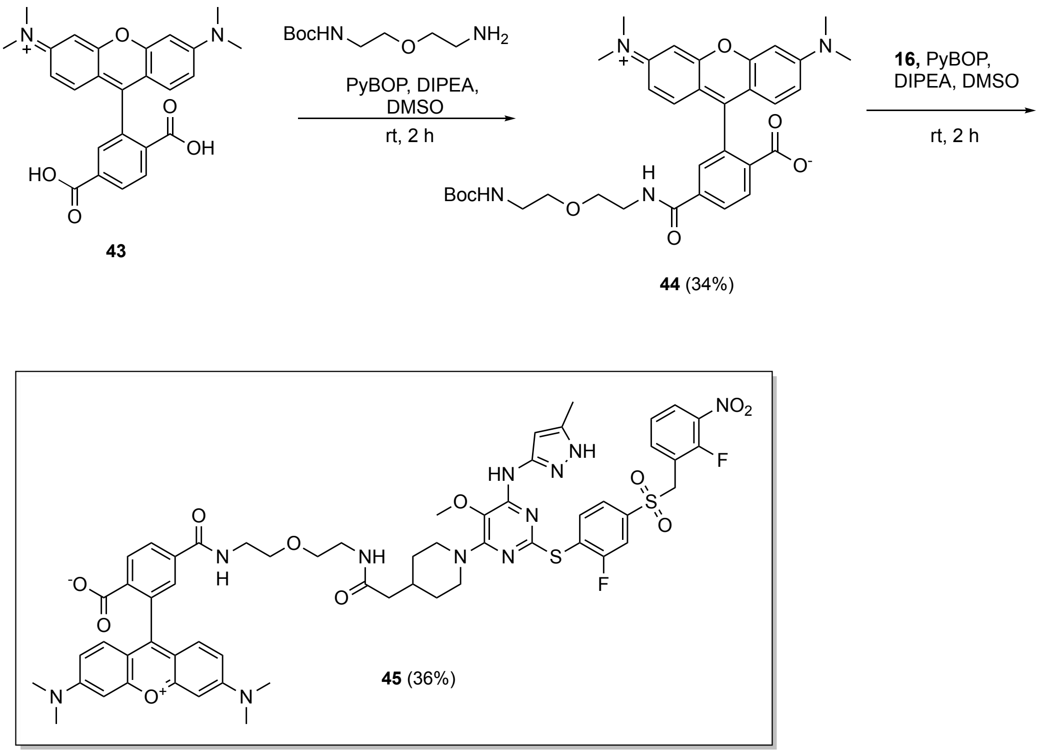


**Supplementary Scheme 10**. Synthesis of centrinone-TMR probe **45**.

#### 3. Biology

##### 3.1 Expression and purification of PLK4

*E. Coli* strain BL21-Gold(DE3)pLysS was transformed with a plasmid encoding the kinase domain of PLK4 (2-275) with a 10xHis-tag in LB medium. Protein expression was induced at 18 °C by the addition of 0.3 mM IPTG and expression was continued for 18 h. After that, the bacterial suspension was centrifuged at 7'000 rpm at 4 °C for 15 min. The bacterial pellet was resuspended in 50 mL of lysis buffer (50 mM Tris (pH 7.5), 400 mM NaCl, 2 mM $MgCl_2$, 5 mM EDTA, 1 mM DTT, 0.5 mM PMSF, 5% glycerol, 0.5% Triton X-100), lysozyme was added and the bacteria were lysed by sonication. The lysate was centrifuged at 4 °C at 15'000 rpm for 30 min and the supernatant was incubated was incubated with Ni-NTA beads for 3 h, with rotation. Prior to this, Ni-NTA beads were spun down and washed twice with wash buffer (20 mM Tris (pH 7.5), 1 mM DTT, 800 mM NaCl, 10% glycerol, 40 mM imidazole). After 3 h, the beads were spun down and they were washed with wash buffer. Beads were loaded onto the column and eluted with elution buffer (20 mM Tris (pH 7.5), 1 mM DTT, 400 mM NaCl, 10% glycerol, 500 mM imidazole).

Buffer was exchanged (20mM Tris (pH 7.5), 1 mM DTT, 200mM NaCl, 10% glycerol) using spin columns and the protein was flash frozen and stored at -80 °C.

Concentration of PLK4 was determined using Pierce assay. Pierce™ Protein Assay Reagent was mixed with the sample at a 15:1 ratio, incubated at room temperature for 5 minutes and absorbance was measured at 660 nm. BSA was used as the standard for the calibration curve.

**3.2 Fluorescence polarization assay**

The solution of 5 nM TMR-centrinone **45** was titrated with increasing concentrations of PLK4-KD dissolved in 20 mM Tris pH 8.0, 200 mM NaCl, 0.05 % 3-[(3-cholamidopropyl)dimethylammonio]-1-propanesulfonate (CHAPS) in a black flat bottom 96-well plate (Greiner Bio-One). The plate was incubated for 3h at room temperature followed by fluorescence polarization measurements. All measurements were done in triplicates. Fluorescence polarization values were determined using a plate reader (TECAN Spark® 20M) using 531 nm as excitation wavelength (bandwidth 20 nm) and 580 nm as emission wavelength (bandwidth 25 nm). The obtained mP values from each well were plotted against the concentration of the protein and the apparent Kd value of TMR-centrinone was obtained by fitting the curve to the following equation[1-2]:

$$FP = FP_{min} + \left(\frac{FP_{max} - FP_{min}}{2c_f}\right)\left(c_f + c_p + K_d - \sqrt{(c_f + c_p + K_d)^2 - 4c_f c_p}\right)$$

where $FP_{min}$ is the minimum polarization value, $FP_{max}$ is the maximum polarization value , $C_f$ is the concentration of TMR-centrinone and $C_p$ is the concentration of the protein (PLK4-KD).

In the competition assay, 5 nM TMR-cen and 10 nM PLK4-KD were used. The solution of the tracer and protein was titrated with increasing amounts of the synthesized probes. Plates were incubated for 3 h at room temperature and fluorescence polarization measurements were performed using the same conditions as before. From the obtained mP values, the concentration of free protein was determined using the following equation[2]:

$$[P_{free}] = K_{d\,(tracer)} \frac{FP_{min} - FP}{FP - FP_{max}}$$

where $[P_{free}]$ is the concentration of the free protein, $K_{d\,(tracer)}$ is the previously determined $K_d$ value of the tracer, $FP_{min}$ is the minimum polarization value and $FP_{max}$ is the maximum polarization value.

The obtained concentration of free protein was plotted against the concentration of the competitor probe and the obtained curve was fitted to a single-site binding isotherm to determine the $K_d$ of the probe:

$$FP = FP_{min} + \frac{FP_{max} - FP_{min}}{1 + \frac{K_{d\,(competitor)}}{[P_{free}]}}$$

##### 3.3 Cell culture

MDA-MB-231 (ATCC HTB-26™), HeLa (ATCC), HeLa Centrin1-GFP (Piel *et al*, 2000)[3] and RPE-1 USP28$^{-/-}$ cell lines (Meitinger *et al*, 2016)[4] were cultured in high-glucose, phenol red DMEM (GlutaMAX) medium containing 10% fetal bovine serum (FBS), 1% sodium pyruvate, 100 U/mL penicillin, and 100 µg/mL streptomycin, in a humidified 5% $CO_2$ incubator at 37 °C. Medium for the acentriolar RPE-1 USP28$^{-/-}$ cell line was supplemented with 300 nM centrinone.[5] Cells were split every 3-4 days or at confluency.

##### 3.4 Transfection with PLK4 (K41M) mutant

Transient transfection of HeLa cells was performed using Lipofectamine™ 2000 reagent (Life Technologies): 15 µg of pEGFP-C3-PLK4 K41M-3xFLAG (Addgene plasmid #69838; http://n2t.net/addgene:69838; RRID:Addgene_69838)[6] was mixed with OptiMEM (500 µL, Life Technologies) and Lipofectamine™ 2000 reagent (30 µL) was mixed with OptiMEM (500 µL). Solutions were then mixed and incubated for 20 minutes at room temperature. The prepared DNA-lipofectamine complex was mixed with 10 mL of growth medium and was added to a 10-cm dish containing cells at 60-90% confluency. The next day, cells were seeded in a 24-well plate for western-blotting (see **3.5 Western blotting**) or a 96-well plate (Ibidi) for microscopy. In all the competition experiments 4-hydroxylthalidomide was used at 50 µM. For microscopy, after 24 h of treatment with compounds at selected concentrations, cells were fixed in methanol at -20 °C for 5 min. After washing once with phosphate-buffered saline (PBS), they were blocked in the blocking solution (1% BSA and 0.1 % Triton X-100 in PBS) for 1 h at room temperature, followed by incubation with the indicated primary antibodies for 16 h at 4 °C. Primary antibodies: anti-Cep152 (rabbit, A302-479A, Bethyl Laboratories) and anti-GFP IgG1 (Developmental Studies Hybridoma Bank, # DSHB-GFP-4C9, 1:100) or anti-PLK4 clone 6H5 (mouse monoclonal, MERCK, MABC544, lot: 3870649, 1:500). After incubation, samples were washed with 0.1 % Triton X-100 in PBS (5 x 1 min) and incubated with secondary antibodies and Hoechst 33342 (1:2’000) for 1 h at room temperature. Secondary antibodies: anti-rabbit AF657 (Alexa Fluor® 647 AffiniPure® Goat Anti-Rabbit IgG (H+L), Jackson Immuno Research, # 111-605-003, 1:500) and anti-mouse AF594 (Alexa Fluor® 594 AffiniPure® Donkey Anti-Mouse IgG (H+L), Jackson Immuno Research, #715-585-150, 1:500) or anti-mouse IgG1 AF594 (Alexa Fluor® 594 AffiniPure® Goat Anti-Mouse IgG, Fcγ subclass 1 specific, #115-585-205, 1:500). Subsequently, cells were washed with the blocking solution (5x1 min), put in PBS and imaged using a 20x water immersion objective and a 40x water immersion objective on a high-content confocal microscope (Molecular Devices™ ImageXpress Micro XL). Image analysis was performed using MetaXpress software (version 6.5.3.247, Molecular Devices, LLC).

##### 3.5 Western blotting

Cells were seeded in 24-well plates and allowed to reattach overnight. The next day, they were treated with compounds at selected concentrations. Acentriolar RPE-1 USP28$^{-/-}$ cells were seeded in the presence of 300 nM centrinone and they were washed with PBS immediately before the compound treatment.[5] After the indicated time of treatment, cells were lysed in SDS sample buffer (50 mM Tris HCl pH 6.8, 8% v/v glycerol, 2% w/v SDS, 100 mM DTT, and 0.1 mg/mL bromophenol blue), boiled, and sonicated. Samples were loaded onto a 4–15% Criterion TGX Stainfree gel (Bio-Rad), and run for 35 min, 210 V in Tris/Glycine/SDS buffer (BioRad). Gels were irradiated for 1 min and stain-free imaged before being transferred onto a PVDF membrane using a Transblot Turbo system (Bio-Rad) and membranes were blocked in 5% milk in 0.1% Tween-20 in Tris-buffered saline (TBST) for 1h. Upon blocking membranes were incubated with a primary antibody

(Anti-PLK-4, clone 6H5 (mouse monoclonal, MERCK, MABC544, lot: 3870649), 1:500) in blocking solution (5% milk in TBST) for 16 h at 4 °C. Membranes were washed with TBST (5 x 5 min), incubated with an HRP-conjugated secondary antibody (Donkey anti-mouse IgG-HRP, Jackson Immuno Research Europe LTD, 1:5’000) in blocking solution (5% milk in TBST) for 1 h at room temperature, washed again and developed using Supersignal West Atto Maximum Sensitivity Substrate (Thermo Fisher) and imaged on Fusion FX geldoc (Vilber). Membranes were stripped using Blue Clear SB for antibody stripping (SERVA) for 20 min and, after 1 h blocking in the blocking solution, incubated with Karyopherin-beta HRP conjugated (Importin beta (IMPβ), Santa Cruz, sc-137016, H7-HRP lot E1519, 1:500) for 16 h at 4 °C or for 2 h at room temperature. After washing, membranes were developed using Supersignal West Pico Maximum Sensitivity Substrate (Thermo Fisher) and imaged on Fusion FX geldoc (Vilber). Band intensities were determined using Fiji Image J2 (National Institute of Health) on background subtracted images and normalized to total protein loaded using Importin beta (Karyopherin-beta HRP conjugated (Importin beta (IMPβ), Santa Cruz, sc-137016, H7-HRP lot E1519), 1:500) or stainfree image as the loading control.

**3.6 Centriole phenotypic scoring**

HeLa Centrin1-GFP cells were seeded in a 96-well plate (10’000 cells/well) and allowed to reattach overnight. The next day, they were treated with selected compounds and after 24 h, they were fixed in methanol at -20 °C for 5 min. For the competition experiments, 4-hydroxythalidomide was used at 50 µM for the degrader concentrations >1 µM and at 10 µM for the degrader concentrations ≤1 µM. After washing once with phosphate-buffered saline (PBS), they were blocked in the blocking solution (1% BSA and 0.1 % Triton X-100 in PBS) for 1 h at room temperature, followed by incubation with the indicated primary antibodies for 16 h at 4 °C. The primary antibody used was anti-GFP IgG1 (Developmental Studies Hybridoma Bank, # DSHB-GFP-4C9, 1:100). After incubation, samples were washed with 0.1 % Triton X-100 in PBS (5 x 1 min) and incubated with the secondary antibody (FITC goat anti-mouse IgG Fcγ subclass 1 specific Jackson Immuno Research, # 115-095-205, 1:500) and Hoechst 33342 (1: 2’000) for 1 h at room temperature. Subsequently, cells were washed with the blocking solution (5 x 1 min), put in PBS and centriole foci were scored on DMi8 inverted microscope (Leica Microsystems,

Wetzlar, Germany) equipped with Evolve 512 Delta EMCCD camera (Photometrics, Tucson, AZ, USA).

#### 4. Chemistry

##### 4.1 General

All chemicals, reagents, and anhydrous solvents were purchased from commercial suppliers (Acros, Apollo, Fluka, Fluorochem, Merck, Sigma-Aldrich and TCI). Column chromatography was carried out on silica gel 60 (SilicaFlash P60, 40-63 μm). Analytical (TLC) and preparative thin layer chromatography (PTLC) were performed on silica gel 60 (Merck, 0.2 mm) and silica gel GF (SiliCycle, 1 or 0.25 mm), respectively. LC-MS was recorded using a Thermo Scientific Accela HPLC equipped with a Thermo C18 Hypersil GOLD column (50 x 2.1 mm, 1.9 μm particle size) coupled with an LCQ Fleet three-dimensional ion trap mass spectrometer (ESI, Thermo Scientific). Linear elution gradients were used from 95:5 to 10:90 $H_2O/CH_3CN$ + 0.1% TFA in 4.0 minutes at a flow rate of 0.75 ml/min (B5), 70:30 to 10:90 $H_2O/CH_3CN$ + 0.1% TFA in 4.0 minutes at a flow rate of 0.75 ml/min (B30) or 40:60 to 10:90 $H_2O/CH_3CN$ + 0.1% TFA in 4.0 minutes at a flow rate of 0.75 ml/min (B60). Reverse phase HPLC purification was performed using an Agilent Technologies 1260 infinity HPLC equipped with a BSE1ICO-2520 Scorpius-C18e-HP column, 100 Å, 5 μm, 21.2 × 250 mm.

All $^{1}H$ and $^{13}C$ NMR spectra were recorded on a Bruker 300 MHz, 400 MHz, or 500 MHz spectrometer at r.t. in $CDCl_3$, DMSO-d6 and $CD_3OD$ and are reported as chemical shifts (δ) in ppm relative to TMS (δ = 0). Spin multiplicities are reported as a singlet (s), doublet (d), triplet (t), quartet (q), quintet (p), sextet (h) or multiplet (m), with coupling constants (J) given in Hz.

##### 4.2 Synthetic procedures

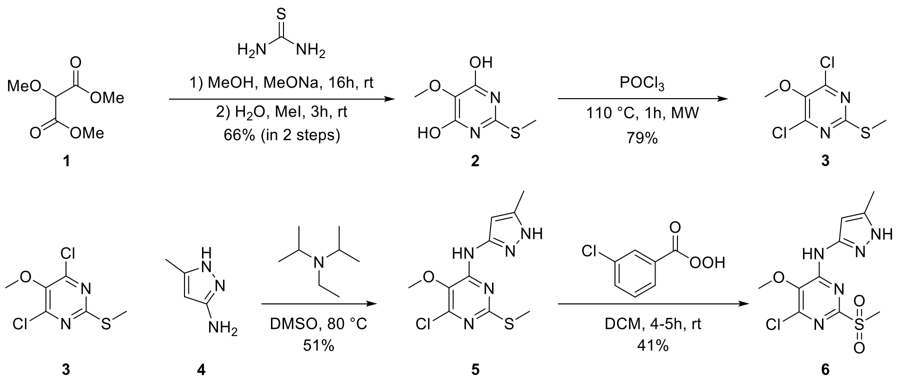


#### 4,6-dihydroxy-5-methoxythiopyrimidine **2**[7]

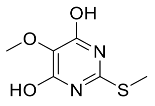

**2**

Dimethyl-malonate **1** (4.07 g, 25.1 mmol, 1.0 eq.) and thiourea (2.87 g, 37.7 mmol, 1.5 eq.) were mixed in MeOH (60 mL) and 25% sodium methoxide (13.6 g, 14.5 mL, 62.8 mmol, 2.5 eq.) solution in MeOH was added dropwise over 15 minutes at 0 °C. The reaction mixture was stirred for 16h at room temperature. Afterwards, MeOH was evaporated and the residue was dissolved in water (40 mL), followed by addition of methyl-iodide (5.34 g, 2.40 mL, 37.7 mmol, 1.5 eq.), upon which the mixture was stirred at room temperature for 3h. The reaction mixture was acidified with a HCl solution (6M) to pH 3-4, which led to precipitation of the product. The precipitate was collected by filtration and the mother liquor was acidified 3 times to allow the complete precipitation of the product. The collected precipitate was washed with water and dried on air to obtain the desired product **2** as a white powder (3.10 g, 66%).

**$^1$H NMR** (400 MHz, MeOD) δ 3.71 (s, 3H), 2.52 (s, 3H).

**$^{13}$C NMR** (75 MHz, MeOD) δ 161.55, 156.01, 123.04, 58.94, 11.78.

#### 4,6-dichloro-5-methoxythiopyirimidine **3**[7]

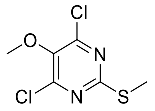

**3**

Product **2** (1.00 g, 5.31 mmol, 1.0 eq) was slowly added into freshly distilled $POCl_3$ (9.84 g, 6 mL, 64.2 mmol, 12.08 eq) in a microwave vessel. The reaction mixture was stirred at

110 °C for 75 minutes at 150 W in a microwave. Afterwards, the mixture was cooled to -20 °C and carefully quenched with cold water. The product was extracted with EtOAc (3 x 20 mL) and combined organic layers were washed with brine, dried over $MgSO_4$ and evaporated. The crude residue was dissolved in $CHCl_3$ and separated by flash column chromatography (pentane/DCM; gradient 0-50%) to obtain the desired product **3** as a beige powder (940 mg, 79%).

**$^1$H NMR** (300 MHz, $CDCl_3$) δ 3.92 (s, 3H), 2.56 (s, 3H).

**$^{13}$C NMR** (75 MHz, $CDCl_3$) δ 166.86, 155.15, 143.44, 61.24, 14.83

6–chloro-5-methoxy-4-((methyl-1*H*-pyrazol-3-yl)amino)-2-(methylthio)pyrimidine **5**[7]

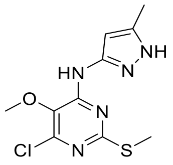


**5**

Compound **3** (430 mg, 1.91 mmol, 1.0 eq) was dissolved in DMSO and 5-methyl-2H-pyrazole-3-yl amine **4** (278 mg, 2.87 mmol, 1.5 eq) and *N,N*-diisopropylethylamine (DIPEA; 741 mg, 1 mL, 3 eq) were added. The reaction mixture was stirred at 80 °C for 5 h. Upon reaction completion, it was diluted by water and the product was extracted with EtOAc (3x20 mL). Combined organic extracts were washed with water and brine, dried over $MgSO_4$ and evaporated. The crude residue was dissolved in EtOAc and purified by flash column chromatography (pentane/EtOAc; gradient 0-100 %) to obtain the desired product **5** as a brown powder (280 mg, 51%).

**$^1$H NMR** (400 MHz, DMSO) δ 12.15 (s, 1H), 9.60 (s, 1H), 6.38 (s, 1H), 3.75 (s, 3H), 2.45 (s, 3H), 2.23 (s, 3H).

**$^{13}$C NMR** (101 MHz, DMSO) δ 164.48, 154.36, 148.33, 146.66, 138.30, 132.16, 97.12, 60.60, 14.10, 10.75.

##### 6-chloro-5-methoxy-4-((5-methyl-1*H*-pyrazol-3-yl)amino)-2-(methylsulfonyl) pyrimidine **6**[7]

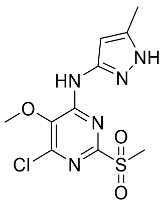


**6**

To a stirred solution of **5** (280 mg, 0.98 mmol, 1.0 eq) in DCM at 0 °C, a solution of mCPBA (77% wt) (505 mg, 2.25 mmol, 2.30 eq) in DCM was slowly added. The reaction mixture was stirred for 4h at room temperature. Upon reaction completion, the mixture was diluted with DCM and washed with 1:1 solution of $Na_2S_2O_3$ and $NaHCO_3$ (3 x 50 mL). The organic phase was washed with water and brine and dried over $MgSO_4$. Prior to evaporation, silica gel was added. After evaporation, the crude was dry-loaded onto the column and purified by flash column chromatography (pentane/EtOAc; gradient 50-100%) to obtain the desired product **6** as a white powder (132 mg, 41%).

**$^1$H NMR** (300 MHz, DMSO) δ 12.28 (s, 1H), 10.35 (s, 1H), 6.46 (s, 1H), 3.87 (s, 3H), 3.31 (s, 3H), 2.25 (s, 3H).

**$^{13}$C NMR** (101 MHz, DMSO) δ 158.44, 158.08, 155.08, 148.08, 145.96, 138.91, 137.22, 96.98, 60.78, 10.87

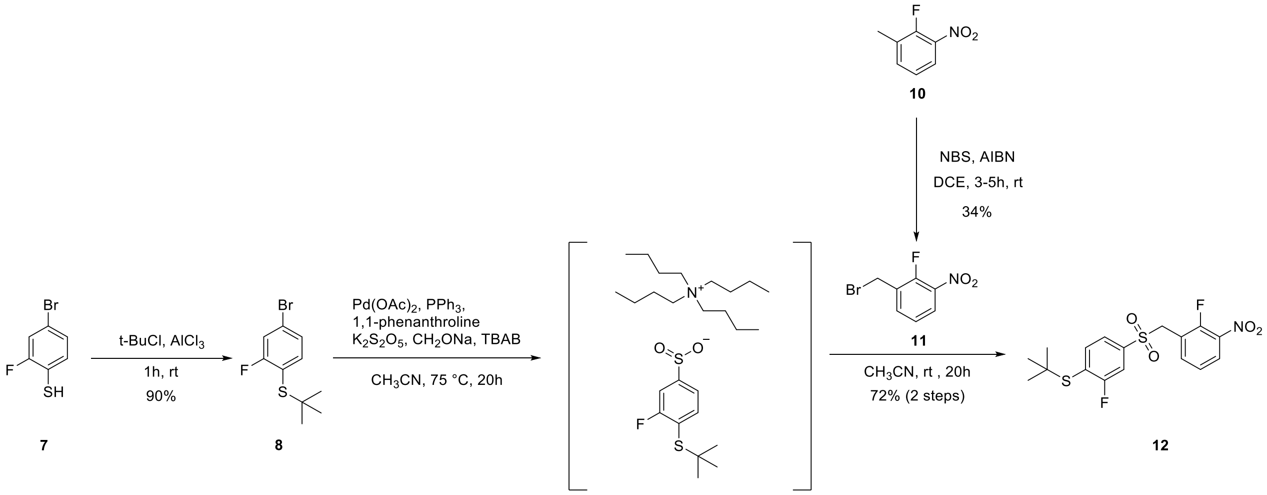

##### 4-bromo-2-fluoro-1-(*tert*-butylthio)benzene **8**[7]

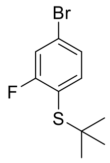

**8**

$AlCl_3$ (65 mg, 0.49 mmol, 0.2 eq.) was added in small portions to the ice-cold solution of thiophenol **7** (500 mg, 2.41 mmol, 1 eq.) in *tert*-butylchloride (5 mL). Evolved HCl was led through a saturated solution of $NaHCO_3$. Upon addition of $AlCl_3$, the ice-bath was removed, and the reaction was stirred at room temperature for 2 h. Afterwards, the reaction mixture was diluted with water and the product was extracted with pentane (3 x 50 mL). The combined organic phases were washed with brine, dried over $MgSO_4$ and filtered. Prior to evaporation, silica gel was added. After evaporation, silica gel loaded with compound was purified by flash column chromatography (pentane; isocratic) to obtain the desired product **8** as a colourless oil (570 mg, 90%).

**$^1$H NMR** (300 MHz, $CDCl_3$) δ 7.39 (t, *J* = 7.9 Hz, 1H), 7.31 (dd, *J* = 7.9, 1.9 Hz, 1H), 7.29 – 7.25 (m, 1H), 1.29 (s, 9H).

**$^{13}$C NMR** (75 MHz, $CDCl_3$) δ 164.08 (d, *J* = 250.7 Hz), 141.13, 127.61 (d, *J* = 4.1 Hz), 124.13 (d, *J* = 8.8 Hz), 119.78 (d, *J* = 27.9 Hz), 119.23 (d, *J* = 19.1 Hz), 47.67, 31.04.

##### 1-(bromomethyl)-2-fluoro-3-nitrobenzene **11**[7]

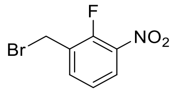

**11**

1-methyl-2-fluoro-3-nitrobenzene **10** (2.00 g, 12.89 mmol, 1 eq.) and *N*-bromosuccinimide (NBS; 2.75 g, 15.47 mmol, 1.2 eq.) were dissolved in 1,2-dichloroethane (100 mL) and to this stirring solution azobisisobutyronitrile (AIBN; 423 mg, 2.58 mmol, 0.2 eq.) was added. The mixture was stirred under reflux for 6 h. After cooling to room temperature, the reaction mixture was diluted with DCM and washed with water, saturated $NaHCO_3$ and brine. Combined organic phases were dried over anh. $MgSO_4$ and evaporated. Crude product was dissolved in pentane/DCM and purified by flash column chromatography (pentane/DCM; gradient 0-50%) to obtain the desired product **11** as a yellow oil (1.03 g, 34%).

$^{1}$**H NMR** (300 MHz, $CDCl_3$) δ 8.02 (ddd, *J* = 8.5, 7.0, 1.8 Hz, 1H), 7.72 (ddd, *J* = 8.0, 6.3, 1.8 Hz, 1H), 7.34 – 7.28 (m, 1H), 4.56 (d, *J* = 1.6 Hz, 2H).

$^{13}$**C NMR** (101 MHz, $CDCl_3$) δ 153.65 (d, *J* = 267.5 Hz), 138.07, 136.62 (d, *J* = 4.0 Hz), 128.78 (d, *J* = 13.9 Hz), 126.39 (d, *J* = 2.5 Hz), 124.63 (d, *J* = 5.0 Hz), 23.70 (d, *J* = 5.4 Hz).

*tert*-butyl (2-fluoro-4-((2-fluoro-3-nitrobenzyl)sulfonyl)phenyl)sulfane **12**

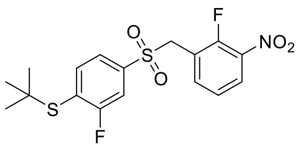


**12**

A glass tube was charged with **8** (230 mg, 0.87 mmol, 1.0 eq), potassium metabisulfite (486 mg, 2.18 mmol, 2.50 eq), tetrabutylammonium bromide (338 mg, 1.05 mmol, 1.20 eq), sodium formate (149 mg, 2.18 mmol, 2.50 eq), palladium (II)-acetate ( 20 mg, 0.09 mmol, 0.10 eq), triphenylphosphine (69 mg, 0.26 mmol, 0.30 eq), 1,10-phenanthroline (47 mg, 0.26 mmol, 0.30 eq) and acetonitrile (5 mL). The mixture was degassed with by bubbling nitrogen gas under stirring for 15 minutes and then stirred at 70 °C for 20 h. Afterwards, the reaction mixture was cooled down to room temperature and **11** (245 mg, 1.05 mmol, 1.20 eq), dissolved in acetonitrile (5 mL), was added. The reaction was stirred at room temperature for 20 h. After that, it was diluted with water and the product was extracted with EtOAc (3 x 20 mL). The combined organic layers were dried over anh. $MgSO_4$, filtered, and evaporated. Crude was dissolved in $CHCl_3$ and purified by flash column chromatography (EtOAc/pentane gradient 0-50%) to obtain the desired product **12** as a pale yellow solid (254 mg, 72%).

$^{1}$**H NMR** (400 MHz, $CDCl_3$) δ 8.06 (ddd, *J* = 8.6, 7.0, 1.8 Hz, 1H), 7.75 (ddd, *J* = 7.7, 6.0, 1.8 Hz, 1H), 7.68 (dd, *J* = 8.0, 6.7 Hz, 1H), 7.47 (dd, *J* = 7.3, 1.9 Hz, 1H), 7.43 (dd, *J* = 7.9, 2.0 Hz, 1H), 7.36 (td, *J* = 8.0, 1.3 Hz, 1H), 4.49 (s, 2H), 1.32 (s, 9H).

$^{13}$**C NMR** (101 MHz, $CDCl_3$) δ 163.58 (d, *J* = 252.8 Hz), 153.82 (d, *J* = 267.0 Hz), 140.54, 139.55 (d, *J* = 6.5 Hz), 138.04 (d, *J* = 3.4 Hz), 129.01 (d, *J* = 19.3 Hz), 127.16 (d, *J* = 2.4 Hz), 124.68 (d, *J* = 5.3 Hz), 123.58 (d, *J* = 4.4 Hz), 119.01 (d, *J* = 14.1 Hz), 115.94 (d, *J* = 28.2 Hz), 55.07 (d, *J* = 2.9 Hz), 48.92, 31.00.

**HRMS (ESI):** calc. for $C_{17}H_{17}F_2NO_4S_2$ $[M+Na]^+$ 424.0465; found 424.0459

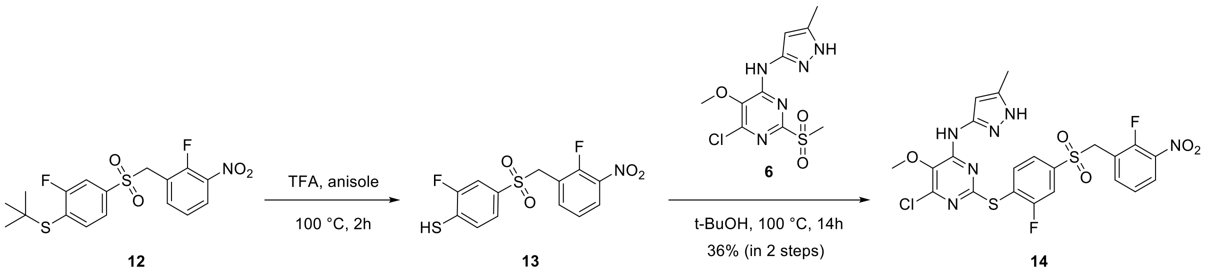


6-chloro-2-((2-fluoro-4-((2-fluoro-3-nitrobenzyl)sulfonyl)phenyl)thio)-5-methoxy-4-((5-methyl-1*H*-pyrazol-3-yl)amino)pyrimidine **14**[7]

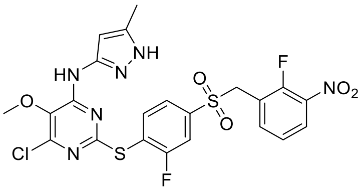

**14**

Sulfone **12** (123 mg, 0.31 mmol, 1.3 eq.) was dissolved in pure trifluoroacetic acid (TFA, 5 mL) and anisole (66 mg, 0.62 mol, 2.6 eq.) was added. The reaction flask was sealed, atmosphere exchanged with argon multiple times and the reaction mixture was stirred at 100 °C for 2 h. Upon reaction completion, the mixture was cooled down to room temperature and TFA was evaporated. The crude containing compound **13** was kept under argon, before adding methyl-sulfone **6** (75 mg, 0.24 mmol, 1.0 eq.) and *tert*-butanol (10 mL). The flask was sealed, air exchanged with argon and reaction was stirred at 100 °C overnight. Upon completion, the solvent was evaporated and the crude purified via RP-HPLC (ACN/$H_2O$ gradient; 10-90%, 60 min) to afford the centrinone precursor **14** as a pale yellow TFA salt (50 mg, 36%).

**$^1$H NMR** (400 MHz, DMSO) δ 9.80 (s, 1H), 8.17 (ddd, *J* = 8.6, 7.1, 1.8 Hz, 1H), 7.99 (dd, *J* = 8.1, 6.7 Hz, 1H), 7.84 (dd, *J* = 8.1, 1.9 Hz, 1H), 7.67 (dd, *J* = 8.1, 2.0 Hz, 1H), 7.61 (ddd, *J* = 7.9, 6.1, 1.8 Hz, 1H), 7.44 (t, *J* = 8.0 Hz, 1H), 5.44 (s, 1H), 5.02 (s, 2H), 3.75 (s, 3H), 2.04 (s, 3H).

**$^{13}$C NMR** (101 MHz, DMSO) δ 161.63 (d, *J* = 252.4 Hz), 161.16, 154.47, 153.47 (d, *J* = 265.9 Hz), 148.41, 145.78, 141.06 (d, *J* = 6.7 Hz), 139.03 (d, *J* = 3.8 Hz), 138.87, 138.04, 137.33 (d, *J* = 8.0 Hz), 133.33, 127.02 (d, *J* = 2.2 Hz), 124.94 (d, *J* = 4.8 Hz), 124.80 (d, *J* = 3.9 Hz), 124.22 (d, *J* = 18.4 Hz), 118.87 (d, *J* = 14.2 Hz), 115.97 (d, *J* = 26.4 Hz), 96.00, 60.69, 54.10, 10.80.

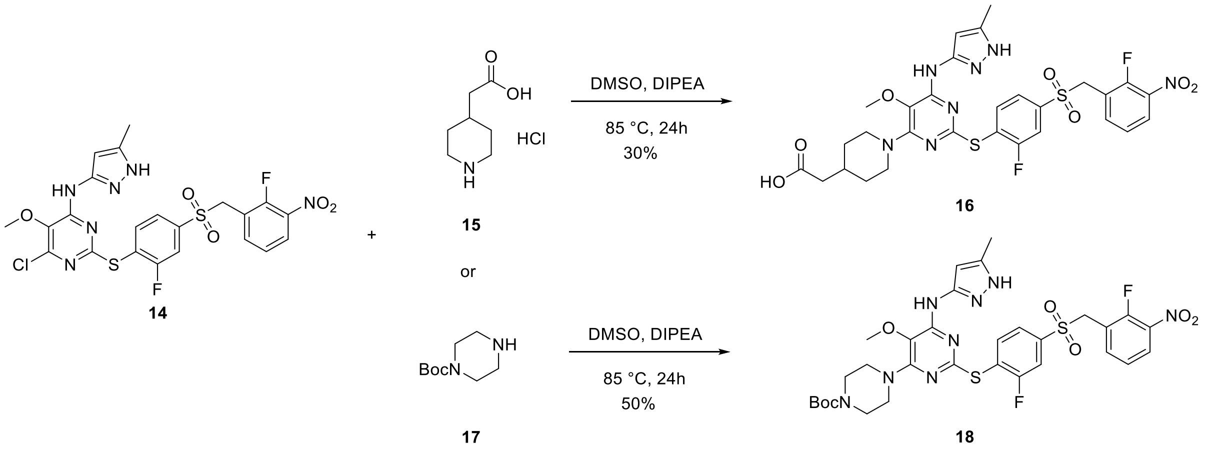


2-(1-(2-((2-fluoro-4-((2-fluoro-3-nitrobenzyl)sulfonyl)phenyl)thio)-5-methoxy-6-((5-methyl-1*H*-pyrazol-3-yl)amino)pyrimidin-4-yl)piperidin-4-yl) acetic acid **16**[7]

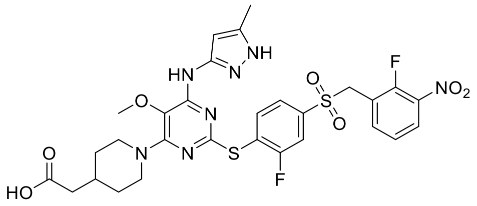

**16**

Centrinone precursor **14** (20 mg, 0.034 mmol, 1.0 eq.) was dissolved in anh. DMSO (0.5 mL) and 2-(piperidin-4-yl)acetic acid hydrochloride **15** (30.8 mg, 0.17 mmol, 5.0 eq.) and DIPEA (44 mg, 60 µL, 0.34 mmol, 10 eq.) were added. The reaction mixture was stirred at 85 °C overnight. After cooling down to room temperature, the mixture was acidified, diluted with water and purified by RP-HPLC (ACN/$H_2O$ gradient; 10-90%, 60 min). The corresponding fractions were collected, frozen and lyophilized overnight to obtain the desired product **16** as a beige powder (7 mg, 30%).

**$^1$H NMR** (400 MHz, DMSO) δ 8.94 (s, 1H), 8.17 (ddd, *J* = 8.6, 7.1, 1.8 Hz, 1H), 7.96 (dd, *J* = 8.1, 6.8 Hz, 1H), 7.81 (dd, *J* = 8.1, 1.9 Hz, 1H), 7.68 – 7.61 (m, 0H), 7.65 (d, *J* = 1.9 Hz, 1H), 7.63 (d, *J* = 2.0 Hz, 1H), 7.45 (t, *J* = 8.0 Hz, 1H), 5.64 (s, 1H), 5.01 (s, 2H), 4.18 (d, *J* = 13.0 Hz, 6H), 3.54 (s, 3H), 2.81 (td, *J* = 12.9, 2.4 Hz, 2H), 2.15 (d, *J* = 6.9 Hz, 2H), 2.09 (s, 3H), 1.88 (ddd, *J* = 11.1, 7.2, 3.8 Hz, 1H), 1.70 – 1.61 (m, 2H), 1.14 (qd, *J* = 12.5, 3.8 Hz, 2H).

**$^{13}$C NMR** (101 MHz, DMSO) δ 173.40, 161.62 (d, *J* = 251.3 Hz), 159.56, 153.66, 153.45 (d, *J* = 265.6 Hz), 153.23, 145.90, 140.64 (d, *J* = 6.7 Hz), 140.41, 139.05 (d, *J* = 3.7 Hz), 138.03, 137.32 (d, *J* = 7.9 Hz), 127.01, 125.46 (d, *J* = 18.4 Hz), 124.95 (d, *J* = 4.7 Hz), 124.45, 122.15,

118.91 (d, *J* = 14.4 Hz), 115.61 (d, *J* = 26.6 Hz), 95.14, 58.66, 54.04, 45.61, 40.50, 32.58, 31.48, 11.04.

*tert*-butyl-4-(2-((2-fluoro-4-((2-fluoro-3-nitrobenzyl) sulfonyl) phenyl) thio)-5-methoxy-6-((5-methyl-1*H*-pyrazol-3-yl) amino) pyrimidin-4-yl)piperazine-1-carboxylate **18**[7]

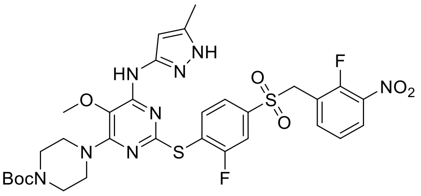


**18**

Centrinone precursor **14** (20 mg, 0.034 mmol, 1.0 eq.) was dissolved in anh. DMSO (0.5 mL) and tert-butyl piperazine-1-carboxylate **17** (32 mg, 0.17 mmol, 5.0 eq) and DIPEA (44 mg, 60 µL, 0.34 mmol, 10 eq.) were added. The reaction mixture was stirred at 85 °C overnight. After cooling down to room temperature, the mixture was acidified, diluted with water and purified by RP-HPLC (ACN/$H_2O$ gradient; 10-90%, 60 min). The corresponding fractions were collected, frozen and lyophilized overnight to obtain the desired product **18** as a yellow powder (12.3 mg, 50%).

**$^1$H NMR** (400 MHz, DMSO) δ 8.92 (d, *J* = 12.7 Hz, 1H), 8.17 (t, *J* = 7.7 Hz, 1H), 7.96 (t, *J* = 7.5 Hz, 1H), 7.83 (d, *J* = 10.0 Hz, 1H), 7.65 (d, *J* = 7.9 Hz, 2H), 7.46 (t, *J* = 8.0 Hz, 1H), 5.61 (d, *J* = 3.8 Hz, 1H), 5.02 (s, 2H), 3.56 (s, 3H), 3.45 (s, 4H), 3.35 (s, 4H), 2.08 (s, 3H), 1.40 (s, 9H).

**$^{13}$C NMR** (126 MHz, DMSO) δ 161.62 (d, *J* = 251.2 Hz), 159.67, 153.88, 153.62 (d, *J* = 3.1 Hz), 153.43 (d, *J* = 266.1 Hz), 146.20, 140.68 (d, *J* = 6.8 Hz), 139.64, 139.06 (d, *J* = 3.7 Hz), 138.04, 137.32 (d, *J* = 8.1 Hz), 126.99, 125.36 (d, *J* = 18.6 Hz), 124.95 (d, *J* = 5.0 Hz), 124.49 (d, *J* = 4.3 Hz), 122.41, 118.92 (d, *J* = 14.3 Hz), 115.65 (d, *J* = 26.7 Hz), 95.26, 79.03, 58.71, 54.01, 45.41, 28.02, 11.00.

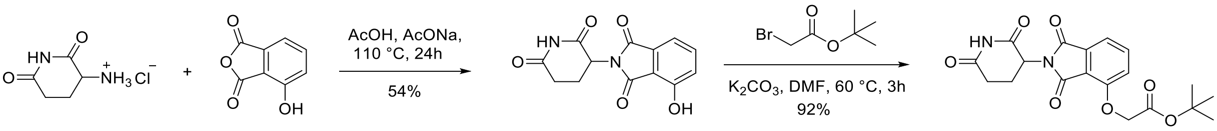


**19** **20** **21** **22**

##### 2-(2,6-dioxopiperidin-3-yl)-4-hydroxyisoindoline-1,3-dione **21**[8]

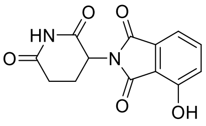

**21**

4-hydroxyisobenzofuran-1,3-dione **20** (250 mg, 1.52 mmol, 1.0 eq) and 2,6-dioxopiperidin-3-aminium chloride **19** (250 mg, 1.52 mmol, 1.0 eq.) were dissolved in acetic acid (6 mL), followed by addition of sodium acetate (250 mg, 3.05 mmol, 2.0 eq). The reaction mixture was stirred under reflux for 24 h. After cooling to room temperature, the precipitate was filtered and washed with ice-cold water and diethylether to obtain the desired product **21** as an off-white powder (223 mg, 54%).

**$^1$H NMR** (400 MHz, DMSO) δ 11.17 (s, 1H), 11.07 (s, 1H), 7.64 (dd, *J* = 8.4, 7.2 Hz, 1H), 7.31 (d, *J* = 6.8 Hz, 1H), 7.24 (d, *J* = 7.8 Hz, 1H), 5.06 (dd, *J* = 12.8, 5.4 Hz, 1H), 2.87 (m, 1H), 2.63 – 2.51 (m, 2H), 2.05 – 1.96 (m, 1H).

**$^{13}$C NMR** (101 MHz, DMSO) δ 173.27, 170.49, 167.50, 166.29, 155.98, 136.85, 133.63, 124.05, 114.83, 114.73, 49.10, 40.67, 40.62, 40.46, 40.41, 40.26, 40.20, 40.00, 39.79, 39.58, 39.37, 31.43, 22.50.

##### *tert*-butyl 2-((2-(2,6-dioxopiperidin-3-yl)-1,3-dioxoisoindolin-4-yl)oxy)acetate **22**

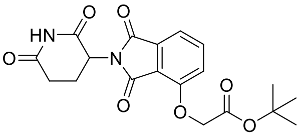

**22**

Hydroxy-thalidomide **21** (100 mg, 0.36 mmol, 1.0 eq) was dissolved in anh. DMF (5 mL) and potassium carbonate (76 mg, 0.55 mmol, 1.5 eq) and *tert*-butyl bromoacetate (71 mg, 0.36 mmol, 1.0 eq) were added. The reaction mixture was stirred at 60 °C for 3 h. Upon completion and cooling to room temperature, the reaction mixture was diluted with EtOAc and washed with water, 5% LiCl and brine. The combined organic phases were dried over anh. $MgSO_4$, filtered, and evaporated. The crude was dissolved in $CHCl_3$ and purified by flash column chromatography (pentane/EtOAc; gradient 0-100%) to afford the product **22** as a white solid (130 mg, 92%).

$^{1}$**H NMR** (400 MHz, DMSO) δ 11.10 (s, 1H), 7.80 (dd, *J* = 8.5, 7.3 Hz, 1H), 7.48 (d, *J* = 7.2 Hz, 1H), 7.38 (d, *J* = 8.5 Hz, 1H), 5.10 (dd, *J* = 12.8, 5.4 Hz, 1H), 4.97 (s, 2H), 2.95 – 2.83 (m, 1H), 2.64 – 2.52 (m, 2H), 2.09 – 1.99 (m, 1H), 1.43 (s, 9H).
$^{13}$**C NMR** (101 MHz, DMSO) δ 172.78, 169.89, 167.14, 166.72, 165.12, 155.04, 136.76, 133.25, 119.97, 116.45, 115.90, 81.91, 65.50, 48.79, 40.15, 39.94, 39.73, 39.52, 39.31, 39.10, 38.89, 30.94, 27.67, 21.96.

**Synthesis of Degraders**

**General Protocol A:**

Compound **22** was dissolved in trifluoroacetic acid (TFA) and left for 2 h at room temperature to allow deprotection. Upon completion, the reaction mixture was diluted with DCM and concentrated to dryness. The obtained product was dissolved in anh. DMSO, followed by the addition of *N,N*-diisopropylethylamine (DIPEA) (20 eq) and *O*-[*N*-succinimidyl)-1,1,3,3-tetramethyluronium tetrafluoroborate (TSTU) (1.1 eq). This mixture was added into the suitable *N*-Boc-protected linker (**23a-c**, **25a-b**) (2 eq), dissolved in DMSO. After 2 h, the reaction was acidified by acetic acid and diluted with water. The final product was obtained by RP-HPLC (ACN/$H_2O$ gradient; 10-90%, 25 min).

**General Protocol B:**

Suitable thalidomide-linker-NHBoc (**24a-c, 26a-b**) was dissolved in TFA and left at room temperature for 2 h to allow deprotection. TFA was evaporated by the flow of nitrogen and the obtained product was used without further purification.

Compound **16** (1 eq) was dissolved in anh. DMSO, followed by the addition of *N,N*-diisopropylethylamine (DIPEA) (20 eq.) and benzotriazole-1-yl-oxy-tris-pyrrolidino-phosphonium hexafluorophosphate (PyBOP) (1.2 eq). This mixture was added into the previously deprotected thalidomide-linker (2 eq), dissolved in DMSO. After 2 h, the reaction was acidified by acetic acid and diluted with water. The final product was obtained by RP-HPLC (ACN/$H_2O$ step gradient, 10-90%, 35 min).

**General Protocol C:**

**21 (**1 eq.) was mixed with **27a-b** (1.50 eq), potassium carbonate (3 eq) and potassium iodide (1 eq) in DMF. The reaction was stirred for 12 h at 70 °C. Afterwards, it was diluted with water and extracted with ethyl acetate. The organic layers were washed with water and brine, dried over $MgSO_4$ and concentrated. Crude was dissolved in DMSO and the product was obtained by RP-HPLC (ACN/$H_2O$ step gradient, 30-90%, 60 min).

*tert*-butyl-(2-(2-(2-((2-(2,6-dioxopiperidin-3-yl)-1,3-dioxoisoindolin-4-yl)oxy)acetamido)ethoxy)ethyl)carbamate **24a**

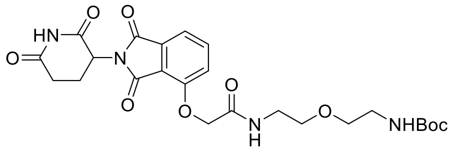


**24a**

General protocol **A** was followed. **24a** was obtained as a white powder (4.0 mg, 51%).

**¹H NMR** (500 MHz, DMSO) δ 11.11 (s, 1H), 8.02 (t, *J* = 5.6 Hz, 1H), 7.81 (dd, *J* = 8.5, 7.3 Hz, 1H), 7.49 (d, *J* = 7.2 Hz, 1H), 7.39 (d, *J* = 8.5 Hz, 1H), 6.75 (t, *J* = 5.8 Hz, 1H), 5.12 (dd, *J* = 12.8, 5.4 Hz, 1H), 4.79 (s, 2H), 3.44 (t, *J* = 5.7 Hz, 2H), 3.38 (t, *J* = 6.2 Hz, 2H), 3.33 – 3.28 (m, 2H), 3.12 – 3.04 (m, 2H), 2.89 (ddd, *J* = 16.8, 13.7, 5.3 Hz, 1H), 2.64 – 2.51 (m, 2H), 2.08 – 1.99 (m, 1H), 1.36 (s, 9H).

**¹³C NMR** (126 MHz, DMSO) δ 172.76, 169.88, 166.90, 166.74, 165.47, 155.61, 155.03, 136.94, 133.05, 120.35, 116.77, 116.03, 77.64, 69.03, 68.56, 67.49, 48.81, 38.39, 30.96, 28.23, 22.00.

*tert*-butyl-(2-(2-(2-(2-((2-(2,6-dioxopiperidin-3-yl)-1,3-dioxoisoindolin-4-yl)oxy)acetamido)ethoxy)ethoxy)ethyl)carbamate **24b**

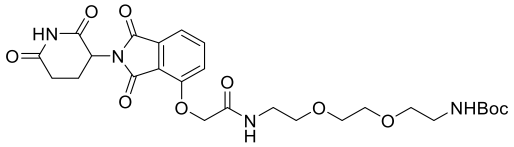


**24b**

General protocol **A** was followed. **24b** was obtained as a white powder (13.8 mg, 82%).

**¹H NMR** (400 MHz, DMSO) δ 11.11 (s, 1H), 8.01 (t, *J* = 5.7 Hz, 1H), 7.81 (dd, *J* = 8.5, 7.3 Hz, 1H), 7.50 (d, *J* = 7.2 Hz, 1H), 7.40 (d, *J* = 8.6 Hz, 1H), 6.74 (s, 0H), 5.11 (dd, *J* = 12.9, 5.4 Hz, 1H), 4.79 (s, 2H), 3.49 (d, *J* = 2.1 Hz, 2H), 3.48 – 3.44 (m, 2H), 3.41 – 3.28 (m, 6H), 3.22 –

3.10 (m, 1H), 3.08 – 3.02 (m, 2H), 2.96 – 2.83 (m, 1H), 2.65 – 2.52 (m, 2H), 2.10 – 1.99 (m, 1H), 1.36 (s, 9H).

**$^{13}$C NMR** (101 MHz, DMSO) δ 172.78, 170.86, 169.88, 166.91, 166.74, 165.45, 155.60, 155.00, 136.94, 133.05, 120.35, 117.00, 116.78, 116.05, 113.42, 77.61, 70.12, 69.58, 69.54, 69.47, 69.18, 69.12, 68.83, 67.51, 48.82, 38.47, 36.16, 35.79, 30.96, 28.23, 22.01.

*tert*-butyl (1-((2-(2,6-dioxopiperidin-3-yl)-1,3-dioxoisoindolin-4-yl) oxy)-2-oxo-6,9,12-trioxa-3-azatetradecan-14-yl) carbamate **24c**

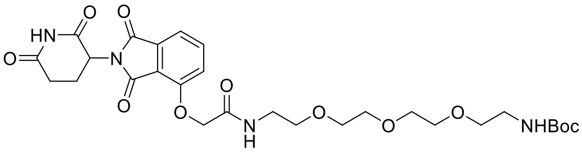


**24c**

General protocol **A** was followed. **24c** was obtained as a white powder (3.8 mg, 42%).

**$^{1}$H NMR** (400 MHz, DMSO) δ 11.10 (s, 1H), 8.00 (t, *J* = 5.7 Hz, 1H), 7.80 (dd, *J* = 8.6, 7.2 Hz, 1H), 7.49 (d, *J* = 7.2 Hz, 1H), 7.39 (d, *J* = 8.4 Hz, 1H), 6.72 (t, *J* = 5.6 Hz, 1H), 5.10 (dd, *J* = 12.8, 5.5 Hz, 1H), 4.78 (s, 2H), 3.49 – 3.43 (m, 10H), 3.39 – 3.28 (m, 6H), 3.21 – 3.08 (m, 1H), 3.07 – 3.00 (m, 2H), 2.95 – 2.82 (m, 1H), 2.63 – 2.52 (m, 2H), 2.09 – 1.97 (m, 1H), 1.38 – 1.33 (m, 9H).

**$^{13}$C NMR** (101 MHz, DMSO) δ 172.78, 170.86, 169.87, 166.91, 166.74, 165.45, 155.59, 154.99, 136.96, 133.05, 120.36, 116.77, 116.06, 77.60, 70.12, 69.74, 69.63, 69.58, 69.50, 69.16, 69.11, 68.83, 67.51, 48.82, 38.41, 30.95, 28.23, 22.01.

2-((2-(2,6-dioxopiperidin-3-yl)-1,3-dioxoisoindolin-4-yl)oxy)-*N*-(2-(2-(2-(1-(2-((2-fluoro-4-((2-fluoro-3-nitrobenzyl)sulfonyl)phenyl)thio)-5-methoxy-6-((5-methyl-1*H*-pyrazol-3-yl)amino)pyrimidin-4-yl)piperidin-4-yl)acetamido)ethoxy)ethyl)acetamide **D1**

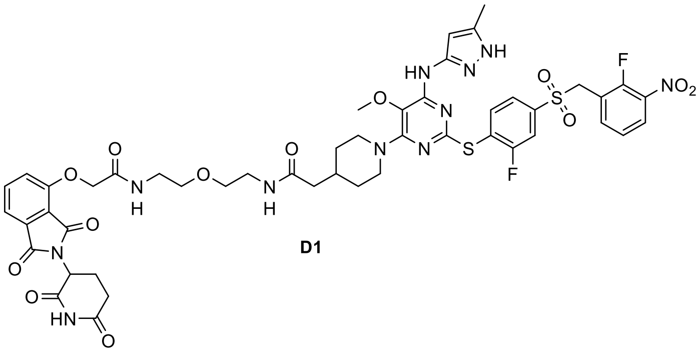


**D1**

General protocol **B** was followed. **D1** was obtained as a white powder (1.5 mg, 47%).

**$^{1}$H NMR** (500 MHz, DMSO) δ 11.11 (s, 1H), 8.65 (s, 1H), 8.20 – 8.14 (m, 1H), 8.03 – 7.99 (m, 1H), 7.98 – 7.92 (m, 1H), 7.85 – 7.77 (m, 3H), 7.67 – 7.61 (m, 2H), 7.49 (d, *J* = 7.3 Hz, 1H), 7.46 (t, 1H), 7.39 (d, *J* = 8.5 Hz, 1H), 5.54 (s, 1H), 5.13 (dd, *J* = 12.7, 5.5 Hz, 1H), 5.00 (s, 2H), 4.79 (s, 2H), 4.15 (d, *J* = 12.7 Hz, 2H), 3.52 (s, 3H), 3.45 (t, *J* = 5.7 Hz, 2H), 3.40 (t, *J* = 6.0 Hz, 2H), 3.32 (q, *J* = 5.7 Hz, 2H), 3.21 (q, *J* = 6.0 Hz, 2H), 2.94 – 2.83 (m, 1H), 2.82 – 2.74 (m, 2H), 2.64 – 2.55 (m, 2H), 2.05 (s, 3H), 2.04 – 2.01 (m, 1H), 1.99 (d, *J* = 7.1 Hz, 2H), 1.86 (s, 1H), 1.14 – 1.04 (m, 2H).

**$^{13}$C NMR** (126 MHz, DMSO) δ 172.79, 171.06, 169.91, 166.95, 166.75, 165.49, 161.64 (d, J = 251.5 Hz), 159.57, 154.99, 153.62, 153.50, 153.46 (d, J = 265.6 Hz), 146.49, 140.60 (d, J = 6.8 Hz), 139.06 (d, J = 3.6 Hz), 138.03, 137.32 (d, J = 7.9 Hz), 136.95, 133.04, 127.01, 125.50 (d, J = 18.3 Hz), 124.95 (d, J = 4.8 Hz), 124.47 (d, J = 3.8 Hz), 122.00, 120.36, 118.91 (d, J = 14.2 Hz), 116.77, 116.06, 115.63 (d, J = 26.7 Hz), 95.08, 68.93, 68.60, 67.51, 58.49, 54.06, 48.81, 45.70, 42.26, 38.33, 33.11, 31.58, 30.95, 22.01, 11.05.

**HRMS** (ESI)**:** calc. for $C_{48}H_{50}F_2N_{11}O_{13}S_2$ [M+H]+: 1090.2999; found 1090.2992

2-((2-(2,6-dioxopiperidin-3-yl)-1,3-dioxoisoindolin-4-yl)oxy)-*N*-(2-(2-(2-(2-(1-(2-((2-fluoro-4-((2-fluoro-3-nitrobenzyl)sulfonyl)phenyl)thio)-5-methoxy-6-((5-methyl-1*H*-pyrazol-3-yl)amino)pyrimidin-4-yl)piperidin-4-yl)acetamido)ethoxy)ethoxy)ethyl)acetamide **D2**

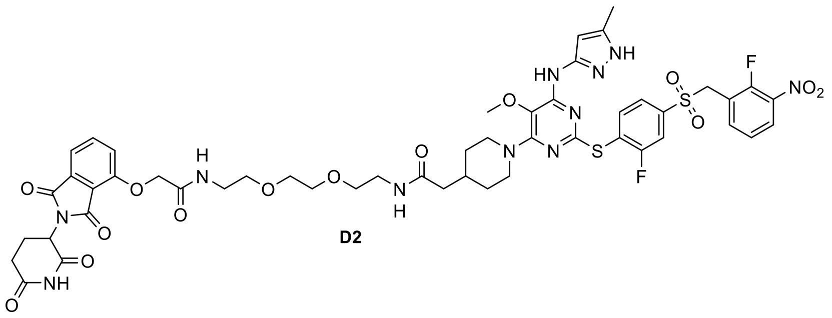

General protocol **B** was followed. **D2** was obtained as a white powder (1.6 mg, 49%).

**$^{1}$H NMR** (500 MHz, DMSO) δ 11.12 (s, 1H), 8.67 (s, 1H), 8.17 (t, *J* = 7.7 Hz, 1H), 8.03 – 7.99 (m, 1H), 7.96 (t, *J* = 7.5 Hz, 1H), 7.87 – 7.83 (m, 1H), 7.79 (d, *J* = 7.7 Hz, 2H), 7.66 – 7.62 (m, 2H), 7.49 (d, *J* = 7.4 Hz, 1H), 7.45 (t, *J* = 8.0 Hz, 1H), 7.39 (d, *J* = 8.6 Hz, 1H), 5.54 (s, 1H), 5.11 (dd, *J* = 12.9, 5.5 Hz, 1H), 5.00 (s, 2H), 4.78 (s, 2H), 4.16 (d, *J* = 12.9 Hz, 2H), 3.18 (q, *J* = 5.9 Hz, 2H), 2.93 – 2.85 (m, 1H), 2.83 – 2.74 (m, 3H), 2.05 (s, 3H), 2.02 (s, 1H), 1.99 (d, *J* = 7.2 Hz, 2H), 1.87 (s, 1H), 1.59 (d, *J* = 13.0 Hz, 2H), 1.13 – 1.08 (m, 2H).

$^{13}$**C NMR** (126 MHz, DMSO) δ 172.83, 171.08, 169.92, 166.96, 166.77, 165.49, 161.66 (d, J = 251.7 Hz), 159.59, 155.00, 153.64, 153.50, 153.48 (d, J = 265.6 Hz), 146.47, 140.62 (d, J = 6.5 Hz), 139.52, 139.09, 138.06, 137.37, 136.98, 133.07, 127.04, 125.52 (d, J = 18.0 Hz), 124.98 (d, J = 5.0 Hz), 124.49, 122.03, 120.37, 118.92 (d, J = 14.0 Hz), 116.79, 116.09, 115.65 (d, J = 26.1 Hz), 95.10, 69.61, 69.54, 69.17, 68.86, 67.53, 58.53, 54.08, 48.84, 45.73, 42.30, 38.44, 33.16, 31.59, 30.97, 22.03, 11.07.

**HRMS** (ESI)**:** calc. for $C_{50}H_{54}F_2N_{11}O_{14}S_2$ [M+H]$^+$: 1134.3261; found 1134.3252

2-((2-(2,6-dioxopiperidin-3-yl)-1,3-dioxoisoindolin-4-yl)oxy)-*N*-(1-(1-(2-((2-fluoro-4-((2-fluoro-3-nitrobenzyl)sulfonyl)phenyl)thio)-5-methoxy-6-((5-methyl-1*H*-pyrazol-3-yl)amino)pyrimidin-4-yl)piperidin-4-yl)-2-oxo-6,9,12-trioxa-3-azatetradecan-14-yl)acetamide **D3**

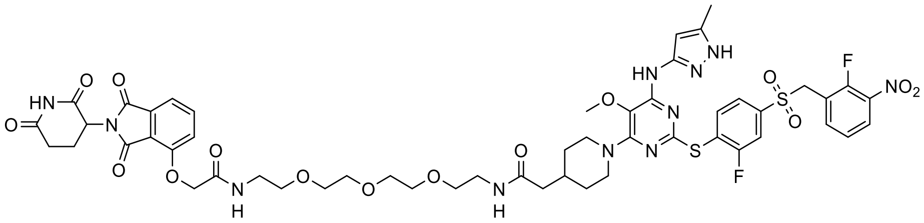

**D3**

General protocol **B** was followed. **D3** was obtained as a white powder (2.1 mg, 61%).

$^1$**H NMR** (400 MHz, DMSO) δ 11.11 (s, 1H), 8.76 (s, 1H), 8.22 – 8.11 (m, 1H), 8.01 (t, J = 5.7 Hz, 1H), 7.96 (dd, J = 8.1, 6.8 Hz, 1H), 7.87 – 7.83 (m, 1H), 7.83 – 7.77 (m, 2H), 7.69 – 7.60 (m, 2H), 7.49 (d, J = 7.3 Hz, 1H), 7.45 (t, J = 8.0 Hz, 1H), 7.39 (d, J = 8.6 Hz, 1H), 5.58 (s, 1H), 5.11 (dd, J = 13.0, 5.4 Hz, 1H), 5.01 (s, 2H), 4.78 (s, 2H), 4.16 (d, J = 13.0 Hz, 2H), 3.51 (s, 3H), 3.53 – 3.44 (m, 10H), 3.38 (t, J = 5.9 Hz, 2H), 3.31 (q, J = 5.7 Hz, 2H), 3.18 (q, J = 5.9 Hz, 2H), 2.95 – 2.84 (m, 1H), 2.79 (t, J = 12.4 Hz, 2H), 2.65 – 2.52 (m, 2H), 2.07 (s, 3H), 2.06 – 2.01 (m, 1H), 1.99 (d, J = 7.2 Hz, 2H), 1.93 – 1.81 (m, 1H), 1.59 (d, J = 12.7 Hz, 2H), 1.16 – 1.03 (m, 2H).

$^{13}$**C NMR** (126 MHz, DMSO) δ 172.78, 170.99, 169.88, 166.88 (d, J = 7.5 Hz), 166.74, 165.45, 161.63 (d, J = 251.5 Hz), 159.57, 154.99, 153.62, 153.45 (d, J = 266.2 Hz), 153.38, 146.22, 140.62 (d, J = 6.6 Hz), 139.81, 139.06 (d, J = 3.7 Hz), 138.03, 137.32 (d, J = 7.9 Hz), 136.95, 133.04, 127.01, 125.48 (d, J = 18.5 Hz), 124.95 (d, J = 4.8 Hz), 124.45, 122.07, 120.35, 118.91 (d, J = 14.1 Hz), 116.76, 116.05, 115.63 (d, J = 26.4 Hz), 95.10, 69.75, 69.63,

69.55, 69.13, 68.83, 67.51, 58.54, 54.04, 48.81, 45.70, 42.26, 38.40, 33.13, 31.57, 30.95, 22.00, 11.04

**HRMS** (ESI)**:** calc. for $C_{52}H_{58}F_2N_{11}O_{15}S_2$ $[M+H]^+$: 1178.3523; found 1178.3515

*tert*-butyl-(2-(2-((2-(2,6-dioxopiperidin-3-yl)-1,3-dioxoisoindolin-4-yl)oxy) acetamido) ethyl) carbamate **26a**

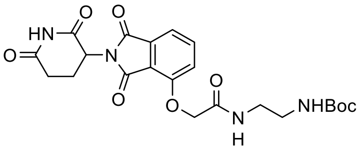

**26a**

General protocol **A** was followed. **26a** was obtained as a white powder (4.8 mg, 67%).

**$^1$H NMR** (400 MHz, DMSO) δ 11.11 (s, 1H), 8.03 (t, *J* = 5.9 Hz, 1H), 7.81 (t, *J* = 7.9 Hz, 1H), 7.50 (d, *J* = 7.2 Hz, 1H), 7.39 (d, *J* = 8.6 Hz, 1H), 6.85 (t, *J* = 5.9 Hz, 1H), 5.12 (dd, *J* = 12.8, 5.4 Hz, 1H), 4.76 (s, 2H), 3.17 (q, *J* = 6.4 Hz, 2H), 3.01 (q, *J* = 6.2 Hz, 2H), 2.90 (ddd, *J* = 19.6, 14.1, 5.5 Hz, 1H), 2.68 – 2.51 (m, 2H), 2.08 – 1.99 (m, 1H), 1.36 (s, 9H).

**$^{13}$C NMR** (101 MHz, DMSO) δ 172.77, 169.86, 167.03, 166.74, 165.42, 155.66, 155.13, 136.93, 133.04, 120.45, 116.79, 116.04, 77.70, 67.59, 48.80, 40.20, 40.15, 39.99, 39.94, 39.78, 39.73, 39.52, 39.31, 39.10, 38.89, 38.61, 30.94, 28.20, 21.99.

*tert*-butyl-(6-(2-((2-(2,6-dioxopiperidin-3-yl)-1,3-dioxoisoindolin-4-yl) oxy) acetamido) hexyl) carbamate **26b**

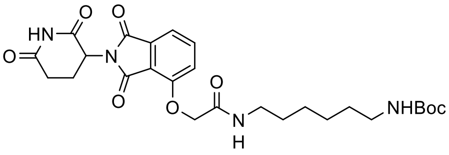

**26b**

General protocol **A** was followed. **26b** was obtained as a white powder (6 mg, 38%).

**$^1$H NMR** (500 MHz, DMSO) δ 11.05 (s, 1H), 7.86 (t, *J* = 5.8 Hz, 1H), 7.74 (dd, *J* = 8.5, 7.3 Hz, 1H), 7.43 (d, *J* = 7.2 Hz, 1H), 7.32 (d, *J* = 8.5 Hz, 1H), 6.69 (t, *J* = 5.7 Hz, 1H), 5.05 (dd, *J* = 12.8, 5.4 Hz, 1H), 4.70 (s, 2H), 3.06 (q, *J* = 6.7 Hz, 2H), 2.84 – 2.78 (m, 2H), 2.58 – 2.46 (m, 2H), 2.01 – 1.94 (m, 1H), 1.38 – 1.32 (m, 2H), 1.30 (s, 9H), 1.29 – 1.24 (m, 2H), 1.23 – 1.10 (m, 4H).

**$^{13}$C NMR** (126 MHz, DMSO) δ 173.25, 170.35, 167.21, 167.08, 165.98, 156.05, 155.52, 137.39, 133.50, 120.85, 117.29, 116.50, 77.77, 68.10, 49.27, 38.75, 31.41, 29.90, 29.45, 28.74, 26.44, 22.47.

2-((2-(2,6-dioxopiperidin-3-yl)-1,3-dioxoisoindolin-4-yl)oxy)-*N*-(2-(2-(1-(2-((2-fluoro-4-((2-fluoro-3-nitrobenzyl)sulfonyl)phenyl)thio)-5-methoxy-6-((5-methyl-1*H*-pyrazol-3-yl)amino)pyrimidin-4-yl)piperidin-4-yl)acetamido)ethyl)acetamide **D4**

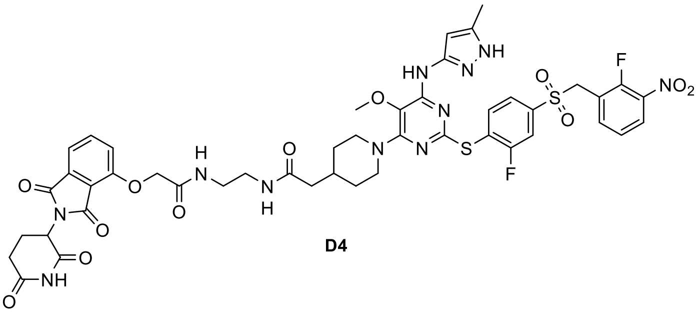


**D4**

General protocol **B** was followed. **D4** was obtained as a white powder (1.2 mg, 40%).

**$^{1}$H NMR** (500 MHz, DMSO) δ 11.12 (s, 1H), 8.71 (s, 1H), 8.17 (t, J = 7.6 Hz, 1H), 8.06 – 8.00 (m, 1H), 7.99 – 7.92 (m, 1H), 7.91 – 7.85 (m, 1H), 7.84 – 7.77 (m, 2H), 7.68 – 7.61 (m, 2H), 7.50 (d, J = 7.1 Hz, 1H), 7.45 (t, J = 7.9 Hz, 1H), 7.39 (d, J = 8.5 Hz, 1H), 5.55 (s, 1H), 5.12 (dd, J = 13.0, 5.4 Hz, 1H), 5.00 (s, 2H), 4.77 (s, 2H), 4.15 (d, J = 12.7 Hz, 2H), 3.51 (s, 3H), 3.23 – 3.10 (m, 4H), 2.94 – 2.83 (m, 1H), 2.81 – 2.71 (m, 2H), 2.64 – 2.53 (m, 2H), 2.06 (s, 3H), 2.03 (s, 1H), 1.96 (d, J = 7.0 Hz, 2H), 1.86 (s, 1H), 1.59 (d, J = 12.4 Hz, 2H), 1.09 (q, J = 12.1, 11.6 Hz, 2H).

**$^{13}$C NMR** (126 MHz, DMSO) δ 172.78, 171.19, 169.88, 167.06, 166.76, 165.45, 161.63 (d, J = 251.4 Hz), 159.57, 155.10, 153.64, 153.45 (d, J = 266.2 Hz), 153.43, 146.35, 140.61 (d, J = 6.7 Hz), 139.61, 139.06 (d, J = 3.7 Hz), 138.03, 137.32 (d, J = 7.7 Hz), 136.96, 133.05, 127.01, 125.49 (d, J = 18.3 Hz), 124.95 (d, J = 4.8 Hz), 124.47 (d, J = 3.9 Hz), 122.03, 120.47, 118.91 (d, J = 14.2 Hz), 116.84, 116.08, 115.63 (d, J = 26.7 Hz), 95.08, 67.62, 58.51, 54.05, 48.81, 45.69, 42.37, 38.38, 38.04, 33.01, 31.60, 30.95, 22.01, 11.04.

**HRMS** (ESI)**:** calc. for $C_{46}H_{46}F_2N_{11}O_{12}S_2$ $[M+H]^+$: 1046.2737; found 1046.2727

2-((2-(2,6-dioxopiperidin-3-yl)-1,3-dioxoisoindolin-4-yl)oxy)-*N*-(6-(2-(1-(2-((2-fluoro-4-((2-fluoro-3-nitrobenzyl)sulfonyl)phenyl)thio)-5-methoxy-6-((5-methyl-1*H*-pyrazol-3-yl)amino)pyrimidin-4-yl)piperidin-4-yl)acetamido)hexyl)acetamide **D5**

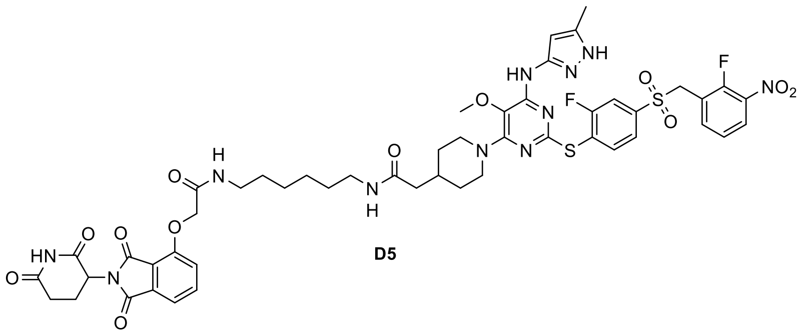

**D5**

General protocol **B** was followed. **D5** was obtained as a white powder (1.8 mg, 56%).

**$^1$H NMR** (500 MHz, DMSO) δ 11.11 (s, 1H), 8.81 (s, 1H), 8.21 – 8.14 (m, 1H), 7.98 – 7.91 (m, 2H), 7.82 – 7.79 (m, 2H), 7.75 (t, J = 5.6 Hz, 1H), 7.69 – 7.60 (m, 2H), 7.49 (d, J = 7.3 Hz, 1H), 7.45 (t, J = 7.9 Hz, 1H), 7.39 (d, J = 8.5 Hz, 1H), 5.59 (s, 1H), 5.12 (dd, J = 12.8, 5.5 Hz, 1H), 5.01 (s, 2H), 4.77 (s, 2H), 4.16 (d, J = 12.8 Hz, 2H), 3.52 (s, 3H), 3.14 (q, J = 6.6 Hz, 2H), 3.00 (q, J = 6.6 Hz, 2H), 2.95 – 2.84 (m, 1H), 2.79 (t, J = 12.4 Hz, 2H), 2.62 – 2.55 (m, 2H), 2.54 (s, 1H), 2.07 (s, 3H), 2.05 – 2.00 (m, 1H), 1.97 (d, J = 7.1 Hz, 2H), 1.92 – 1.83 (m, 1H), 1.59 (d, J = 12.5 Hz, 2H), 1.46 – 1.39 (m, 2H), 1.39 – 1.33 (m, 2H), 1.30 – 1.20 (m, 4H), 1.15 – 1.05 (m, 2H).

**$^{13}$C NMR** (126 MHz, DMSO) δ 172.78, 170.63, 169.88, 166.74, 166.62, 165.51, 161.62 (d, J = 251.4 Hz), 159.55, 155.05, 153.65, 153.44 (d, J = 265.6 Hz), 153.36, 146.21, 140.62 (d, J = 6.4 Hz), 139.86, 139.05 (d, J = 3.5 Hz), 138.02, 137.31 (d, J = 7.7 Hz), 136.92, 133.03, 127.00, 125.47 (d, J = 18.3 Hz), 124.94 (d, J = 4.6 Hz), 124.44 (d, J = 3.8 Hz), 122.07, 120.37, 118.90 (d, J = 14.4 Hz), 116.82, 116.04, 115.61 (d, J = 26.6 Hz), 95.10, 67.64, 58.53, 54.03, 48.80, 45.69, 42.39, 38.26, 33.11, 31.59, 30.94, 29.11, 28.97, 26.08, 25.96, 22.00, 11.03.

**HRMS** (ESI)**:** calc. for $C_{46}H_{46}F_2N_{11}O_{12}S_2$ $[M+H]^+$: 1102.3363; found 1102.3356

*tert*-butyl-(4-((2-(2,6-dioxopiperidin-3-yl)-1,3-dioxoisoindolin-4-yl)oxy)butyl)carbamate **28b**

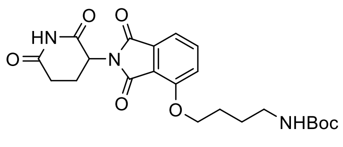

**28b**

General protocol **C** was followed. **28b** was obtained as a light yellow powder (28 mg, 34%).

**$^1$H NMR** (400 MHz, DMSO) δ 11.09 (s, 1H), 7.81 (dd, *J* = 8.5, 7.3 Hz, 1H), 7.54 – 7.47 (m, 1H), 7.44 (dd, *J* = 7.2, 0.6 Hz, 1H), 6.84 (t, *J* = 5.8 Hz, 1H), 5.07 (dd, *J* = 12.8, 5.4 Hz, 1H), 4.21 (t, *J* = 6.4 Hz, 2H), 2.98 (q, *J* = 6.6 Hz, 2H), 2.88 (ddd, *J* = 17.0, 13.8, 5.3 Hz, 1H), 2.62 – 2.52 (m, 2H), 2.08 – 1.98 (m, 1H), 1.79 – 1.68 (m, 2H), 1.60 – 1.51 (m, 2H), 1.37 (s, 9H).

**$^{13}$C NMR** (101 MHz, DMSO) δ 172.77, 169.93, 166.84, 165.32, 155.94, 155.62, 137.03, 133.24, 119.79, 116.22, 115.16, 77.37, 68.55, 48.72, 40.15, 39.94, 39.73, 39.52, 39.31, 39.10, 38.89, 30.94, 28.26, 27.06, 25.94, 25.83, 22.00.

*N*-(2-((2-(2,6-dioxopiperidin-3-yl)-1,3-dioxoisoindolin-4-yl) oxy) ethyl)-2-(1-(2-((2-fluoro-4-((2-fluoro-3-nitrobenzyl) sulfonyl) phenyl) thio)-5-methoxy-6-((5-methyl-1*H*-pyrazol-3-yl) amino) pyrimidin-4-yl)piperidin-4-yl) acetamide **D6**

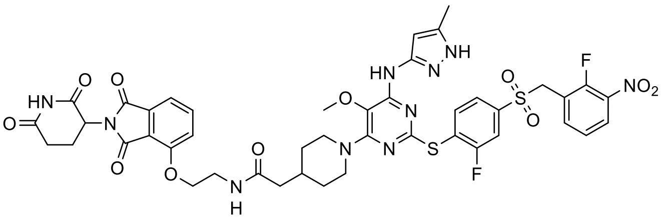


**D6**

General protocol **B** was followed. **D6** was obtained as a yellow powder (1.4 mg, 49%).

**$^1$H NMR** (500 MHz, DMSO) δ 11.10 (s, 1H), 8.64 (s, 1H), 8.16 (t, *J* = 7.6 Hz, 1H), 8.06 (t, *J* = 5.6 Hz, 1H), 8.01 – 7.92 (m, 1H), 7.85 – 7.78 (m, 2H), 7.68 – 7.61 (m, 2H), 7.55 (d, *J* = 8.6 Hz, 1H), 7.49 – 7.41 (m, 2H), 5.55 (s, 1H), 5.08 (dd, *J* = 12.7, 5.5 Hz, 1H), 5.00 (s, 2H), 4.25 (t, *J* = 5.9 Hz, 2H), 4.14 (d, *J* = 12.7 Hz, 2H), 3.51 (s, 3H), 2.93 – 2.83 (m, 1H), 2.77 (t, *J* = 12.5 Hz, 2H), 2.62 – 2.54 (m, 2H), 2.05 (s, 3H), 2.02 (d, *J* = 7.4 Hz, 2H), 1.93 – 1.83 (m, 1H), 1.60 (d, *J* = 12.5 Hz, 2H), 1.14 – 1.05 (m, 2H).

**$^{13}$C NMR** (126 MHz, DMSO) δ 172.81, 171.47, 169.94, 166.82, 165.25, 161.65 (d, *J* = 251.6 Hz), 159.57, 155.73, 153.59, 153.50, 153.46 (d, *J* = 266.2 Hz), 146.48, 140.61 (d, *J* = 6.7 Hz), 139.38, 139.07 (d, *J* = 3.7 Hz), 138.04, 137.31 (d, *J* = 8.0 Hz), 137.05, 133.29, 127.01, 125.51 (d, *J* = 18.4 Hz), 124.95 (d, *J* = 5.0 Hz), 124.47 (d, *J* = 3.8 Hz), 121.98, 120.13, 118.90 (d, *J* = 14.4 Hz), 116.46, 115.74, 115.54, 95.09, 67.39, 58.49, 54.06, 48.77, 45.69, 42.29, 37.87, 33.12, 31.59, 30.96, 22.04, 11.05.

**HRMS** (ESI): calc. for $C_{44}H_{43}F_2N_{10}O_{11}S_2$ $[M+H]^+$: 989.2522; found 989.2510

*N*-(4-((2-(2,6-dioxopiperidin-3-yl)-1,3-dioxoisoindolin-4-yl) oxy) butyl)-2-(1-(2-((2-fluoro-4-((2-fluoro-3-nitrobenzyl) sulfonyl) phenyl) thio)-5-methoxy-6-((5-methyl-1*H*-pyrazol-3-yl) amino) pyrimidin-4-yl)piperidin-4-yl) acetamide **D7**

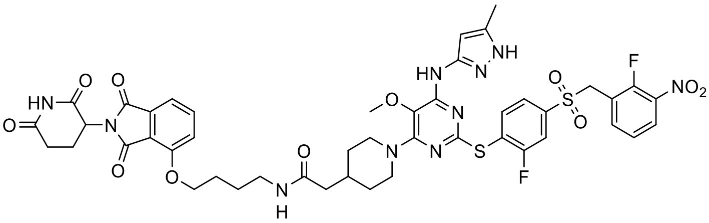

**D7**

General protocol **B** was followed. **D7** was obtained as a yellow powder (1.05 mg, 34%).

**$^1$H NMR** (400 MHz, DMSO) δ 11.10 (s, 1H), 8.72 (s, 1H), 8.20 – 8.14 (m, 1H), 7.96 (t, 1H), 7.86 – 7.78 (m, 3H), 7.67 – 7.62 (m, 2H), 7.51 (d, *J* = 8.6 Hz, 1H), 7.48 – 7.42 (m, 2H), 5.58 (s, 1H), 5.08 (dd, *J* = 12.9, 5.3 Hz, 1H), 5.01 (s, 2H), 4.22 (t, *J* = 6.4 Hz, 2H), 4.15 (d, *J* = 12.7 Hz, 2H), 3.52 (s, 3H), 3.14 – 3.08 (m, 2H), 2.93 – 2.84 (m, 1H), 2.79 (t, *J* = 13.4 Hz, 2H), 2.62 – 2.52 (m, 2H), 2.06 (s, 3H), 2.05 – 2.02 (m, 1H), 2.01 – 1.97 (m, 2H), 1.89 (s, 1H), 1.78 – 1.71 (m, 2H), 1.59 (q, *J* = 7.4, 6.8 Hz, 5H), 1.16 – 1.04 (m, 2H).

**$^{13}$C NMR** (126 MHz, DMSO) δ 172.79, 170.76, 169.98, 166.86, 165.37, 161.63 (d, *J* = 251.6 Hz), 159.56, 155.94, 153.63, 153.45 (d, *J* = 266.2 Hz), 153.43, 146.35, 140.61 (d, *J* = 6.8 Hz), 139.64, 139.05, 138.03, 137.29, 137.06, 133.25, 127.01, 125.49 (d, *J* = 18.5 Hz), 124.95 (d, *J* = 4.3 Hz), 124.45, 122.04, 119.76, 118.90 (d, *J* = 14.1 Hz), 116.21, 115.62 (d, *J* = 26.8 Hz), 115.19, 95.11, 68.50, 58.52, 54.04, 48.74, 45.70, 42.41, 37.93, 33.12, 31.61, 30.96, 25.94, 25.67, 22.02, 11.04.

**HRMS** (ESI): calc. for $C_{46}H_{46}F_2N_{10}O_{11}S_2$ $[M+H]^+$: 1017.2835; found 1017.2830

2-(2,6-dioxopiperidin-3-yl)-4-fluoroisoindoline-1,3-dione **30**

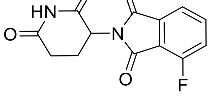

**30**

4-fluoroisobenzofuran-1,3-dione **29** (252 mg, 1.52 mmol, 1 eq) and 2,6-dioxopiperidin-3-aminium chloride **19** (250 mg, 1.52 mmol, 1 eq) were dissolved in acetic acid (6 mL), followed by addition of sodium-acetate (249 mg, 3.04 mmol, 2 eq). The reaction mixture was stirred under reflux for 24h. After cooling to room temperature, the precipitate was filtered and washed with ice-cold water and diethyl-ether to obtain the desired product **30** as off-white powder (191 mg, 46 %).

**$^1$H NMR** (400 MHz, DMSO) δ 11.13 (s, 1H), 7.94 (td, *J* = 8.3, 7.8, 4.5 Hz, 1H), 7.78 (d, *J* = 7.3 Hz, 1H), 7.73 (t, *J* = 8.9 Hz, 1H), 5.15 (dd, *J* = 12.8, 5.4 Hz, 1H), 2.88 (ddd, *J* = 17.1, 13.9, 5.5 Hz, 1H), 2.64 – 2.50 (m, 2H), 2.46 (d, *J* = 4.5 Hz, 0H), 2.10 – 2.01 (m, 1H).

**$^{13}$C NMR** (101 MHz, DMSO) δ 172.72, 169.67, 166.09 (d, *J* = 2.9 Hz), 163.95, 156.80 (d, *J* = 262.4 Hz), 138.04 (d, *J* = 7.9 Hz), 133.44, 122.99 (d, *J* = 19.5 Hz), 120.04 (d, *J* = 3.4 Hz), 117.03 (d, *J* = 12.6 Hz), 49.08, 30.89, 21.83.

##### 2-(2,6-dioxopiperidin-3-yl)-4-(4-(2-((2-fluoro-4-((2-fluoro-3-nitrobenzyl) sulfonyl) phenyl) thio)-5-methoxy-6-((5-methyl-1*H*-pyrazol-3-yl)amino)pyrimidin-4-yl)piperazin-1-yl)isoindoline-1,3-dione **D8**

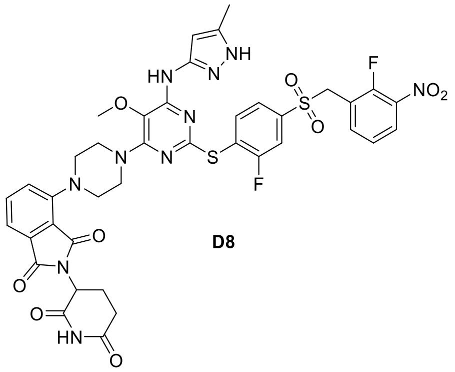


**D8**

Compound **18** (2.5 mg, 3.41 µmol, 1 eq) was dissolved in TFA and left for 2 h to allow deprotection. Upon completion, TFA was evaporated by the flow of nitrogen and the product **31** was used without further purification.

**31** (2 mg, 3.2 µmol, 1 eq) and **30** (1.14 mg, 4.1 µmol, 1.30 eq) were dissolved in anh. DMSO (0.5 mL) and DIPEA (4.1 mg, 5.5 µL, 32 µmol, 10 eq) was added. The reaction mixture was stirred at 90 °C for 4 h. Afterwards, the reaction was acidified and diluted with water. The final product was obtained by RP-HPLC (ACN/$H_2O$ step gradient 10-90%, 35 min) as yellow powder (0.7 mg, 25%).

**$^1$H NMR** (500 MHz, DMSO) δ 11.09 (s, 1H), 8.74 (s, 1H), 8.20 – 8.13 (m, 1H), 8.01 – 7.94 (m, 1H), 7.83 (dd, *J* = 8.0, 2.0 Hz, 1H), 7.76 – 7.69 (m, 1H), 7.67 – 7.60 (m, 2H), 7.45 (t, *J* = 8.0 Hz, 1H), 7.41 – 7.34 (m, 2H), 5.54 (s, 1H), 5.09 (dd, *J* = 12.8, 5.4 Hz, 1H), 5.01 (s, 2H), 3.75 – 3.69 (m, 4H), 3.59 (s, 3H), 3.38 – 3.32 (m, 4H), 2.91 – 2.82 (m, 1H), 2.61 – 2.54 (m, 2H), 2.05 (s, 3H), 2.01 (t, *J* = 5.3 Hz, 1H).

**$^{13}$C NMR** (126 MHz, DMSO) δ 172.79, 169.97, 167.03, 166.32, 161.65 (d, *J* = 251.4 Hz), 159.73, 154.51, 153.75, 153.59, 149.50, 140.64 (d, *J* = 6.7 Hz), 139.06 (d, *J* = 3.6 Hz), 138.03, 137.31 (d, *J* = 8.1 Hz), 135.92, 133.64, 127.00, 125.44 (d, *J* = 18.4 Hz), 124.95 (d, *J* = 4.7 Hz), 124.53 (d, *J* = 3.8 Hz), 123.79, 122.38, 118.92 (d, *J* = 14.3 Hz), 116.82, 115.69 (d, *J* = 26.4 Hz), 115.13, 95.20, 58.74, 54.09, 50.42, 48.81, 45.60, 30.96, 22.01, 11.04.

**HRMS** (ESI)**:** calc. for $C_{39}H_{35}F_2N_{10}O_9S_2$ $[M+H]^+$: 889.1998 ; found 889.1990

*tert*-butyl 4-(2-(2,6-dioxopiperidin-3-yl)-1,3-dioxoisoindolin-4-yl) piperazine-1-carboxylate **32**

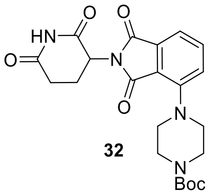


32

**31** (10 mg, 36.2 µmol, 1 eq) and **17** (13.5 mg, 72.4 µmol, 2 eq) were dissolved in anh. DMSO (0.5 mL) and DIPEA (46.8 mg, 63 µL, 362 µmol, 10 eq) was added. The reaction mixture was stirred at 90 °C for 4 h. Afterwards, the reaction was acidified and diluted with water. The final product **32** was obtained by RP-HPLC (ACN/$H_2O$ step gradient 10-90%, 35 min) as yellow powder (7 mg, 44%).

**$^1$H NMR** (400 MHz, DMSO) δ 11.08 (s, 1H), 7.71 (dd, *J* = 8.4, 7.2 Hz, 1H), 7.42 – 7.31 (m, 2H), 5.09 (dd, *J* = 12.9, 5.4 Hz, 1H), 3.50 (t, *J* = 4.9 Hz, 4H), 3.24 (t, *J* = 5.3 Hz, 4H), 2.93 – 2.81 (m, 1H), 2.63 – 2.51 (m, 2H), 2.06 – 1.97 (m, 1H), 1.42 (s, 9H).

**$^{13}$C NMR** (101 MHz, DMSO) δ 172.79, 169.96, 167.00, 166.31, 153.87, 149.57, 135.93, 133.59, 123.93, 116.98, 115.25, 79.07, 50.39, 48.81, 40.15, 39.94, 39.73, 39.52, 39.31, 39.10, 38.89, 30.94, 28.05, 22.03.

*tert*-butyl-(2-(4-(2-(2,6-dioxopiperidin-3-yl)-1,3-dioxoisoindolin-4-yl)piperazin-1-yl)-2-oxoethyl)carbamate **33**

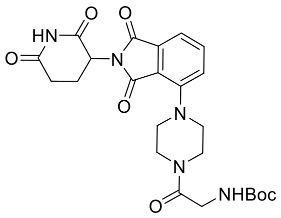


**32** (12.1 mg, 27.4 μmol, 1 eq) was dissolved in TFA and left for 2 h to allow deprotection. Upon completion, TFA was evaporated by the flow of nitrogen and the product **32a** was used without further purification.

(*tert*-butoxycarbonyl) glycine (4 mg, 23 μmol, 1 eq) was dissolved in anh. DMSO (0.5 mL), followed by the addition of *N,N*-diisopropylethylamine, DIPEA (59 mg, 80 μL, 80 μmol, 20 eq.) and PyBOP (14.3 mg, 27 μmol, 1.2 eq). This mixture was added into **32a** (9.4 mg, 27 μmol, 1.2 eq), dissolved in DMSO. After 2 h, the reaction was acidified by acetic acid and diluted with water. The final product **33** was obtained by RP-HPLC (ACN/$H_2O$ step gradient, 10-90%, 35 min) as yellow powder (4 mg, 35%).

**$^1$H NMR** (400 MHz, DMSO) δ 11.08 (s, 0H), 7.73 (dd, *J* = 8.4, 7.1 Hz, 1H), 7.40 (d, *J* = 7.1 Hz, 1H), 7.35 (d, *J* = 8.2 Hz, 1H), 6.78 (t, *J* = 6.0 Hz, 1H), 5.11 (dd, *J* = 12.9, 5.4 Hz, 1H), 3.84 (d, *J* = 5.9 Hz, 2H), 3.62 (d, 4H), 3.31 (s, 4H), 2.95 – 2.81 (m, 1H), 2.65 – 2.52 (m, 2H), 2.09 – 1.98 (m, 1H), 1.39 (s, 9H).

**$^{13}$C NMR** (101 MHz, DMSO) δ 172.78, 169.96, 167.59, 167.00, 166.35, 155.79, 149.38, 135.96, 133.60, 123.88, 116.96, 115.27, 77.91, 50.78, 50.11, 48.82, 43.90, 41.74, 41.30, 40.15, 39.94, 39.73, 39.52, 39.31, 39.10, 38.89, 30.94, 28.21, 22.03.

2-(2,6-dioxopiperidin-3-yl)-4-(4-(2-(1-(2-((2-fluoro-4-((2-fluoro-3-nitrobenzyl)sulfonyl) phenyl)thio)-5-methoxy-6-((5-methyl-1*H*-pyrazol-3-yl)amino)pyrimidin-4-yl)piperidin-4-yl) acetyl)piperazin-1-yl)isoindoline-1,3-dione **D9**

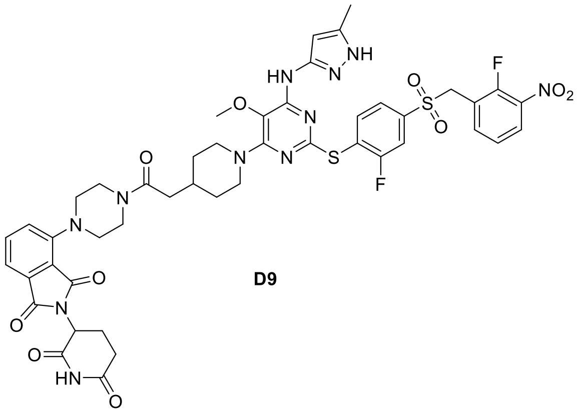


**D9**

General protocol **B** was followed. **D9** was obtained as a yellow powder (1.8 mg, 61%).

**$^1$H NMR** (500 MHz, DMSO) δ 11.09 (s, 1H), 8.66 (s, 1H), 8.20 – 8.13 (m, 1H), 7.99 – 7.95 (m, 1H), 7.80 (dd, *J* = 8.0, 1.9 Hz, 1H), 7.76 – 7.69 (m, 1H), 7.67 – 7.61 (m, 2H), 7.45 (t, *J* = 8.0 Hz, 1H), 7.39 (d, *J* = 7.1 Hz, 1H), 7.34 (d, *J* = 8.4 Hz, 1H), 5.57 (s, 1H), 5.11 (dd, *J* = 12.7, 5.5 Hz, 1H), 5.00 (s, 2H), 4.19 (d, *J* = 12.6 Hz, 2H), 3.67 – 3.63 (m, 4H), 3.53 (s, 3H), 3.31 – 3.22 (m, 4H), 2.92 – 2.86 (m, 1H), 2.84 – 2.78 (m, 2H), 2.65 – 2.54 (m, 2H), 2.31 (d, *J* = 6.7 Hz, 2H), 2.06 (s, 3H), 2.04 – 2.00 (m, 1H), 1.99 – 1.91 (m, 1H), 1.68 (d, *J* = 12.4 Hz, 2H), 1.20 – 1.12 (m, 2H).

**$^{13}$C NMR** (126 MHz, DMSO) δ 172.82, 169.99, 169.72, 167.03, 166.37, 161.63 (d, *J* = 251.4 Hz), 159.54, 153.59, 153.53, 153.45 (d, *J* = 265.6 Hz), 149.44, 146.51, 140.59 (d, *J* = 6.8 Hz), 139.06 (d, *J* = 3.7 Hz), 138.04, 137.32 (d, *J* = 7.9 Hz), 135.95, 133.62, 127.01, 125.53 (d, *J* = 18.2 Hz), 124.95 (d, *J* = 4.6 Hz), 124.44, 123.90, 121.97, 118.90 (d, *J* = 14.2 Hz), 116.92, 115.61 (d, *J* = 26.4 Hz), 115.25, 95.12, 58.53, 54.05, 51.04, 50.28, 48.83, 45.74, 45.15, 40.94, 32.83, 31.74, 30.96, 22.05, 11.06.

**HRMS** (ESI): calc. for $C_{46}H_{46}F_2N_{11}O_{10}S_2$ $[M+H]^+$: 1014.2838; found 1014.2828

*N*-(2-(4-(2-(2,6-dioxopiperidin-3-yl)-1,3-dioxoisoindolin-4-yl)piperazin-1-yl)-2-oxoethyl)-2-(1-(2-((2-fluoro-4-((2-fluoro-3-nitrobenzyl)sulfonyl)phenyl)thio)-5-methoxy-6-((5-methyl-1*H*-pyrazol-3-yl)amino)pyrimidin-4-yl)piperidin-4-yl)acetamide **D10**

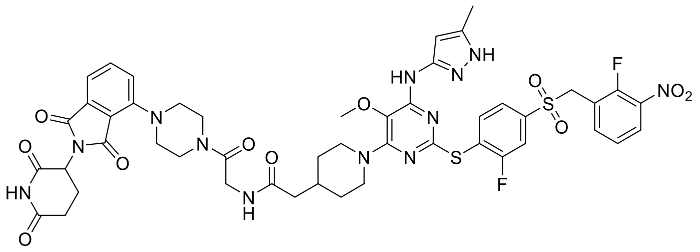

**D10**

General protocol **B** was followed. **D10** was obtained as a yellow powder (1.4 mg, 45%).

**$^1$H NMR** (500 MHz, DMSO) δ 11.09 (s, 1H), 8.66 (s, 1H), 8.19 – 8.15 (m, 1H), 7.99 – 7.95 (m, 2H), 7.81 (dd, J = 8.0, 1.9 Hz, 1H), 7.75 – 7.71 (m, 1H), 7.66 – 7.63 (m, 2H), 7.45 (t, J = 7.9 Hz, 1H), 7.40 (d, J = 7.1 Hz, 1H), 7.36 (d, J = 8.5 Hz, 1H), 5.56 (s, 1H), 5.12 (dd, J = 12.7, 5.5 Hz, 1H), 5.01 (s, 2H), 4.17 (d, J = 12.7 Hz, 2H), 4.00 (d, J = 5.6 Hz, 2H), 3.65 – 3.62 (m, 4H), 3.53 (s, 3H), 3.29 (d, J = 27.7 Hz, 4H), 2.92 – 2.84 (m, 1H), 2.83 – 2.77 (m, 2H), 2.62 – 2.54 (m, 2H), 2.10 (d, J = 7.1 Hz, 2H), 2.06 (s, 3H), 2.04 – 2.01 (m, 1H), 1.91 (s, 1H), 1.67 (d, J = 12.6 Hz, 2H), 1.19 – 1.11 (m, 2H).

**$^{13}$C NMR** (126 MHz, DMSO) δ 172.79, 171.17, 169.98, 167.28, 167.01, 166.36, 161.61 (d, J = 251.3 Hz), 159.53, 153.61, 153.50, 153.45 (d, J = 265.6 Hz), 149.39, 146.61, 140.55, 139.04, 138.01, 137.31 (d, J = 7.9 Hz), 135.96, 133.61, 127.00, 125.50 (d, J = 18.2 Hz), 124.93, 124.44, 123.88, 122.00, 118.90 (d, J = 14.5 Hz), 116.97, 115.62 (d, J = 26.5 Hz), 115.28, 95.10, 58.49, 54.04, 50.76, 50.14, 48.83, 45.73, 44.09, 42.13, 41.30, 40.37, 33.24, 31.55, 30.95, 22.04, 11.04.

**HRMS** (ESI)**:** calc. for $C_{48}H_{48}F_2N_{12}O_{11}S_2$ $[M+H]^+$:1071.3053; found 1071.3055

2-(2,6-dioxopiperidin-3-yl)-5-fluoroisoindoline-1,3-dione **35**

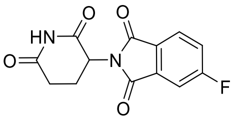

**35**

5-fluoroisobenzofuran-1,3-dione **34** (250 mg, 1.51 mmol, 1 eq) and 2,6-dioxopiperidin-3-aminium chloride **19** (248 mg, 1.51 mmol, 1 eq) were dissolved in acetic acid (6 mL), followed by addition of sodium-acetate (247 mg, 3.01 mmol, 2 eq). The reaction mixture was stirred under reflux for 24 h. After cooling to room temperature, the precipitate was filtered and washed with ice-cold water and diethyl-ether to obtain the desired product **35** as off-white powder (112 mg, 28 %).

$^{1}$**H NMR** (400 MHz, DMSO) δ 11.14 (s, 1H), 8.01 (dd, *J* = 8.3, 4.5 Hz, 1H), 7.85 (dd, *J* = 7.5, 2.3 Hz, 1H), 7.72 (ddd, *J* = 9.4, 8.2, 2.4 Hz, 1H), 5.16 (dd, *J* = 12.8, 5.4 Hz, 1H), 2.89 (ddd, *J* = 16.9, 13.8, 5.3 Hz, 1H), 2.64 – 2.52 (m, 2H), 2.47 (d, *J* = 4.5 Hz, 0H), 2.11 – 2.03 (m, 1H).

$^{13}$**C NMR** (101 MHz, DMSO) δ 172.73, 169.74, 167.25, 166.18, 165.89 (d, *J* = 2.9 Hz), 164.72, 134.20 (d, *J* = 9.9 Hz), 127.43 (d, *J* = 2.7 Hz), 126.27 (d, *J* = 9.8 Hz), 121.75 (d, *J* = 23.7 Hz), 111.43 (d, *J* = 25.3 Hz), 49.18, 30.91, 21.92.

*tert*-butyl-4-(2-(2,6-dioxopiperidin-3-yl)-1,3-dioxoisoindolin-5-yl)piperazine-1-carboxylate **36**

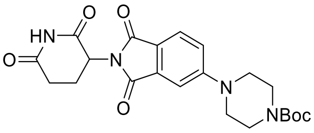


**36**

**35** (10.9 mg, 39.5 µmol, 1 eq) and **16** (14.7 mg, 78.9 µmol, 2 eq) were dissolved in anh. DMSO (0.5 mL) and DIPEA (51 mg, 69 µL, 395 µmol, 10 eq) was added. The reaction mixture was stirred at 90 °C for 4 h. Afterwards, the reaction was acidified and diluted by water. The final product **36** was obtained by RP-HPLC (ACN/$H_2O$ step gradient 10-90%, 35 min) as yellow powder (5.4 mg, 31%).

$^{1}$**H NMR** (500 MHz, DMSO) δ 11.07 (s, 1H), 7.68 (d, *J* = 8.5 Hz, 1H), 7.33 (d, *J* = 2.3 Hz, 1H), 7.23 (dd, *J* = 8.6, 2.4 Hz, 1H), 5.06 (dd, *J* = 12.8, 5.4 Hz, 1H), 2.87 (m, *J* = 16.7, 13.7, 5.4 Hz, 1H), 2.64 – 2.49 (m, 2H), 2.01 (m, *J* = 12.8, 5.7, 3.3 Hz, 1H), 1.41 (s, 9H).

$^{13}$**C NMR** (126 MHz, DMSO) δ 172.81, 170.07, 167.52, 166.97, 154.98, 153.85, 133.86, 124.94, 118.53, 117.91, 108.07, 79.17, 48.79, 46.58, 30.98, 28.07, 22.17.

2-(2,6-dioxopiperidin-3-yl)-5-(4-(2-(1-(2-((2-fluoro-4-((2-fluoro-3-nitrobenzyl) sulfonyl) phenyl) thio)-5-methoxy-6-((5-methyl-1*H*-pyrazol-3-yl) amino) pyrimidin-4-yl) piperidin-4-yl) acetyl) piperazin-1-yl) isoindoline-1,3-dione **D11**

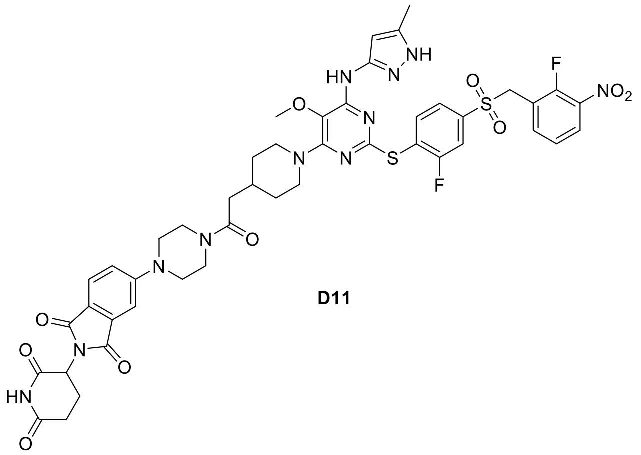


**D11**

General protocol **B** was followed. **D11** was obtained as a yellow powder (0.7 mg, 23%).

**$^{1}$H NMR** (500 MHz, DMSO) δ 11.08 (s, 1H), 8.57 (s, 1H), 8.17 (t, J = 7.4 Hz, 1H), 7.99 – 7.93 (m, 1H), 7.80 (dd, J = 8.0, 2.0 Hz, 1H), 7.70 (d, J = 8.5 Hz, 1H), 7.66 – 7.61 (m, 2H), 7.45 (t, J = 8.0 Hz, 1H), 7.36 – 7.32 (m, 1H), 7.27 – 7.21 (m, 1H), 5.55 (s, 1H), 5.08 (dd, J = 12.7, 5.6 Hz, 1H), 5.00 (s, 2H), 4.19 (d, J = 12.6 Hz, 2H), 3.65 – 3.56 (m, 4H), 3.53 (s, 3H), 2.93 – 2.74 (m, 2H), 2.61 – 2.53 (m, 1H), 2.31 (d, J = 6.9 Hz, 2H), 2.05 (s, 3H), 1.68 (d, J = 12.4 Hz, 2H), 1.19 – 1.14 (m, 2H).

**$^{13}$C NMR** (126 MHz, DMSO) δ 172.82, 170.09, 169.78, 167.54, 166.99, 161.86 (d, J = 241.3 Hz), 159.53, 154.89, 153.55, 140.57, 139.04, 138.03, 137.39, 133.85, 129.66, 127.01, 124.94, 124.43, 121.91, 118.48, 117.78, 115.61 (d, J = 27.0 Hz), 107.95, 95.10, 58.51, 54.06, 48.80, 46.88, 46.57, 45.75, 44.27, 35.12, 32.73, 31.75, 30.99, 22.19, 11.49.

**HRMS** (ESI)**:** calc. for $C_{46}H_{46}F_2N_{11}O_{10}S_2$ $[M+H]^+$: 1014.2838; found 1014.2834

*tert*-butyl-(*E*)-4-(4-(4-methoxyphenyl)-4-oxobut-2-enoyl)-1λ4-piperazine-1-carboxylate **41**[9]

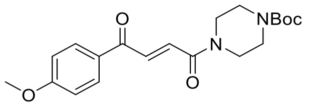


**41**

(*E*)-4-(4-methoxyphenyl)-4-oxobut-2-enoic (250 mg, 1.21 mmol, 1 eq) acid **40** was dissolved in anhydrous DMF (20 mL) and DIPEA (784 mg, 6.06 mmol, 5 eq) and 1-[Bis(dimethylamino)methylene]-1*H*-1,2,3-triazolo[4,5-b]pyridinium-3-oxidehexafluoro phosphate (HATU, 507 mg, 1.33 mmol, 1.10 eq) were added. After stirring for 30 min, 1-Boc piperazine (248.4 mg, 1.33 mmol, 1.10 eq) was added and reaction mixture was stirred at room temperature for 2h. Upon completion, the reaction mixture was washed with 5% LiCl solution and extracted with EtOAc (3x50 mL). The combined organic phase was dried over $MgSO_4$, vacuum filtered and concentrated. The product was obtained by flash column chromatography (pentane/EtOAc, gradient 0-50%) as a yellow powder (400 mg, 88%, purity: 90%).

**$^1$H NMR** (400 MHz, $CDCl_3$) δ 8.07 – 8.03 (m, 2H), 8.00 – 7.95 (m, 1H), 7.47 (d, *J* = 14.9 Hz, 1H), 7.01 – 6.96 (m, 2H), 3.90 (s, 3H), 3.75 – 3.70 (m, 2H), 3.65 – 3.60 (m, 2H), 3.52 – 3.47 (m, 4H), 1.49 (s, 9H).

(*E*)-1-(4-methoxyphenyl)-4-(piperazin-1-yl) but-2-ene-1,4-dione **42**[9]

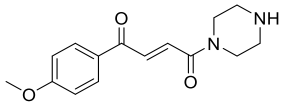


**42**

Product **41** (100 mg, 0.27 mmol, 1 eq) was dissolved in TFA (3 mL) and the mixture was stirred for 2 h at room temperature. The final product **42** was obtained by RP-HPLC (ACN/$H_2O$ gradient; 10-90%, 60 min) as white powder (45 mg, 61%).

**$^1$H NMR** (400 MHz, $CDCl_3$) δ 8.06 – 8.02 (m, 2H), 8.00 (d, *J* = 14.9 Hz, 1H), 7.40 (d, *J* = 14.9 Hz, 1H), 7.02 – 6.95 (m, 2H), 4.00 (d, *J* = 27.7 Hz, 4H), 3.90 (s, 3H).

**$^{13}$C NMR** (101 MHz, $CDCl_3$) δ 187.25, 164.56, 164.36, 136.22, 131.53, 129.77, 114.37, 55.77, 39.06.

(*E*)-1-(4-(2-((2-fluoro-4-((2-fluoro-3-nitrobenzyl) sulfonyl) phenyl) thio)-5-methoxy-6-((5-methyl-1H-pyrazol-3-yl) amino) pyrimidin-4-yl) piperazin-1-yl)-4-(4-methoxyphenyl) but-2-ene-1,4-dione **D12**

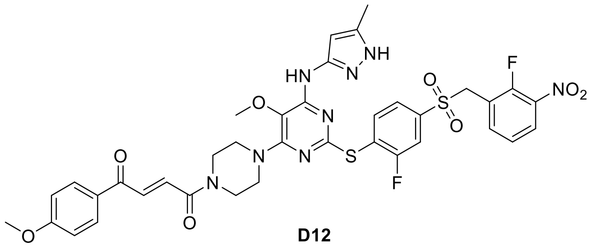


**D12**

Compound **18** (2 mg, 2.7 µmol, 1 eq) was dissolved in TFA and left for 2h at room temperature to allow deprotection. Upon completion, TFA was evaporated by the flow of nitrogen and the obtained product **31** was used without further purification.

(*E*)-4-(4-methoxyphenyl)-4-oxobut-2-enoic acid **40** (0.7 mg, 3.2 µmol, 1.2 eq) was dissolved in anh. DMSO (0.5 mL), followed by addition of DIPEA (8.7 mg, 12 µL, 67 µmol, 25 eq) and TSTU (1.1 mg, 3.6 µmol, 1.3 eq). **31** (1.7 mg, 2.7 µmol, 1 eq), dissolved in anh. DMSO, was added to this mixture and, after 2 h, the reaction was acidified by acetic acid and diluted with water. The final product **D12** was obtained by RP-HPLC (ACN/$H_2O$ gradient; 10-90%, 25 min) as yellow powder (1.5 mg, 69%).

**$^1$H NMR** (500 MHz, DMSO) δ 8.84 (s, 1H), 8.20 – 8.13 (m, 1H), 8.07 – 8.00 (m, 2H), 7.99 – 7.95 (m, 1H), 7.83 (dd, J = 8.1, 1.9 Hz, 1H), 7.78 (d, J = 15.1 Hz, 1H), 7.68 – 7.60 (m, 2H), 7.48 – 7.38 (m, 2H), 7.11 – 7.07 (m, 2H), 5.56 (s, 1H), 5.01 (s, 2H), 3.87 (s, 3H), 3.57 (s, 3H), 2.06 (s, 3H).

**$^{13}$C NMR** (126 MHz, DMSO) δ 187.52, 163.73, 163.66, 161.61 (d, J = 251.5 Hz), 153.71, 153.59, 153.46 (d, J = 265.6 Hz), 146.38, 140.68 (d, J = 6.7 Hz), 139.07, 137.97, 137.29 (d, J = 7.9 Hz), 133.48, 132.64, 131.16, 129.44, 127.00, 125.35 (d, J = 18.4 Hz), 124.93 (d, J = 4.8 Hz), 124.52, 122.46, 118.89 (d, J = 14.2 Hz), 115.70 (d, J = 26.5 Hz), 114.31, 95.27, 58.80, 55.64 (d, J = 8.3 Hz), 54.04, 45.95, 45.35, 41.49, 11.01.

**HRMS** (ESI)**:** calc. for $C_{37}H_{35}F_2N_8O_8S_2$ $[M+H]^+$: 821.1987; found 821.1981

(*E*)-1-(4-(2-(1-(2-((2-fluoro-4-((2-fluoro-3-nitrobenzyl)sulfonyl)phenyl)thio)-5-methoxy-6-((5-methyl-1H-pyrazol-3-yl)amino)pyrimidin-4-yl)piperidin-4-yl)acetyl)piperazin-1-yl)-4-(4-methoxyphenyl)but-2-ene-1,4-dione **D13**

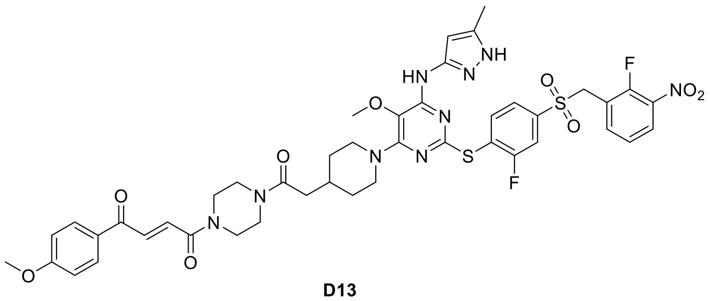


**D13**

Compound **42** (1.31 mg, 4.8 µmol, 1.50 eq) was dissolved in anh. DMSO (0.3 mL) and DIPEA (8.3 mg, 12 µL, 63.8 µmol, 20 eq) and PyBOP (2.0 mg, 3.8 µmol, 1.1 eq) were added.

After 15 minutes, **16** (2.2 mg, 3.2 μmol, 1 eq) was added to the reaction mixture. The reaction was left for 2 h at room temperature and afterwards acidified with acetic acid (50 μL) and diluted with water (20 μL). The final product **D13** was obtained by RP-HPLC (ACN/$H_2O$ step gradient, 10-90%, 35 min) as a pale yellow solid (1.38 mg, 46%).

**$^1$H NMR** (400 MHz, DMSO) δ 8.77 (s, 1H), 8.20 – 8.14 (m, 1H), 8.07 – 8.02 (m, 2H), 7.99 – 7.93 (m, 1H), 7.82 – 7.76 (m, 2H), 7.67 – 7.61 (m, 2H), 7.48 – 7.39 (m, 2H), 7.09 (d, J = 8.6 Hz, 2H), 5.61 (s, 1H), 5.01 (s, 2H), 4.18 (d, J = 12.6 Hz, 2H), 3.87 (s, 3H), 3.53 (s, 3H), 2.82 (t, J = 12.5 Hz, 2H), 2.30 – 2.25 (m, 2H), 2.07 (s, 3H), 1.95 (s, 1H), 1.66 (d, J = 12.5 Hz, 2H), 1.20 – 1.09 (m, 2H).

**$^{13}$C NMR** (126 MHz, DMSO) δ 187.56, 169.73, 163.74, 161.62 (d, J = 251.7 Hz), 159.53, 153.58, 153.44 (d, J = 266.2 Hz), 153.39, 146.23, 140.60 (d, J = 6.8 Hz), 139.80, 139.04, 138.02, 137.31 (d, J = 8.1 Hz), 133.54 (d, J = 11.2 Hz), 132.63 (d, J = 18.3 Hz), 131.18, 129.44, 127.00, 125.50 (d, J = 18.4 Hz), 124.93, 124.43, 122.01, 118.89 (d, J = 14.0 Hz), 115.60 (d, J = 26.4 Hz), 114.32, 95.13, 58.57, 55.67, 54.02, 45.70, 45.24 (d, J = 18.2 Hz), 44.64, 41.89, 41.36 (d, J = 25.7 Hz), 40.61, 39.02, 32.68, 31.70, 11.03.

**HRMS** (ESI)**:** calc. for $C_{44}H_{46}F_2N_9O_9S_2$ $[M+H]^+$: 946.2828; found 946.2821

4-((2-(2-((tert-butoxycarbonyl) amino) ethoxy) ethyl) carbamoyl)-2-(6-(dimethylamino)-3-(dimethyliminio)-3H-xanthen-9-yl) benzoate **44**

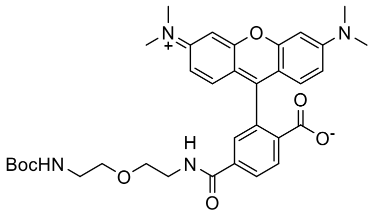


**44**

Compound **43** (20 mg, 46.5 μmol, 1 eq) was dissolved in anh. DMSO (0.5 mL), followed by the addition of DIPEA (120 mg, 162 μL, 0.93 mmol, 20 eq.) and PyBOP (29 mg, 55.8 μmol, 1.2 eq). This mixture was added into **23a** (14.2 mg, 69.7 μmol, 1.5 eq), dissolved in DMSO. After 2h, the reaction was acidified by acetic acid (50 μL) and diluted with water (30 μL). The final product was obtained by RP-HPLC (ACN/$H_2O$ step gradient, 30-90%, 35 min), as a pink powder (9.80 mg, 40%).

**¹H NMR** (400 MHz, DMSO) δ 8.81 (t, *J* = 5.5 Hz, 1H), 8.33 – 8.21 (m, 2H), 7.88 (d, *J* = 1.8 Hz, 1H), 7.11 – 7.01 (m, 4H), 6.97 (s, 2H), 6.75 (t, *J* = 5.5 Hz, 1H), 3.51 (t, *J* = 5.8 Hz, 2H), 3.43 (t, *J* = 5.5 Hz, 2H), 3.38 (t, *J* = 6.1 Hz, 2H), 3.27 (s, 12H), 3.11 – 2.97 (m, 2H), 1.34 (s, 9H).

**¹³C NMR** (126 MHz, DMSO) δ 166.37, 165.04, 157.28, 156.05, 138.04, 133.60, 131.53, 131.17, 129.49, 129.24, 115.12, 113.29, 96.75, 78.06, 69.38, 68.97, 46.34, 46.31, 40.99, 28.67, 26.41, 26.35.

2-(3,6-bis(dimethylamino) xanthylium-9-yl)-4-((2-(2-(2-(1-(2-((2-fluoro-4-((2-fluoro-3-nitrobenzyl) sulfonyl) phenyl) thio)-5-methoxy-6-((5-methyl-1H-pyrazol-3-yl) amino) pyrimidin-4-yl) piperidin-4-yl) acetamido) ethoxy) ethyl) carbamoyl) benzoate **45**

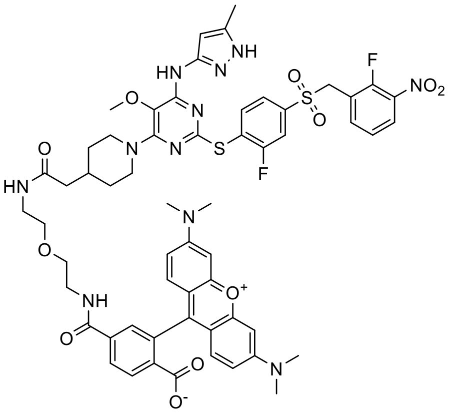


**45**

Compound **16** (2.10 mg, 3.0 μmol, 1 eq) was dissolved in anh. DMSO (0.5 mL), followed by the addition of DIPEA (7.9 mg, 11 μL, 60.9 μmol, 20 eq.) and PyBOP (1.9 mg, 3.7 μmol, 1.2 eq). This mixture was added into **44** (3.15 mg, 6.1 μmol, 2 eq), dissolved in DMSO. After 2h, the reaction was acidified by acetic acid (50 μL) and diluted with water (30 μL). The final product was obtained by RP-HPLC (ACN/$H_2O$ step gradient, 30-90%, 35 min), as a pink powder (1.3 mg, 36%).

**¹H NMR** (500 MHz, DMSO) δ 8.80 (s, 1H), 8.57 (s, 1H), 8.30 – 8.22 (m, 2H), 8.17 (s, 1H), 7.99 – 7.92 (m, 1H), 7.90 (s, 1H), 7.85 – 7.77 (m, 2H), 7.64 (s, 2H), 7.47 – 7.42 (m, 1H), 7.04 (s, 4H), 6.94 (s, 2H), 5.53 (s, 1H), 5.00 (s, 2H), 4.13 (d, *J* = 13.1 Hz, 2H), 3.25 (d, *J* = 3.8 Hz, 12H), 2.77 – 2.69 (m, 1H), 2.07 – 2.02 (m, 3H), 1.97 – 1.91 (m, 2H), 1.58 – 1.51 (m, 2H), 1.11 – 1.00 (m, 2H).

$^{13}$**C NMR** (126 MHz, DMSO) δ 171.04, 164.64, 153.57 (d, *J* = 5.6 Hz), 153.45 (d, *J* = 265.5 Hz), 140.58 (d, *J* = 6.8 Hz), 139.05, 137.99, 137.28, 130.51, 129.03, 127.01, 125.53 (d, *J* = 18.0 Hz), 124.95 (d, *J* = 4.3 Hz), 124.47, 121.93, 118.91 (d, *J* = 14.3 Hz), 118.37, 115.99, 115.64 (d, *J* = 26.1 Hz), 110.15, 96.41, 95.06, 68.85, 58.46, 54.07, 46.63, 45.69, 42.29, 40.46, 38.34, 33.14, 31.57, 25.95, 22.52, 11.04.

**HRMS** (ESI): calc. for $C_{58}H_{59}F_2N_{11}O_{11}S_2$ $[M+H]^+$ 1188.3883; found 1188.3848

#### 6. Full western blots

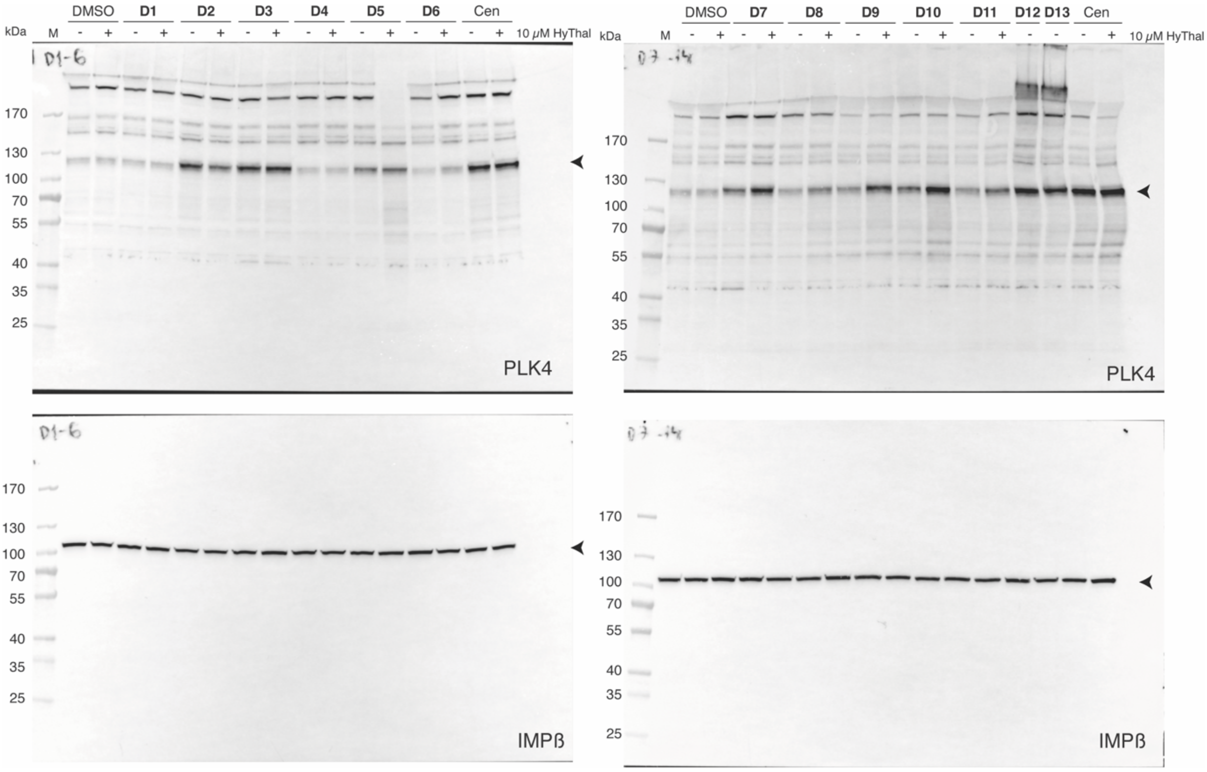


Figure 3A

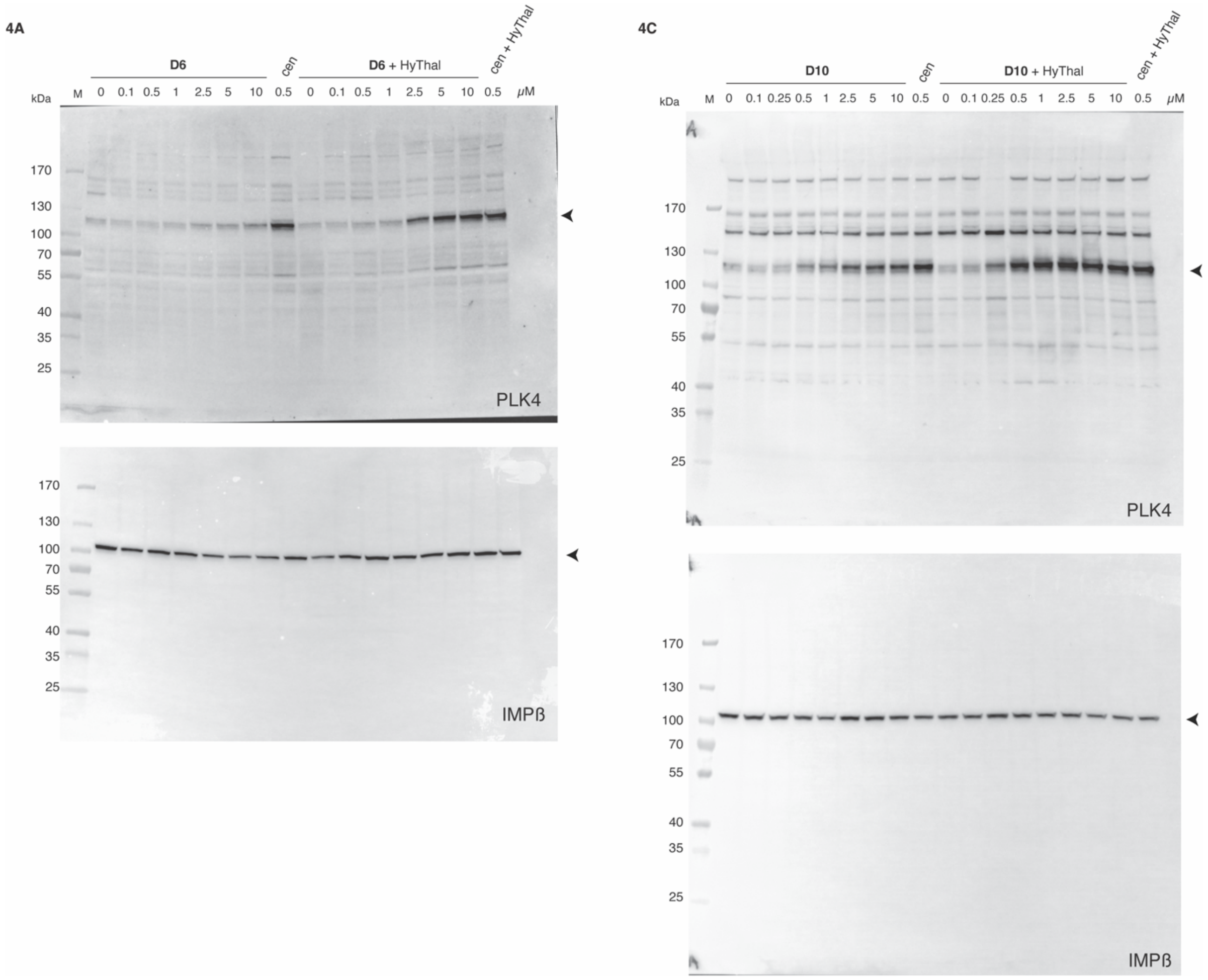


Figure 4A and 4C

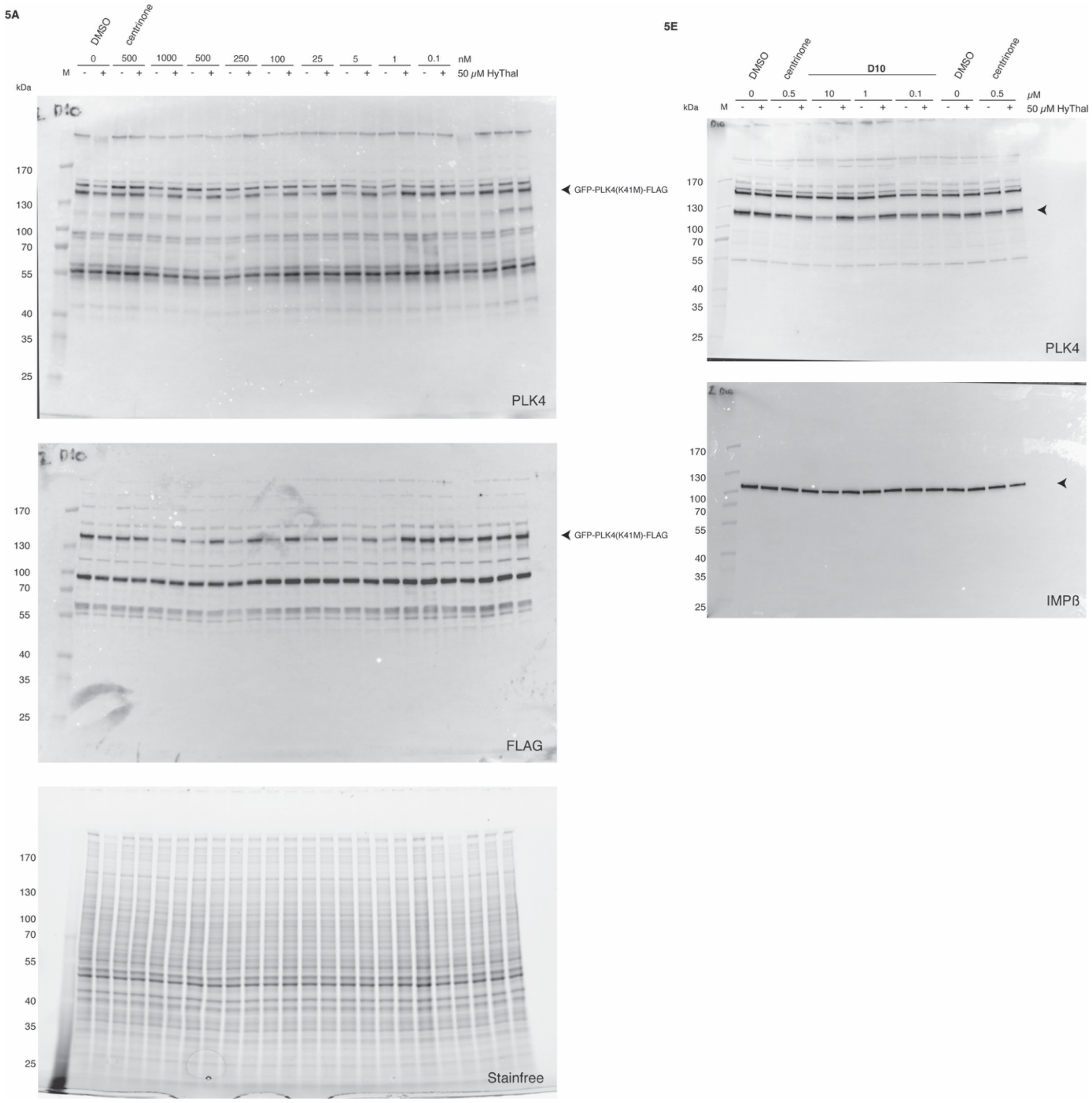


Figure 5A and 5E

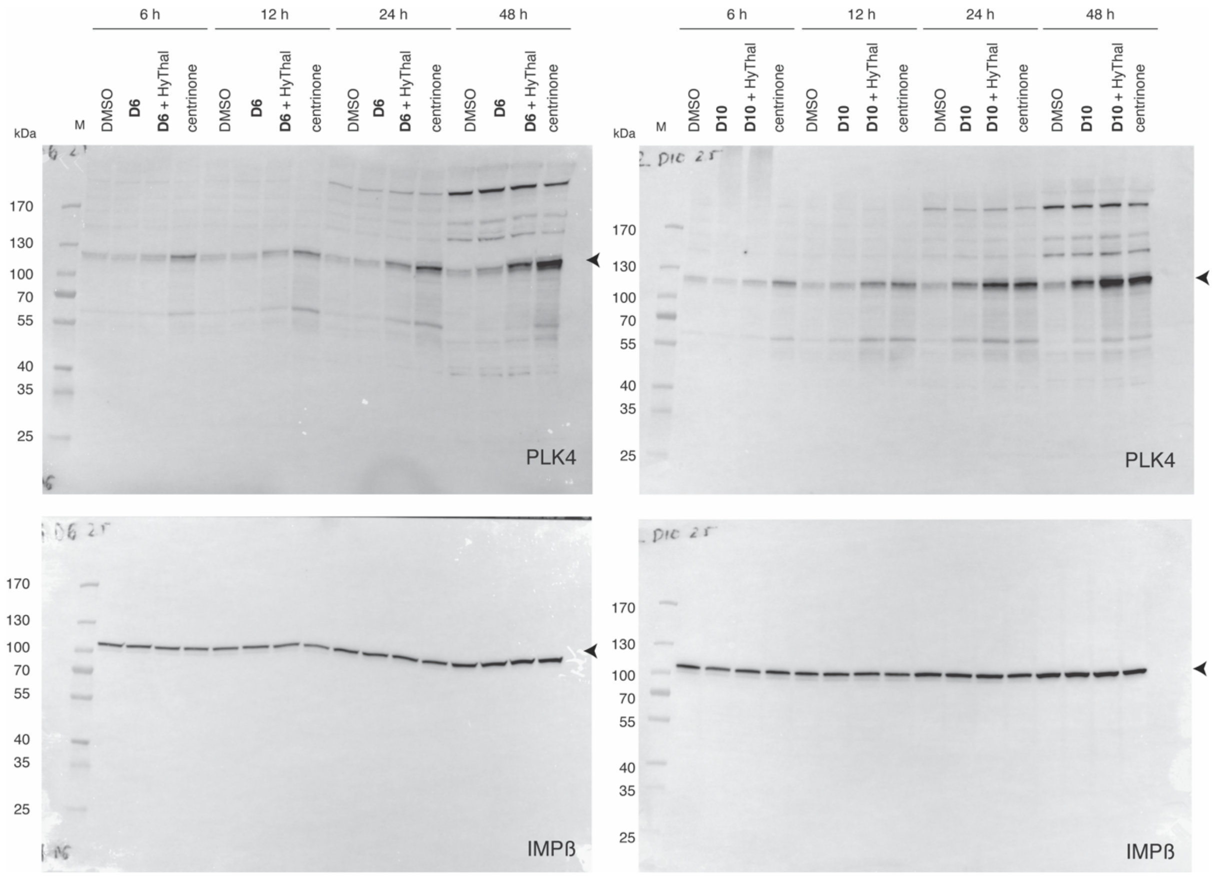

Supplementary Figure 1

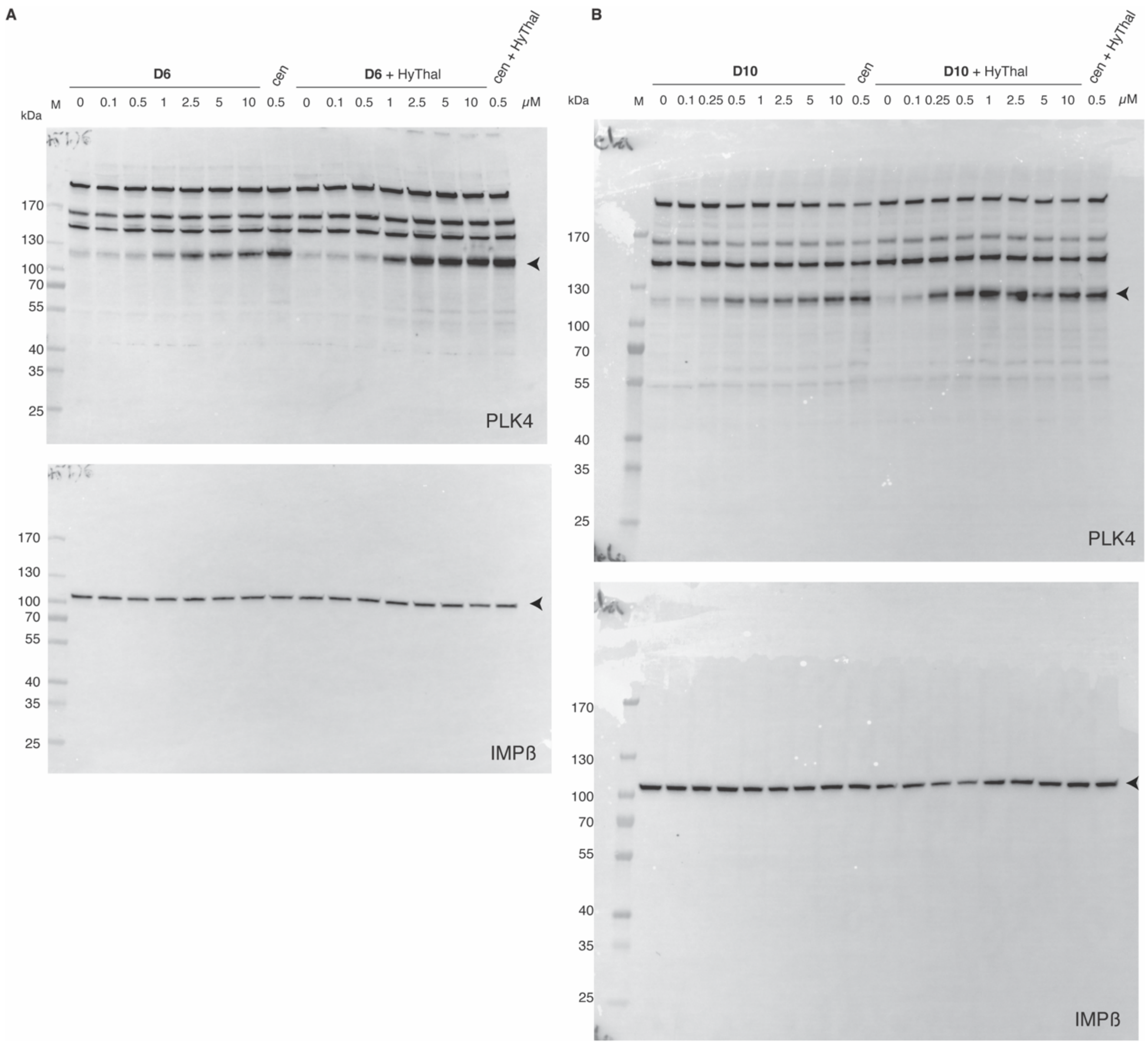


Supplementary Figure 2

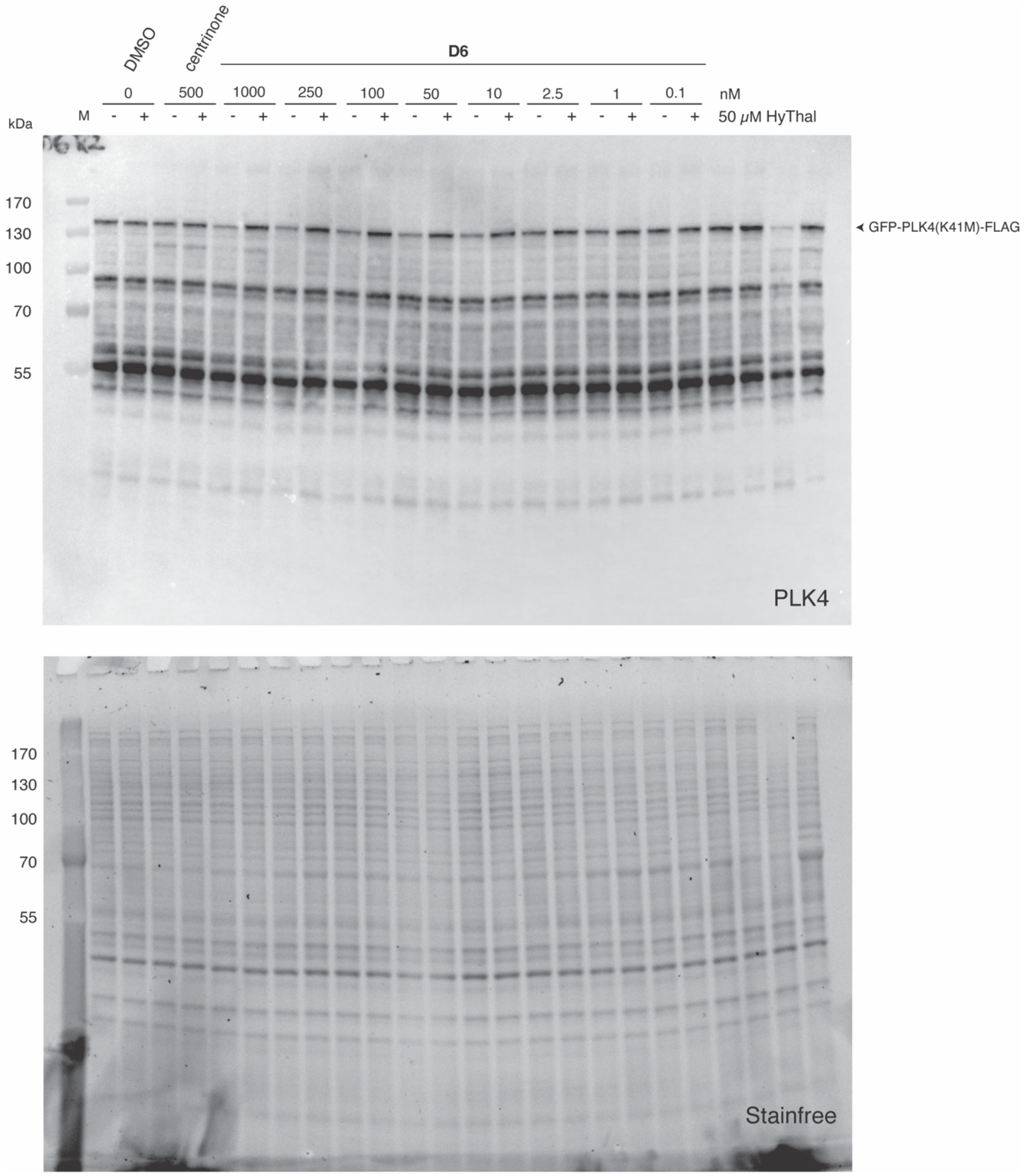


Supplementary Figure 4A

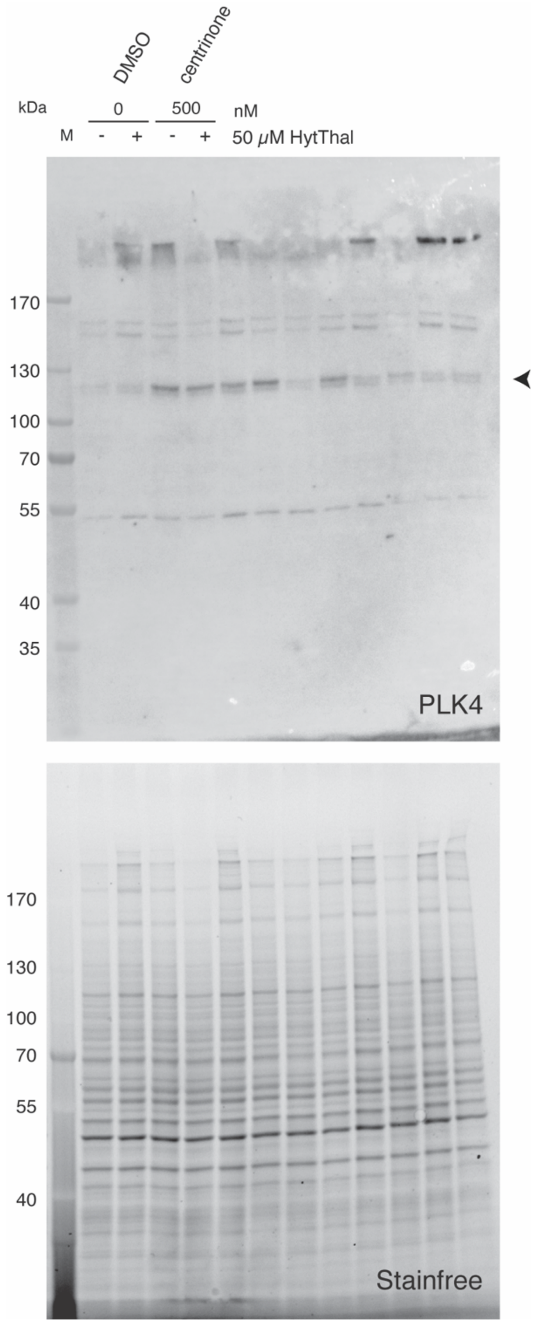

Supplementary Figure 5A

### 7. NMR Spectra

#### 4,6-Dihydroxy-5-methoxy thiopyrimidine **2**

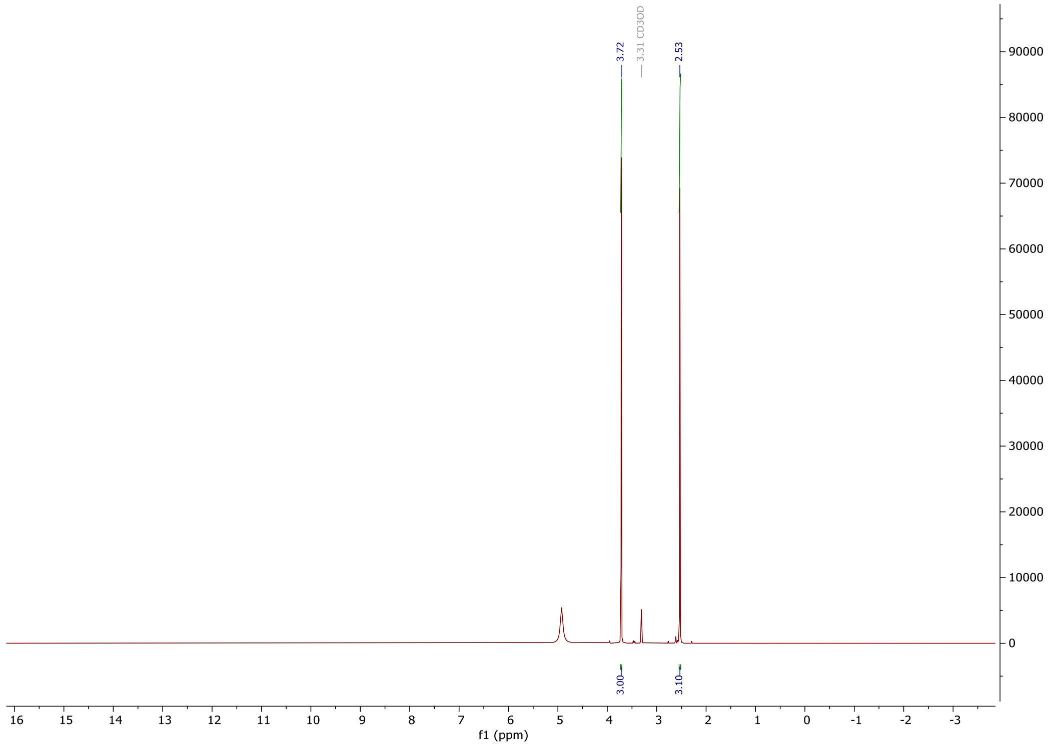


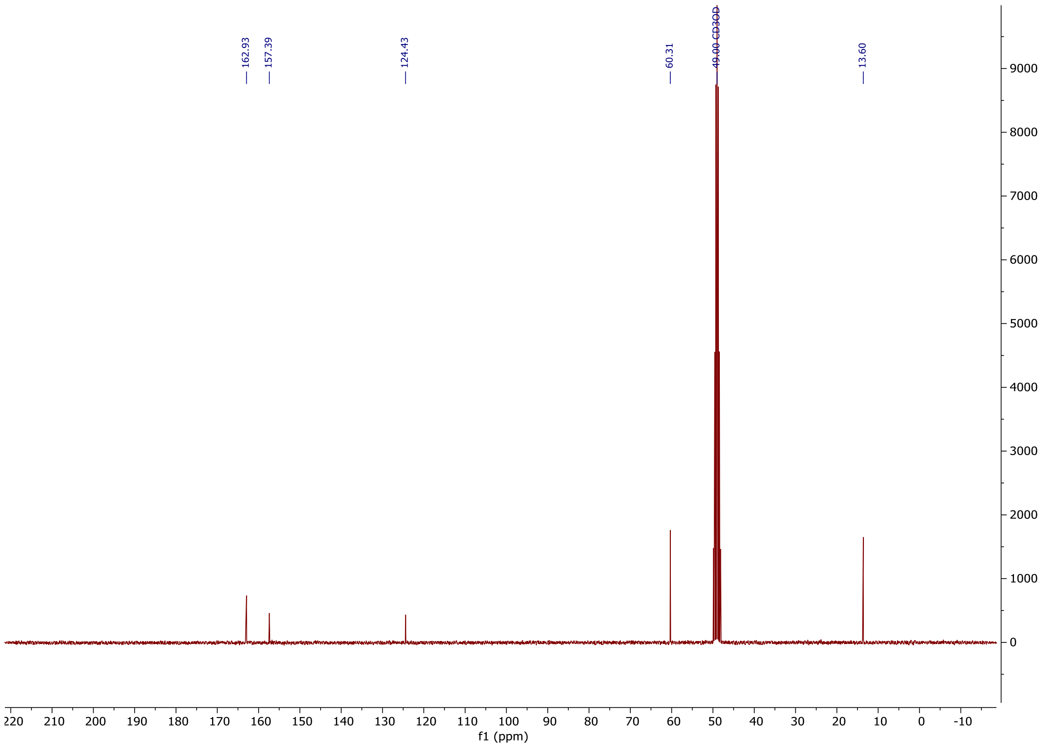

### 4,6-Dichloro-5-methoxy thiopyirimidine **3**

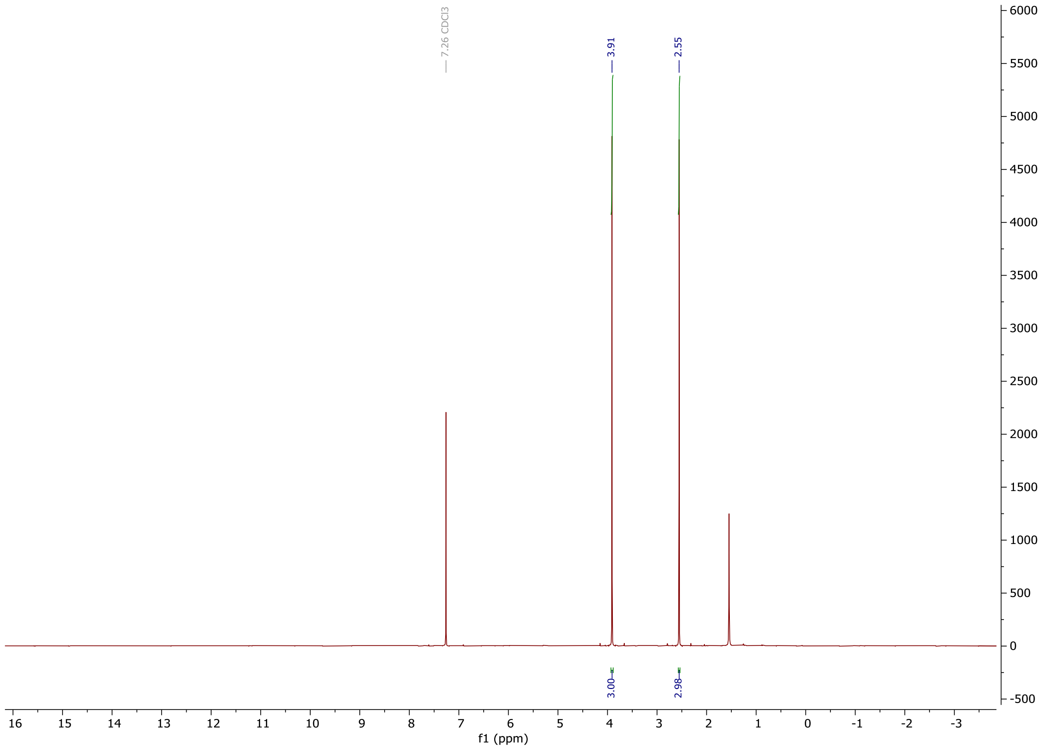


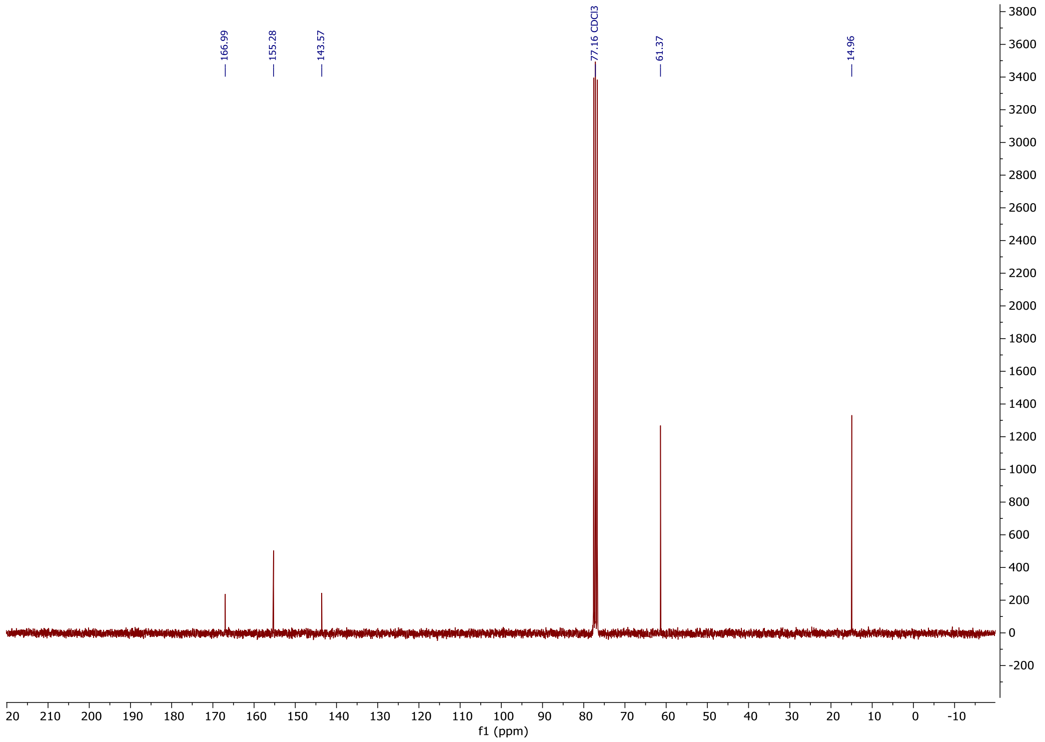

6-Chloro-5-methoxy-4-((methyl-1H-pyrazol-3-yl)amino)-2-(methylthio)pyrimidine **5**

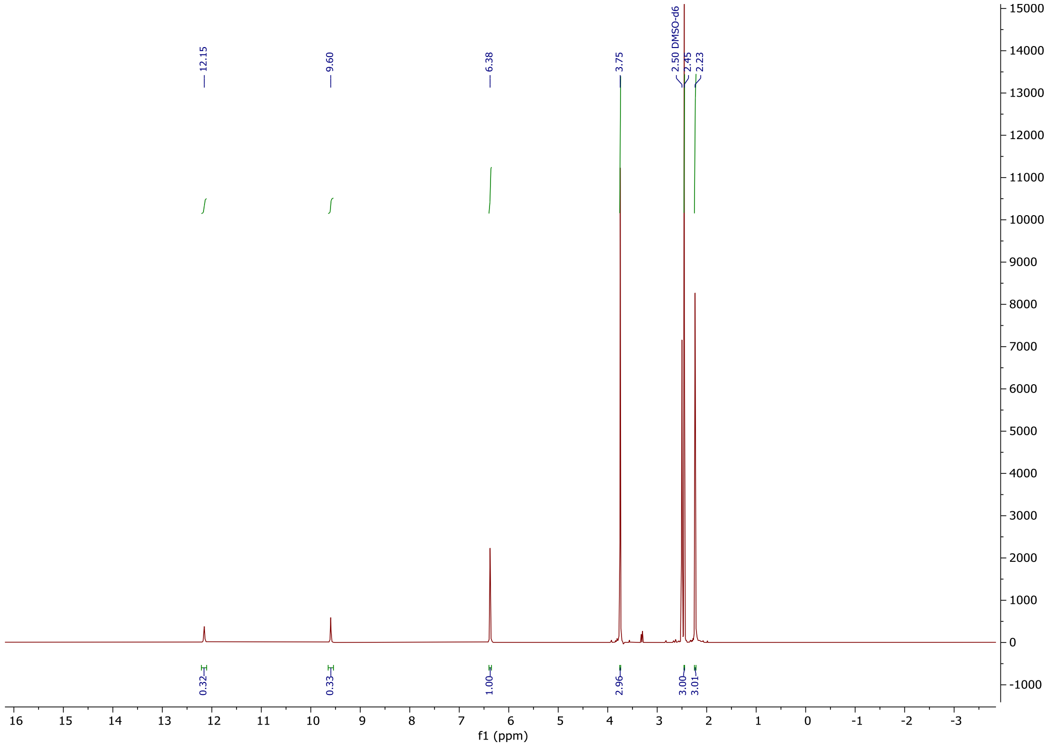


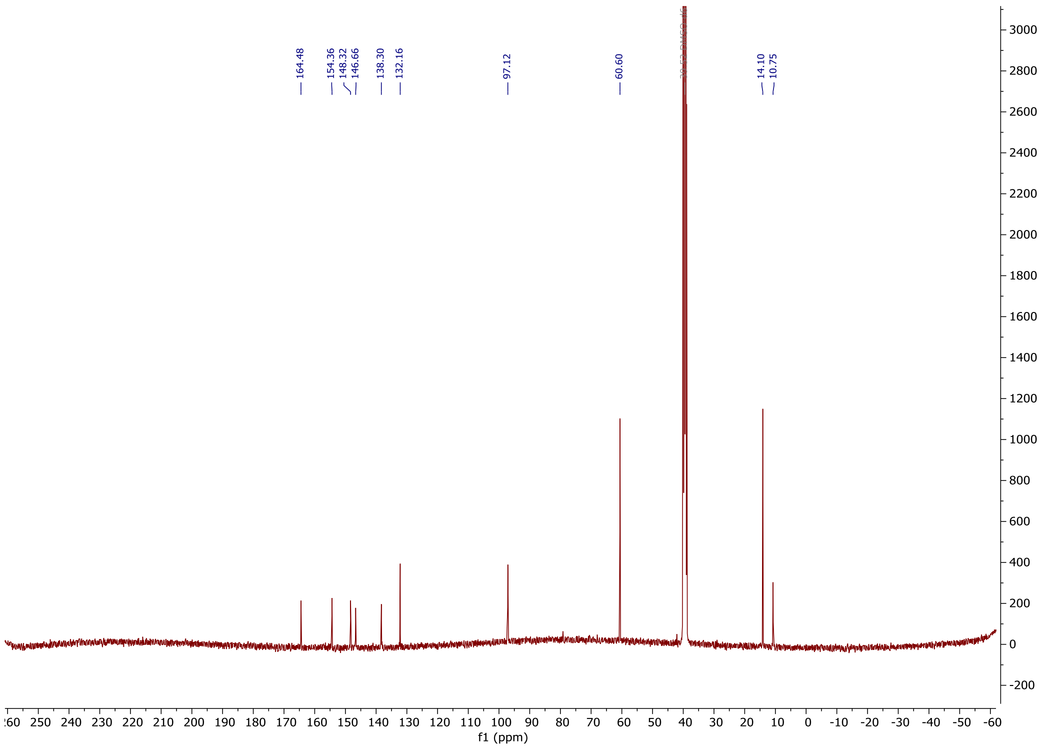

#### 6-Chloro-5-methoxy-4-((5-methyl-1H-pyrazol-3-yl)amino)-2-(methylsulfonyl) pyrimidine **6**

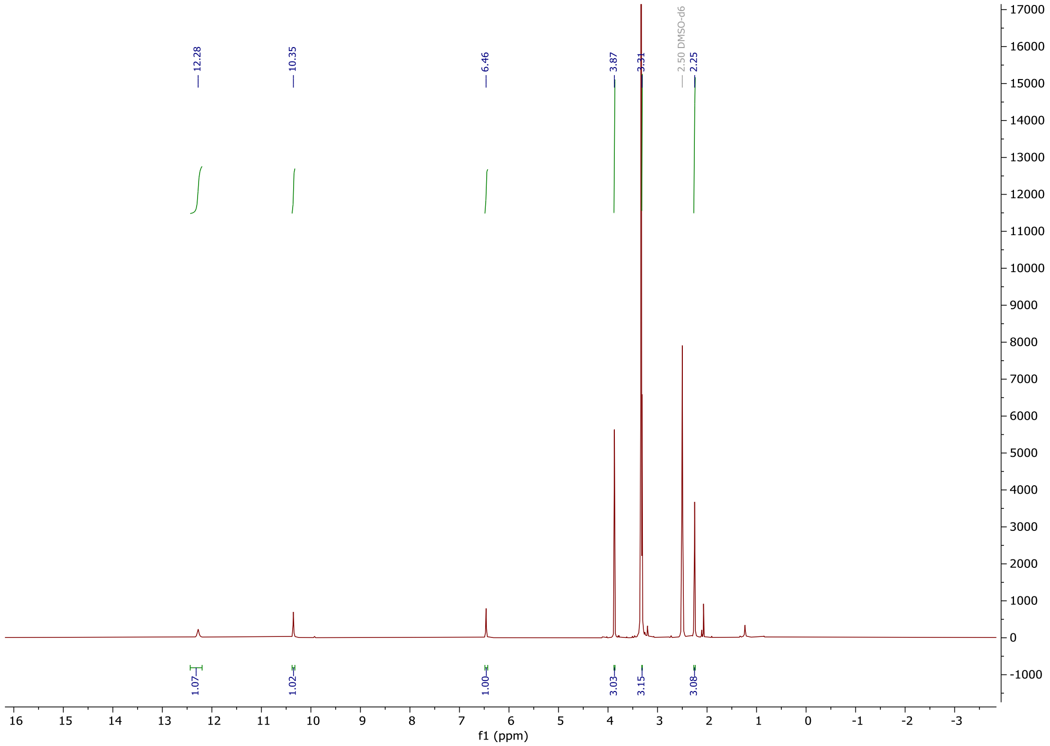


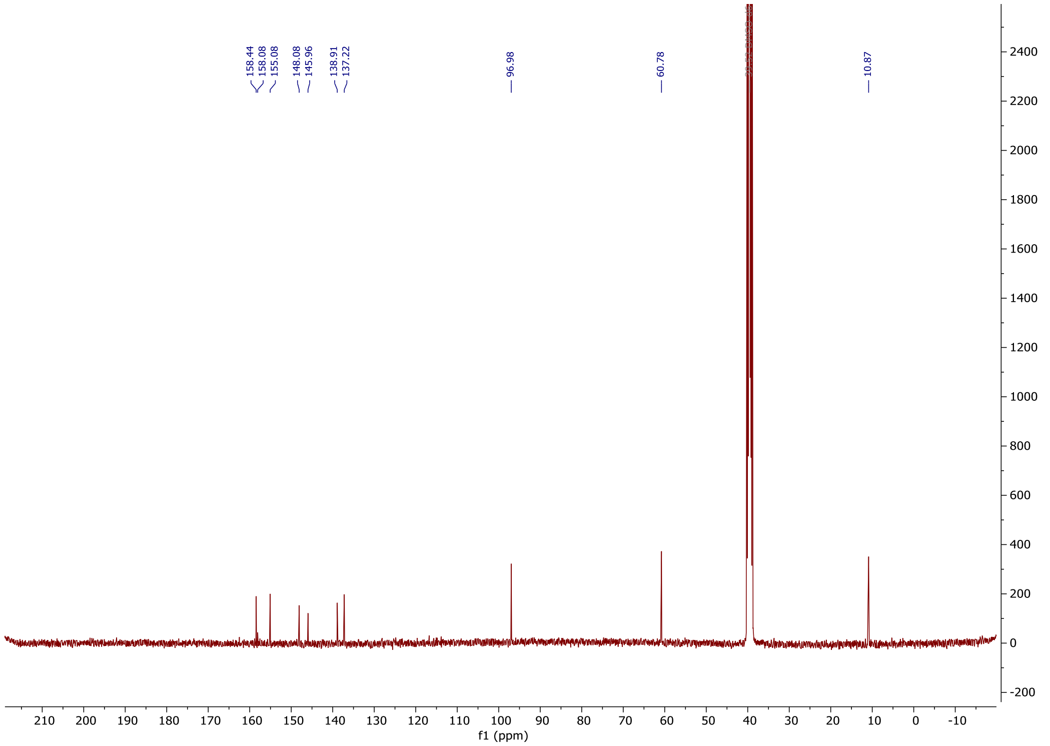

4-Bromo-2-fluoro-1-(tert-butylthio)benzene **8**

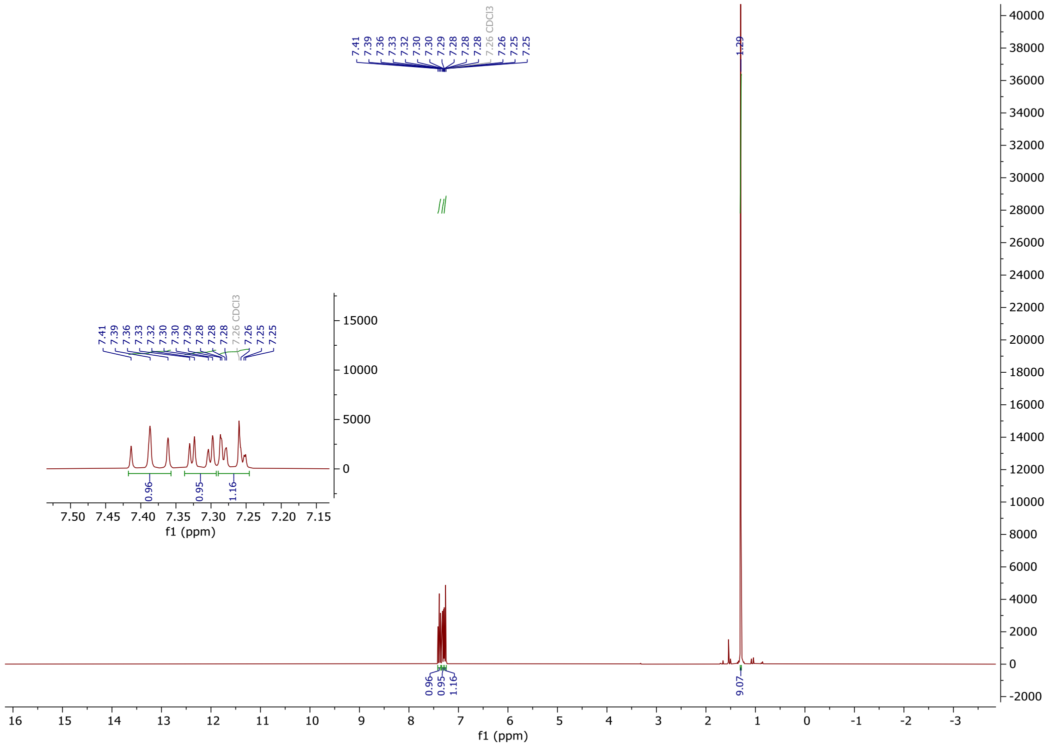


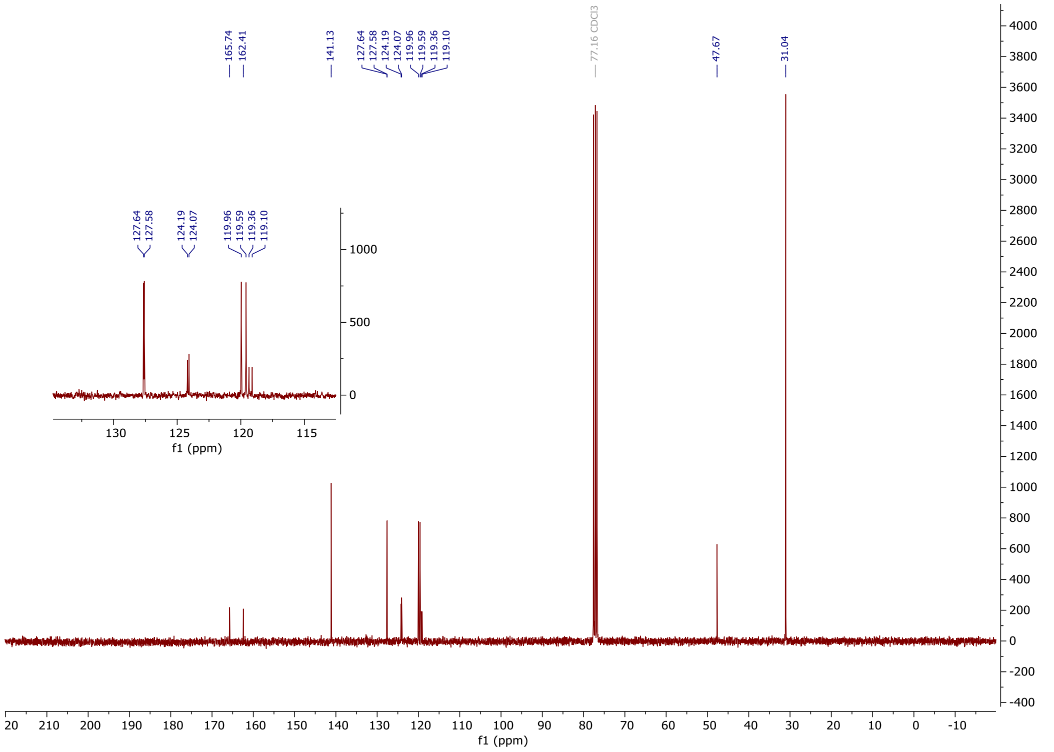

1-(Bromomethyl)-2-fluoro-3-nitrobenzene **11**

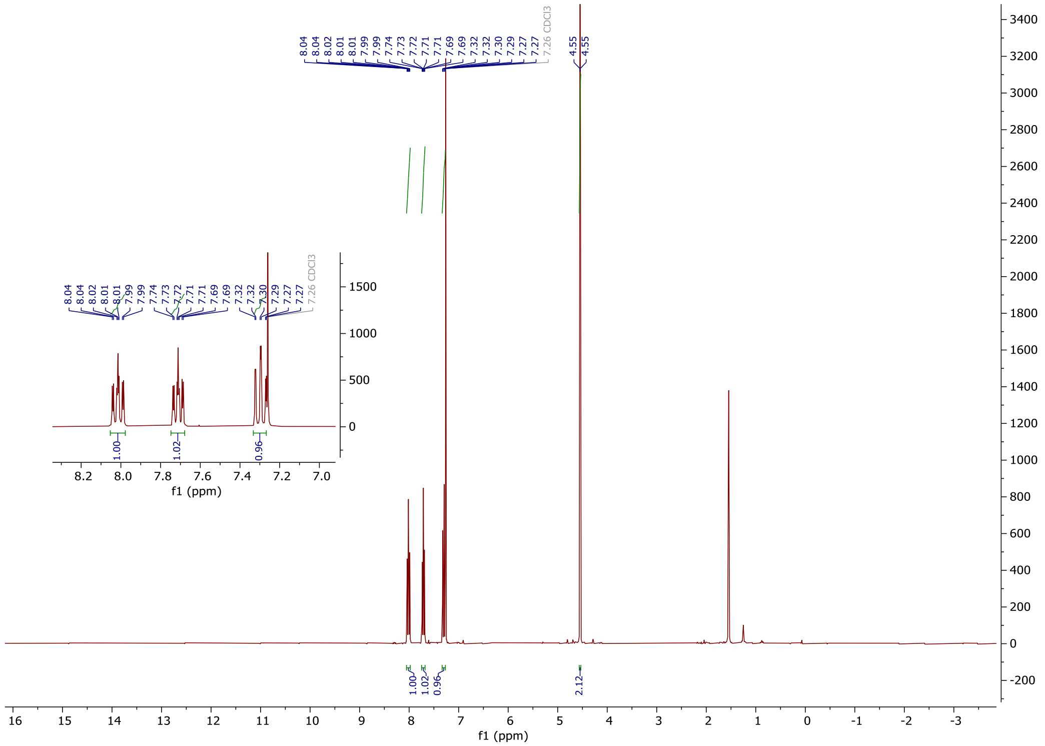

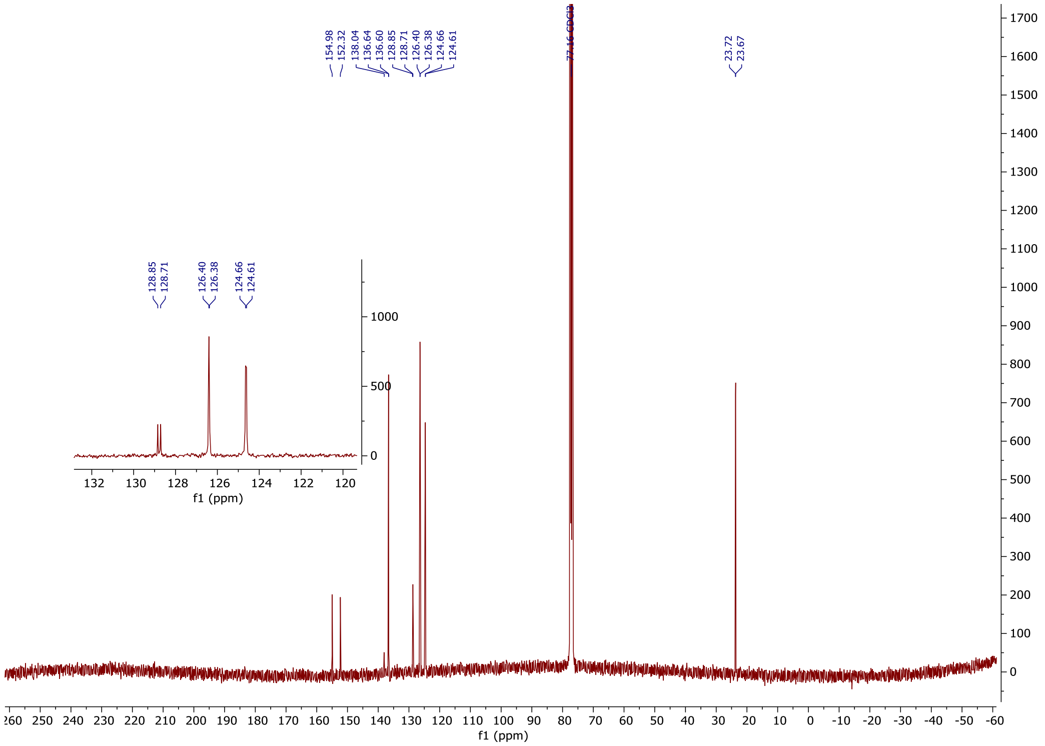

tert-Butyl (2-fluoro-4-((2-fluoro-3-nitrobenzyl) sulfonyl) phenyl) sulfane **12**

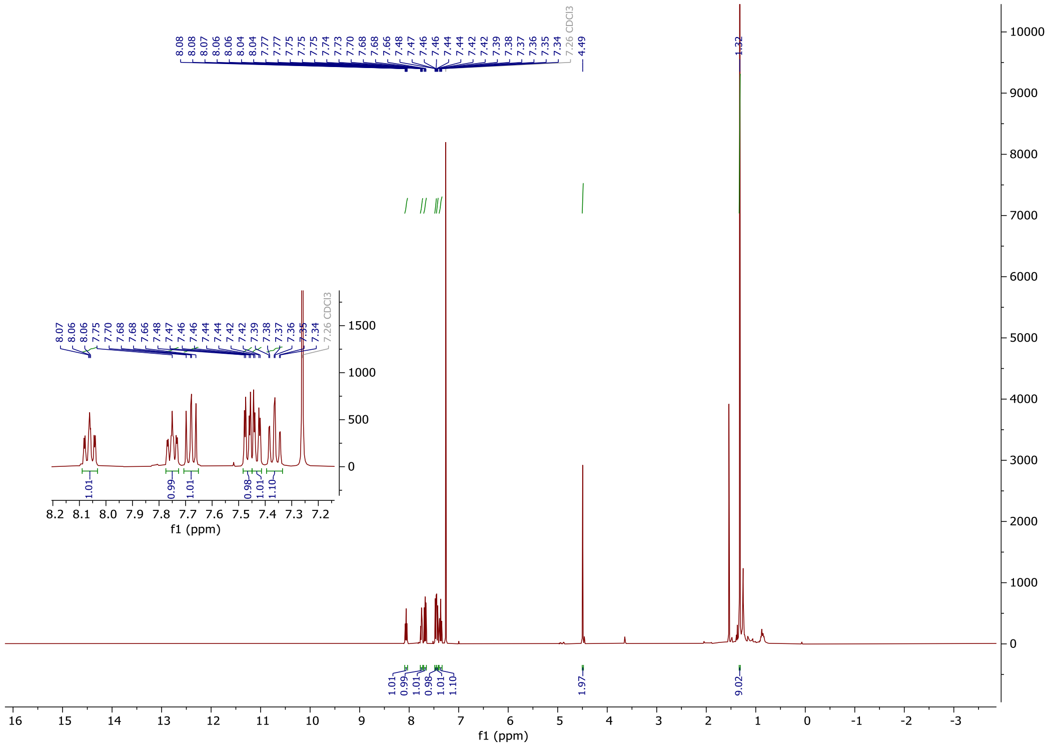


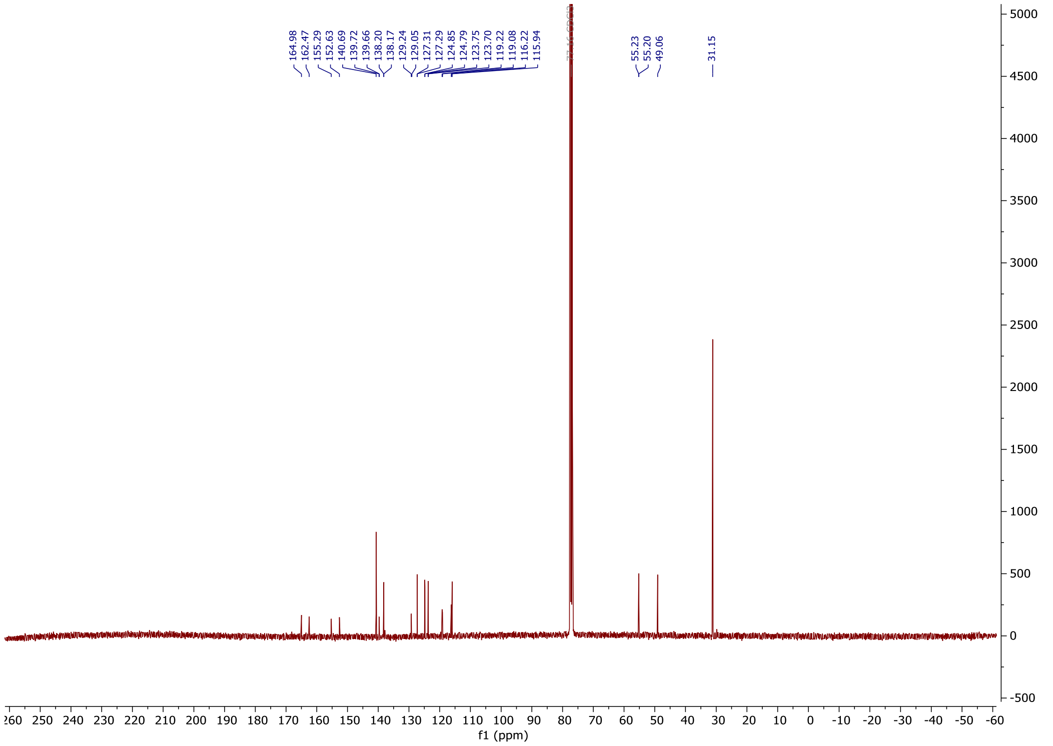

6-chloro-2-((2-fluoro-4-((2-fluoro-3-nitrobenzyl)sulfonyl)phenyl)thio)-5-methoxy-4-((5-methyl-1H-pyrazol-3-yl)amino) pyrimidine **14**

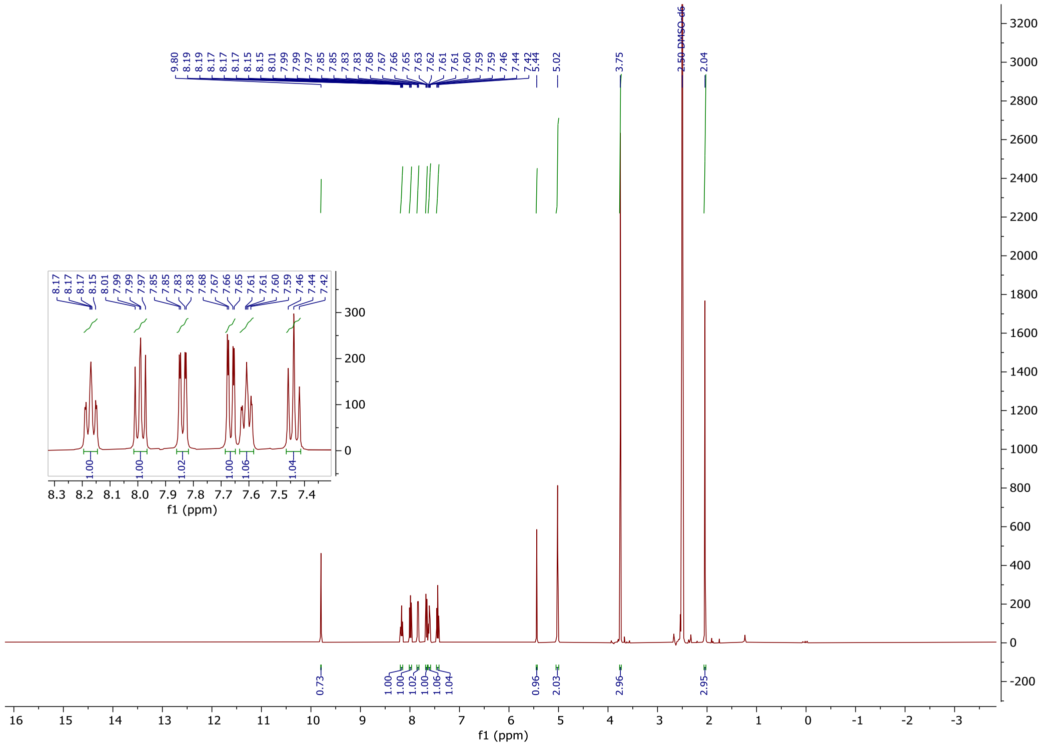


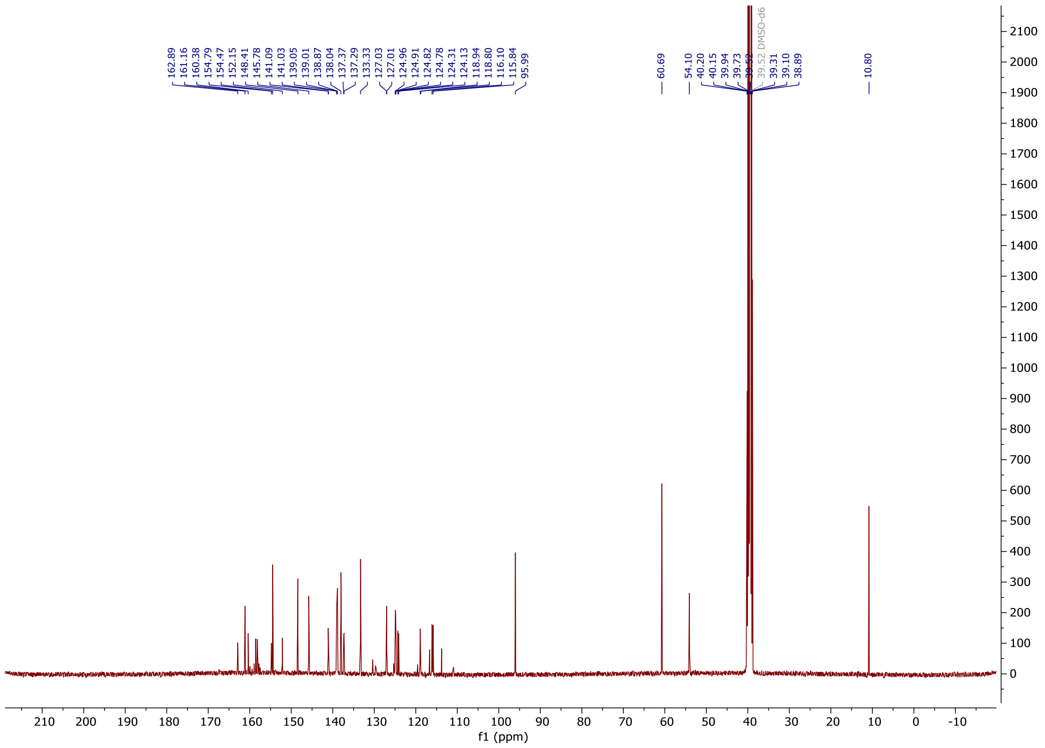

2-(1-(2-((2-fluoro-4-((2-fluoro-3-nitrobenzyl)sulfonyl)phenyl)thio)-5-methoxy6-((5-methyl-1H-pyrazol-3-yl)amino)pyrimidin-4-yl)piperidin-4-yl) acetic acid **16**

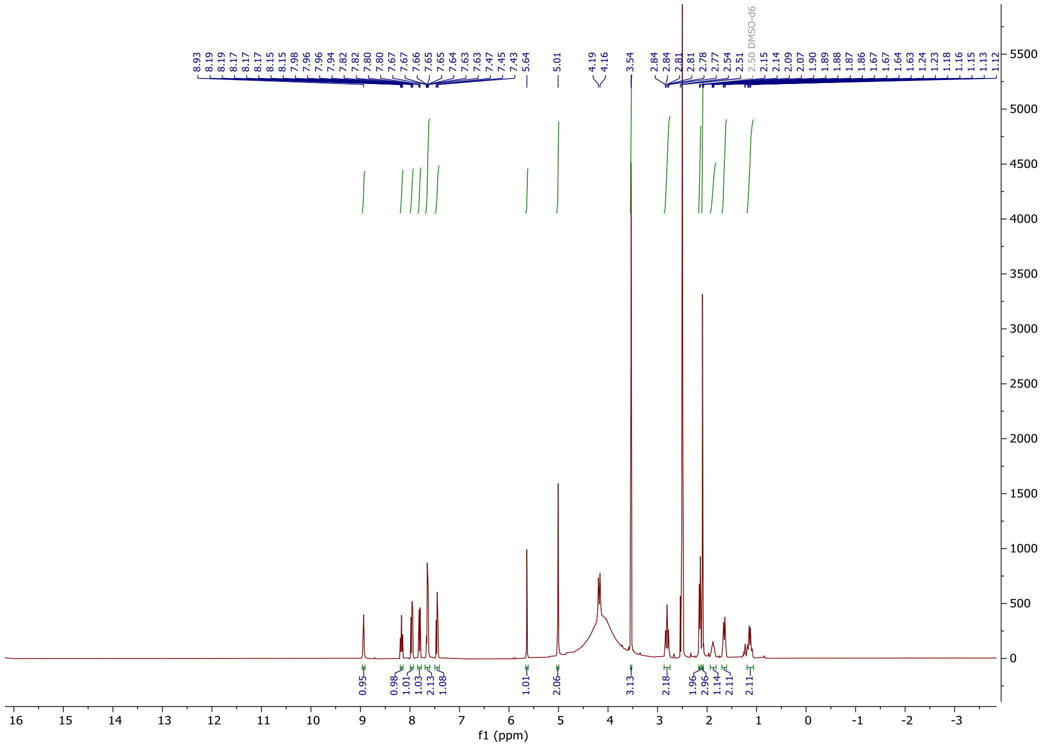

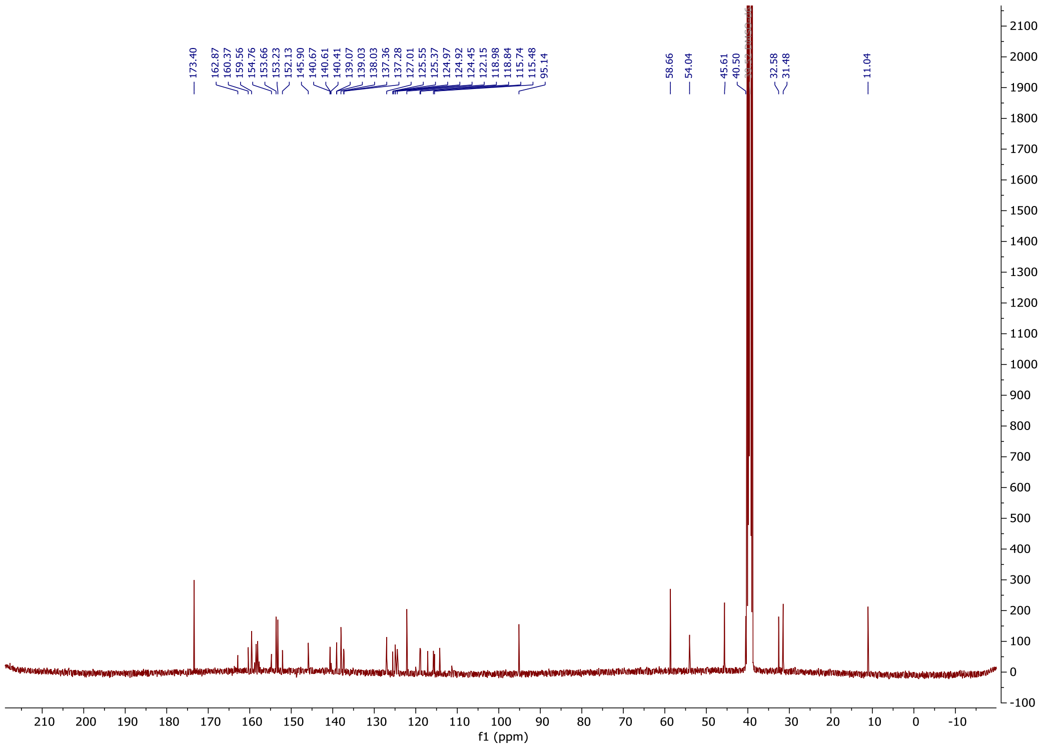

*tert*-butyl-4-(2-((2-fluoro-4-((2-fluoro-3-nitrobenzyl)sulfonyl)phenyl)thio)-5-methoxy-6-((5-methyl-1H-pyrazol-3-yl)amino)pyrimidin-4-yl)piperazine-1-carboxylate **18**

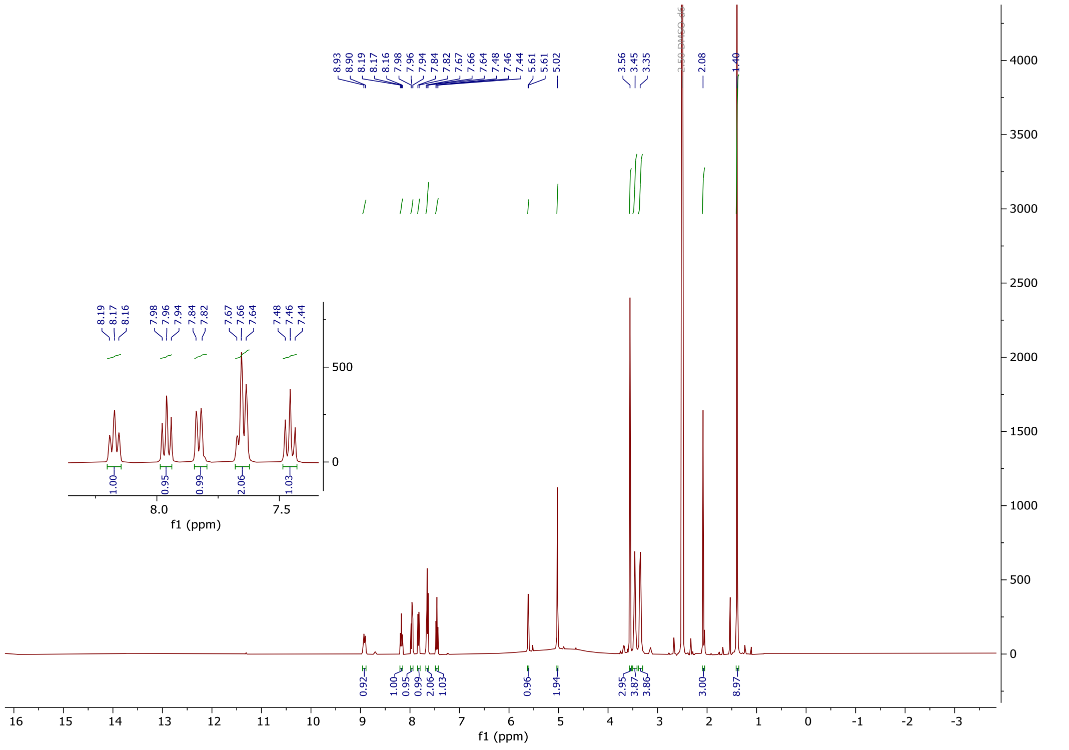


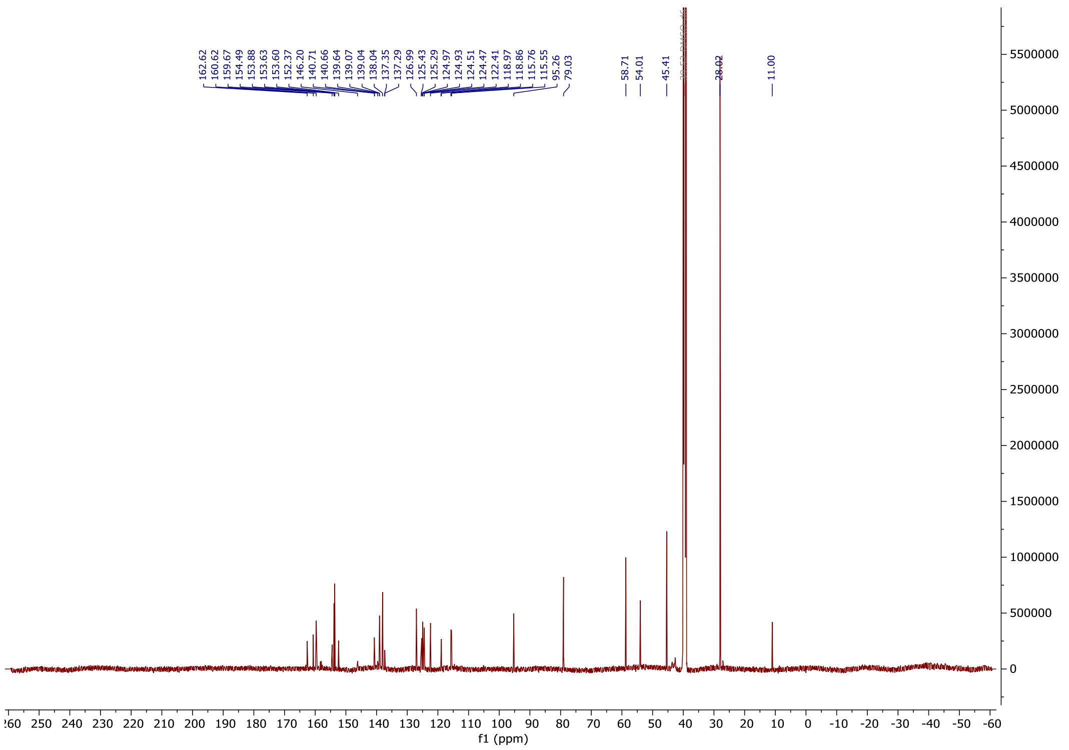

2-(2,6-dioxopiperidin-3-yl)-4-hydroxyisoindoline-1,3-dione **21**

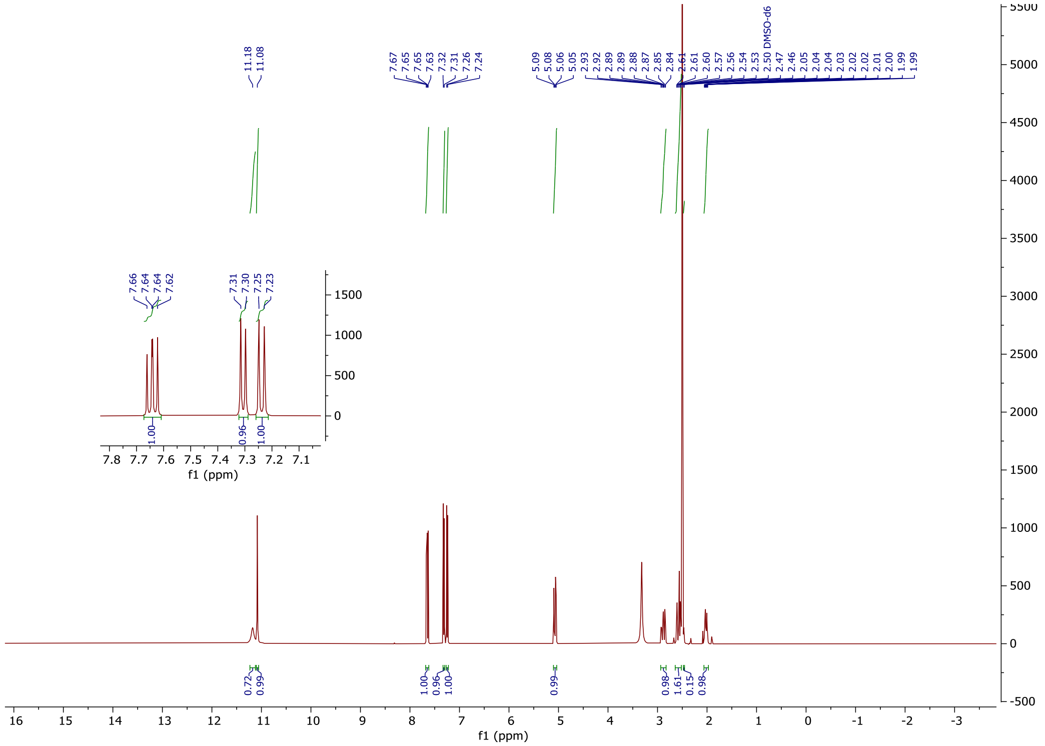


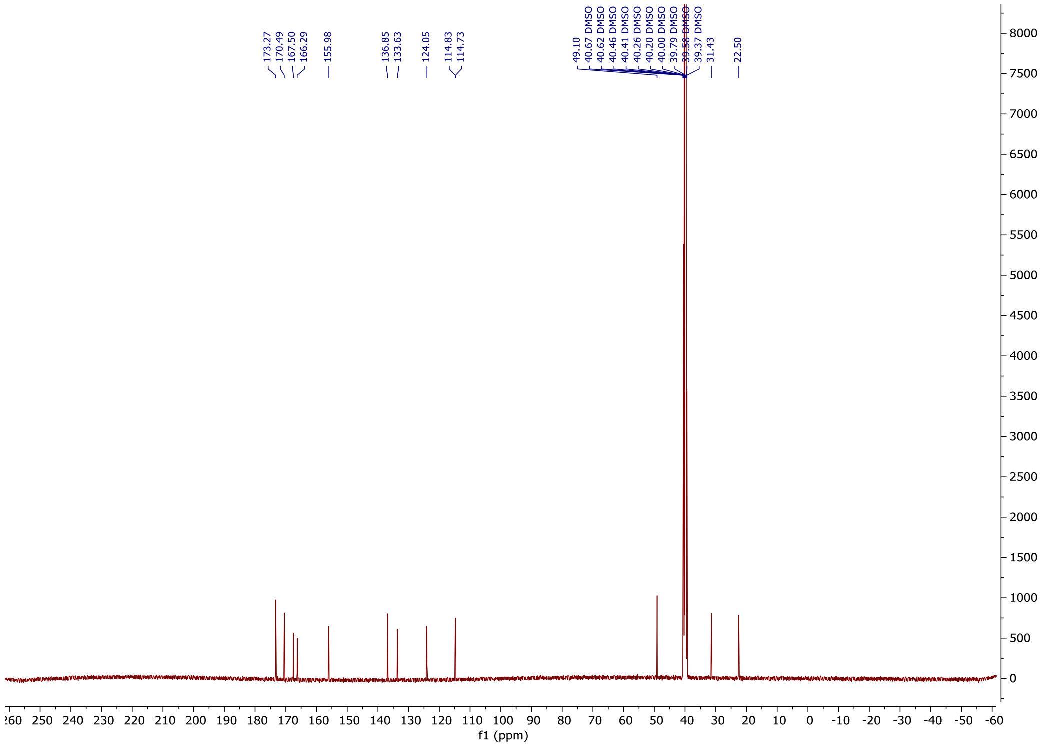

tert-butyl (2-(2-((2-(2,6-dioxopiperidin-3-yl)-1,3-dioxoisoindolin-4-yl)oxy)acetamido)ethyl)carbamate **22**

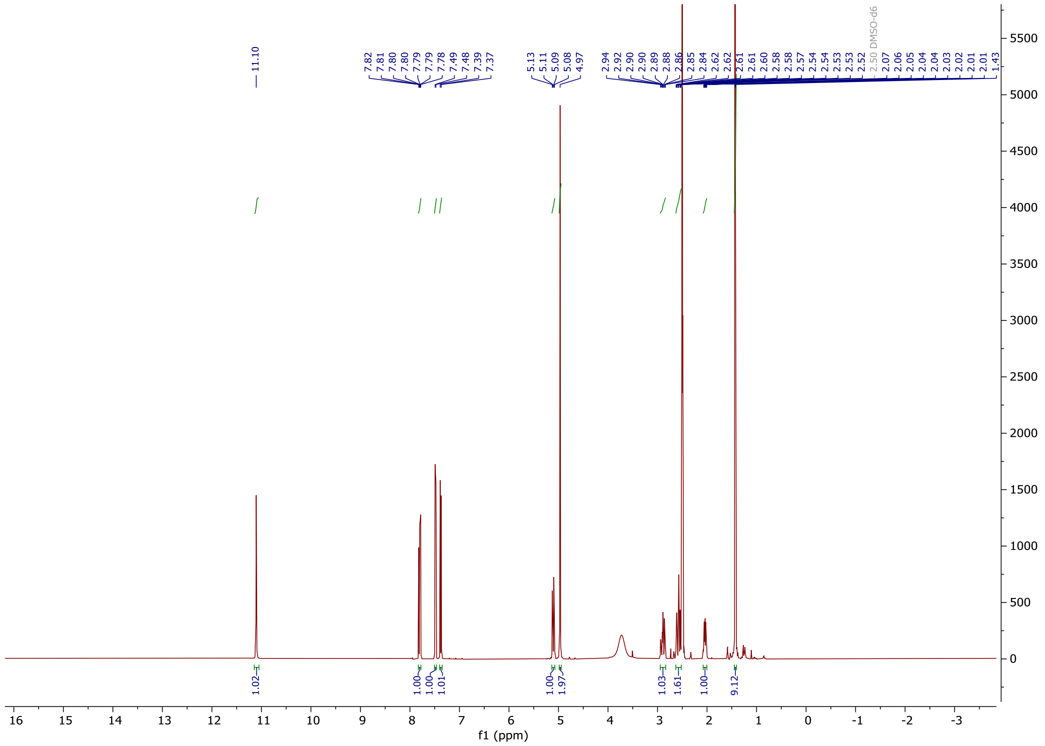


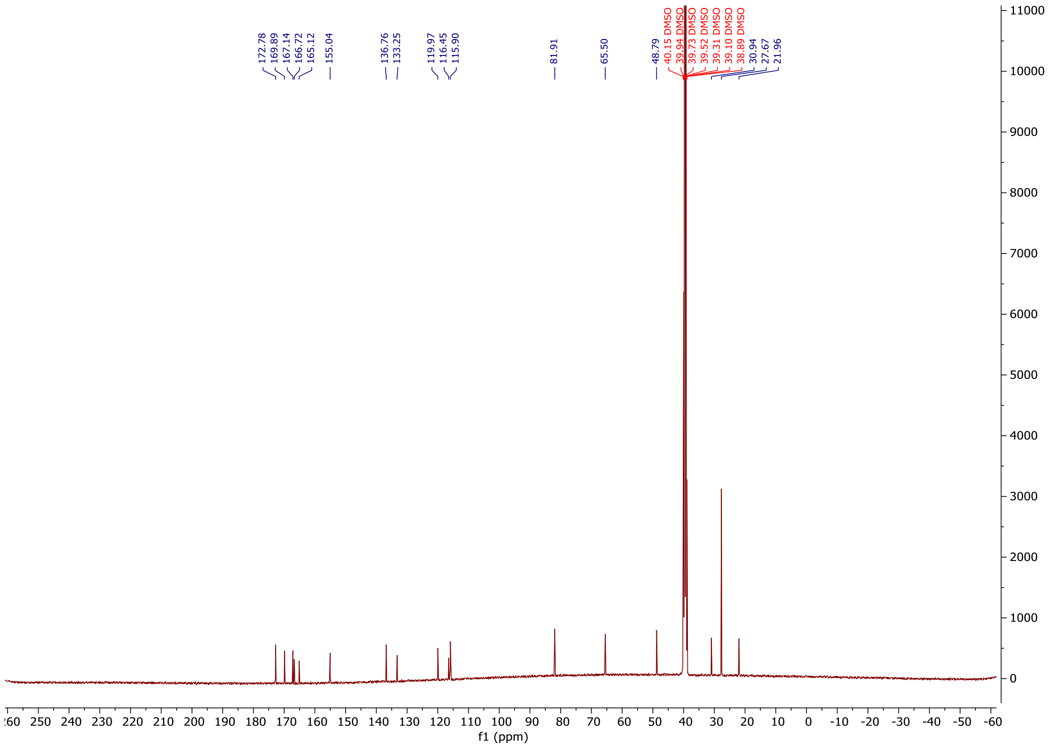

*tert*-butyl-(2-(2-(2-((2-(2,6-dioxopiperidin-3-yl)-1,3-dioxoisoindolin-4-yl)oxy)acetamido)ethoxy)ethyl)carbamate **24a**

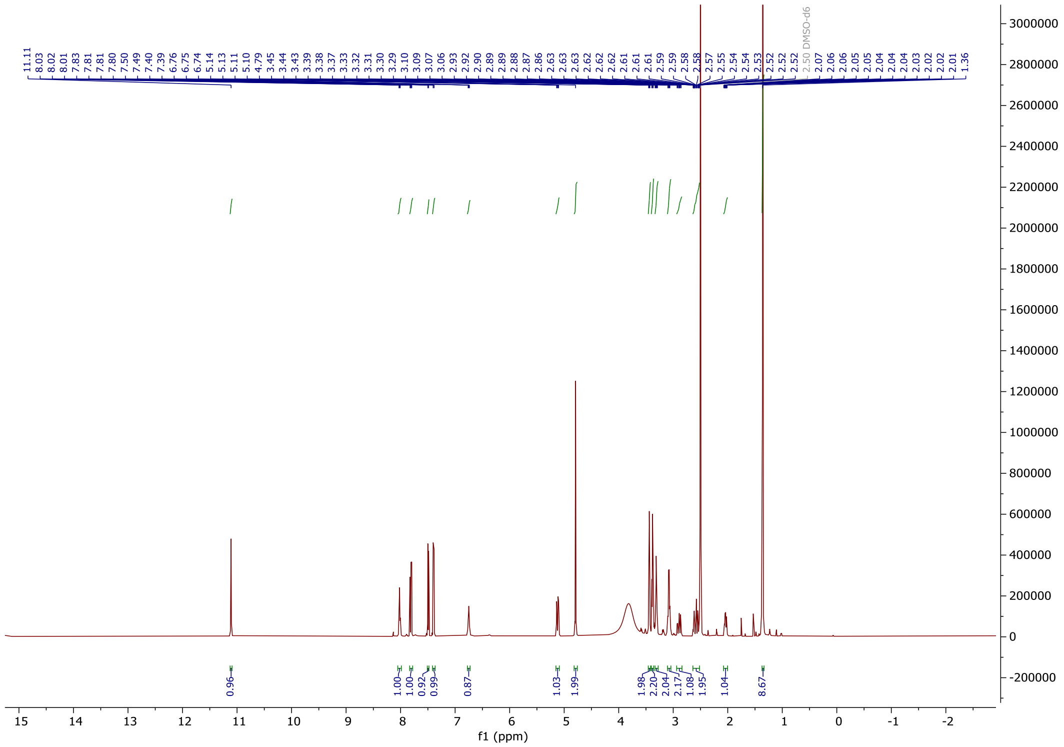


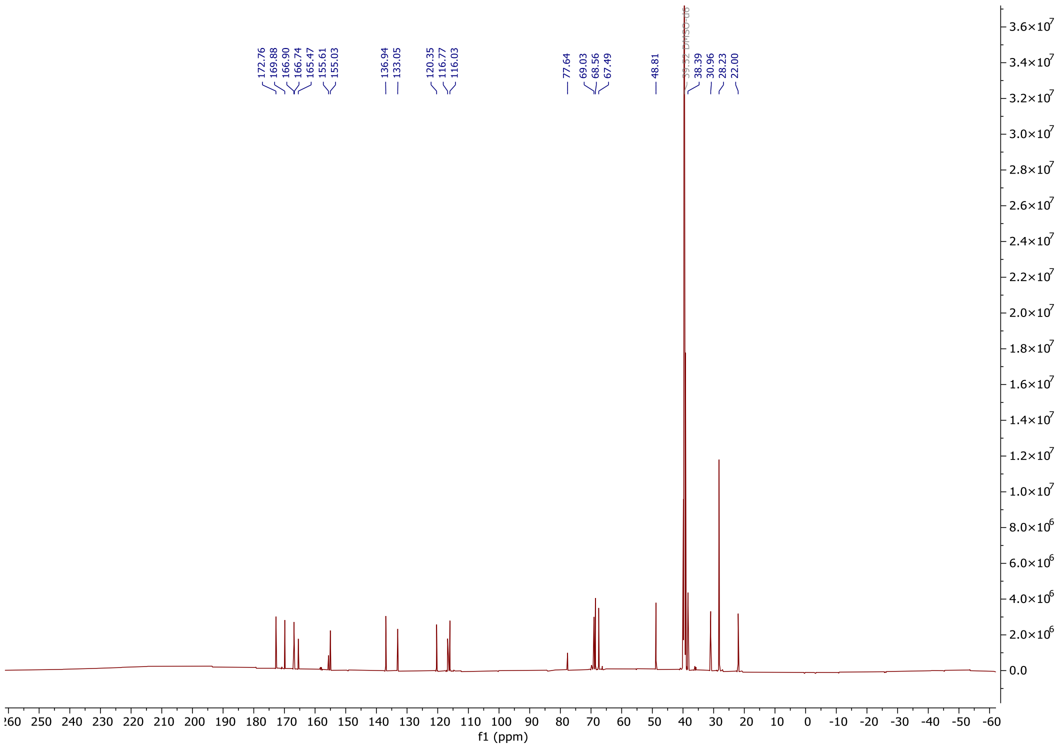

*tert*-butyl-(2-(2-(2-(2-((2-(2,6-dioxopiperidin-3-yl)-1,3-dioxoisoindolin-4-yl)oxy)acetamido)ethoxy)ethoxy)ethyl)carbamate **24b**

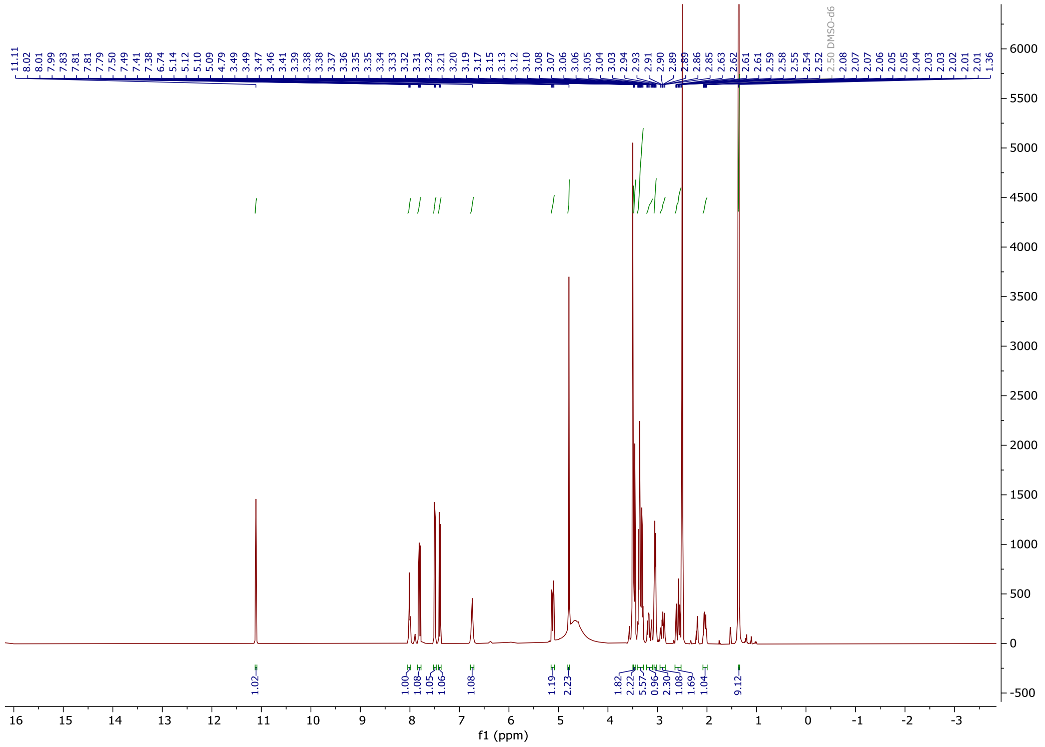

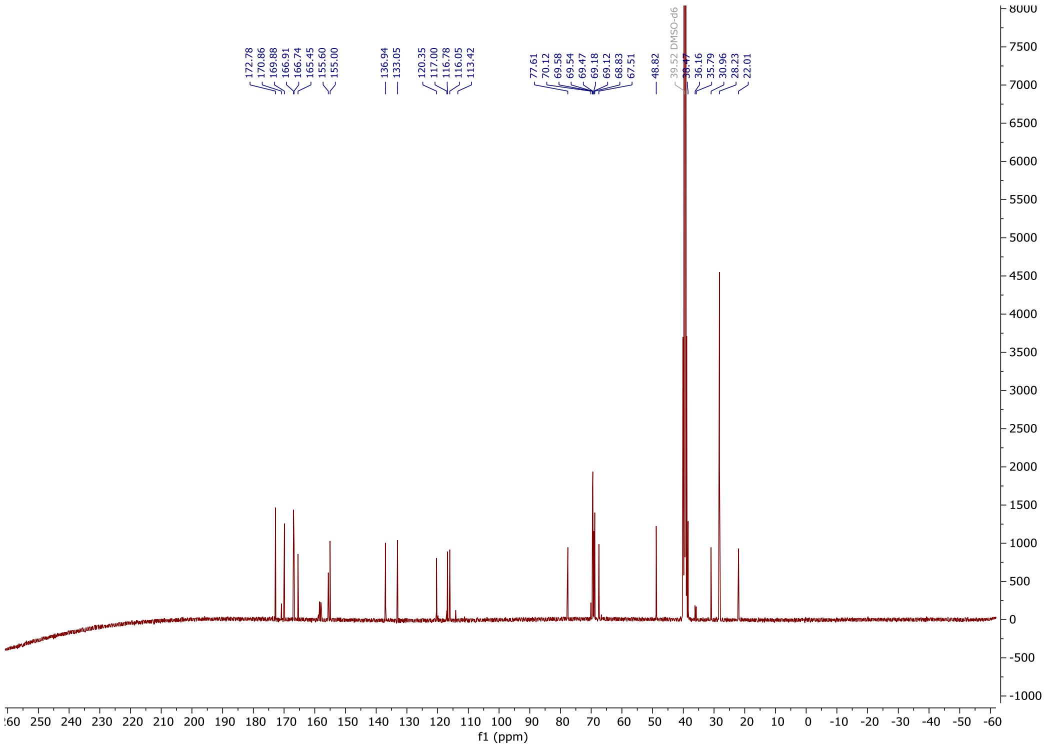

*tert*-butyl (1-((2-(2,6-dioxopiperidin-3-yl)-1,3-dioxoisoindolin-4-yl) oxy)-2-oxo-6,9,12-trioxa-3-azatetradecan-14-yl) carbamate **24c**

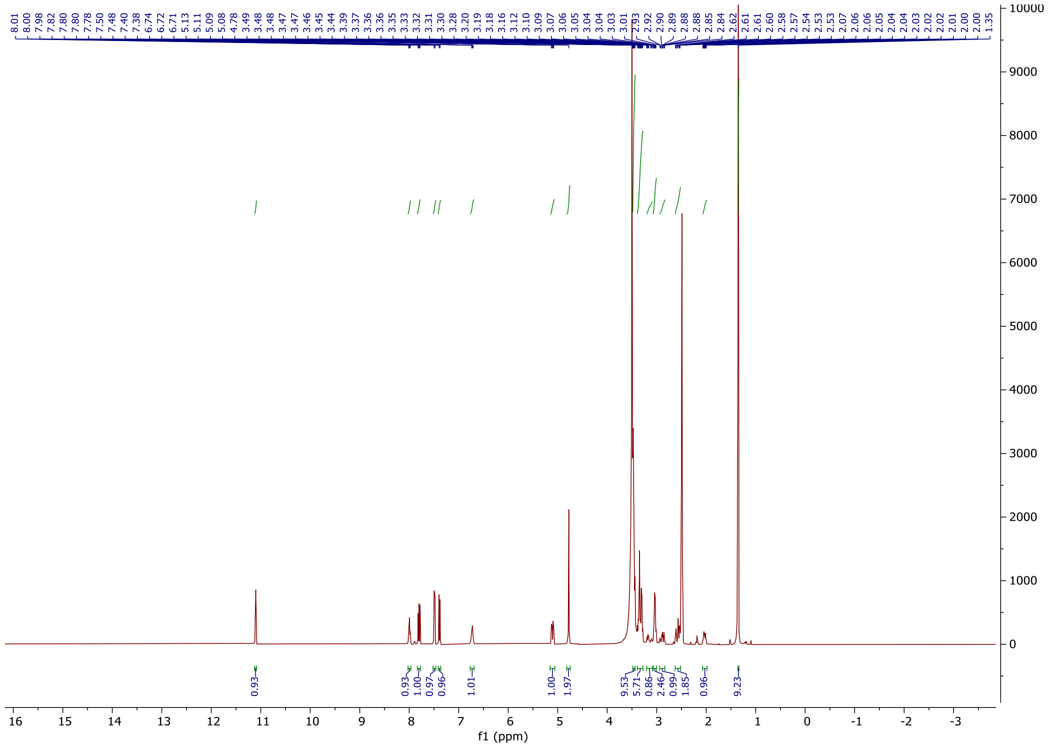

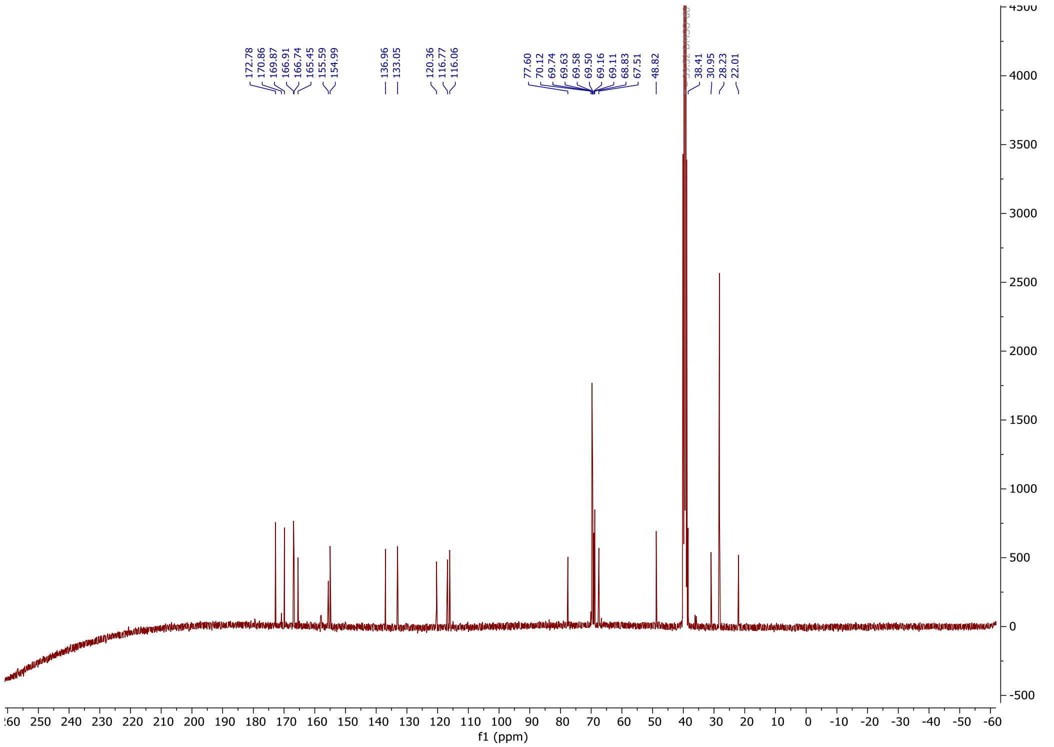

2-((2-(2,6-dioxopiperidin-3-yl)-1,3-dioxoisoindolin-4-yl)oxy)-N-(2-(2-(2-(1-(2-((2-fluoro-4-((2-fluoro-3-nitrobenzyl)sulfonyl)phenyl)thio)-5-methoxy-6-((5-methyl-1H-pyrazol-3-yl)amino)pyrimidin-4-yl)piperidin-4-yl)acetamido)ethoxy)ethyl)acetamide **D1**

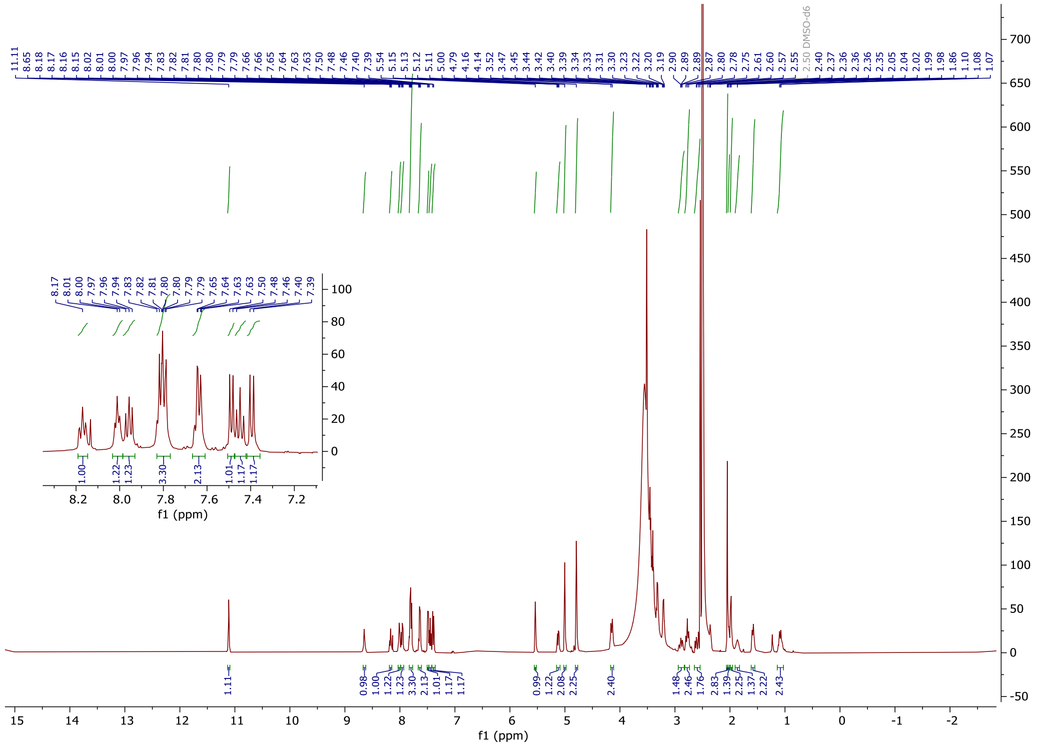


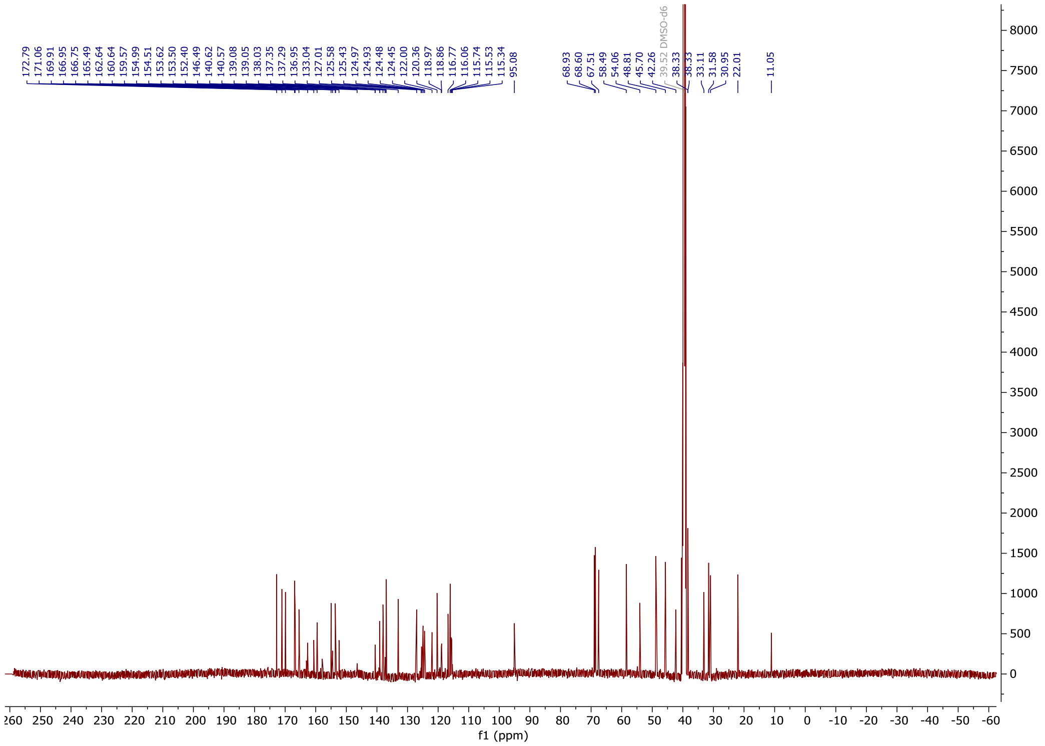

2-((2-(2,6-dioxopiperidin-3-yl)-1,3-dioxoisoindolin-4-yl)oxy)-N-(2-(2-(2-(2-(1-(2-((2-fluoro-4-((2-fluoro-3-nitrobenzyl)sulfonyl)phenyl)thio)-5-methoxy-6-((5-methyl-1H-pyrazol-3-yl)amino)pyrimidin-4-yl)piperidin-4-yl)acetamido)ethoxy)ethoxy)ethyl)acetamide **D2**

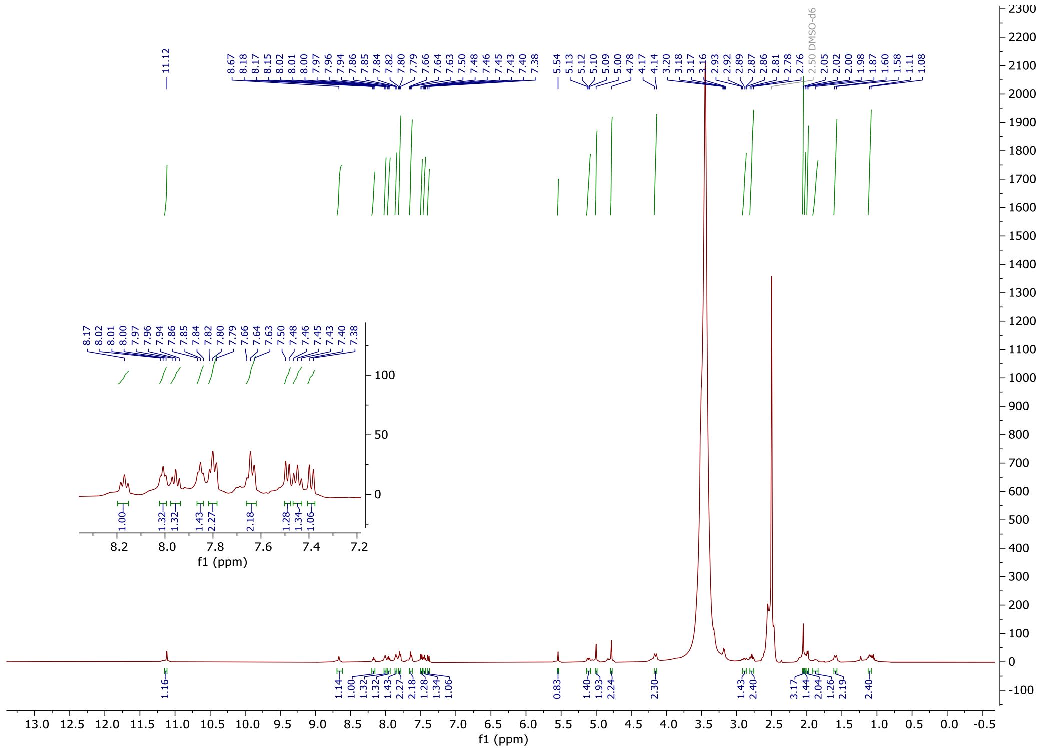


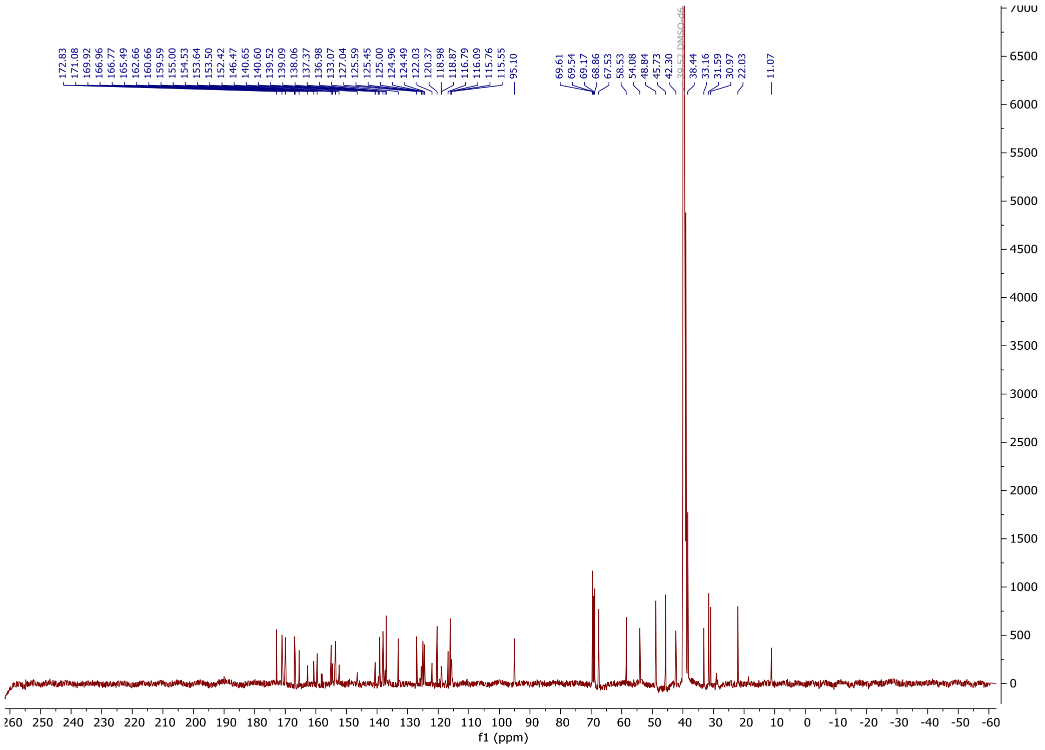

2-((2-(2,6-dioxopiperidin-3-yl)-1,3-dioxoisoindolin-4-yl)oxy)-N-(1-(1-(2-((2-fluoro-4-((2-fluoro-3-nitrobenzyl)sulfonyl)phenyl)thio)-5-methoxy-6-((5-methyl-1H-pyrazol-3-yl)amino)pyrimidin-4-yl)piperidin-4-yl)-2-oxo-6,9,12-trioxa-3-azatetradecan-14-yl)acetamide **D3**

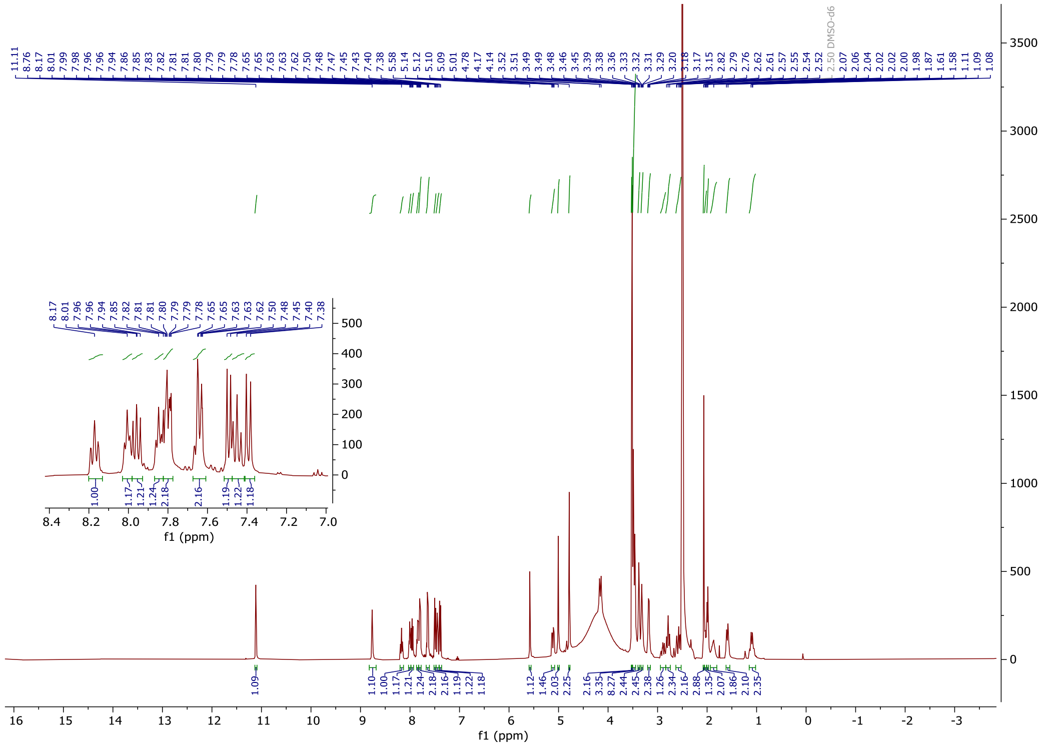

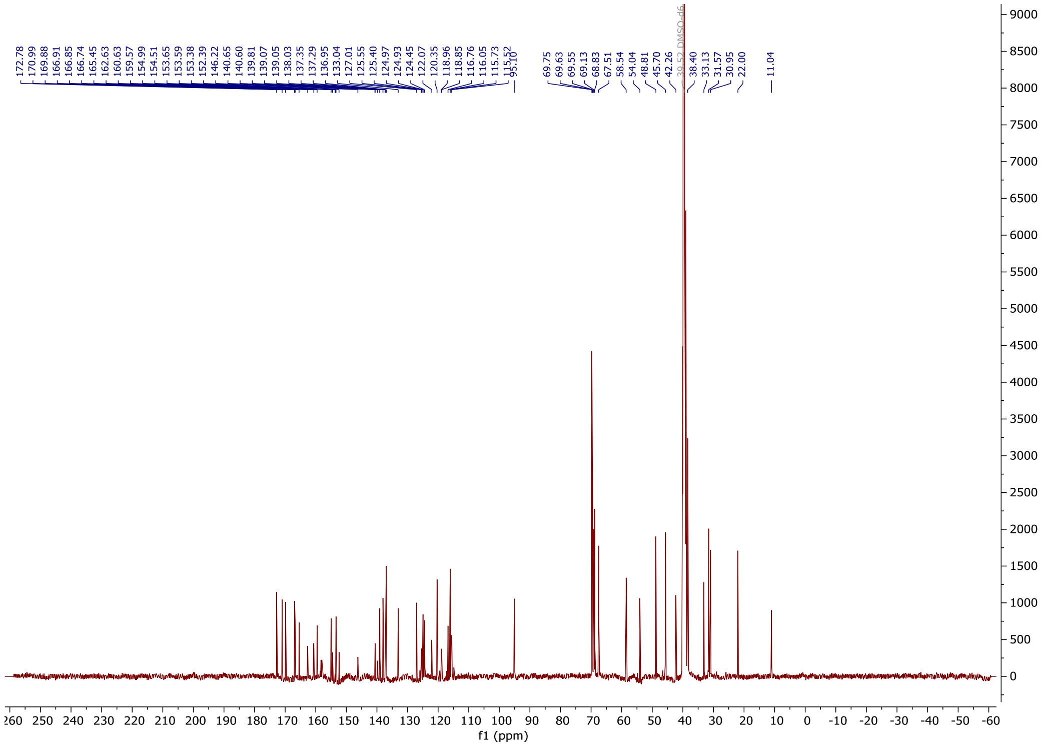


*tert*-butyl (2-(2-((2-(2,6-dioxopiperidin-3-yl)-1,3-dioxoisoindolin-4-yl)oxy)acetamido)ethyl)carbamate **26a**

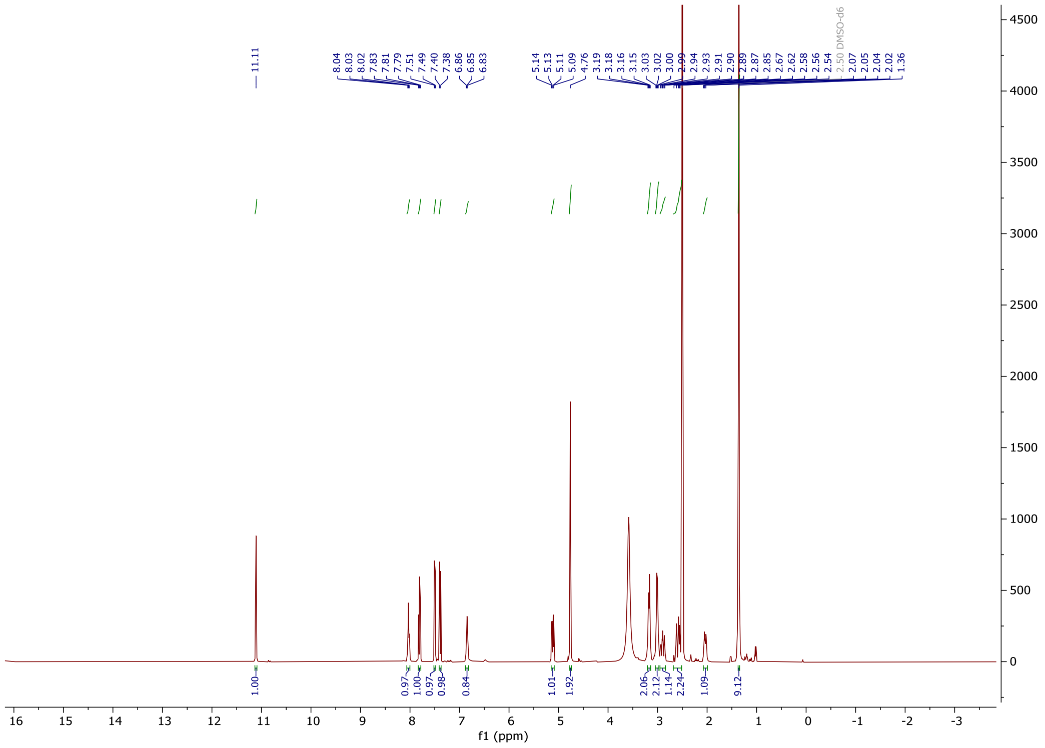

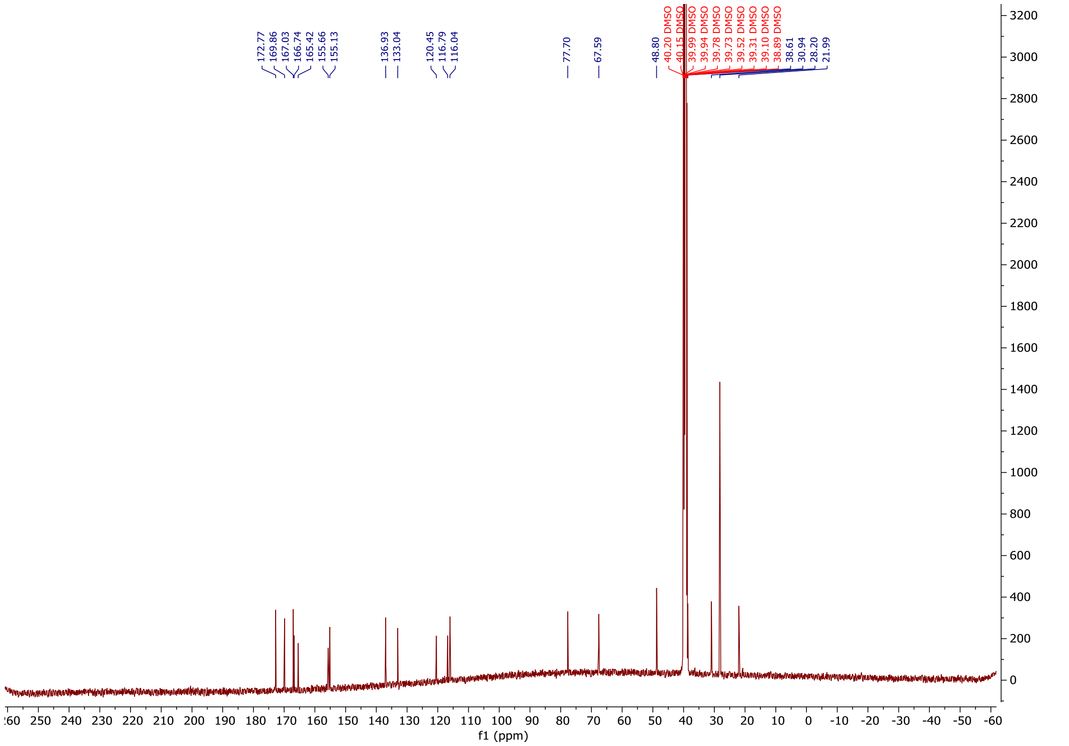

*tert*-butyl-(6-(2-((2-(2,6-dioxopiperidin-3-yl)-1,3-dioxoisoindolin-4-yl)oxy)acetamido)hexyl)carbamate **26b**

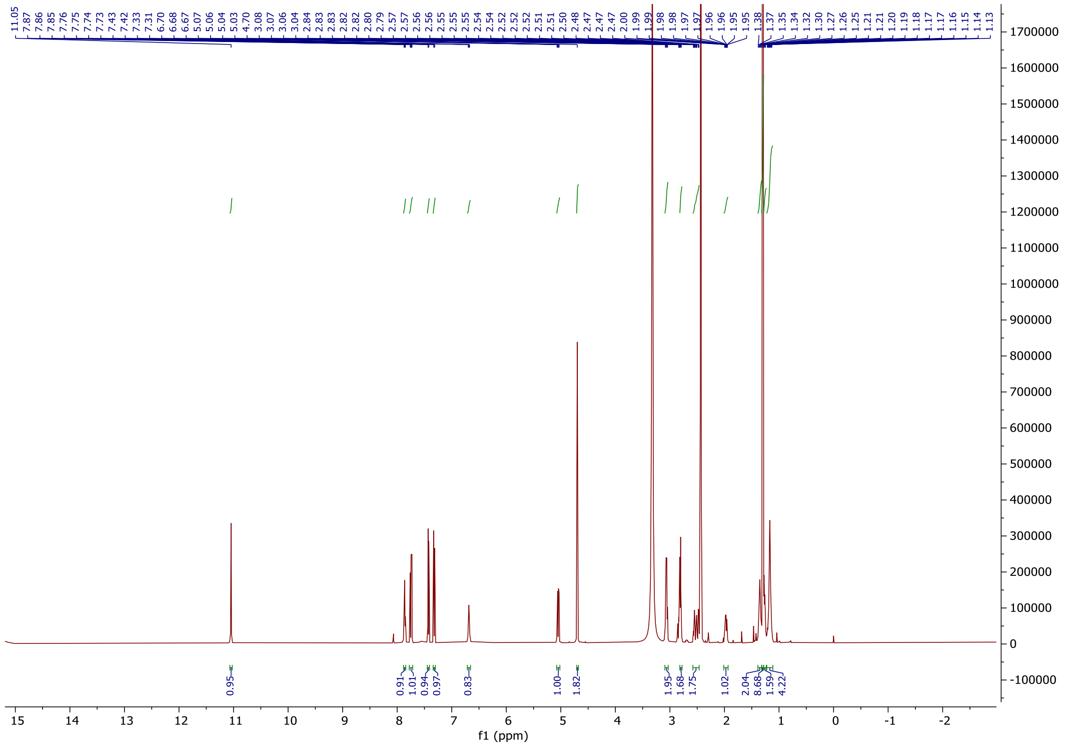

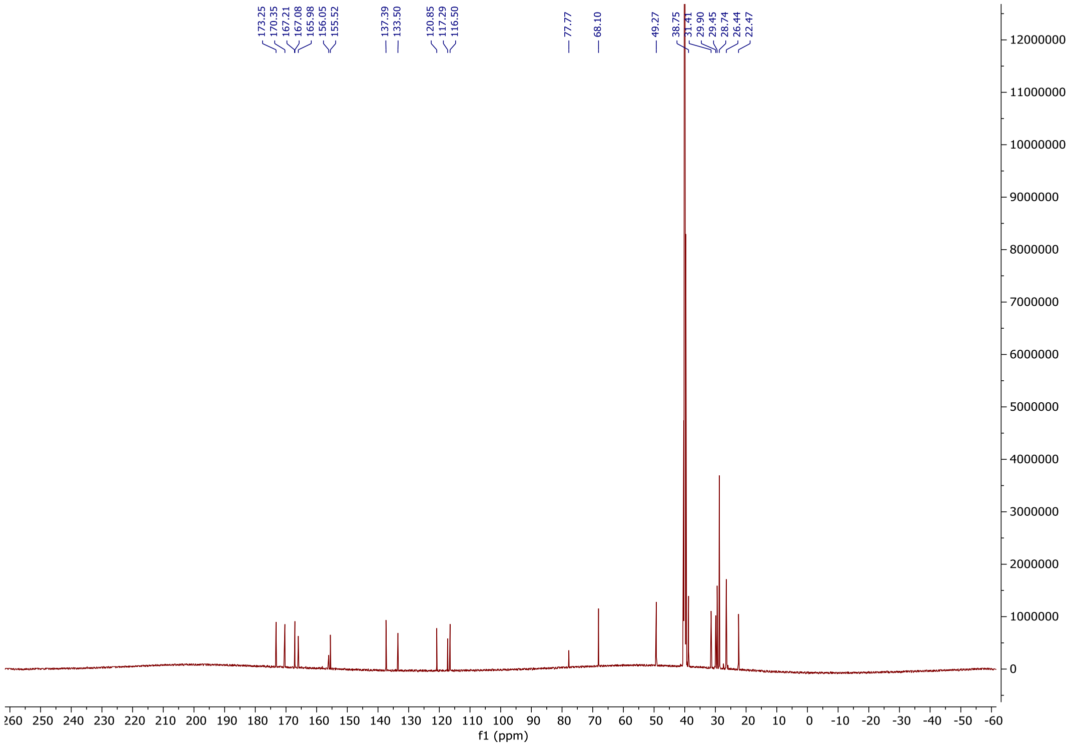


2-((2-(2,6-dioxopiperidin-3-yl)-1,3-dioxoisoindolin-4-yl)oxy)-N-(2-(2-(1-(2-((2-fluoro-4-((2-fluoro-3-nitrobenzyl)sulfonyl)phenyl)thio)-5-methoxy-6-((5-methyl-1H-pyrazol-3-yl)amino)pyrimidin-4-yl)piperidin-4-yl)acetamido)ethyl)acetamide **D4**

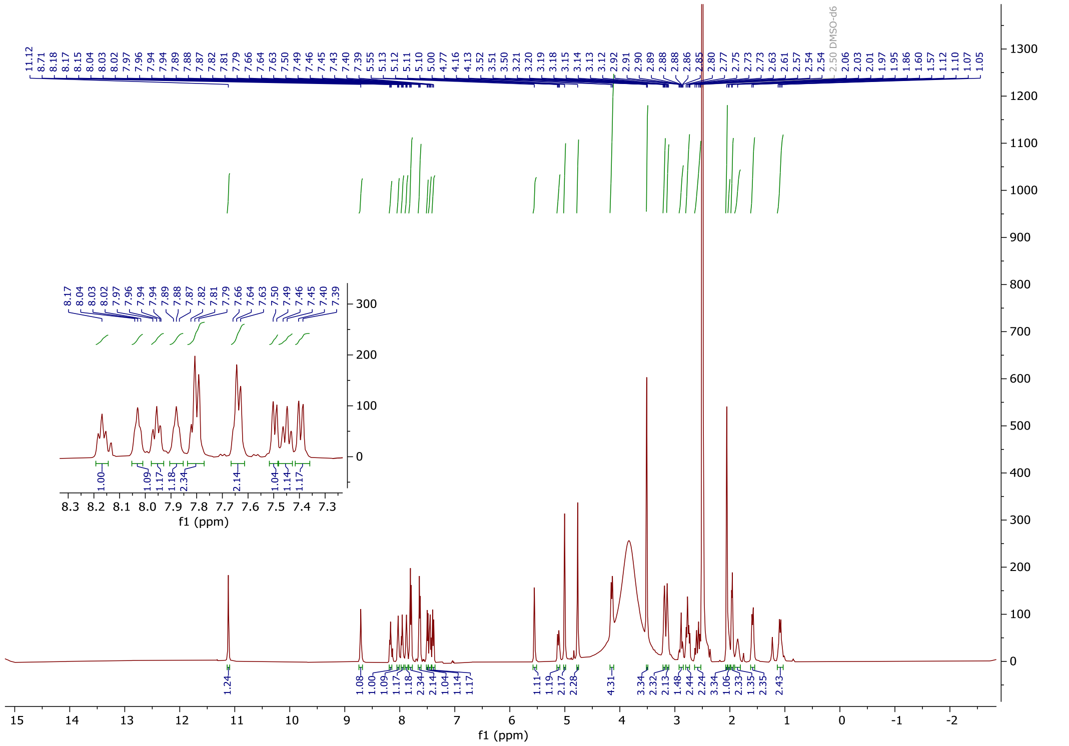

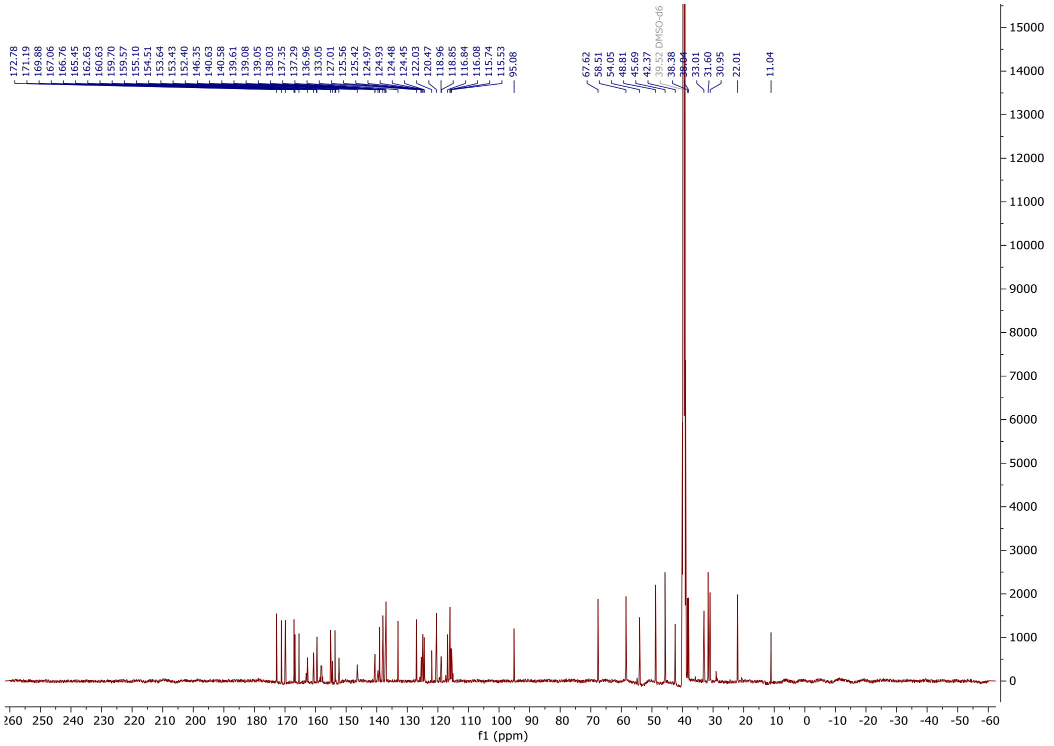

2-((2-(2,6-dioxopiperidin-3-yl)-1,3-dioxoisoindolin-4-yl)oxy)-N-(6-(2-(1-(2-((2-fluoro-4-((2-fluoro-3-nitrobenzyl)sulfonyl)phenyl)thio)-5-methoxy-6-((5-methyl-1H-pyrazol-3-yl)amino)pyrimidin-4-yl)piperidin-4-yl)acetamido)hexyl)acetamide **D5**

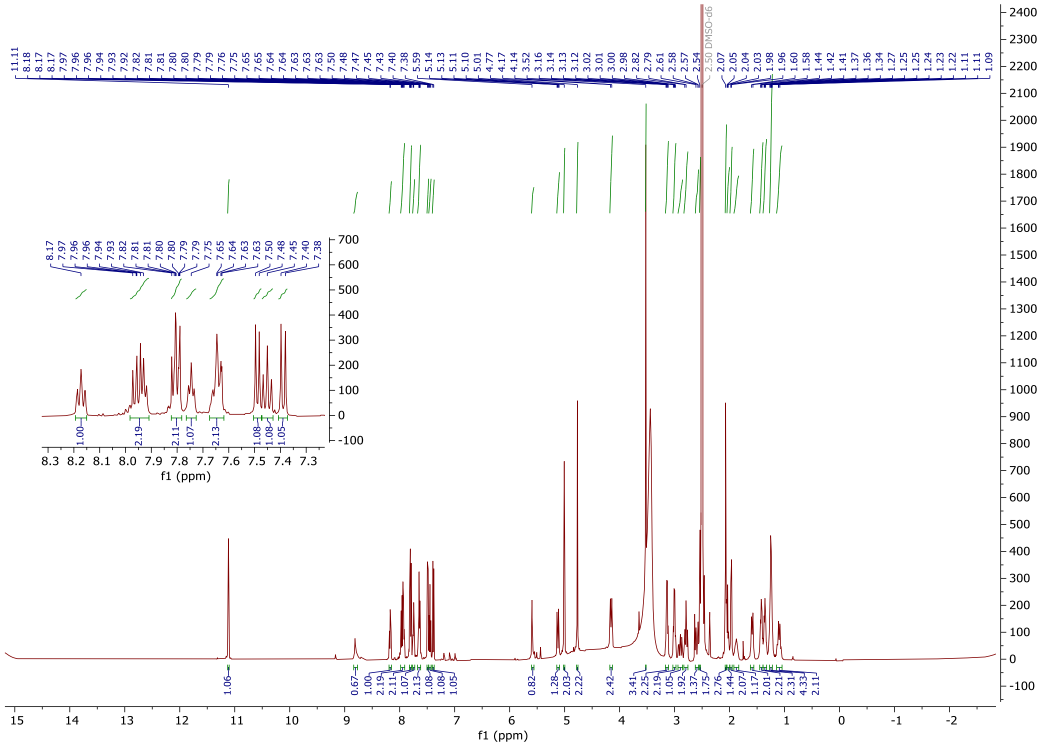

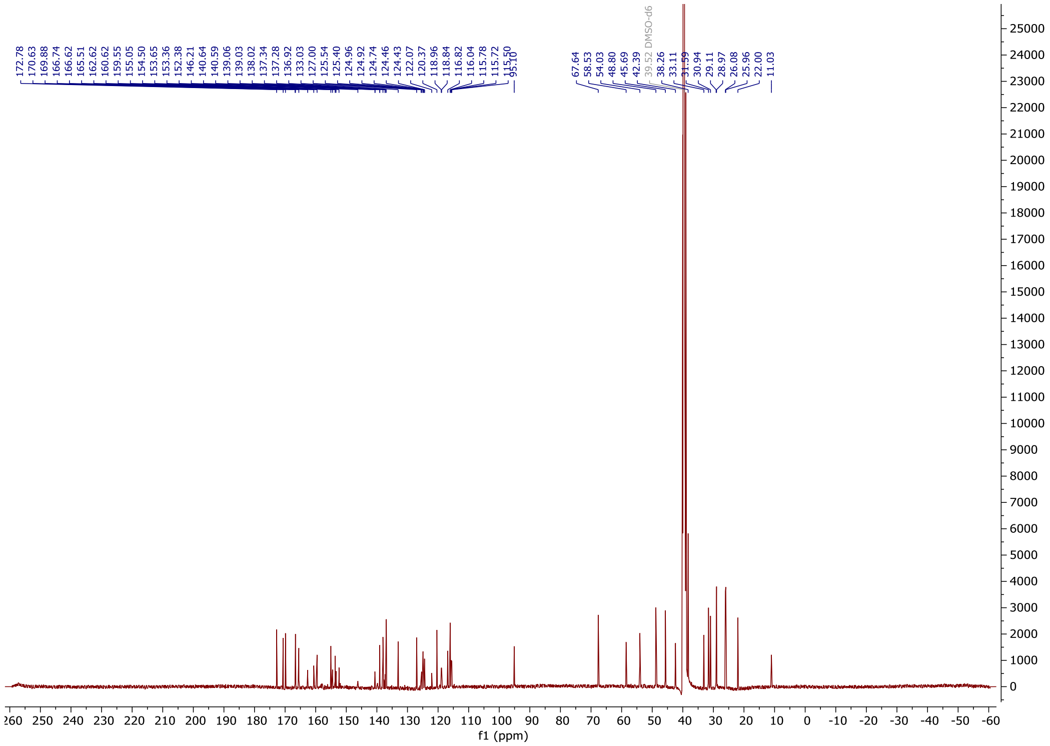

*tert*-butyl-(4-((2-(2,6-dioxopiperidin-3-yl)-1,3-dioxoisoindolin-4-yl)oxy)butyl)carbamate **28b**

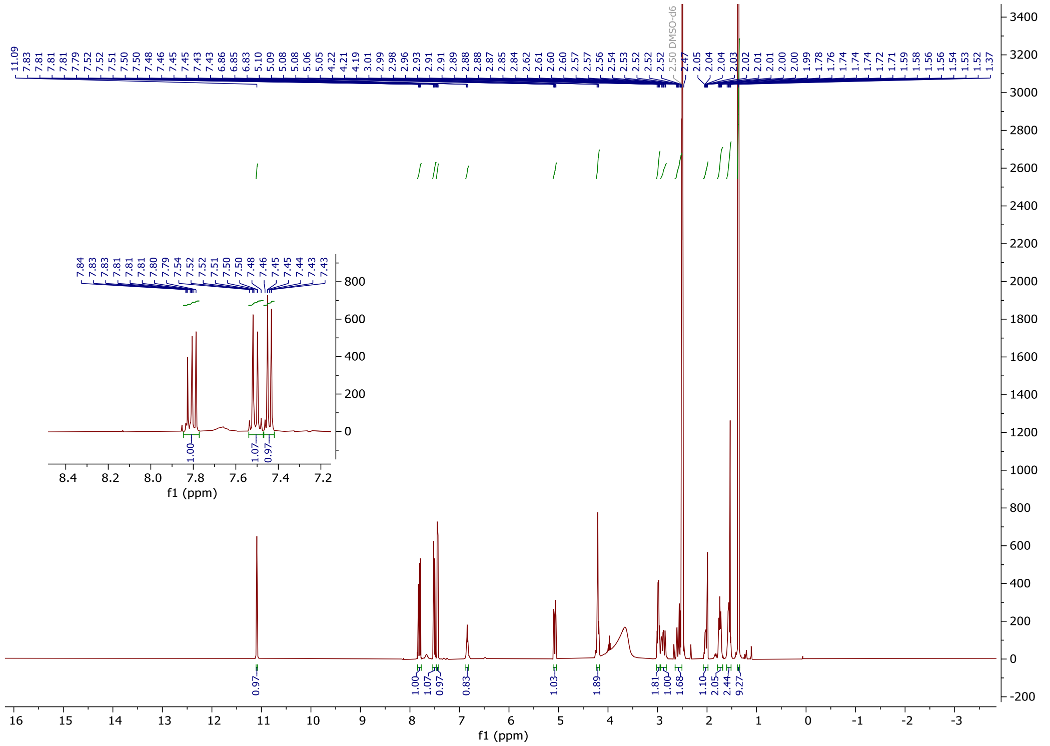

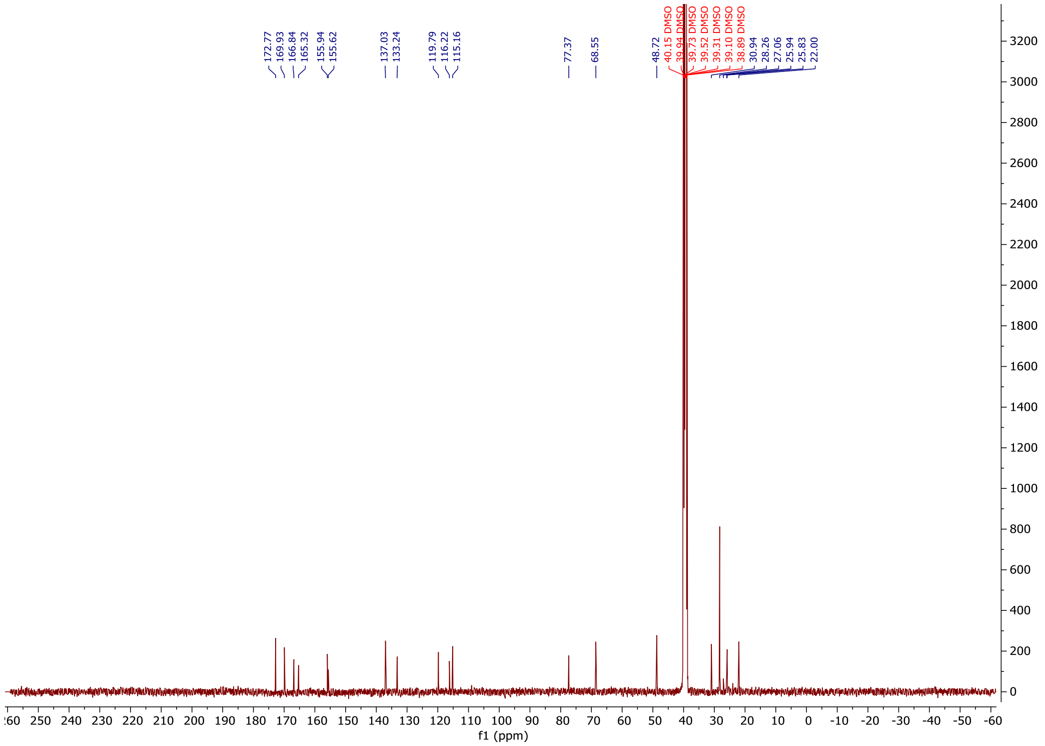

*N*-(2-((2-(2,6-dioxopiperidin-3-yl)-1,3-dioxoisoindolin-4-yl) oxy) ethyl)-2-(1-(2-((2-fluoro-4-((2-fluoro-3-nitrobenzyl) sulfonyl) phenyl) thio)-5-methoxy-6-((5-methyl-1H-pyrazol-3-yl) amino) pyrimidin-4-yl)piperidin-4-yl) acetamide **D6**

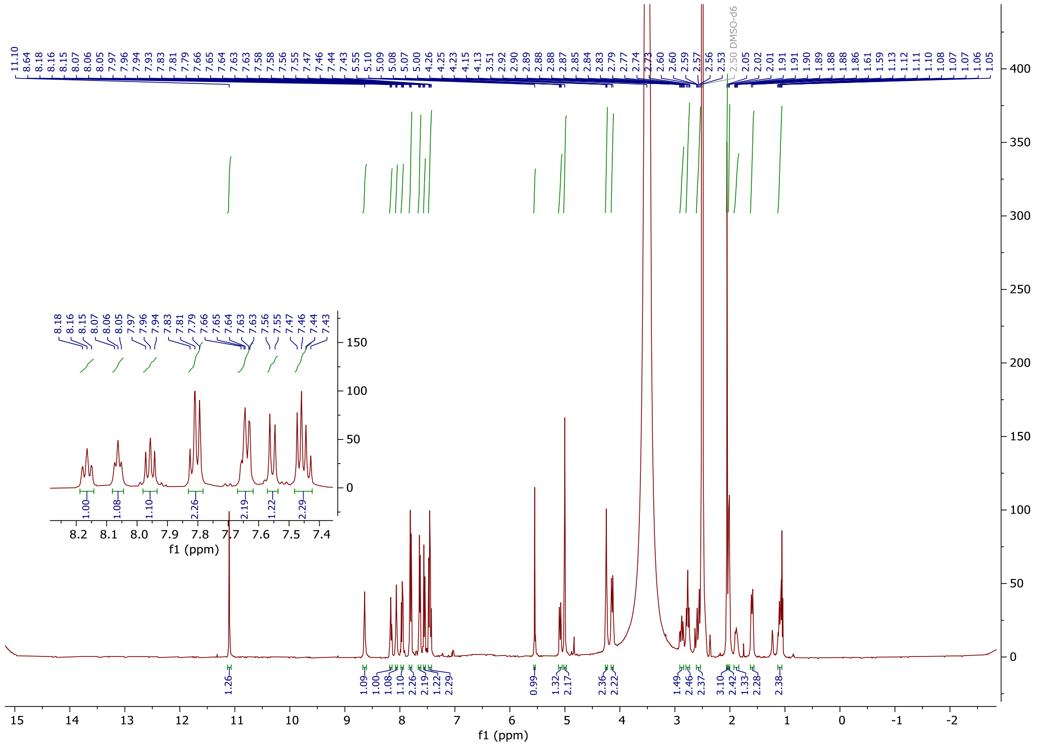

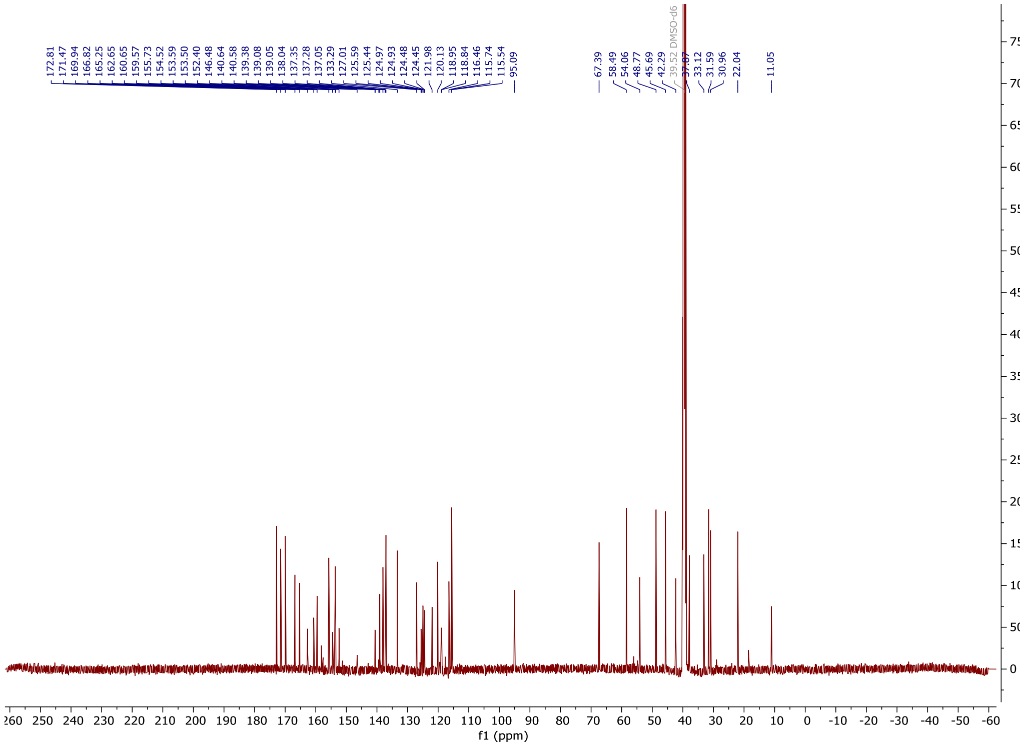


*N*-(4-((2-(2,6-dioxopiperidin-3-yl)-1,3-dioxoisoindolin-4-yl) oxy) butyl)-2-(1-(2-((2-fluoro-4-((2-fluoro-3-nitrobenzyl) sulfonyl) phenyl) thio)-5-methoxy-6-((5-methyl-1H-pyrazol-3-yl) amino) pyrimidin-4-yl)piperidin-4-yl) acetamide **D7**

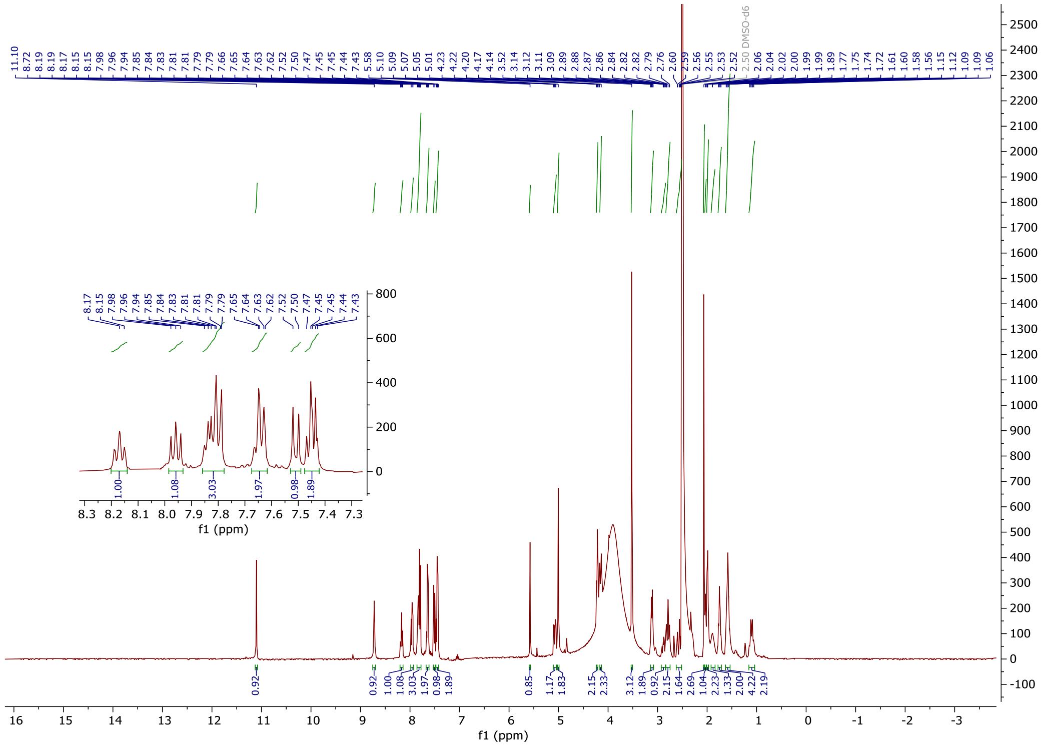

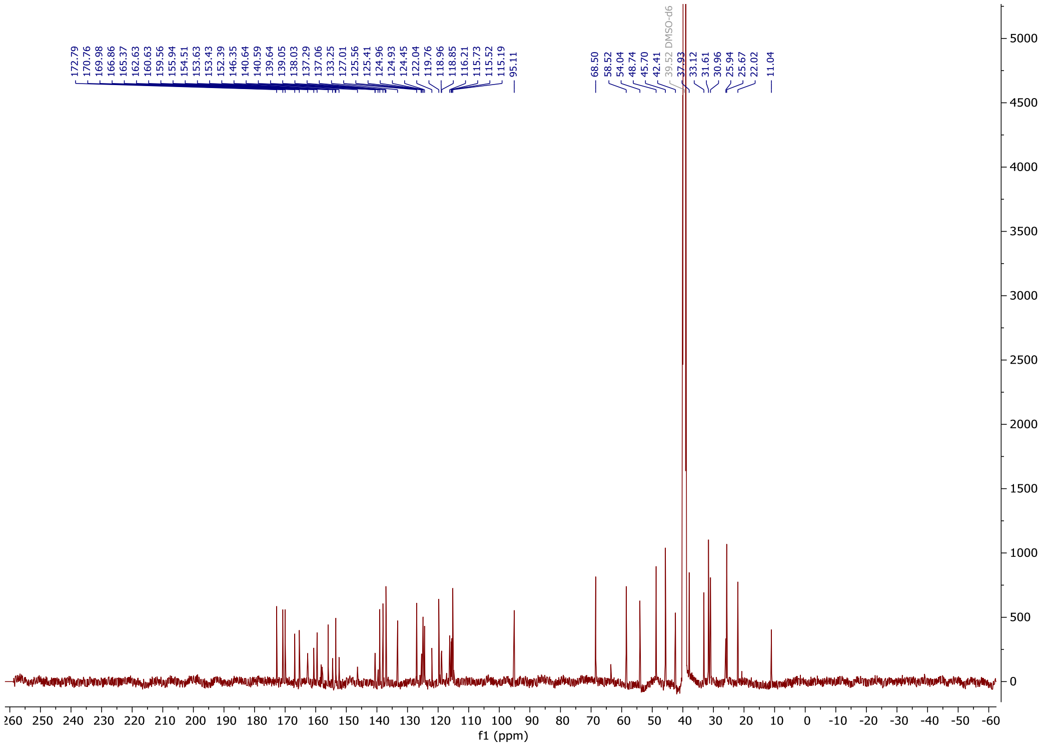


2-(2,6-dioxopiperidin-3-yl)-4-(4-(2-((2-fluoro-4-((2-fluoro-3-nitrobenzyl) sulfonyl) phenyl) thio)-5-methoxy-6-((5-methyl-1H-pyrazol-3-yl)amino) pyrimidin-4-yl) piperazin-1-yl) isoindoline-1,3-dione **30**

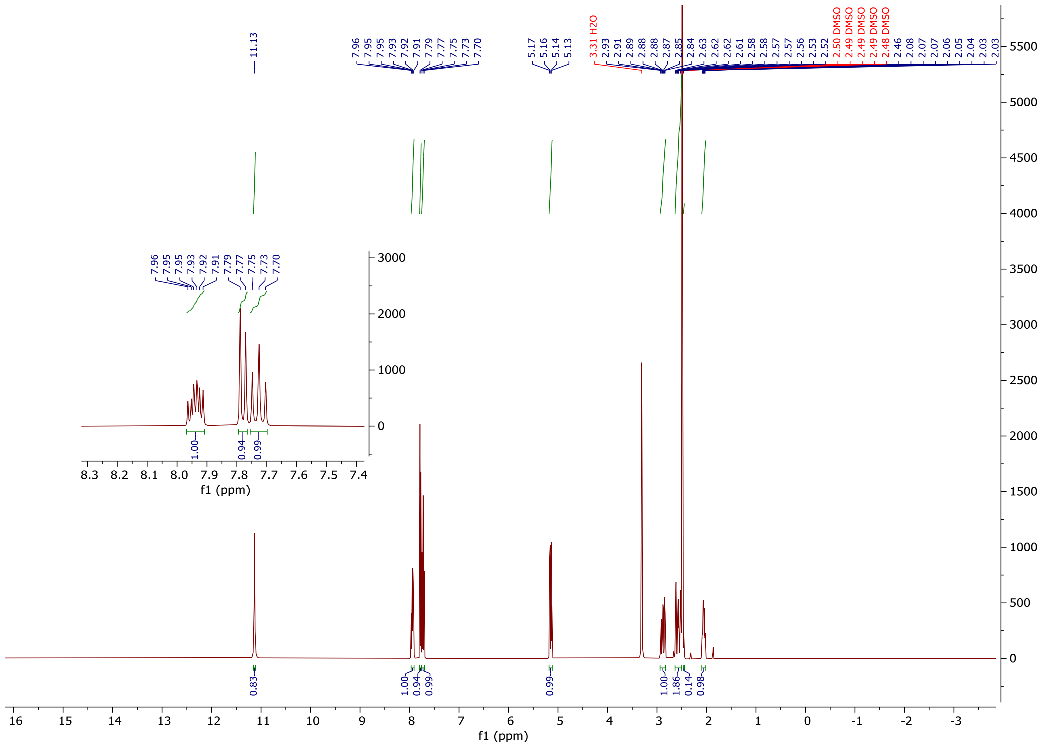

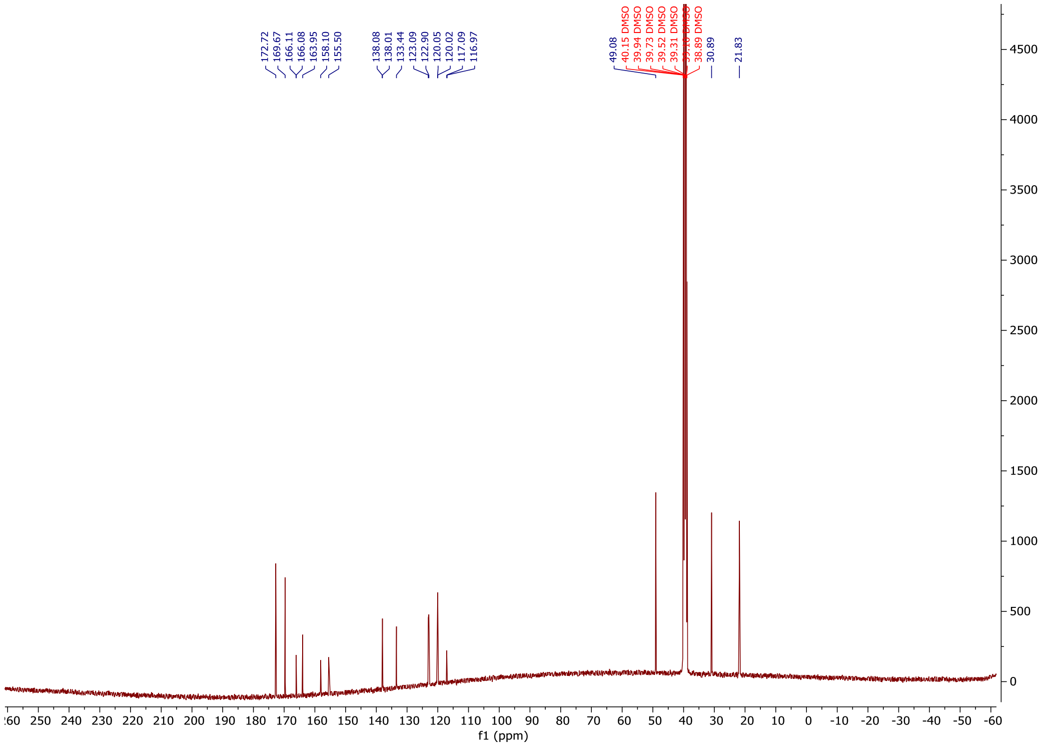


2-(2,6-dioxopiperidin-3-yl)-4-(4-(2-((2-fluoro-4-((2-fluoro-3-nitrobenzyl) sulfonyl) phenyl) thio)-5-methoxy-6-((5-methyl-1H-pyrazol-3-yl)amino) pyrimidin-4-yl) piperazin-1-yl) isoindoline-1,3-dione **D8**

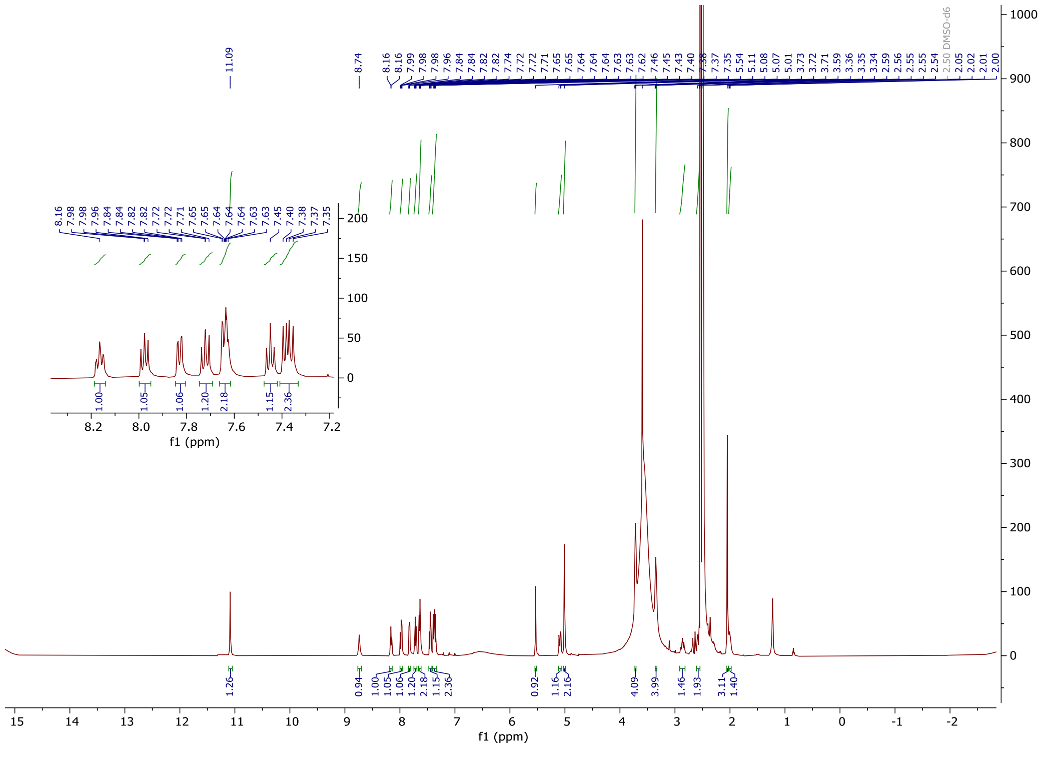

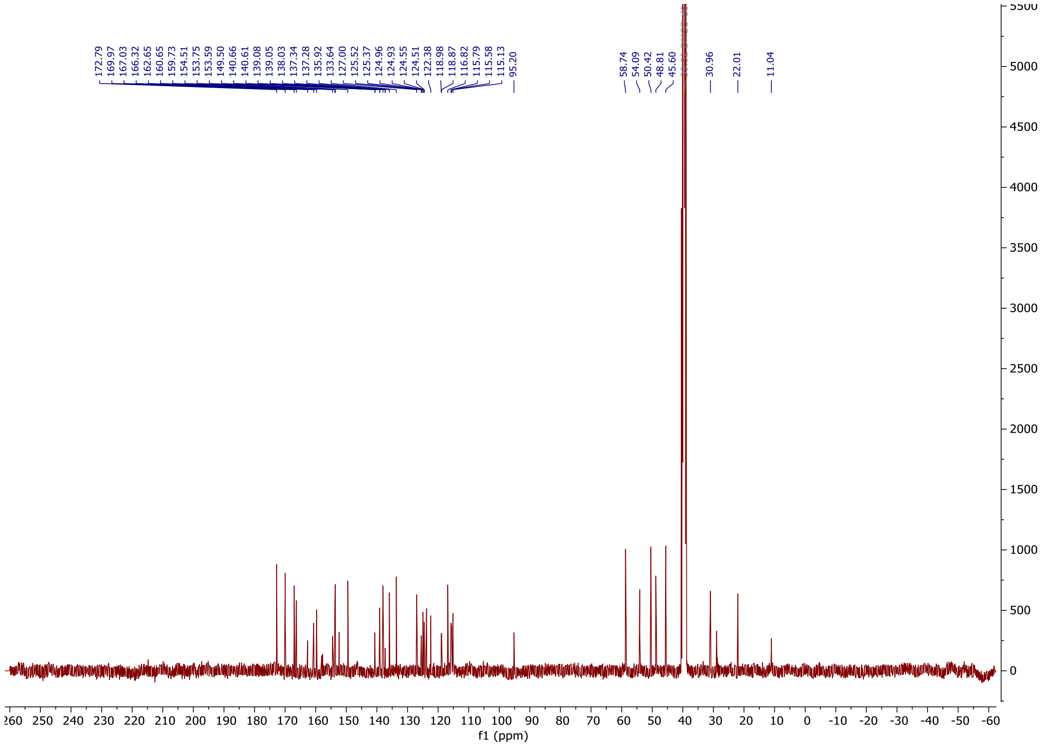

*tert*-butyl-4-(2-(2,6-dioxopiperidin-3-yl)-1,3-dioxoisoindolin-4-yl)-piperazine-1-carboxylate

**32**

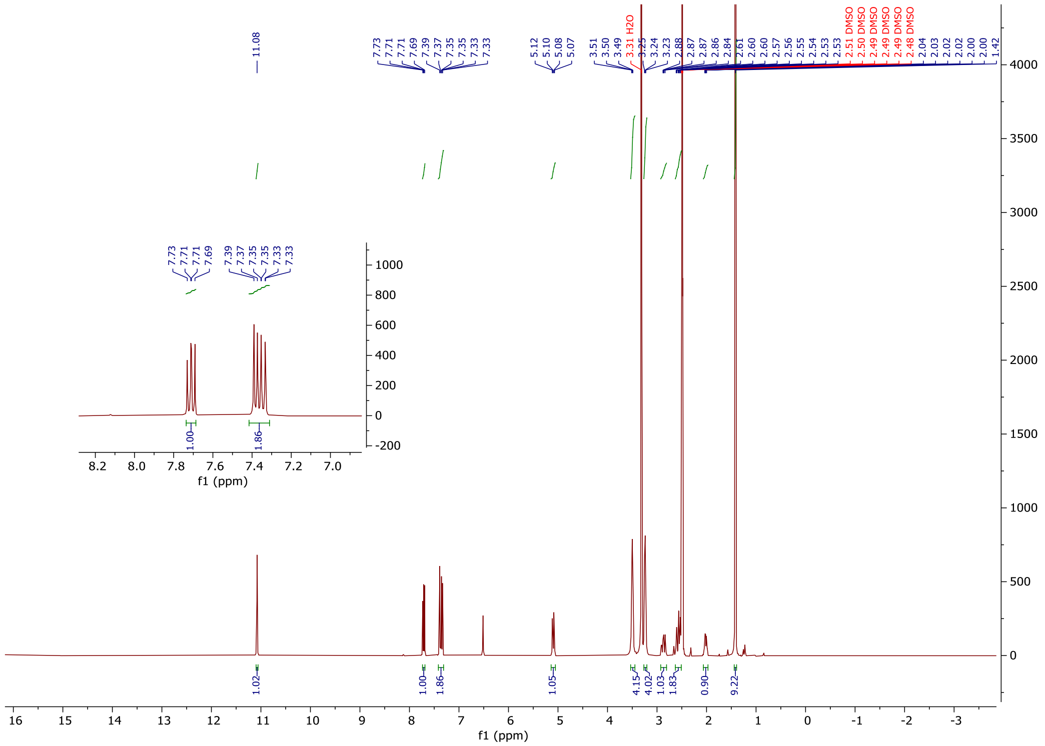

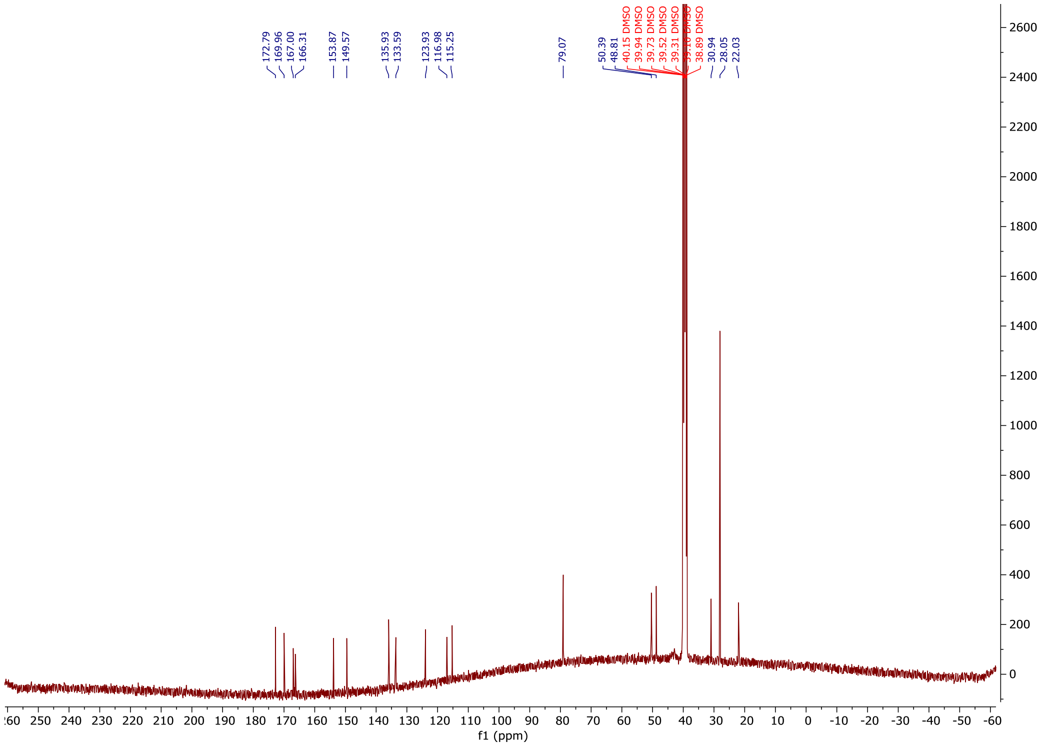

*tert*-butyl-(2-(4-(2-(2,6-dioxopiperidin-3-yl)-1,3-dioxoisoindolin-4-yl)piperazin-1-yl)-2-oxoethyl)carbamate **33**

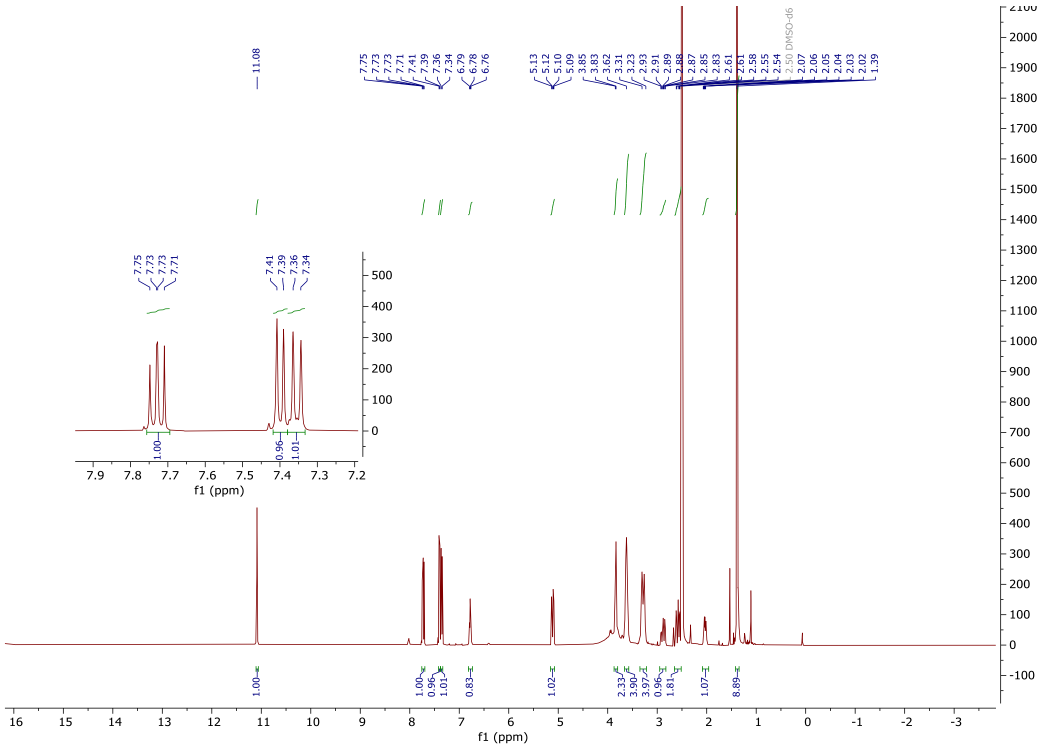

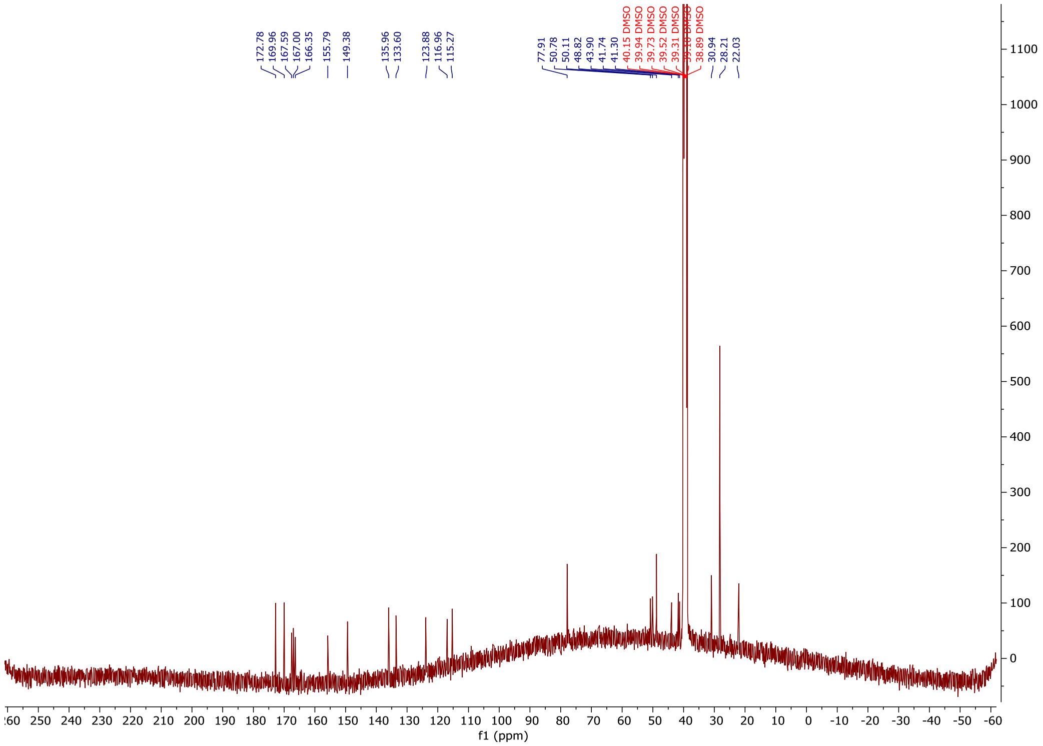


2-(2,6-dioxopiperidin-3-yl)-4-(4-(2-(1-(2-((2-fluoro-4-((2-fluoro-3-nitrobenzyl) sulfonyl) phenyl) thio)-5-methoxy-6-((5-methyl-1H-pyrazol-3-yl) amino) pyrimidin-4-yl) piperidin-4-yl) acetyl) piperazin-1-yl) isoindoline-1,3-dione **D9**

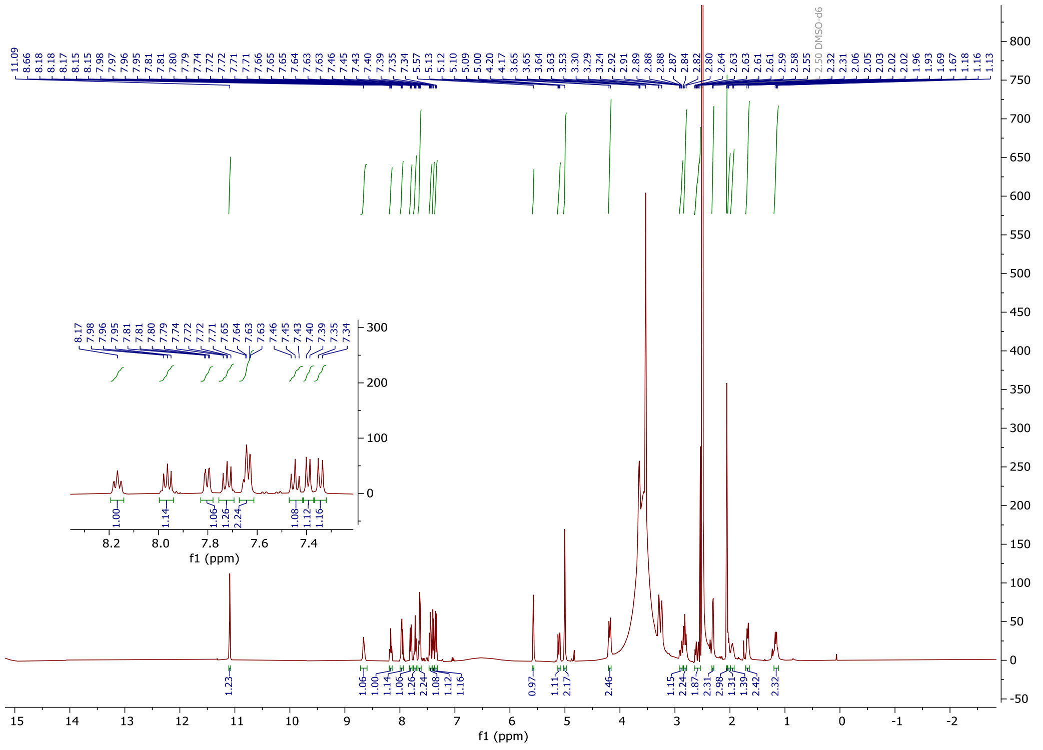

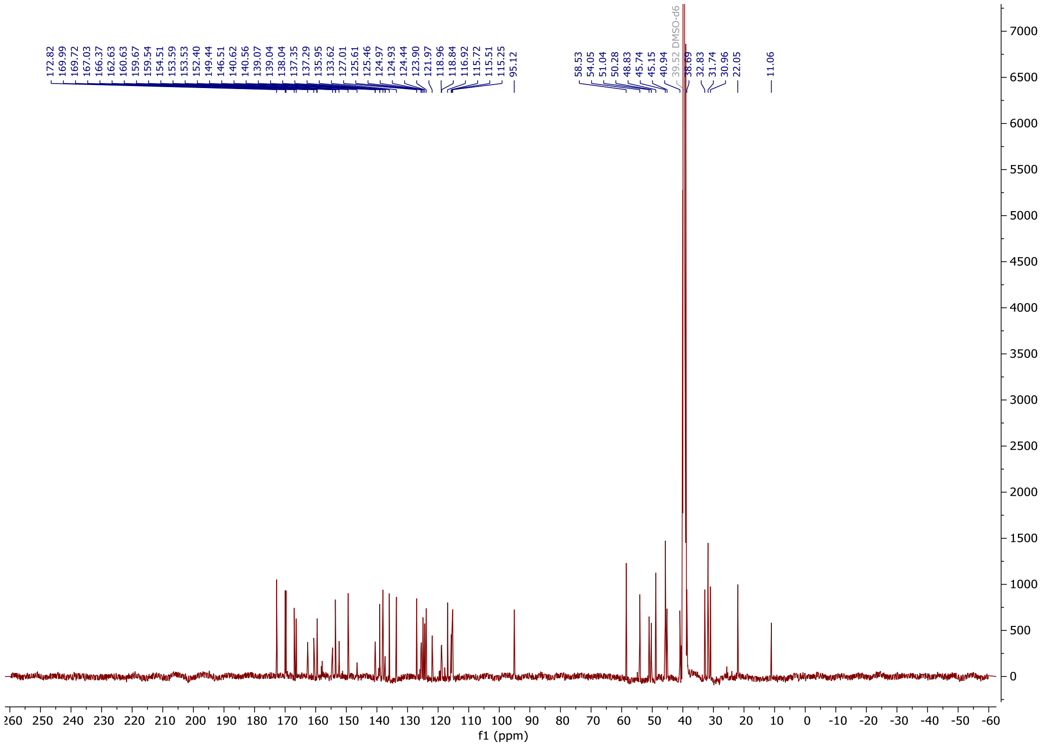

*N-(*2-(4-(2-(2,6-dioxopiperidin-3-yl)-1,3-dioxoisoindolin-4-yl)piperazin-1-yl)-2-oxoethyl)-2-(1-(2-((2-fluoro-4-((2-fluoro-3-nitrobenzyl)sulfonyl)phenyl)thio)-5-methoxy-6-((5-methyl-1H-pyrazol-3-yl)amino)pyrimidin-4-yl)piperidin-4-yl)acetamide **D10**

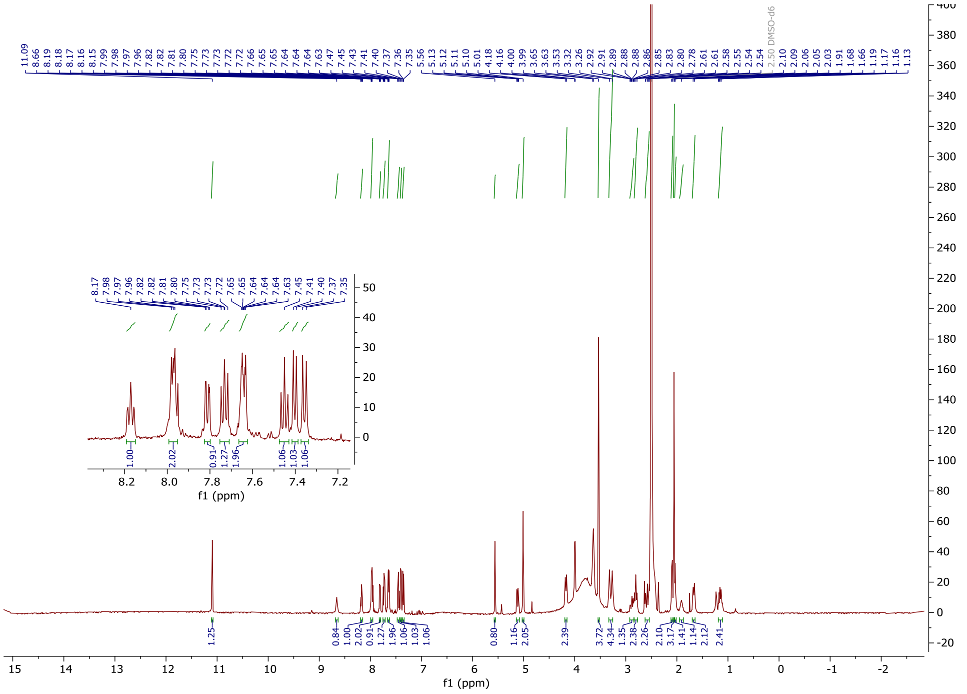

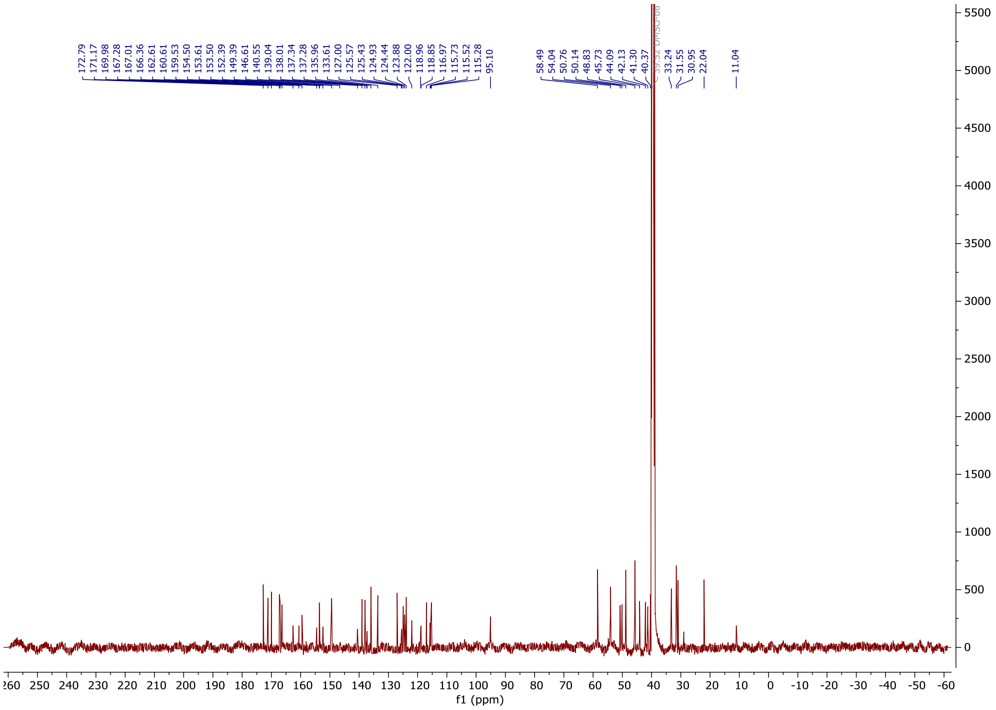


2-(2,6-dioxopiperidin-3-yl)-5-fluoroisoindoline-1,3-dione **35**

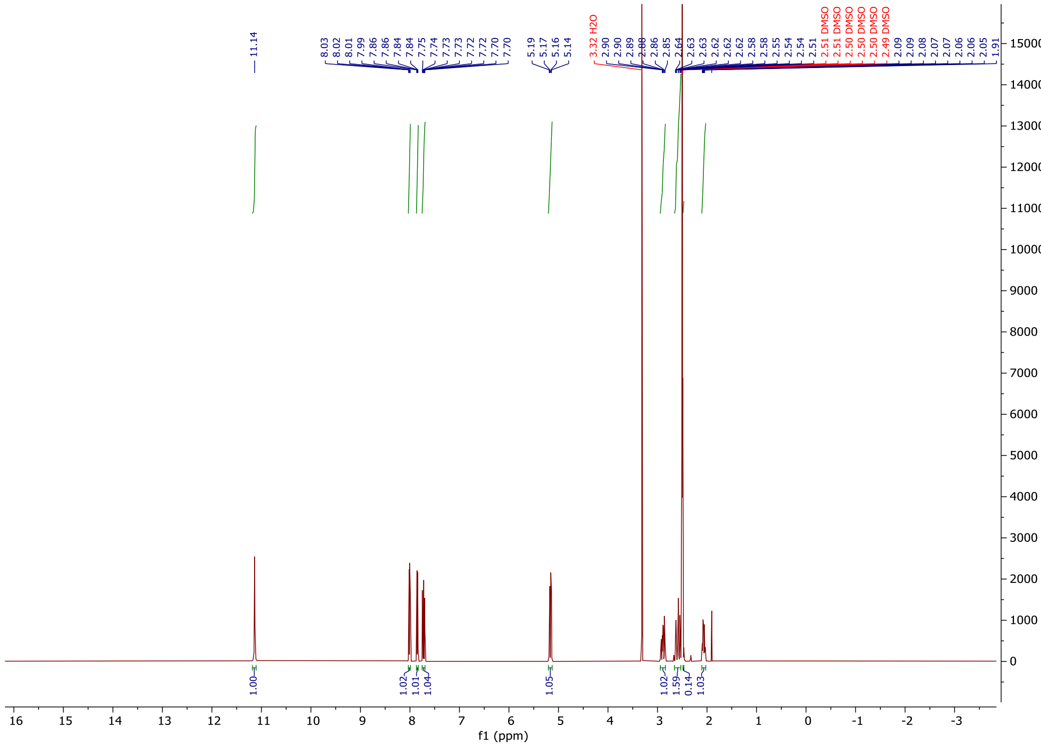

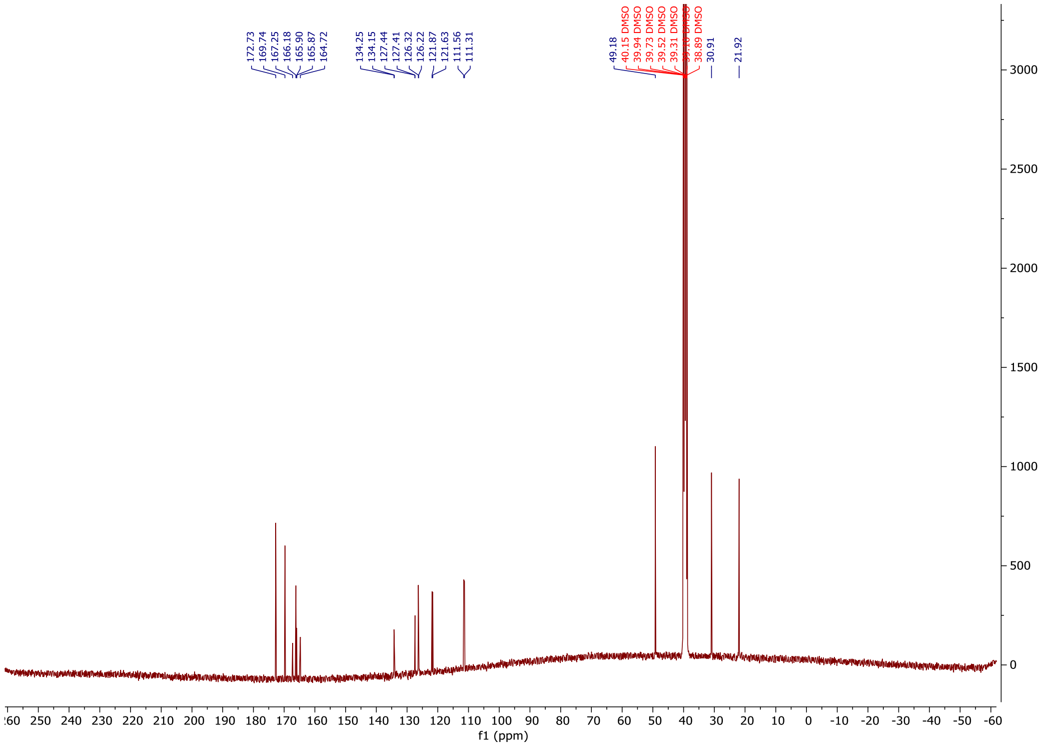

*tert*-butyl-4-(2-(2,6-dioxopiperidin-3-yl)-1,3-dioxoisoindolin-5-yl)piperazine-1-carboxylate

**36**

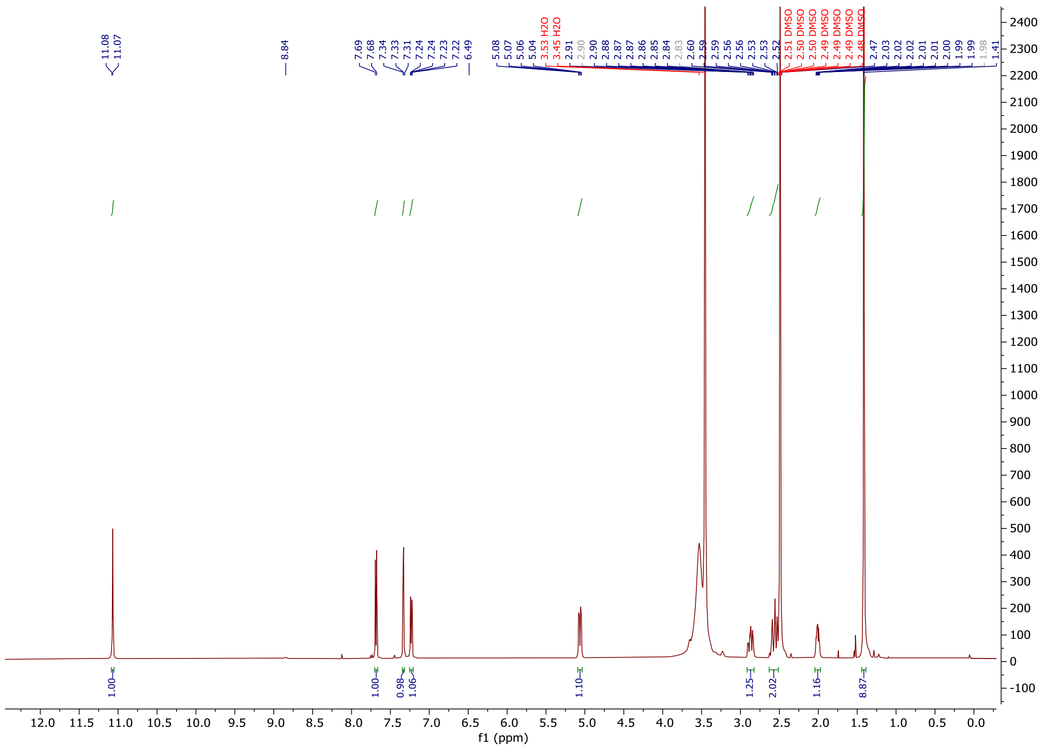

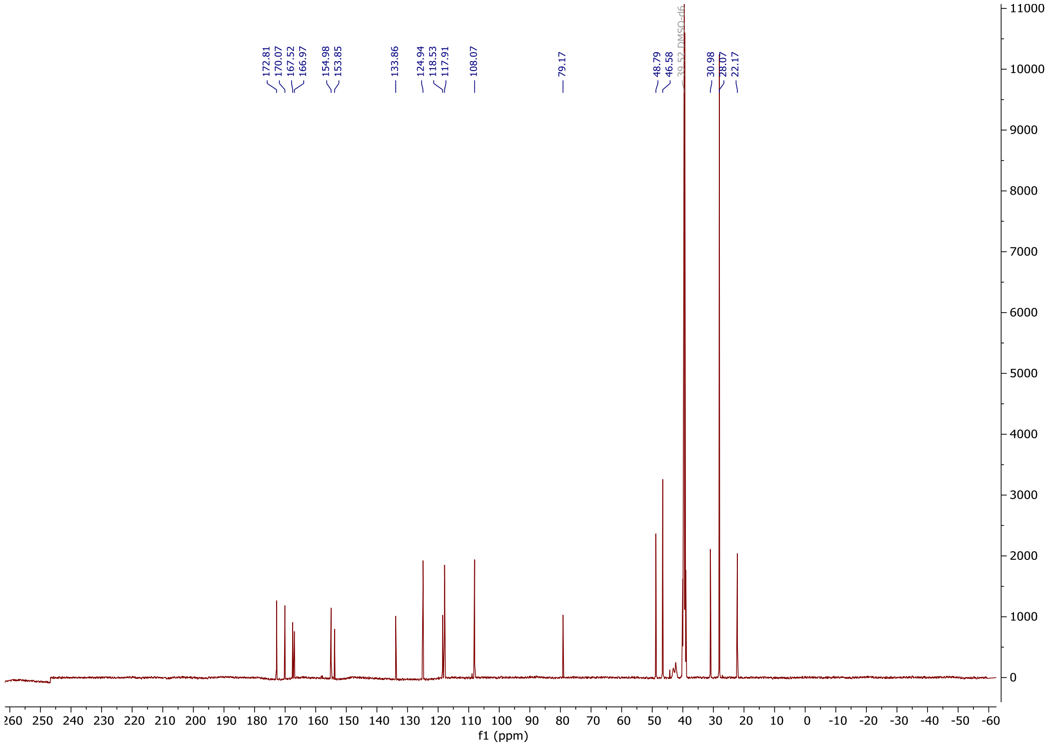
172.81
170.07
167.52
166.97
154.98
153.85
133.86
124.94
118.53
117.91
108.07
79.17
48.79
46.58
39.52 DMSO-d6
30.98
28.07
22.17
f1 (ppm)

2-(2,6-dioxopiperidin-3-yl)-5-(4-(2-(1-(2-((2-fluoro-4-((2-fluoro-3-nitrobenzyl) sulfonyl) phenyl) thio)-5-methoxy-6-((5-methyl-1H-pyrazol-3-yl) amino) pyrimidin-4-yl) piperidin-4-yl) acetyl) piperazin-1-yl) isoindoline-1,3-dione **D11**

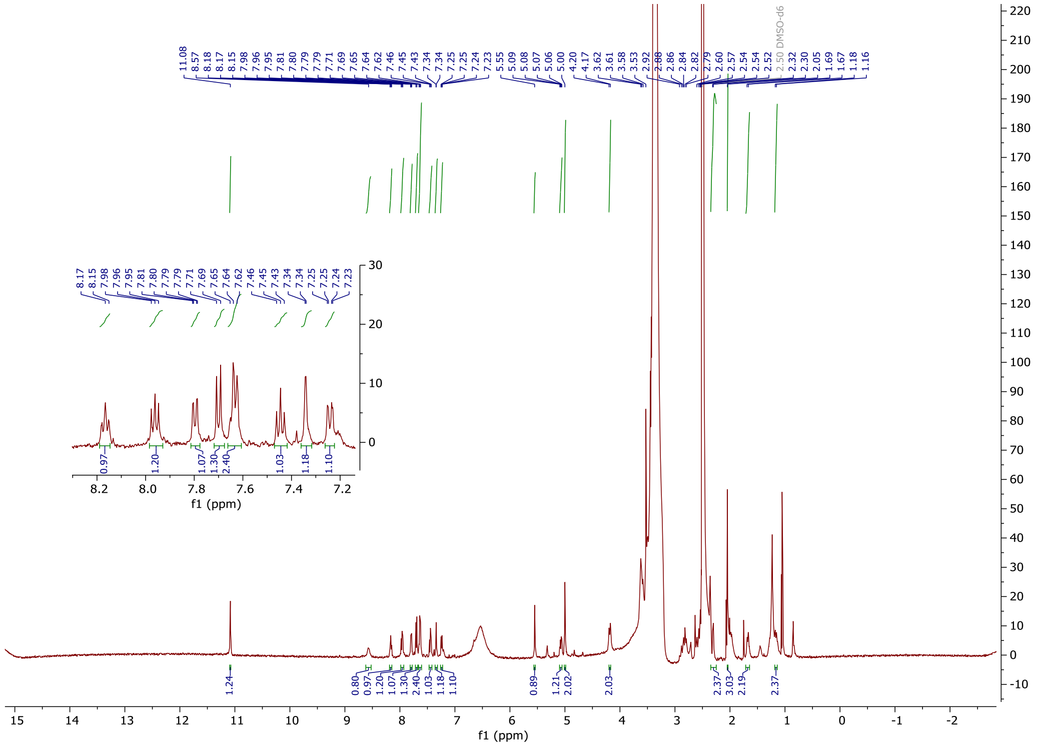


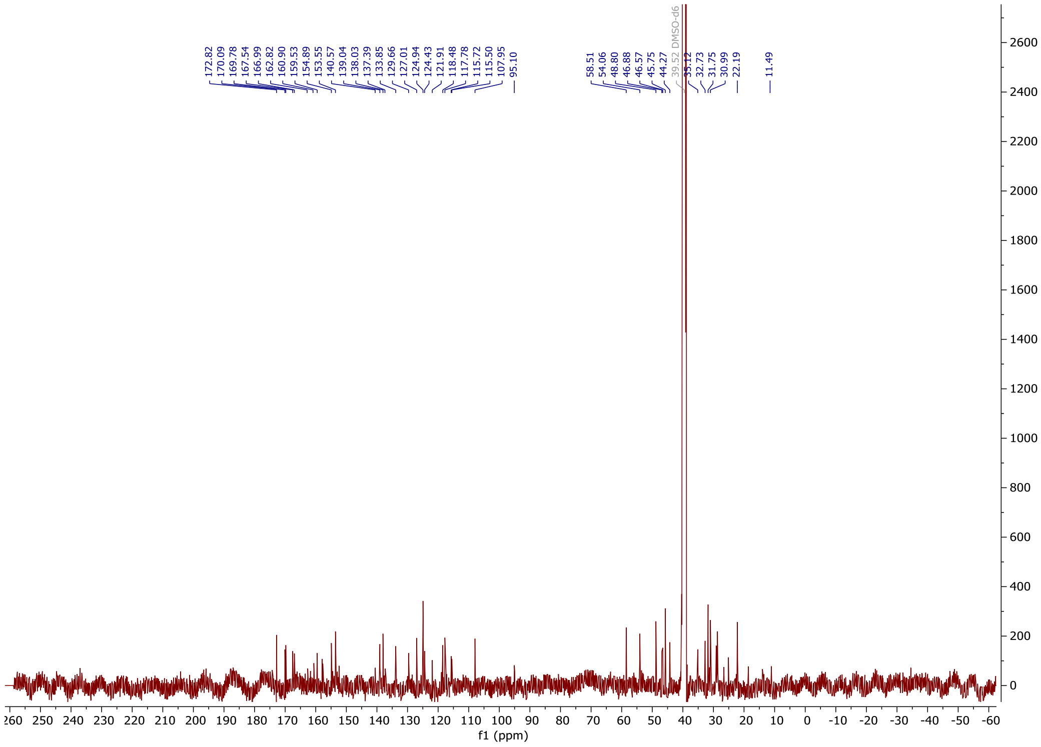

(*E*)-1-(4-(2-((2-fluoro-4-((2-fluoro-3-nitrobenzyl) sulfonyl) phenyl) thio)-5-methoxy-6-((5-methyl-1H-pyrazol-3-yl) amino) pyrimidin-4-yl) piperazin-1-yl)-4-(4-methoxyphenyl) but-2-ene-1,4-dione **D12**

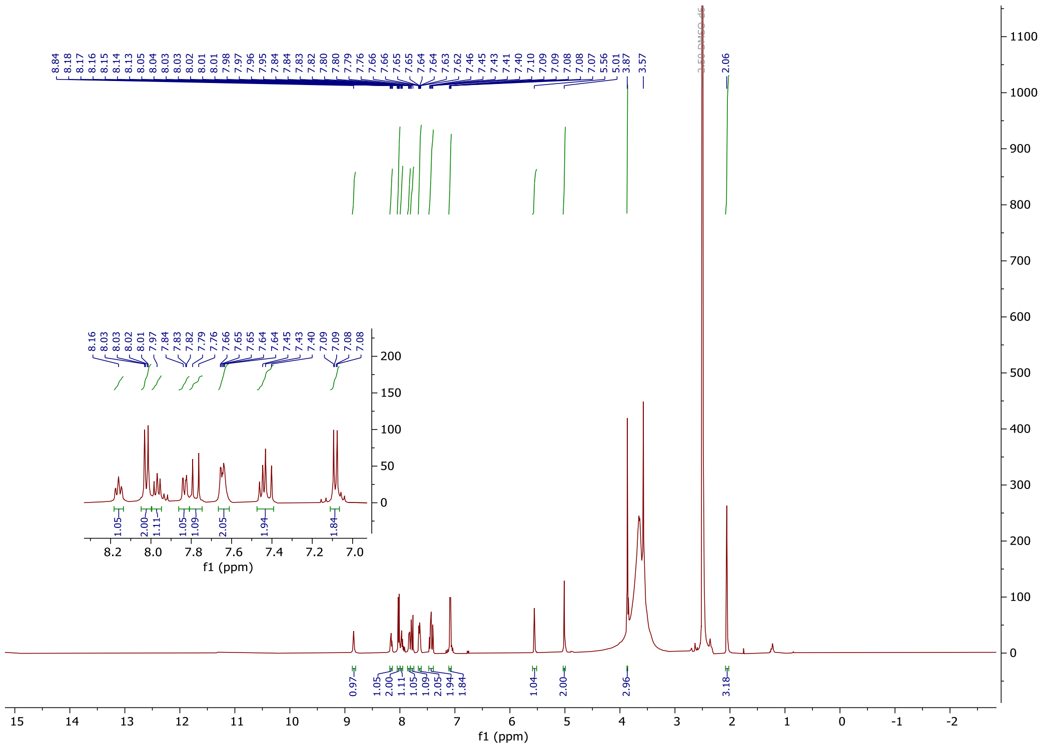

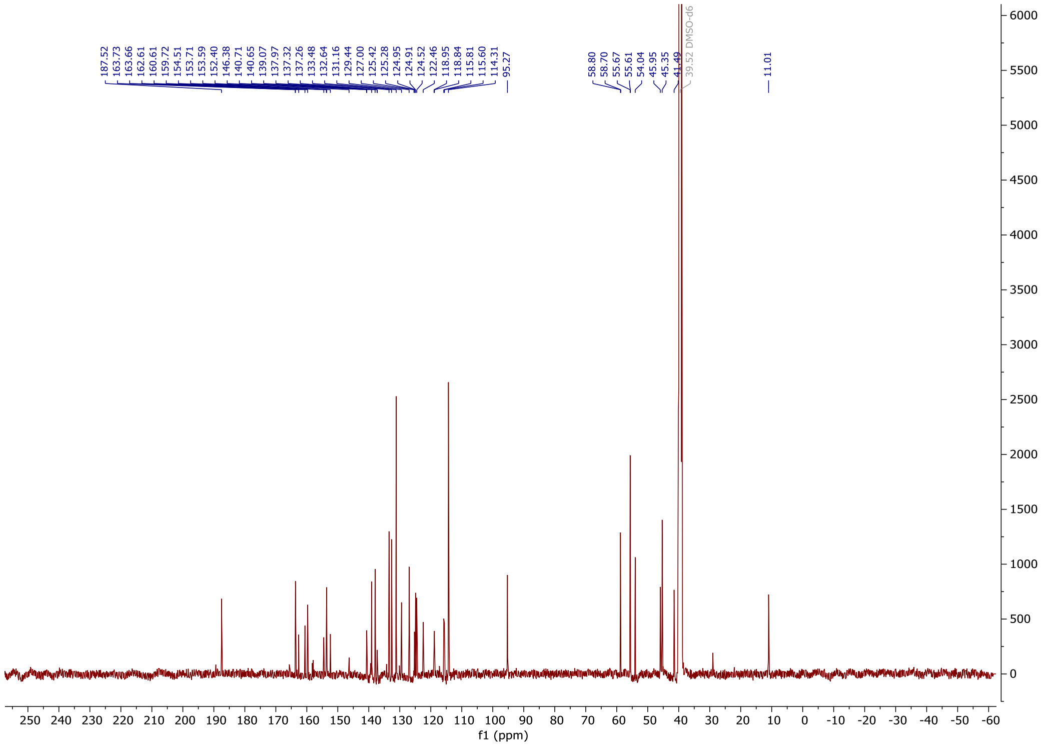

*tert*-butyl-(*E*)-4-(4-(4-methoxyphenyl)-4-oxobut-2-enoyl)-1λ4-piperazine-1-carboxylate **41**

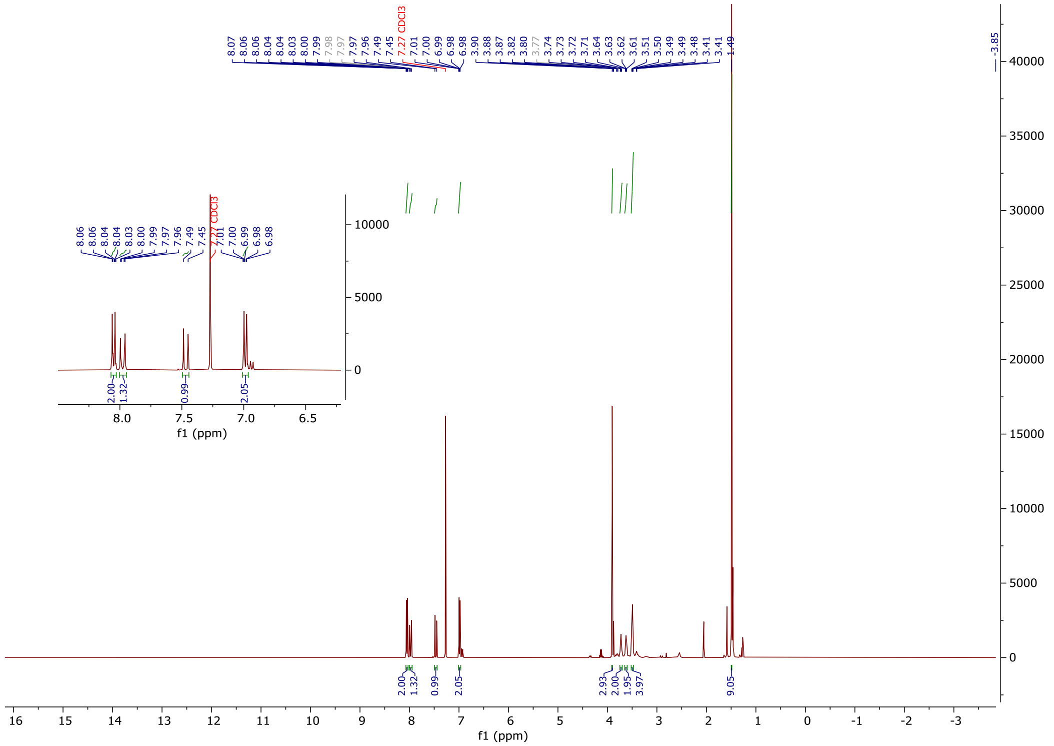

(*E*)-1-(4-methoxyphenyl)-4-(piperazin-1-yl) but-2-ene-1,4-dione **42**

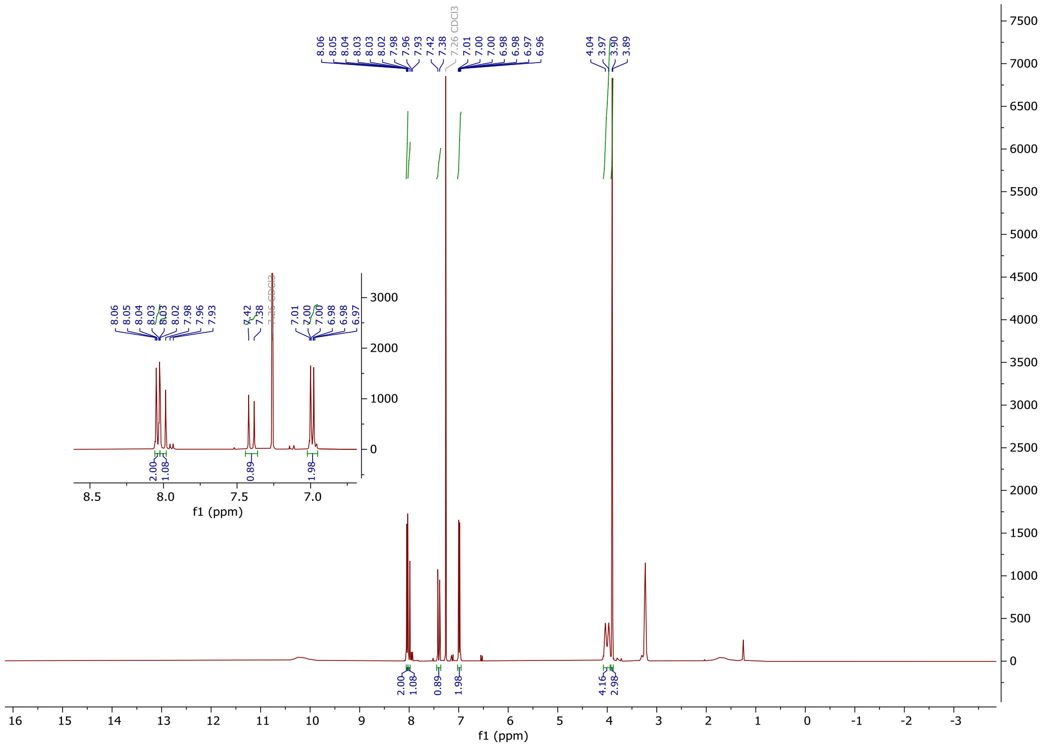


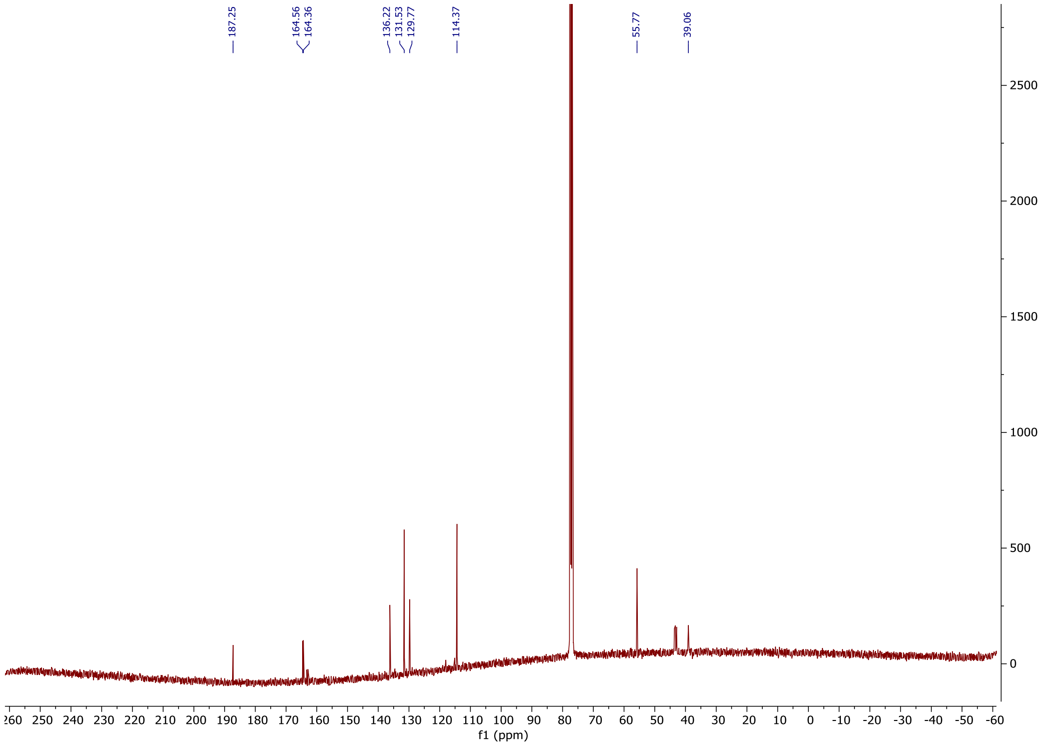

(*E*)-1-(4-(2-(1-(2-((2-fluoro-4-((2-fluoro-3-nitrobenzyl)sulfonyl)phenyl)thio)-5-methoxy-6-((5-methyl-1H-pyrazol-3-yl)amino)pyrimidin-4-yl)piperidin-4-yl)acetyl)piperazin-1-yl)-4-(4-methoxyphenyl)but-2-ene-1,4-dione **D13**

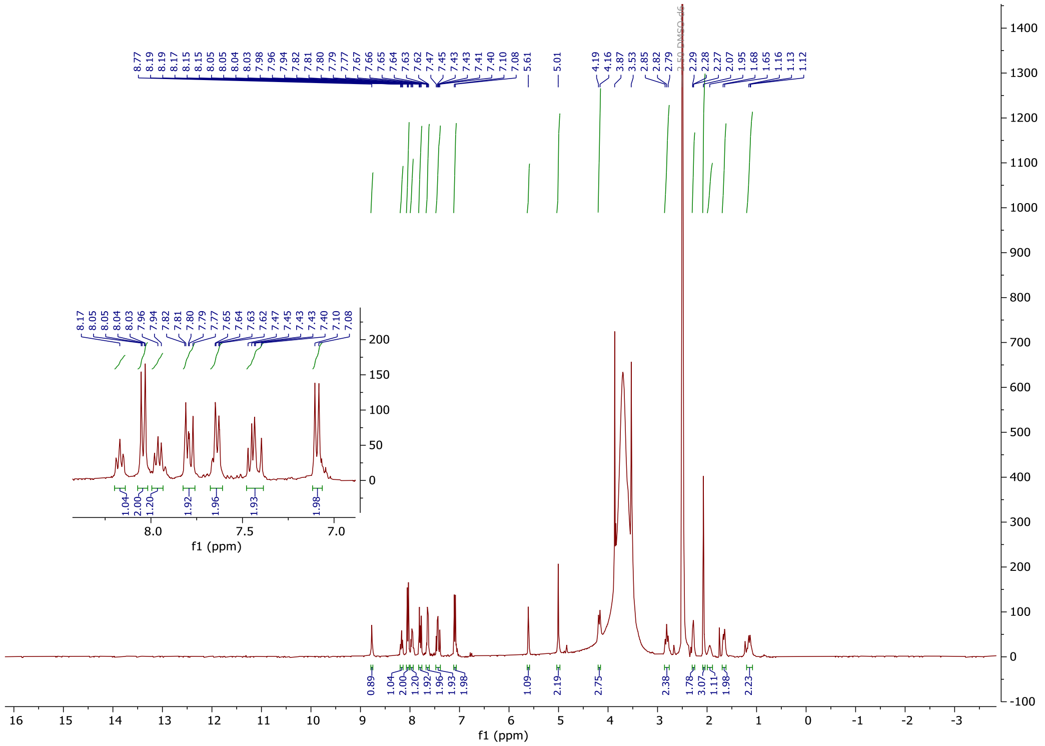

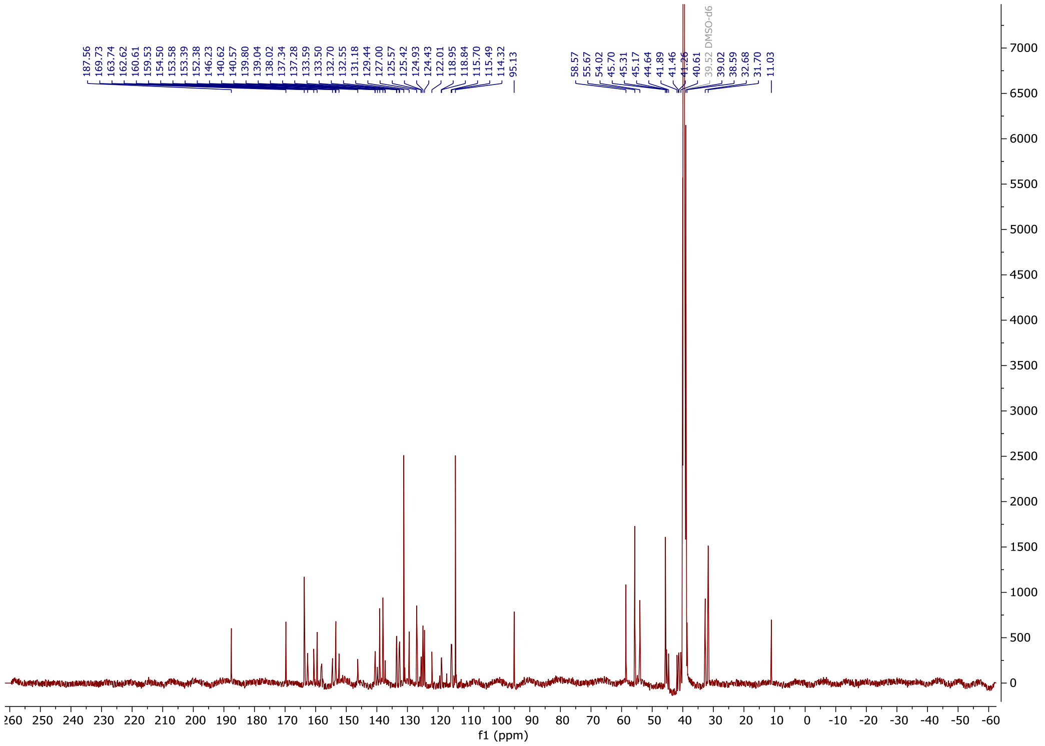

4-((2-(2-((tert-butoxycarbonyl) amino) ethoxy) ethyl) carbamoyl)-2-(6-(dimethylamino)-3-(dimethyliminio)-3H-xanthen-9-yl) benzoate **44**

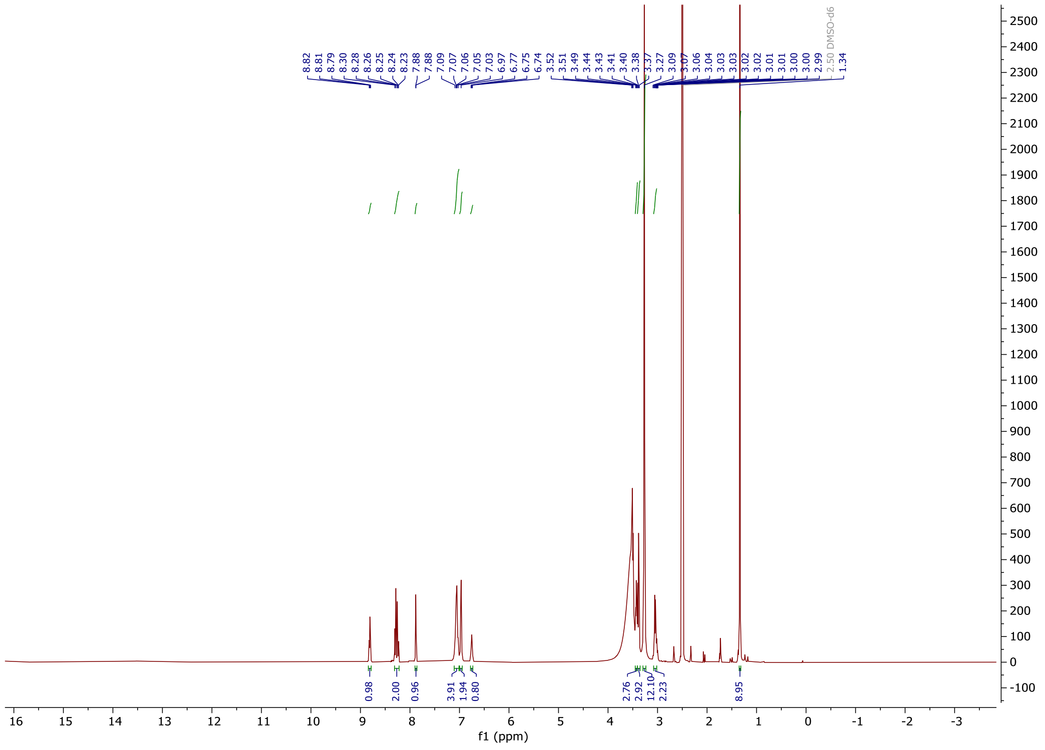

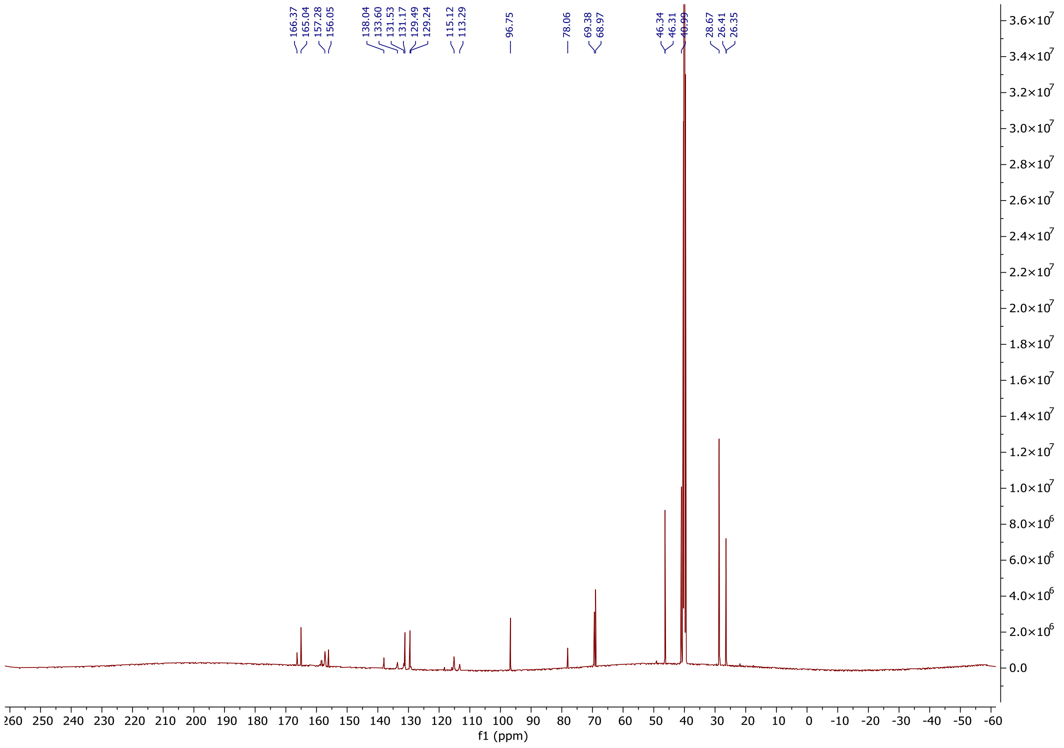

2-(3,6-bis(dimethylamino) xanthylium-9-yl)-4-((2-(2-(2-(1-(2-((2-fluoro-4-((2-fluoro-3-nitrobenzyl) sulfonyl) phenyl) thio)-5-methoxy-6-((5-methyl-1H-pyrazol-3-yl) amino) pyrimidin-4-yl) piperidin-4-yl) acetamido) ethoxy) ethyl) carbamoyl) benzoate **45**

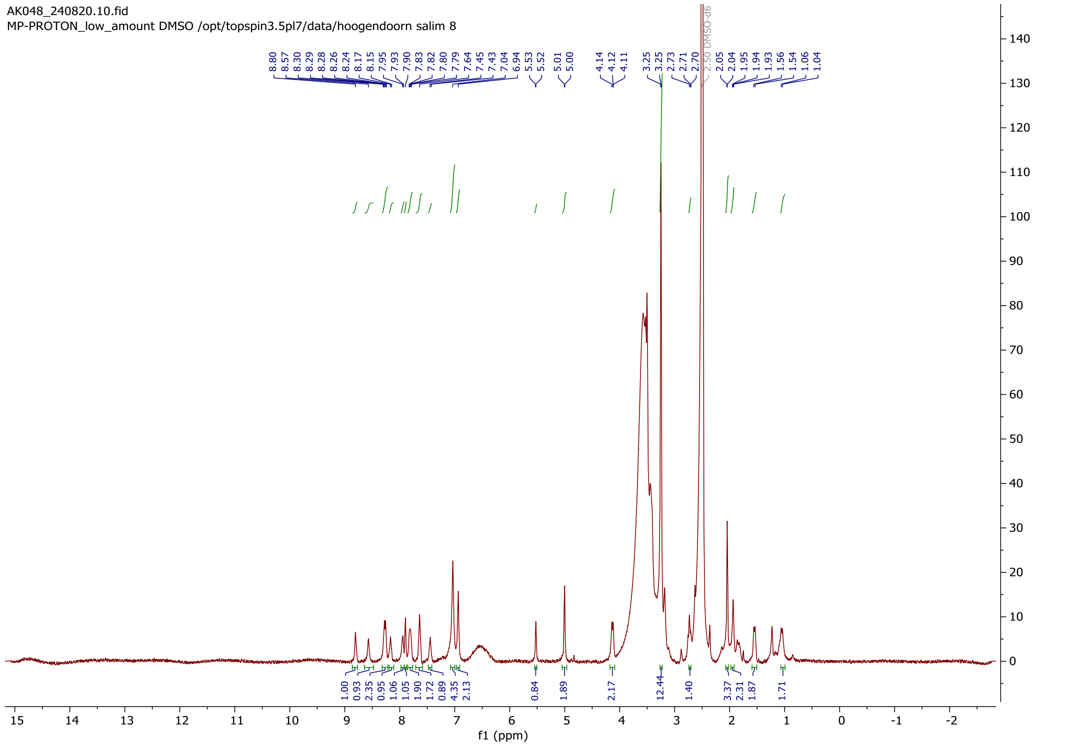


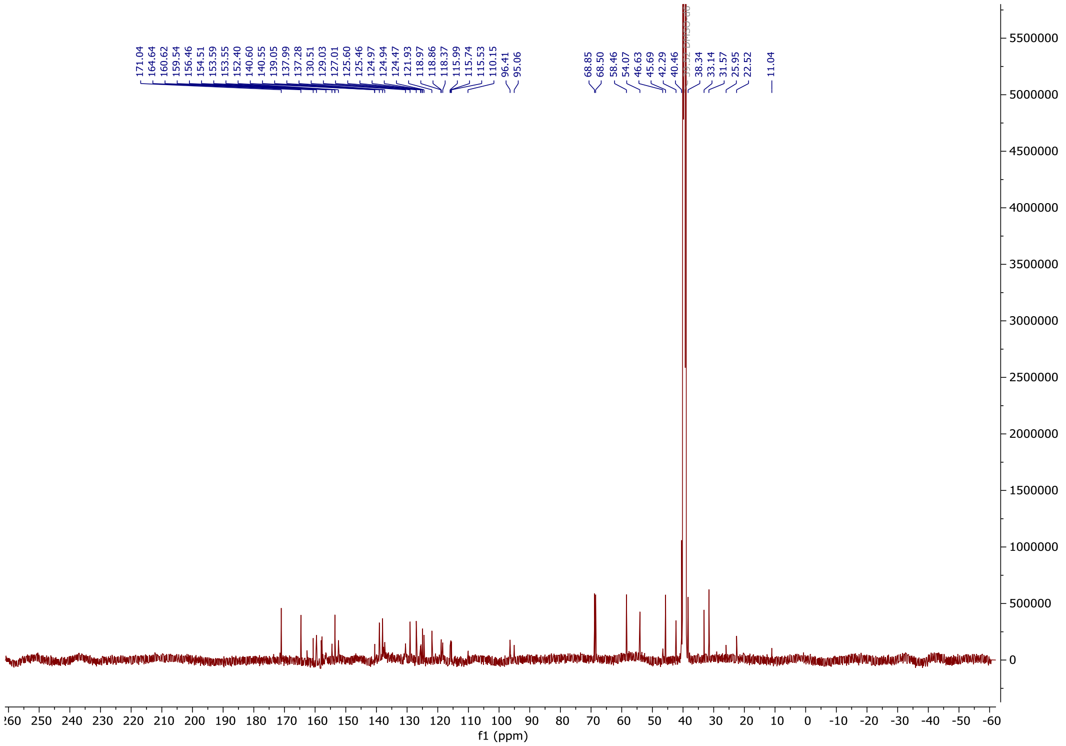
